# Nucleolar Essential Protein 1 (Nep1): Elucidation of Enzymatic Catalysis Mechanism by Combined Molecular Dynamics Simulation and Quantum Chemical Calculations

**DOI:** 10.1101/2023.03.21.532383

**Authors:** Mateusz Jedrzejewski, Barbara Bełza, Iwona Lewandowska, Marta Sadlej, Agata P. Perlinska, Rafal Augustyniak, Thomas Christian, Ya–Ming Hou, Marcin Kalek, Joanna I. Sulkowska

## Abstract

Nep1 is a protein essential for the formation of the eukaryotic and archaeal small ribosomal subunit. It is an enzyme responsible for the site–specific SAM–dependent methylation of pseudouridine (Ψ) during the pre–rRNA processing. It possesses a non–trivial topology, namely, a 3_1_ knot in the active site. Herein, we investigate the structure and mechanism of catalysis of Nep1 using a combination of bioinformatics, computational, and experimental methods. In particular, we address the issue of seemingly unfeasible deprotonation of Ψ nucleobase in the active site of Nep1 by a distant aspartate residue (e.g., D101 in Nep1 of *S. cerevisiae*). Sequence alignment analysis across different organisms identifies a conserved serine/threonine residue that may play a role of a proton–transfer mediator (e.g., S233 in Nep1 from *S. cerevisiae*), facilitating the reaction. Two enzyme–substrate complexes, one based on an available crystal structure and the other generated by molecular docking, of representative eukaryotic (from *S. cerevisiae*) and archaeal (from *A. fulgidus*) Nep1 homologs are subjected to molecular dynamics (MD) simulations. The resulting trajectories confirm that the hydroxyl–containing amino acid can indeed adopt a position suitable for proton–shuttling, with the OH group located in between the proton donor and acceptor. However, during the MD simulations, a water molecule emerges from arrangements of the active site, which can assume the role of the proton–transfer mediator instead. To discern between these two alternative pathways, we evaluate the possible methylation mechanisms by quantum–chemical calculations based on density functional theory, using the cluster approach. The obtained energy profiles indicate that the most facile course of the reaction for both the yeast and archaeal enzymes is to engage the water molecule. These results are corroborated by agreement of the computed energy barriers with experimentally measured enzyme kinetics. Moreover, mutational studies show that, while aspartate D101 is crucial for the catalytic activity, serine S233 is irrelevant in this context, indirectly supporting the water–mediated proton transfer. Our findings comprehensively elucidate the mode of action of Nep1 and provide implication for understanding the catalytic mechanisms of other enzymes that involve a proton transfer in the active site over extended distances.

## Introduction

Methylation is one of the most common post–synthetic modifications of DNA and RNA. It plays an important role in the regulation of crucial processes, such as gene expression, cell cycle, tumor progression, or genome repair. A large group of enzymes is dedicated to synthesis of methylated nucleic acids, each typically showing specificity for a given target site. Accordingly, they represent multiple distinct folds and use different binding modes to their substrates.^1,2^ Conversely, the great majority of methyltransferases utilize a common methyl donor, S–adenosyl–methionine (SAM). Methyltransferases are divided into five distinct superfamilies, among which the second largest is the SPOUT (SpoU–TrmD).^3,4^ SPOUT proteins modify mainly RNA, especially rRNA and tRNA.^5^ Most of the SPOUT proteins are homodimers and in all SPOUT members, the protein backbone forms a non–trivial topology, a so–called trefoil knot (Figure 1).^6^

**Figure 1:**
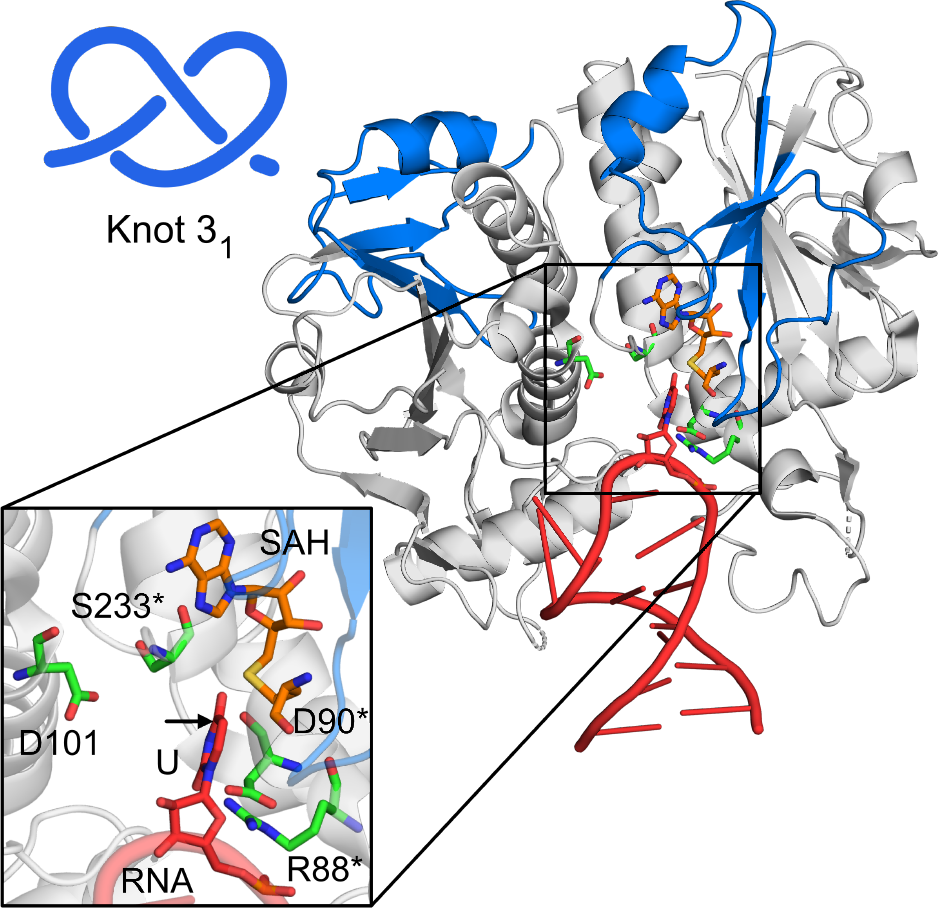
Crystal structure of *S. cerevisiae* Nep1 (grey) in complex with 18S rRNA (red); PDB code: 3oin. The 3_1_ knot is colored blue. The inset shows the details of the active site with the key residues indicated (in the crystal SAM was replaced by inhibitor, Sadenosylhomocysteine (SAH), and Ψ1191 was replaced by U). The C5 of U, corresponding to the N1 of Ψ (the position of methylation) is indicated with an arrow.

In SPOUT proteins, the role of the knot is generally not well known. However, given that the knot is strictly structurally conserved and necessary for function, it is likely a vital part of their active site.^7,8^ In particular, the knot residues are involved in the binding of both the nucleic acid substrate and the methyl donor, as well as in the interactions between the two subunits of the homodimeric protein. Moreover, a mutation in the knot can break the intricate signaling within the protein and decrease or even completely shut down the enzymatic activity.^9^ Given that the knot is a crucial region of a SPOUT protein, its potential role has been discussed in several studies.^5,10,11^

In this work, we focus on a knotted SPOUT methyltransferase, the Nucleolar Essential Protein 1 (Nep1, also known as EMG1). Although the majority of SPOUT family members are bacterial proteins, Nep1 is only present in Eukarya and Archaea.^12^ It exhibits a typical SPOUT quaternary and tertiary structure, including being a homodimer and having a 3_1_ knot in the active site (Figure 1).^13^ Nep1 plays an important role in the ribosomal biogenesis and is necessary for the formation of the small ribosomal subunit.^14^ Specifically, Nep1 is responsible for the site–specific N1–methylation of pseudouridine (Ψ) in archaeal 16S rRNA and eukaryotic 18S rRNA (Ψ1191; all residues numbers in the text are given for *Saccharomyces cerevisiae*, unless specified otherwise).^12,15^ It recognizes a highly conserved 5’–(C/U)ΨCAAC–3’ consensus motif.^12^ While this sequence appears three times in the eukaryotic and twice in the archaeal rRNA, the motif located in helix 35 was experimentally confirmed as the exclusive Nep1 methylation target. ^12^ Importantly, the D86G mutation in the human Nep1 (equivalent to D90 in *S. cerevisiae*) was identified as the cause of Bowen–Conradi syndrome – a severe ribosomopathic disease, in most cases lethal in early childhood.^16,17^

As for all SPOUT proteins, the methylation catalyzed by Nep1 is assumed to occur in two chemical steps: (1) a deprotonation of the nitrogen atom in the nucleobase (the N1 of Ψ) and (2) a methyl transfer from SAM.^12^ The binding of SAM and rRNA to Nep1 and the methyl transfer mechanism have been explored through structural and sequential comparisons with other SPOUT family members, as well as by in vitro enzymatic activity studies. Four highly conserved arginine residues: R88, R129, R132, and R136, were found to be engaged in rRNA binding. Alteration of any of these residues results in a loss of the enzymatic function of Nep1 in vitro. ^5,12,13,18^ Among them, R88 was proposed to form hydrogen bonds to the O4 oxygen of Ψ.^19^ This interaction would not only be responsible for substrate recognition and binding, but it would also promote the enzymatic reaction by facilitating the deprotonation of N1 and stabilizing the negative charge in the intermediate (Figure 2).^19^ Mutations D214R, L232S, and A237D were found to weaken SAM binding.^13^ Importantly, based on partial conservation analysis and structural data, aspartate D101 was suggested to be the most probable proton acceptor during the deprotonation step. However, in the only available crystal structure of Nep1 complex with rRNA (from *S. cerevisiae*, PDB code: 3oin; Figure 1), the residue is located as much as 6.8 Å away from the supposed position of the N1 atom of Ψ (corresponding to the C5 of U in the crystal structure).^18^ An additional complication arises from a considerable variability of the amino–acid sequences within the active sites of Nep1 homologs originating from different organisms, which may have implications for the methylation mechanism and even lead to divergent pathways. Overall, the current understanding of the catalysis by Nep1 is only superficial and its fundamental details remain unclear.

**Figure 2:**
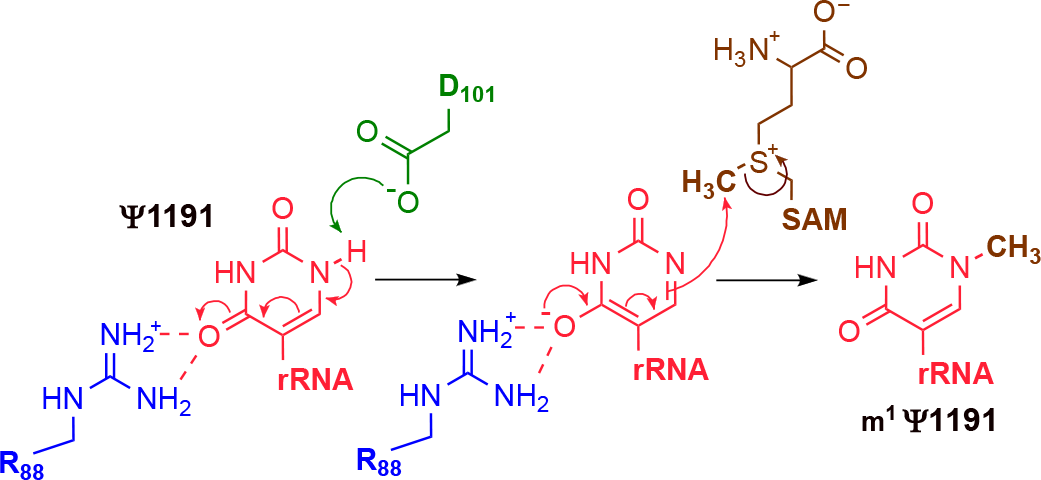
Overview of the proposed mechanism for the N1–methylation of pseudouridine catalyzed by Nep1.

Due to the biological importance of Nep1, herein, we use an array of computational meth-ods, corroborated by experimental validation, to study its structure and to elucidate the mechanism of enzymatic catalysis. In particular, through a sequence alignment analysis of Nep1 homologs from multiple organisms, we identify potential proton transfer mediators that could allow for the deprotonation of pseudouridine by the distant base in all eukaryotic and some archaeal homologs of Nep1. To gain insight into the mechanism of methyl transfer, two representative proteins, from *S. cerevisiae* and *A. fulgidus*, are subjected to a hybrid workflow that includes: molecular docking of the rRNA fragment (only for *A. fulgidus* Nep1), molecular dynamics (MD) simulations of the full enzyme–substrate complex, and quantum chemical (QM) modelling of the methyl transfer reaction in the active site. A similar sequential MD/QM approach has enabled us recently to uncover the mechanism of action for another SPOUT methyltransferase, an Mg^2+^–dependent TrmD.^20^ This methodology is particularly suited for situations when available crystal structures are incomplete, such as in the case of TrmD, wherein the position of the catalytic Mg^2+^ ion is unknown.^21^ For the *A. fulgidus* Nep1 homolog, we take it even a step further, assembling first the enzyme–substrate complex by the computational docking of an rRNA strand. This comprehensive in silico characterization of both the structure and dynamics of the enzyme–substrate complexes of Nep1 proteins and their catalytic mechanisms establishes that the proton–transfer step indeed requires a mediator, whose role may be assumed either by a suitable amino acid residue or, preferably, by a water molecule present in the active site. Thus, the current results not only provide new and significant insights into understanding the important Nep1 enzyme, but they also advance the protein modelling methodology and showcase its potential.

## Results and discussion

### Identification and diversity of potential catalytic residues in the active site of Nep1 from different organisms

First, we performed a bioinformatics analysis to explore the conservation of amino acids in the active site of Nep1 across organisms. We conducted a multiple sequence local alignment using the Clustal Omega algorithm^22^ for selected species (Figure 3). The sequence data was mapped to the crystal structure of the active site of Nep1 from *S. cerevisiae* containing the rRNA substrate and the SAH analog (Figure 1) to identify the residues that may be relevant for promoting the methyl transfer reaction. To this end, aspartate D90, mutated in the Bowen–Conradi syndrome, remains present in all the examined sequences, along with the nearby arginine R88. Although it may be tempting to assume that D90 could be the acceptor during the proton–transfer step, it is positioned in the active site on the opposite side of the pseudouracil ring from the supposed position of the N1 proton, precluding its involvement as the general base in the mechanism. Based on the crystal structure, D90 may work together with R88 to create a binding pocket for the Ψ nucleobase. In contrast, the supposed N1 of Ψ is directed toward aspartate D101, making the latter a viable candidate as the proton acceptor, as previously suggested.^18^ Due to the considerable distance separating these two moieties, we focused on the region of the active site in between them. Interestingly, there is a serine moiety (S223) located midway, whose OH group could possibly act as a proton shuttle, enabling deprotonation over the extended distance. Moreover, aspartate D101, proposed as the proton acceptor, and serine S233, proposed here as the proton–transfer mediator, remain conserved throughout the examined eukaryotic species (Figure 3), supporting their involvement in the enzymatic mechanism.

**Figure 3:**
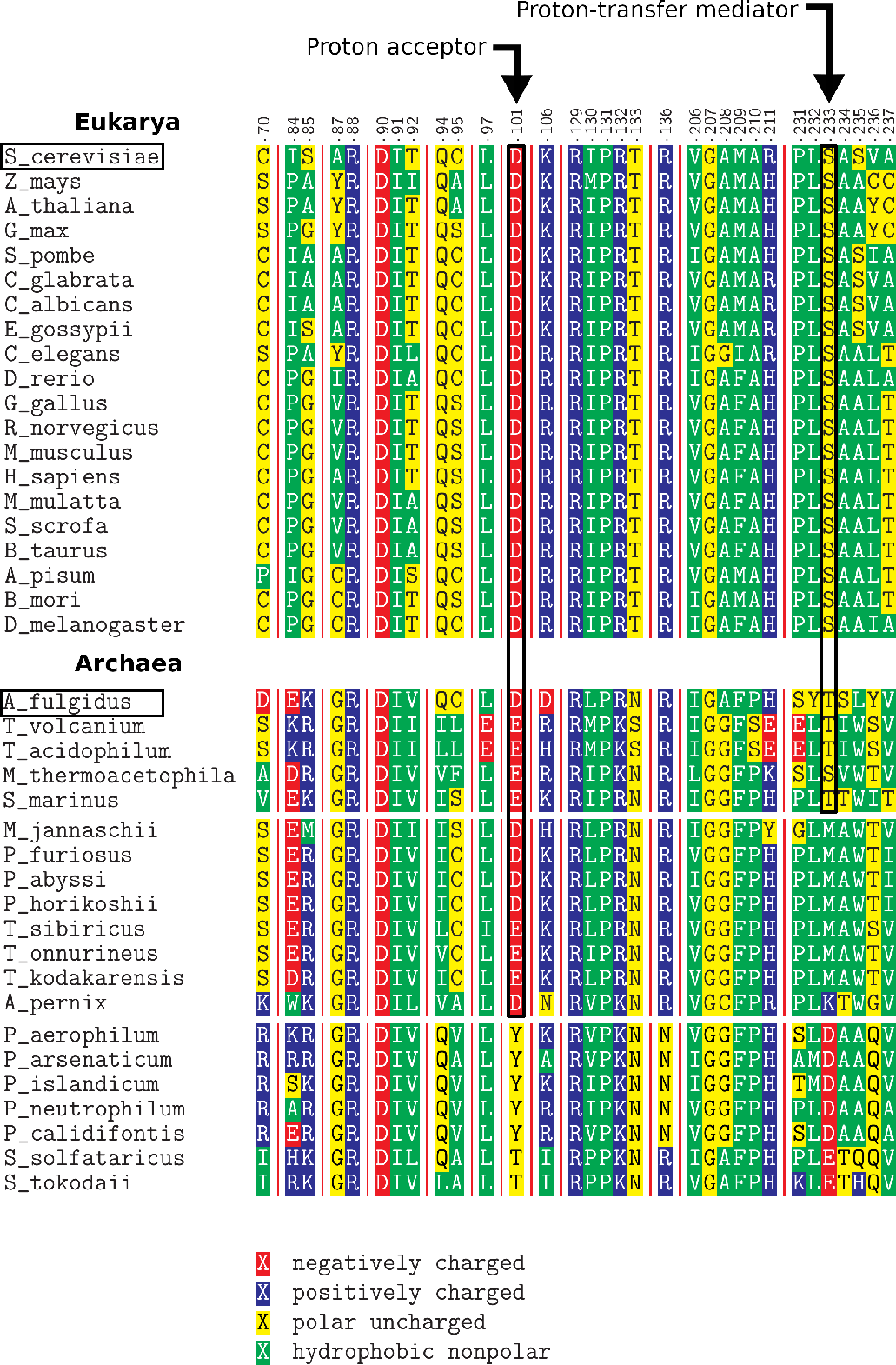
Fragment of the alignment of Nep1 proteins from Eukarya and Archaea. The numbering is based on Nep1 from *S. cerevisiae*. Arrows point to the residues we propose as catalytically important (the proton acceptor and the proton–transfer mediator). The sequences in Archaea are grouped based on the identity of amino acids in these two positions. Black boxes indicate *S. cerevisiae* and *A. fulgidus*, whose Nep1 proteins were further investigated with MD and QM.

Despite the overall high conservation of the protein sequence, contrary to eukaryotes, the archaeal species exhibit a considerable diversity among the amino acids occupying these two important sites. Specifically, we can distinguish three classes within the Archaea domain. The first class closely resembles the eukaryotic sequence with either aspartate or glutamate as the putative proton acceptor and either serine or threonine as the putative proton transfer mediator. In the second class, a methionine is present (or a lysine, in the single case of *A. pernix*) in place of serine or threonine as the proton transfer mediator. The third class differs even more – the proposed proton acceptor site is occupied by a threonine or tyrosine, aspartate or glutamate occupies the putative mediator site (Figure 3). Here, a proton–transfer mediator is deemed unnecessary, as the carboxylate group of these residues is close to the N1 of Ψ and could likely directly contribute to the deprotonation.

The variability of the proposed catalytic residues implies that the hypothesized hydroxyl–mediated deprotonation cannot be a universal pathway for all the archaeal Nep1 homologs. Nevertheless, without considering two latter classes, we decided to investigate in detail the mechanism of the methylation for two representative enzymes: Nep1 of *S. cerevisiae* (eukaryotic, containing a serine as the supposed proton–transfer mediator) and Nep1 of *A. fulgidus* (archaeal, containing a threonine, instead). The respective proteins will be referred to as *Sc*Nep1 and *Af* Nep1 in the following text.

### Docking of rRNA to *Af* Nep1

As mentioned above, the only available crystal structure of the Nep1 complex with rRNA is the one for *Sc*Nep1 (PDB code: 3oin). Therefore, in the case of *Af* Nep1, we first carried out molecular docking between the protein and the RNA substrate using the NPDock server.^23^

A survey of the available crystal structures of SPOUT methyltransferases reveals that the protein surface in contact with RNA is very much alike across the family, suggesting a similar binding mode to the nucleic acid substrate. In the specific case of *Sc*Nep1, four arginine residues engaged in rRNA binding (R88, R129, R132, and R136) are highly conserved throughout almost all organism that contain this enzyme.^12,13,18^ Moreover, the C_*α*_ superposition of the *Af* Nep1 and the *Sc*Nep1 crystal structures reveals small root–mean–square deviation (RMSD) values, confirming structural similarity of the two proteins (Table S1). Finally, the target rRNA of Nep1 is relatively similar in both Eukarya and Archaea. In particular, the secondary structure of 18S (*S. cerevisiae*) and 16S (*A. fulgidus*) rRNA exhibits high resemblance within the loop that contains the Ψ for methylation (Figure S1).^24^ Therefore, considering the high degree of homology between the eukaryotic and archaeal proteins, as well as between the respective rRNA substrates, we decided to dock the nucleic acid fragment cropped from the crystal structure of the enzyme–substrate complex with *Sc*Nep1 (PDB code: 3oin) onto the holoenzyme *Af* Nep1 (PDB code: 3o7b).

Five clusters of structures were generated by the server, in two of which the rRNA was docked at the correct site. The homology between yeast and archaeal proteins was used for the evaluation and selection of obtained docked complexes. First, we superimposed the output structures from these two clusters with the crystal structure of *Sc*Nep1 (by the C_*α*_ atoms) and rejected the complexes in which the RMSD calculated for the phosphorus atoms of rRNA was over 4 Å. At this point, eight structures from a single cluster remained. Next, we examined the surroundings of the four conserved arginine residues that are crucial for the rRNA binding^19^ and selected four complexes with the distances to the rRNA suitable for forming hydrogen bonds (the full list of examined hydrogen bonds can be found in Table S2). Finally, we discarded any structures containing heavy atom clashes, which could disturb the stability of the complex during the subsequent step of the workflow – the full–atom MD simulations. From the remaining two structures, a single one was selected and used as a starting point for the MD simulations.

Following the docking, the uracil moiety present in the crystal structure was manually replaced with pseudouracil in the appropriate position. Analogously, SAH from the crystal of *Af* Nep1 was modified to the actual methyl donor SAM.

### Structure and dynamics of the active site by MD simulations

We conducted multiple all–atom molecular dynamics simulations for each of the enzyme–substrate complexes in order to investigate the dynamics of each structure, especially within the active site region. Unlike crystal structural analysis, MD simulations allow for probing and analysis of the conformational landscape of a protein, giving insight into the conformational state that most likely represents the one in physiological condition and providing implications for catalysis. This issue becomes even more vital for the enzyme–substrate complex of *Af* Nep1, which was obtained through molecular docking. As its structure was generated by fitting together two rigid fragments taken from separate crystals, it is clearly far from optimal at this stage. Therefore, the MD simulations will not only explore the conformational details of the *Af* Nep1–RNA complex, but will also enable the complex to adopt a more relaxed structure.

Overall 1.2*µ*s and 2.3*µ*s of independent trajectories were run for *Sc*Nep1 and *Af* Nep1, respectively (Table S4). In the former case, the starting structure for MD simulations was the known crystal structure (PDB code: 3oin, with the uracil moiety manually replaced with pseudouracil in the appropriate position and SAH changed to SAM). In the latter, the enzyme–substrate complex obtained by the docking was used.

The RMSD for C_*α*_ atoms in all the simulations was below 4 Å and the enzyme–substrate complexes were stable throughout the simulation periods (Figure S2). Hydrogen bonds between the rRNA and protein chains A and B were generally preserved throughout the trajectories. Moreover, the number of these bonds was similar for both studied organisms (Figures S3–4). Furthermore, we performed a quantitative occurrence analysis for the most important protein–RNA hydrogen bonds, confirming that the interactions with the four conserved RNA–binding arginines are preserved in the MD simulations (Table S5). The stability testing in multiple trajectories for *Af* Nep1 also provides a solid validation of successful docking.

Regarding the active sites themselves, the distances between the key moieties, such as Ψ and SAM, Ψ and S233, and Ψ and D101 (Figures S6–9), remained stable throughout the MD simulations. Nevertheless, the active sites experienced slight fluctuations, resulting in a range of conformations. Importantly, these included the arrangements that aligned the proposed proton–transfer mediators (the OH groups of S233* and T198* for *Sc*Nep1 and *Af* Nep1, respectively) in between the N1 of Ψ and the carboxylic group of D101 (D74 in *Af* Nep1). This alignment this connects them with two contiguous hydrogen bonds (Figure 4A–B). It is worth noting that in the crystal structure of *Sc*Nep1, although S233* is positioned in between the nucleobase and D101, its OH group does not directly interact with either of these moieties (Figure 1). This may be in part caused by the presence of a regular uridine in the RNA in the crystal, instead of the actual pseudouridine substrate. Thus, the MD simulations identified a conformational state of the active site that is putatively better suited for catalysis, which is indeed confirmed by the following QM calculations (see below).

**Figure 4:**
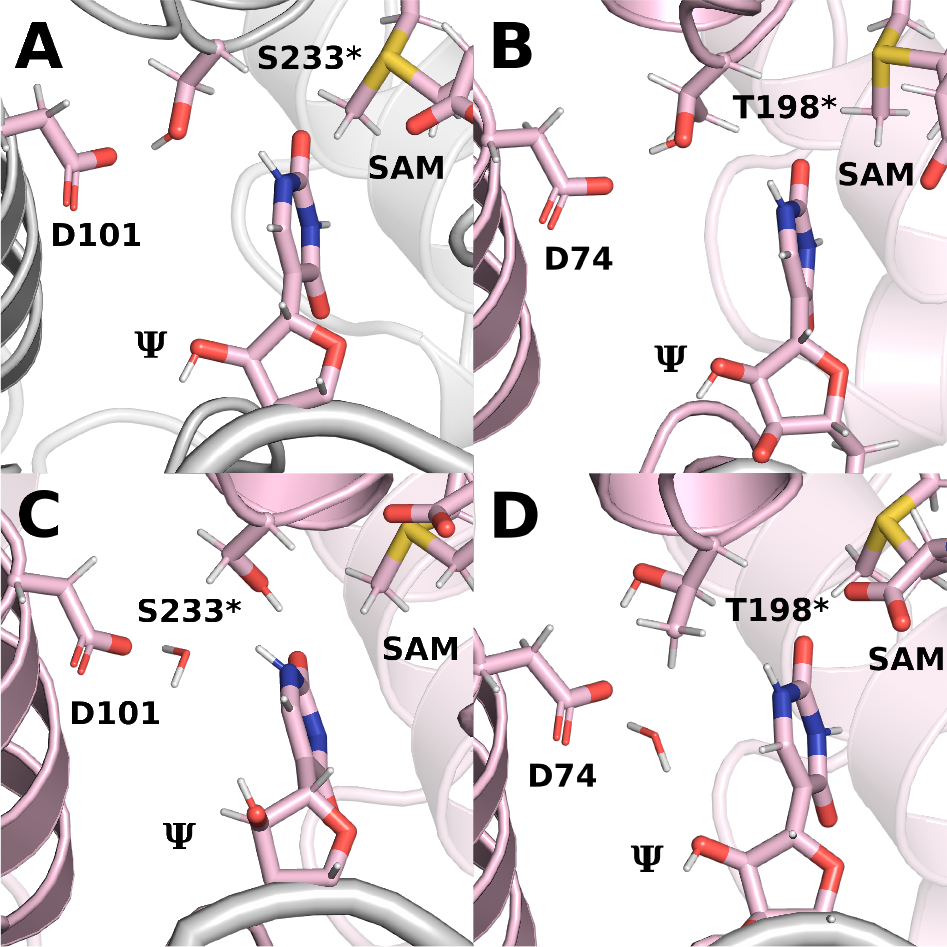
The snapshots of each active site from the MD simulations. In A and B, the N1 of Ψ and the proton acceptor are bridged by the OH group of the putative proton–transfer mediator residue (S233* and T198* for *Sc*Nep1 and *Af* Nep1, respectively). In C and D, these moieties are bridged by a water molecule instead, indicating a water–mediated proton transfer as an alternative mechanistic pathway

Very interestingly, in addition to the conformation that bridges the N1 of Ψ and the proton acceptor by the OH group of serine or threonine, we also identified, for both *Sc*Nep1 and *Af* Nep1, the MD snapshots wherein the proton donor and acceptor are each hydrogen–bonded by a water molecule instead (Figure 4C–D). Although the abundance of such geometries was rather low (below 1% of the simulation time), their spontaneous and repeated occurrence during the nanosecond simulations makes the water–mediated proton transfer a valid possibility during the methyl transfer mechanism (the chemical reaction occurs within the order of seconds).

In summary, the MD simulations have confirmed that the active site of *Sc*Nep1 and *Af* Nep1 can adopt a conformation that bridges the OH group of the putative proton–transfer mediator residue (S233* and T198*, respectively) with both the N1 of Ψ and the carboxylic group of the proton acceptor (D101 and D74, respectively) with hydrogen bonds. Such an arrangement renders the active site primed for the first step of the methylation reaction, i.e. deprotonation of N1, despite the large distance separating it from the proton acceptor. Furthermore, the simulations have revealed conformations of the active site that can bridge a water molecule with the proton donor and the proton acceptor, indicating hat a water–mediated proton transfer may be possible.

### Comparison of the mechanistic pathways by QM calculations

To characterize the methylation mechanism and evaluate the two alternative reaction pathways, we conducted QM calculations based on the density functional theory using the cluster approach.^25–28^ In this method, an enzyme–substrate complex is truncated to leave only the active site region. The steric effects of the remainder of the structure are modelled by fixing the positions of selected atoms (typically at the truncation points), while the electrostatic effects are represented by the implicit continuous solvent. The cluster approach has been successfully applied to study several methyl transfer reactions in SAM–dependent enzymes, employing both crystallographic and MD data to construct the active site model.^20,29–31^

The computations were carried out at B3LYP–D3BJ(CPCM)/Def2–QZVP//B3LYP–D3BJ/Def2–SVP level of theory. Energy profiles for the methylation of Ψ for the following cases were calculated: Model 1: *Sc*Nep1 with S233* as the proton–transfer mediator, the arrangement of the active site based on the crystal structure; Model 2: *Sc*Nep1 with S233* as the proton–transfer mediator, the arrangement of the active site based on the MD simulations; Model 3: *Sc*Nep1 with a water molecule as the proton–transfer mediator (MD arrangement); Model 4: *Af* Nep1 with T198* as the proton–transfer mediator (MD arrangement); Model 5: *Af* Nep1 with a water molecule as the proton–transfer mediator (MD arrangement). A general two–step mechanism was assumed, as depicted in Figures 5A–B, for the pathways engaging serine/threonine and water in the proton transfer step, respectively.

**Figure 5:**
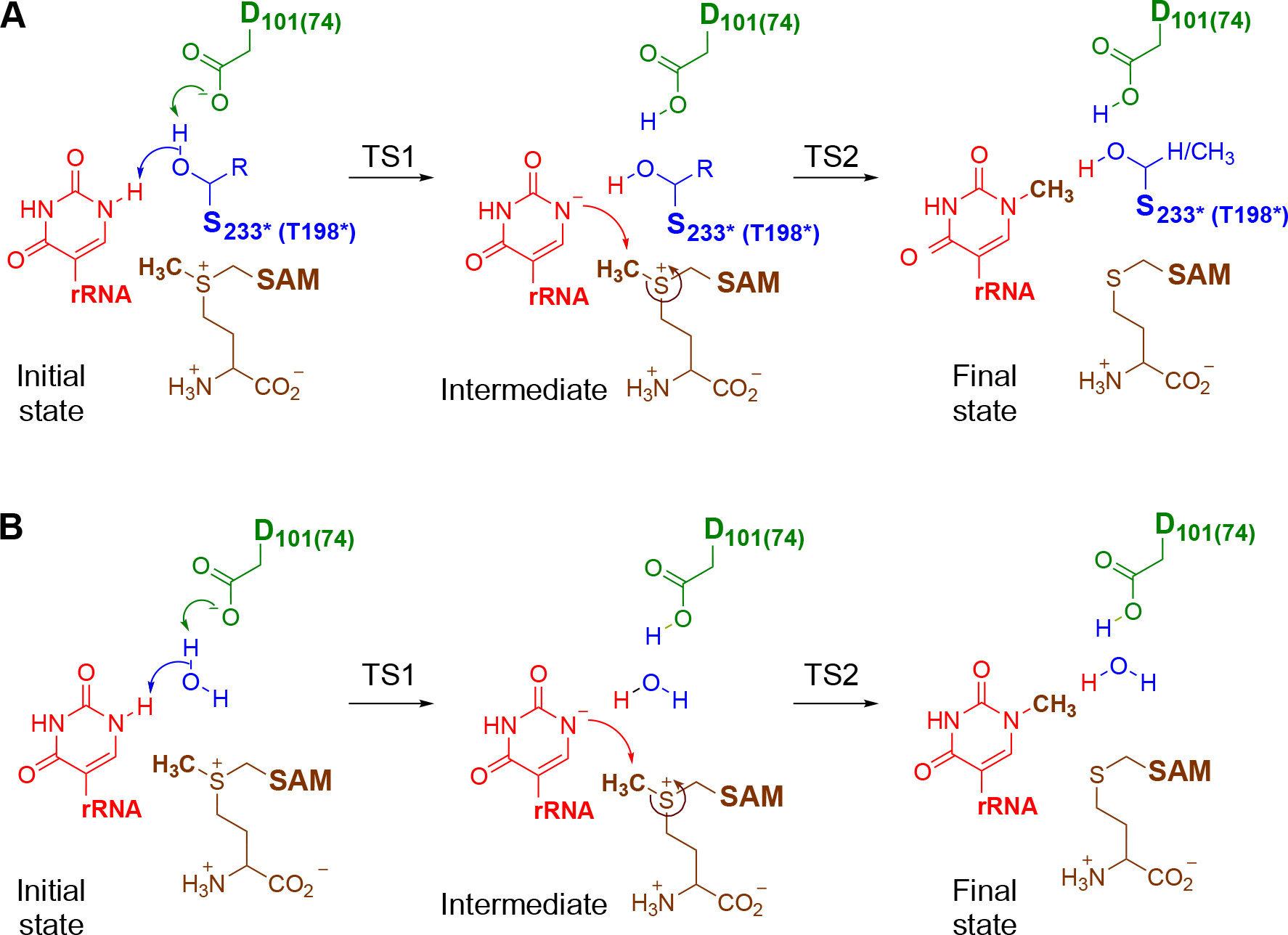
Overview of the evaluated mechanisms of the methylation of Ψ by Nep1, consisting of two chemical steps: the proton transfer (TS1) from N1 to aspartate (D101/D74), mediated by either (A) the hydroxyl–containing amino acids (S233*/T198*) or (B) the molecule of water, and the methyl transfer (TS2) from SAM to N1.

#### Model 1: *Sc*Nep1, S233* as the proton–transfer mediator (crystal structure arrangement)

The first cluster model was built based on the crystal structure of the complex of *Sc*Nep1 with RNA (PDB code: 3oin). After cropping the active side region (see Table S6 for the list of residues incorporated in each of the investigated cluster models), the uracil nucleobase was replaced with pseudouracil and a methyl group was added at the sulfur atom of SAH to generate SAM. Additionally, the side chain of S233* was manually rotated around the C_*α*_–C_*β*_ bond, so that the OH group is located between the carboxylate of D101 and the N1 of Ψ.

The prepared model was subjected to a geometry optimization with constraints imposed on the positions of appropriate atoms. In the resulting initial state, the N1 of Ψ, the OH group of S233*, and the carboxylate of D101 form a continuous chain of two hydrogen bonds, aligned properly for proton transfer. Indeed, the transition state for the concerted movement of the two protons, mediated by the OH of S233* as the proton shuttle, is found to have a barrier of only 2.1 kcal/mol (TS1; see Figure 6 for the energy profiles and Figure 7 for the structures of the proton–transfer transition states). It furnishes an intermediate that is 1.0 kcal/mol higher in energy relative to the initial state. The subsequent methyl transfer from the sulfur atom of SAM to the N1 of Ψ takes place via a transition state having a barrier of 29.1 kcal/mol (TS2). This is by far too high for an enzymatic reaction, suggesting that the arrangement of the active site present in the crystal structure is not catalytically competent.

#### Model 2: *Sc*Nep1, S233* as the proton–transfer mediator (MD arrangement)

During the MD simulations, the OH group of S233* spontaneously comes into interaction via hydrogen bonds with the N1 proton of Ψ and the carboxylic group of D101 (Figure 4A). We constructed an active site model based on an MD snapshot containing such an arrangement. Upon geometry optimization, the hydrogen bond bridge is maintained. Interestingly, the computations established that for this model the proton transfer occurs in a stepwise fashion (not explicitly shown in Figure 5). First, the OH of S233* is deprotonated by D101, providing an anionic alkoxide intermediate (Inter. 1). Next, the N1 proton of Ψ is abstracted by the anionic form of S233*. The transition states for the consecutive proton transfers (TS11 and TS12) are proper first–order saddle points on the electronic energy surface, but upon inclusion of the zero–point energy correction their relative energies fall below those of the respective flanking intermediates. Thus, the barrier for the overall proton transfer can be estimated through the energy of Intermediate 1, i.e., it is very low (0.9 kcal/mol). The resulting deprotonated Ψ is stabilized by interactions with R88*, S233*, and A234*, rendering the deprotonation slightly exothermic (by 2.1 kcal/mol). The subsequent methyl transfer step has the energy barrier of 21.9 kcal/mol, a lower value than that calculated above for the same pathway, employing the crystal structure arrangement. This demonstrates that the MD simulations allow for the active site to adopt a structure that is better suited for catalysis than the structure present in the crystal. The insufficiency of the crystal structure may in part originate from the application of substrate and cofactor analogs to grow the crystal (U, instead of Ψ and SAH, instead of SAM, respectively). In contrast, the actual reactive compounds were used in the simulations.

#### Model 3: *Sc*Nep1, water as the proton–transfer mediator

The MD simulations showed that a water molecule can enter the active site and create a hydrogen–bond bridge between the N1 proton of Ψ and the carboxylate of D101, replacing the OH group of S233* as the mediator (Figure 4C). Notably, the active site in this model has displayed some extra conformational flexibility in the simulations. In particular, the water molecule (contrary to the OH of S233*) could form a hydrogen bond with either of the two oxygen atoms of the carboxylate anion of D101. In order to account for this structural flexibility, we constructed models for QM calculations from the MD snapshots representing both of these arrangements of the active site. For one of the arrangements, there was an additional divergence due to different orientations of arginine R132, raising the total number of evaluated models to three. We computed the energy profiles of the methyl transfer for each of the models. Below we describe only the results for the model showing the lowest calculated energy barrier, while the others are presented in Supporting Information (Table S7 and Figure S11).

The results showed that, in the most favorable model, the protons are transferred through a concerted transition state that involves a water molecule as the proton shuttle (TS1). The energy barrier for the proton transfer is calculated to be 4.0 kcal/mol. After the proton transfer, D101 forms hydrogen bonds with the water molecule and R132. The negatively charged Ψ is stabilized by S233* and R88*, which interact with the carbonyl groups of Ψ via hydrogen bonding.

In the subsequent transition state (TS2), the methyl group is located between the sulfur of SAM and the N1 nitrogen of Ψ, approaching nearly in the plane of the nucleobase (S–CH_3_–N1 angle equals 172^*◦*^). As noted in previous studies, methyl transfer is the rate–determining step of the mechanism, constituting the upper bound of the overall barrier of 18.5 kcal/mol. This is the lowest barrier established among the different pathways for the *Sc*Nep1 enzyme.

#### Model 4: *Af* Nep1, T198* as the proton–transfer mediator

In the second set of QM calculations, we aimed to investigate the methylation mechanism for the archaeal homolog *Af* Nep1. The first cluster model was built based on the MD snapshot, wherein the OH group of T198* is positioned to serve as the proton–transfer mediator in active site of *Af* Nep1 (Figure 4B), analogous to S233* in *Sc*Nep1 (Model 2).

The proton transfer step (TS1) was found to be concerted, with a relatively high, but still permissible, barrier of 20.5 kcal/mol. This is nonetheless not critical for the overall reaction rate, as the methyl transfer proceeds through a transition state having a 24.2 kcal/mol barrier (TS2), which will thus determine the kinetics. Hence, the calculated barrier height is somewhat larger, but comparable to that obtained for the equivalent mechanism in *Sc*Nep1 (21.9 kcal/mol, Model 2).

#### Model 5: *Af* Nep1, water as the proton–transfer mediator

Similar to *Sc*Nep1, a water molecule was observed to enter the active site of *Af* Nep1 during the MD simulations and position itself between the proton donor and acceptor (Figure 4D). Two cluster models were constructed to account for the possibility of transferring the proton to either of the oxygen atoms of the carboxylate group of the acceptor aspartate D74. The more energetically favorable model is described here, while the other is presented in the SI.

The proton transfer takes place in a concerted fashion via the water molecule through a 15.8 kcal/mol barrier (TS1). The subsequent transfer of the methyl group requires crossing a relatively low energy barrier of 19.1 kcal/mol (TS2).

In summary, alternative reaction pathways for N1 methylation of pseudouridine by Nep1 enzymes were evaluated by QM modelling. In all cases, the first step of the reaction, i.e., the proton transfer to the distant aspartate acceptor, turned out to be perfectly feasible, provided that the process is mediated by either the OH group of serine or threonine, or by a molecule of water. Notably, while the proton transfer barrier in *Af* Nep1 is higher relative to *Sc*Nep1, it can be overcome. In turn, the calculations establish that the second step of the mechanism, the methyl transfer, constitutes the kinetic bottleneck of the overall reaction, discerning the two alternative pathways. The computations clearly indicate that the arrangement of the active site present in the available crystal structure of the enzyme–substrate complex is not catalytically competent (Model 1). Conversely, the structure of the active site derived from the MD simulations is much better–suited for promoting the reaction. Interestingly, for both enzymes, the methyl transfer step was found to be noticeably easier (by 3–5 kcal/mol), when the preceding proton transfer had been mediated by a water molecule (Models 3 and 5), rather than by the hydroxyl group of the adjacent serine or threonine (Models 2 and 4). Such an outcome can be explained by a slightly different arrangement of the active site when the water molecule is present, which seems to be suited slightly better for promoting the methyl transfer step. However, the energy difference of 3–5 kcal/mol is within the error of the applied computational methodology, therefore, the two alternatives cannot be definitively discriminated based on the results of the QM calculations alone.

### Experimental kinetics of the enzymatic reactions

To validate the computational results and gain additional insights, we measured the kinetic parameters for the methylation of Ψ in 5’–AAU–UUG–ACΨ–CAA–CAC–GGG–GA–3’ oligonucleotide by both *Sc*Nep1 and *Af* Nep1, as well as by variants of *Sc*Nep1 having mutated amino acids in the relevant key positions (Table 1). The measured rate constants for the chemical step (*k* _*cat*_) in the case of wild–type *Sc*Nep1 and *Af* Nep1 are 0.88 s^^1^ at 30^*◦*^C and 0.20 s^^1^ at 55^*◦*^C, respectively. These values correspond to the energy barriers of approximately 18 kcal/mol for *Sc*Nep1 and 20 kcal/mol for *Af* Nep1. Therefore, there is a very good agreement between the computed and experimental data, especially for the models wherein the water molecule plays the role of the proton–transfer mediator (Figure 6, Models 3 and 5 for *Sc*Nep1 and *Af* Nep1, respectively).

**Table 1:**
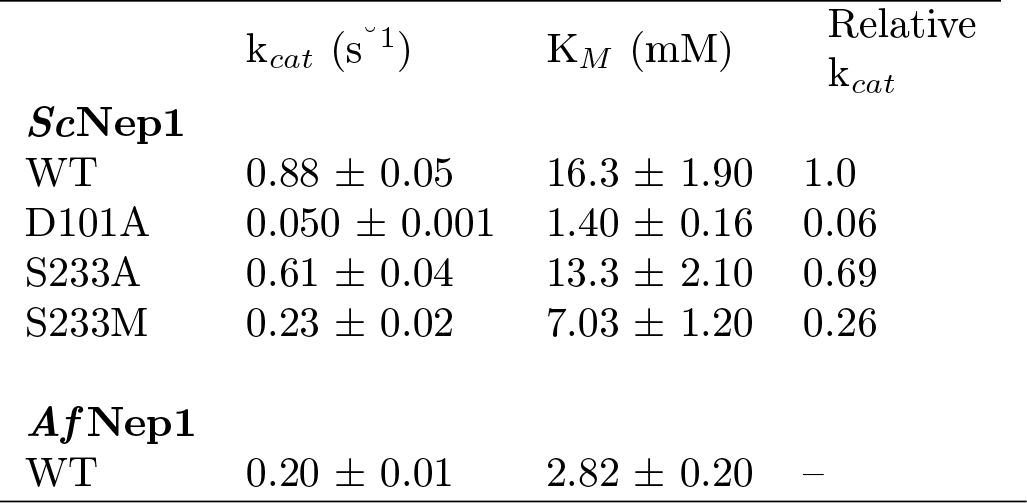
Experimental kinetic parameters for *Sc*Nep1, its three mutated variants (all measured at 30^*◦*^C), and *Af* Nep1 (measured at 55^*◦*^C).

| | $k_{cat}$ ($s^{-1}$) | $K_M$ (mM) | Relative $k_{cat}$ |
| --- | --- | --- | --- |
| ***ScNep1*** |  |  |  |
| WT | $0.88 \pm 0.05$ | $16.3 \pm 1.90$ | 1.0 |
| D101A | $0.050 \pm 0.001$ | $1.40 \pm 0.16$ | 0.06 |
| S233A | $0.61 \pm 0.04$ | $13.3 \pm 2.10$ | 0.69 |
| S233M | $0.23 \pm 0.02$ | $7.03 \pm 1.20$ | 0.26 |
| ***AfNep1*** |  |  |  |
| WT | $0.20 \pm 0.01$ | $2.82 \pm 0.20$ | – |

**Figure 6:**
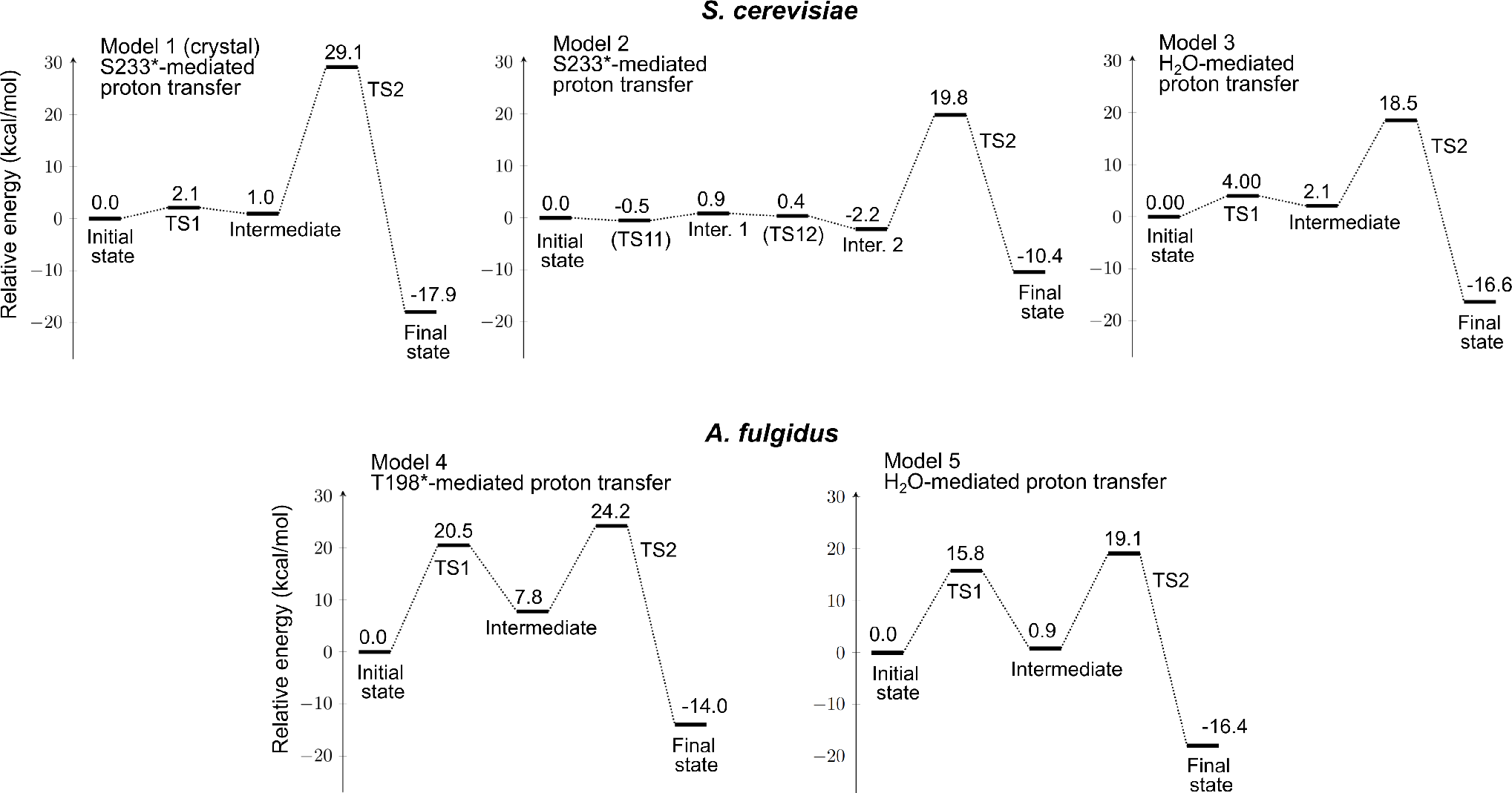
Energy profiles for the methylation of Ψ in the active site of Nep1 proceeding via different evaluated pathways.

**Figure 7:**
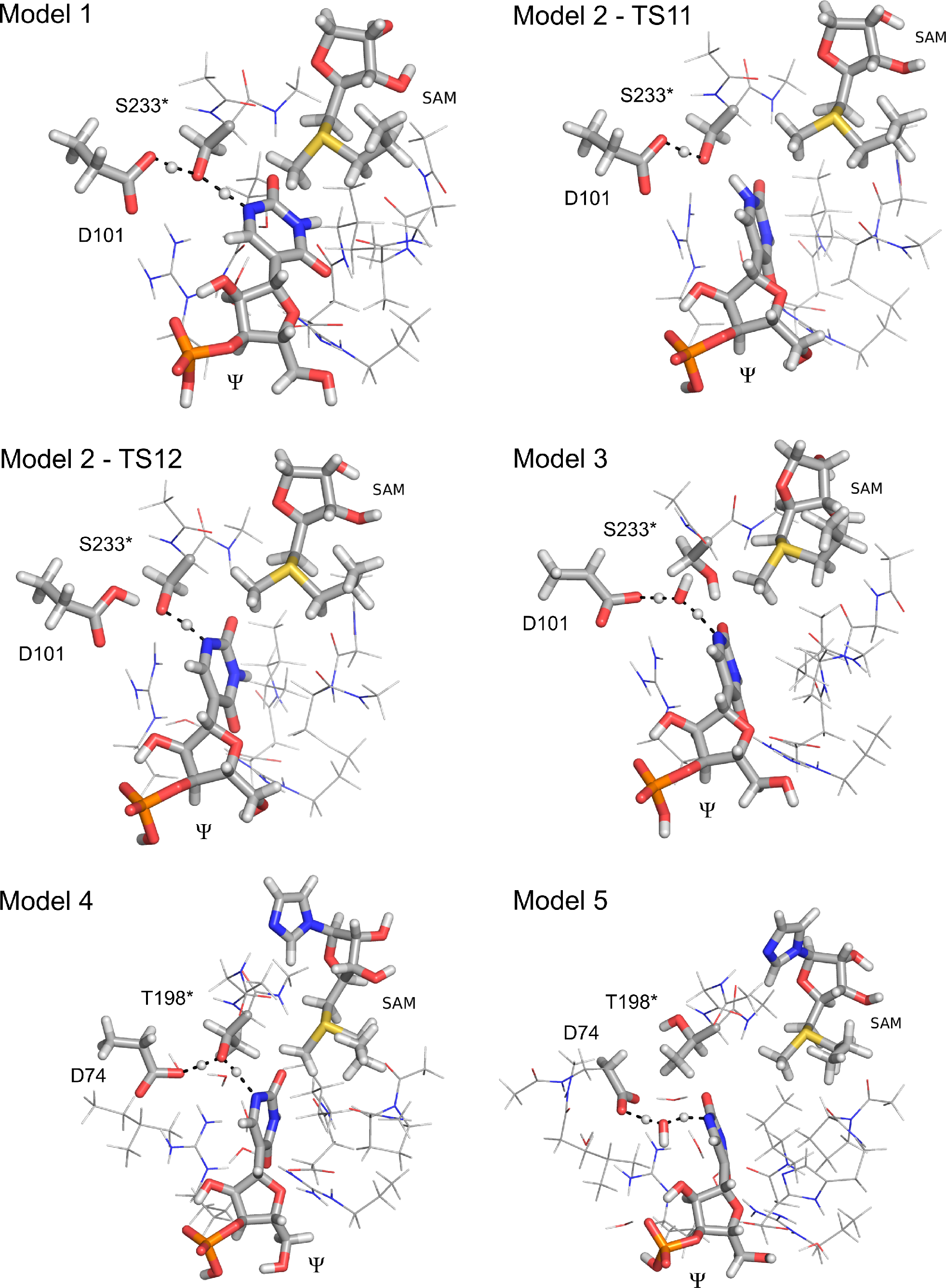
Optimized structures of the transition states for the proton–transfer step (TS1) during the different evaluated pathways.

The mutational studies provide very interesting results. First, the mutation of aspartate D101 to alanine causes an almost 20–fold drop in the reaction rate constant (k_*cat*_), confirming the importance of this residue for catalysis and substantiating its role as the proton acceptor. The concurrent increase in the affinity of the enzyme toward RNA substrate (over 10–fold decrease in K_*M*_) most likely originates from rendering the active site less negatively charged, thus lowering the repulsion with the RNA phosphate backbone. Secondly, regarding the supposed proton–transfer mediator, the k_*cat*_ value shows an insignificant decrease upon the mutation of S233 to alanine, whereas it decreases only moderately when methionine is introduced into this position. Therefore, the hydroxyl–containing amino acid does not seem to be critical for maintaining the catalytic efficiency of the enzyme, implying that it likely does not act as the proton–transfer mediator. The results thus lend support to the water-mediated pathway, in which a water molecule serves as the mediator between proton donor and acceptor, corroborating the QM data (Models 3 and 5).

Importantly, a similar water–mediated proton transfer mode can be applied to rationalize the mechanism of catalysis in the archaeal Nep1 homologs, which do not have a hydroxyl–containing amino acid located in the active site in between Ψ and the proton acceptor (the second class of archaeal proteins in Figure 3). This hypothesis is also supported by the retention of the catalytic activity of the *Sc*Nep1 S233M mutant or S233A mutant of the yeast enzyme (Table 1).

## Conclusions

In summary, using a hybrid workflow encompassing bioinformatics analysis, molecular docking, MD simulations, and QM calculations we have established the mechanism of catalysis by methyltransferase Nep1, responsible for the N1 methylation of pseudouridine during the processing of rRNA.

Our initial hypothesis, based on the alignment of Nep1 sequences from different organisms and crystallographic data, that the first, proton–transfer, step of the reaction is mediated by a hydroxyl–containing amino acid residue, has proven to be incorrect. Namely, the MD simulations, consistent in both examined Nep1 homologs, identified the active site arrangements with a water molecule bridging the N1 of Ψ and the putative aspartate proton acceptor through two contiguous hydrogen bonds. In turn, the QM calculations established that the methylation reactions proceed via lower energy barriers when the water is mediating the proton transfer compared to the pathways involving the OH group of serine/threonine being the proton–shuttle.

The results of QM computations were confirmed by experimental kinetics and mutational analysis, which also validated the identity of the above mentioned aspartate as the proton acceptor. Interestingly, it is not the proton transfer that distinguishes between the mechanistic alternatives, but rather the barrier for the subsequent methyl transfer, instead. The water molecule not only mediates the proton transfer, but its presence also introduces conformational changes of the active site that facilitate the overall methylation reaction. Importantly, the uncovered water–mediated proton transfer may be a general pathway implicated in the catalytic mechanisms of other enzymes, which require a translocation of protons across extended distances in the active site.

From the methodological perspective, current work clearly demonstrates the strengths and advantages of a comprehensive approach employing multiple complementary computational techniques, supplemented by experimental verification of the key conclusions. In particular, the combination of MD simulations and QM modeling has proven to be exceptionally powerful, allowing for the discovery of the unexpected water–mediated proton transfer pathway, which would not be possible to detect using separate methods and isolated pieces of data.

## Methods

### Topology analysis

The Topoly package was used to identify the location of the knot in all Nep1 structures.^32^ In the case of Nep1 from *S. cerevisiae*, the knot core consists of 47 amino acids and involves residues from T179 to V225 in both chains. The N– and C–tails are between residues 1–151 and 199–225, respectively.^7^

### Sequence alignment analysis

PSI–BLAST was used to find *Sc*Nep1 homologs in the UniProtKB/Swiss–Prot database. ^33,34^ For Nep1 sequences from eukaryotic organisms, the Reference Sequence Database was additionally used. Multiple sequence alignments were carried out with Clustal Omega with default parameters.^22^

### Docking of rRNA to *Af* Nep1

Full rRNA fragment, 5’–GGG–CUU–CAA–CGC–CC–3’, was cut out from the crystal structure of the complex with *Sc*Nep1 (PDB code: 3oin). From the crystal structure of *Af* Nep1 holoenzyme (PDB code: 3o7b), water molecules and ligands (tyrosine and SAH molecules) were removed and any atoms missing from the crystal structure were added using the Maestro Schrödinger software.^35^ The docking was performed using the NPDock server.^23^ Obtained conformations were screened and selected according to their RMSD after alignment with the 3oin crystal structure (RMSD below 4 Å), and the remaining complexes were analyzed for distances suitable to form key hydrogen bonds between the protein (specifically, with arginines R88*, R129, R132, and R136) and the rRNA (cutoff 4 Å, Table S2 lists the specific hydrogen bonds that were evaluated). The resulting structures were then checked for any heavy–atom overlaps, and a single complex was chosen to proceed to the next stage. The SAH ligands were added back to the protein structure. A single backbone atom was missing, which was fixed using the PDB2PQR server (the hydrogen atoms were then removed).^36^ Finally, SAH was modified to SAM, by the addition of the methyl group at sulfur and U6 was modified to Ψ.

### Molecular Dynamics Simulations

The reconstructed complexes were protonated in pH 7.0 using PDB2PQR server^36^ and neutralized by adding sodium cations and chloride anions. The explicit water TIP3P model was used. The MD simulations were performed in GROMACS 2018.1 software package with the CHARMM36 force field.^37,38^ All performed production runs shared common first steps of the system preparations, including the energy minimization and the relaxation.

The equilibration prior to the production runs included an NVT ensemble and two (for *Sc*Nep1) or three (for *Af* Nep1) phases of NPT ensemble (Table S3). For the temperature equilibration, a V–rescale temperature coupling was used, at 310K for the yeast protein and at 338K for the archeal protein. For the pressure equilibration, we employed Berendsen barostat.^39^ The restraints on the positions of atoms were gradually loosened to provide better relaxation. Specifically, in the respective equilibration steps, a position restraining force was applied on the heavy atoms of the complex (NVT and NPT1), then on the main–chain atoms of the protein and selected heavy atoms of the rRNA (NPT2), and finally on the *C*_*α*_ atoms of the protein and the P atoms of the rRNA (NPT3).

All–atom MD production stages were performed independently (Table S4). All trajectories were evaluated by a standard stability analysis including: RMSD (root–mean–square deviation) for monitoring the *C*_*α*_ (Figure S2), the side chains of protein, as well as the movements of SAM and rRNA; RMSF (root–mean–square fluctuation) to identify most mobile residues in the protein and the nucleic acid chain; hydrogen bond number to check the stability of the complex (Figure S3).

To select snapshots for the construction of the active site models for the QM calculations, first, the MD trajectories were screened using a Python script, evaluating the distances between the atoms supposed to participate in the proton and methyl transfers (Figure S5): between the OH group of S233*/T198* and the N1 of Ψ; (2) between the carboxylate oxygens of D101/D74 and the OH group of S233*/T198*; (3) between the carboxylate oxygens of D101/D74 and the N1 of Ψ (for the water–mediated proton transfer); (4) between the methyl group of SAM and the N1 of Ψ. Additionally, for *Af* Nep1, the dihedral angle around the bond between *C*_*α*_ and *C*_*β*_ of T198* was also evaluated, so that the OH was in a proper orientation (Figure S10). Next, the snapshots with the distances and the dihedral angle meeting the assigned cutoff criteria were inspected and a representative one was selected manually for each of the distinct arrangements of the active site.

### Quantum chemical calculations

All calculations were carried out using Gaussian 16 package.^40^ The active site models were constructed based on the snapshots from MD simulations (see Table S6 for the list of residues incorporated in each of the investigated cluster models). Geometry optimizations were performed with B3LYP functional^41–45^ including DFT–D3 dispersion correction with BJ dumping,^46^ using Def2–SVP basis set. ^47^ Coordinates of selected atoms were fixed during geometry optimizations to prevent unrealistic movements of residues (Figures S12–19). At the same level of theory, frequency calculations were performed to obtain ZPE corrections and single–point CPCM solvation^48,49^ energies were calculated (with *ϵ* = 4) to model the effects of the rest of the enzyme. For optimized geometries of the stationary points, single–point calculations with the Def2–QZVP basis set were carried out to obtain more accurate energies. The final reported energy values are those for the large basis set corrected for ZPE and solvation effects.

### Expression and purification of proteins

Five proteins were prepared: *Af* Nep1 and *Sc*Nep1 along with its three mutants (D101A, S233A, S233M). The appropriate mutants were prepared by PCR reaction through mutagenesis. Each protein was subcloned into *E. coli* BL21 DE3 RiL and grown in LB medium under the control of 1 mM IPTG. Proteins were purified by ion affinity chromatography and size exclusion chromatography. The details are given in the Supporting Information.

### Kinetic measurements

Steady–state conditions were achieved for Nep1 at 75 nM and RNA from 1 to 30 *µ*M. All *S. cerevisiae* protein assays were performed at 30^*◦*^C and *A. fulgidus* Nep1 was assayed at 55^*◦*^C.

## Acknowledgement

This work was supported by EMBO Installation Grant 2057 (to JIS), the Polish Ministry for Science and Higher Education 0003/ID3/2016/64 (to JIS). This research was carried out with the support of PLGrid Infrastructure.

## References

(1) Cantoni, G. L. Biological methylation: selected aspects. Annu. Rev. Biochem. 1975, 44, 435–451.

(2) Perlinska, A. P.; Stasiulewicz, A.; Nawrocka, E. K.; Kazimierczuk, K.; Setny, P.; Sulkowska, J. I. Restriction of S-adenosylmethionine conformational freedom by knotted protein binding sites. PLoS Comput. Biol. 2020, 16, e1007904.

(3) Schubert, H. L.; Blumenthal, R. M.; Cheng, X. Many paths to methyltransfer: a chronicle of convergence. Trends Biochem. Sci. 2003, 28, 329–335.

(4) Anantharaman, V.; Koonin, E. V.; Aravind, L. SPOUT: a class of methyltransferases that includes spoU and trmD RNA methylase superfamilies, and novel superfamilies of predicted prokaryotic RNA methylases. J. Mol. Microbiol. Biotechnol. 2002, 4, 71–76.

(5) Krishnamohan, A.; Jackman, J. E. A family divided: distinct structural and mechanistic features of the SpoU-TrmD (SPOUT) methyltransferase superfamily. Biochemistry 2018, 58, 336–345.

(6) Jarmolinska, A. I.; Perlinska, A. P.; Runkel, R.; Trefz, B.; Ginn, H. M.; Virnau, P.; Sulkowska, J. I. Proteins’ knotty problems. J. Mol. Biol. 2019, 431, 244–257.

(7) Dabrowski-Tumanski, P.; Rubach, P.; Goundaroulis, D.; Dorier, J.; Sułkowski, P.; Millett, K. C.; Rawdon, E. J.; Stasiak, A.; Sulkowska, J. I. KnotProt 2.0: a database of proteins with knots and other entangled structures. Nucleic Acids Res. 2019, 47, D367–D375.

(8) Nureki, O.; Watanabe, K.; Fukai, S.; Ishii, R.; Endo, Y.; Hori, H.; Yokoyama, S. Deep knot structure for construction of active site and cofactor binding site of tRNA modification enzyme. Structure 2004, 12, 593–602.

(9) Christian, T.; Sakaguchi, R.; Perlinska, A. P.; Lahoud, G.; Ito, T.; Taylor, E. A.; Yokoyama, S.; Sulkowska, J. I.; Hou, Y.-M. Methyl transfer by substrate signaling from a knotted protein fold. Nat. Struct. Mol. Biol. 2016, 23, 941–948.

(10) Dabrowski-Tumanski, P.; Stasiak, A.; Sulkowska, J. I. In search of functional advantages of knots in proteins. PloS one 2016, 11, e0165986.

(11) Virnau, P.; Mirny, L. A.; Kardar, M. Intricate knots in proteins: Function and evolution. PLoS Comput. Biol. 2006, 2, e122.

(12) Wurm, J. P.; Meyer, B.; Bahr, U.; Held, M.; Frolow, O.; Kötter, P.; Engels, J. W.; Heckel, A.; Karas, M.; Entian, K.-D., et al. The ribosome assembly factor Nep1 responsible for Bowen– Conradi syndrome is a pseudouridine-N1specific methyltransferase. Nucleic Acids Res. 2010, 38, 2387–2398.

(13) Leulliot, N.; Bohnsack, M. T.; Graille, M.; Tollervey, D.; Van Tilbeurgh, H. The yeast ribosome synthesis factor Emg1 is a novel member of the superfamily of alpha/beta knot fold methyltransferases. Nucleic Acids Res. 2008, 36, 629–639.

(14) Eschrich, D.; Buchhaupt, M.; Kötter, P.; Entian, K.-D. Nep1p (Emg1p), a novel protein conserved in eukaryotes and archaea, is involved in ribosome biogenesis. Curr. Genet. 2002, 40, 326–338.

(15) Meyer, B.; Wurm, J. P.; Kötter, P.; Leisegang, M. S.; Schilling, V.; Buchhaupt, M. Held, M.; Bahr, U.; Karas, M.; Heckel, A., et al. The Bowen–Conradi syndrome protein Nep1 (Emg1) has a dual role in eukaryotic ribosome biogenesis, as an essential assembly factor and in the methylation of Ψ1191 in yeast 18S rRNA. Nucleic Acids Res. 2011, 39, 1526–1537.

(16) Armistead, J.; Khatkar, S.; Meyer, B.; Mark, B. L.; Patel, N.; Coghlan, G.; Lamont, R. E.; Liu, S.; Wiechert, J.; Cattini, P. A., et al. Mutation of a gene essential for ribosome biogenesis, EMG1, causes Bowen-Conradi syndrome. Am. J. Hum. Genet. 2009, 84, 728–739.

(17) Hunter, A. G.; Woerner, S. J.; Montalvo-Hicks, L. D.; Fowlow, S. B.; Haslam, R. H.; Metcalf, P. J.; Lowry, R. B.; Optiz, J. M. The Bowen-Conradi syndrome—A highly lethal autosomal recessive syndrome of microcephaly, micrognathia, low birth weight, and joint deformities. Am. J. Med. Genet. 1979, 3, 269–279.

(18) Taylor, A. B.; Meyer, B.; Leal, B. Z.; Kötter, P.; Schirf, V.; Demeler, B.; Hart, P. J.; Entian, K.-D.; Wöhnert, J. The crystal structure of Nep1 reveals an extended SPOUT-class methyltransferase fold and a pre-organized SAM-binding site. Nucleic Acids Res. 2008, 36, 1542–1554.

(19) Thomas, S. R.; Keller, C. A.; Szyk, A.; Cannon, J. R.; LaRonde-LeBlanc, N. A. Structural insight into the functional mechanism of Nep1/Emg1 N1-specific pseudouridine methyltransferase in ribosome biogenesis. Nucleic Acids Res. 2011, 39, 2445–2457.

(20) Perlinska, A. P.; Kalek, M.; Christian, T.; Hou, Y.-M.; Sulkowska, J. I. Mg2+Dependent Methyl Transfer by a Knotted Protein: A Molecular Dynamics Simulation and Quantum Mechanics Study. ACS Catal. 2020, 10, 8058–8068.

(21) Ito, T.; Masuda, I.; Yoshida, K.-i.; GotoIto, S. Sekine, S.-i.; Suh, S. W.; Hou, Y.- M.; Yokoyama, S. Structural basis for methyl-donor–dependent and sequencespecific binding to tRNA substrates by knotted methyltransferase TrmD. Proc. Natl. Acad. Sci. 2015, 112, E4197–E4205.

(22) Madeira, F.; Park, Y. M.; Lee, J.; Buso, N.; Gur, T.; Madhusoodanan, N.; Basutkar, P.; Tivey, A. R.; Potter, S. C.; Finn, R. D., et al. The EMBL-EBI search and sequence analysis tools APIs in 2019. Nucleic Acids Res. 2019, 47, W636–W641.

(23) Tuszynska, I.; Magnus, M.; Jonak, K.; Dawson, W.; Bujnicki, J. M. NPDock: a web server for protein–nucleic acid docking. Nucleic Acids Res. 2015, 43, W425–W430.

(24) EMBL-EBI, RNAcentral DataBase.

(25) Himo, F. Quantum chemical modeling of enzyme active sites and reaction mechanisms. Theor. Chem. Acc. 2006, 116, 232–240.

(26) Himo, F. Recent trends in quantum chemical modeling of enzymatic reactions. J. Am. Chem. Soc. 2017, 139, 6780–6786.

(27) Siegbahn, P. E.; Himo, F. Recent developments of the quantum chemical cluster approach for modeling enzyme reactions. J. Biol. Inorg. Chem. 2009, 14, 643–651.

(28) Siegbahn, P. E.; Himo, F. The quantum chemical cluster approach for modeling enzyme reactions. Wiley Interdiscip. Rev. Comput. Mol. Sci. 2011, 1, 323–336.

(29) Velichkova, P.; Himo, F. Methyl transfer in glycine N-methyltransferase. A theoretical study. J. Phys. Chem. B 2005, 109, 8216–8219.

(30) Velichkova, P.; Himo, F. Theoretical study of the methyl transfer in guanidinoacetate methyltransferase. J. Phys. Chem. B 2006, 110, 16–19.

(31) Georgieva, P.; Wu, Q.; McLeish, M. J.; Himo, F. The reaction mechanism of phenylethanolamine N-methyltransferase: a density functional theory study. Biochim Biophys Acta Proteins Proteom. 2009, 1794, 1831–1837.

(32) Dabrowski-Tumanski, P.; Rubach, P.; Niemyska, W.; Gren, B. A.; Sulkowska, J. I. Topoly: Python package to analyze topology of polymers. Brief. Bioinformatics 2021, 22, bbaa196.

(33) Altschul, S. F.; Madden, T. L.; Schäffer, A. A.; Zhang, J.; Zhang, Z.; Miller, W.; Lipman, D. J. Gapped BLAST and PSI-BLAST: a new generation of protein database search programs. Nucleic Acids Res. 1997, 25, 3389–3402.

(34) Altschul, S. F.; Wootton, J. C.; Gertz, E. M.; Agarwala, R.; Morgulis, A.; Schäffer, A. A.; Yu, Y.-K. Protein database searches using compositionally adjusted substitution matrices. FEBS J. 2005, 272, 5101–5109.

(35) Schrödinger, LLC: New York, NY, USA, (Schrödinger Release 2020–4. https://www.schrodinger.com/.

(36) Dolinsky, T. J.; Nielsen, J. E.; McCammon, J. A.; Baker, N. A. PDB2PQR: an automated pipeline for the setup of Poisson–Boltzmann electrostatics calculations. Nucleic Acids Res. 2004, 32, W665–W667.

(37) Rb, B.; X, Z.; J, S.; Pe, L.; J, M.; Mf Jr., M. A. Optimization of the additive CHARMM all-atom protein force field targeting improved sampling of the backbone ϕ, ψ and side-chain χ1 and χ2 dihedral angles. J. Chem. Theory Comput. 2012, 8, 3257–3273.

(38) Best, R. B.; Zhu, X.; Shim, J.; Lopes, P. E.; Mittal, J.; Feig, M.; MacKerell Jr, A. D. Optimization of the additive CHARMM all-atom protein force field targeting improved sampling of the backbone ϕ, ψ and side-chain χ1 and χ2 dihedral angles. J. Chem. Theory Comput. 2012, 8, 3257–3273.

(39) Berendsen, H. J.; Postma, J. v.; van Gunsteren, W. F.; DiNola, A.; Haak, J. R. Molecular dynamics with coupling to an external bath. J. Chem. Phys. 1984, 81, 3684–3690.

(40) Frisch, M. J. et al. Gaussian 16 Revision C.01. 2016; Gaussian Inc. Wallingford CT.

(41) Lee, C.; Yang, W.; Parr, R. G. Development of the Colle-Salvetti correlation-energy formula into a functional of the electron density. Phys. Rev. B 1988, 37, 785–789.

(42) Becke, A. D. Density-functional exchange-energy approximation with correct asymptotic behavior. Phys. Rev. A 1988, 38, 3098–3100.

(43) Becke, A. Density-functional thermochemistry. I. The effect of the exchange-only gradient correction. J. Chem. Phys. 1992, 96, 2155–2160.

(44) Becke, A. D. Density-functional thermochemistry. II. The effect of the Perdew-Wang generalized-gradient correlation correction. J. Chem. Phys. 1992, 97, 9173–9177.

(45) Becke, A. D. A new mixing of Hartree-Fock and local density-functional theories. J. Chem. Phys. 1993, 98, 1372–1377.

(46) Grimme, S.; Ehrlich, S.; Goerigk, L. Effect of the Damping Function in Dispersion Corrected Density Functional Theory. J. Comput. Chem. 2011, 32, 1456–65.

(47) Weigend, F.; Ahlrichs, R. Balanced basis sets of split valence, triple zeta valence and quadruple zeta valence quality for H to Rn: Design and assessment of accuracy. Phys. Chem. Chem. Phys. 2005, 7, 3297–3305.

(48) Barone, M., Vincenzo aRestriction of S-adenosylmethionine conformational freedom by knotted protein binding sitesnd Cossi Quantum Calculation of Molecular Energies and Energy Gradients in Solution by a Conductor Solvent Model. J. Phys. Chem. A 1998, 102, 1995–2001.

(49) Cossi, M.; Rega, N.; Scalmani, G.; Barone, V. Energies, structures, and electronic properties of molecules in solution with the C-PCM solvation model. J. Comput. Chem. 2003, 24, 669–681.

