## Supporting Information for "Nucleolar Essential Protein 1 (Nep1): Elucidation of Enzymatic Catalysis Mechanism by Combined Molecular Dynamics Simulation and Quantum Chemical Calculations"

#### Contents

**1 Crystal structure analysis** **S1**

**2 MD simulations** **S2**
2.1 Hydrogen bonds analysis . . . . . . . . . . S4

**3 QM calculations** **S8**

**4 Experimental procedures** **S85**
4.1 Purification and expression of Nep1 . . . . . . . . . . S85
4.2 Sequences of expressed proteins . . . . . . . . . . S85

### 1 Crystal structure analysis

Table S1: The values of $C_\alpha$ RMSD (Å) between the available crystal structures of Nep1: *S. cerevisiae* (3oin, 3oii, 3oij, 2v3k, 2v3j), *A. fulgidus* (3o7b),*M. jannaschii* (3bbd, 3bbe, 3bbh) and *H.sapiens* (5fai). Structures used for further studies are indicated in bold.

| | **3oin** | 3oii | 3oij | 2v3j | 2v3k | **3o7b** | 5fai | 3bbd | 3bbe |
|---|---|---|---|---|---|---|---|---|---|
| **3oin** | | | | | | | | | |
| 3oii | 1.5 | | | | | | | | |
| 3oij | 1.7 | 1.3 | | | | | | | |
| 2v3j | 0.6 | 0.5 | 0.7 | | | | | | |
| 2v3k | 1.2 | 0.8 | 1.4 | 0.3 | | | | | |
| **3o7b** | 2.6 | 2.6 | 2.7 | 3.1 | 2.6 | | | | |
| 5fai | 1.8 | 1.9 | 2.0 | 1.8 | 1.9 | 2.8 | | | |
| 3bbd | 4.1 | 4.2 | 3.7 | 3.0 | 2.9 | 2.9 | 3.2 | | |
| 3bbe | 3.5 | 3.6 | 3.3 | 3.2 | 3.2 | 2.9 | 3.6 | 0.2 | |
| 3BBH | 5.6 | 3.8 | 3.5 | 3.0 | 2.9 | 2.9 | 3.2 | 0.2 | 0.2 |

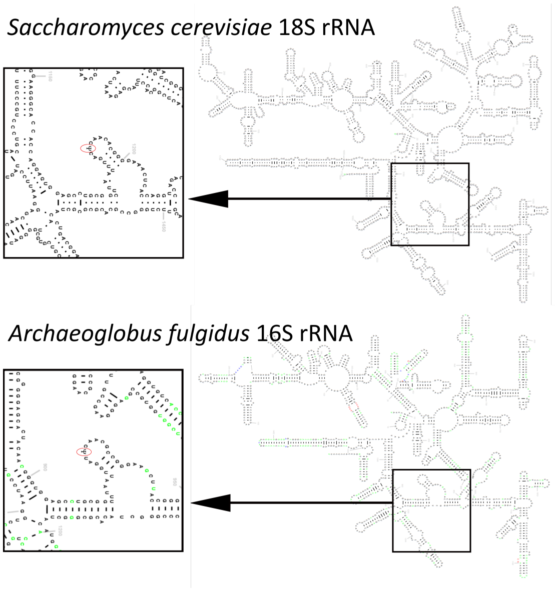


Figure S1: Comparison of the secondary structure of 18S rRNA from *S. cerevisiae* and 16S rRNA from *A. fulgidus* (EMBL-EBI, RNAcentral DataBase). The loops, in which the methylated Ψ is located exhibit high similarity between the two organisms investigated in this work.

Table S2: Key interactions between *Sc*Nep1 and rRNA in crystal structure 3oin (NEF atom names). The hydrogen bonds used during the evaluation of the complexes obtained by the docking between holoenzyme *Af*Nep1 and rRNA fragment are indicated in bold.

| Residue | Nucleotide | Type of interaction |
|---|---|---|
| R136 | U5 | **Hydrogen bonds (NH1, NH2 – O2)** |
| R132 | U6 | **Hydrogen bond (NE – O2)** |
| S235* | U6 | Hydrogen bonding via a water molecule (N – O4) |
| Q94* | U6 | Hydrogen bond via a water molecule (OE1 – O4, N3) |
| R88* | U6 | **Hydrogen bonds (NH1, NH2 – O2)** |
| R88* | U6 | **Hydrogen bond (NH1 – O4')** |
| R132 | C7 | **Hydrogen bond (N – O2)** |
| R136 | C7 | Stacking |
| R129 | A8 | Stacking |
| L159 | A8 | Van der Waals interaction |
| T127 | A8 | Hydrogen bond (O' – N6) |
| T127 | A8 | Hydrogen bonding via a water molecule (OG1 – N6) |
| R129 | A8 | **Hydrogen bond (N – N1)** |
| R129 | A9 | **Hydrogen bond (O' – N6)** |
| V128 | A9 | Van der Waals interaction with the side chain |
| L140 | A9 | Stacking |
| R136 | C10 | **Hydrogen bond (NH2, NE, O' – N3, O2, O2')** |

### 2 MD simulations

Table S3: Simulation times for each step of the relaxation procedure, before the MD production stage. For temperature equilibration, a V–rescale description was used, and for pressure equilibration, the Berendsen model.

| Time (ns) | NVT | NPT1 | NPT2 | NPT3 |
|---|---|---|---|---|
| *Sc*Nep1 | 1 | 1 | – | 10 |
| *Af*Nep1 | 1 | 1 | 30 | 10 |

Table S4: Duration of MD simulations for the complexes of *Sc*Nep1 and *Af*Nep1 with rRNA. Trajectories from which snapshots were taken to construct the cluster models for QM calculations are indicated in bold.

| Time (ns) | md1 | md2 | md3 | md4 | md5 | md6 |
|---|---|---|---|---|---|---|
| *Sc*Nep1 | **200** | 200 | 800 | – | – | – |
| *Af*Nep1 | **500** | 300 | 500 | 300 | 300 | 500 |

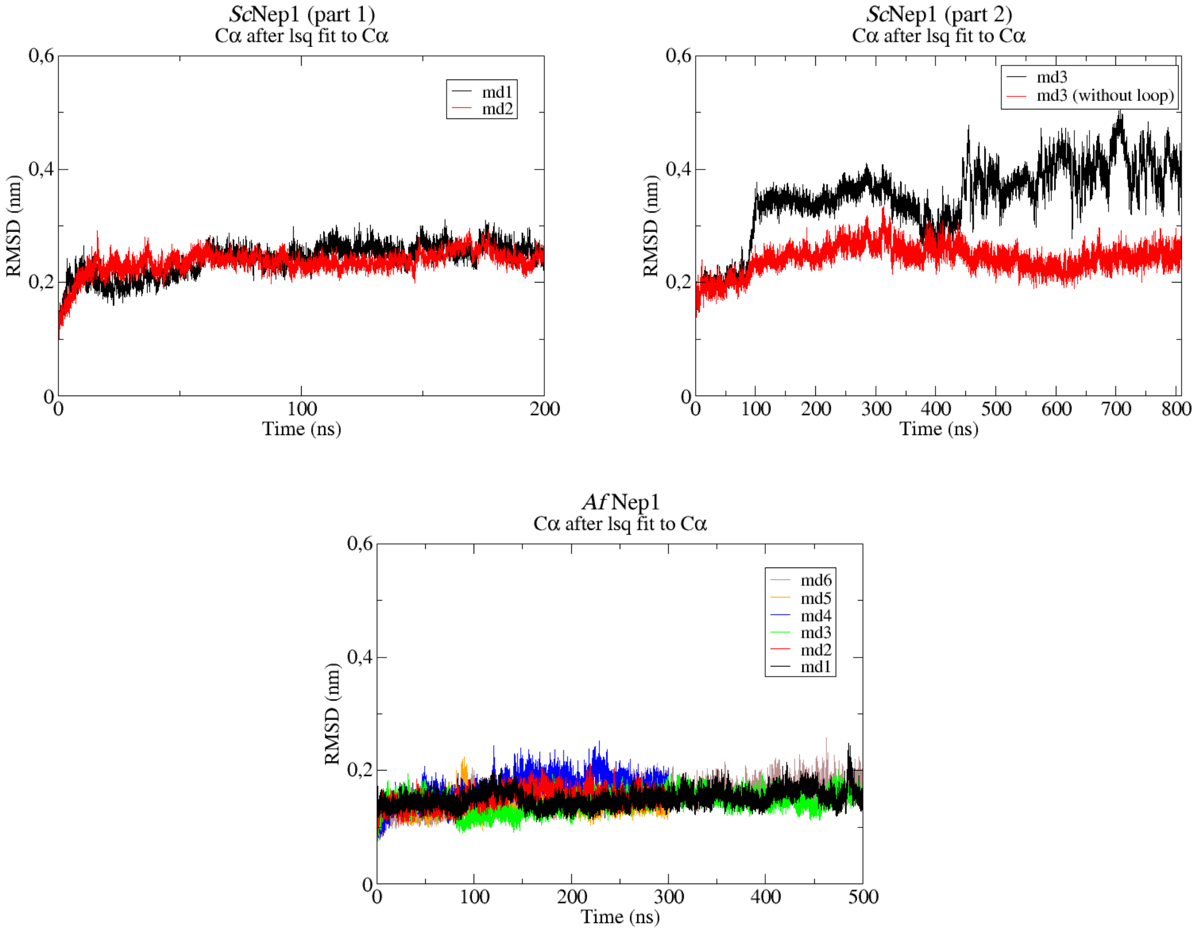


Figure S2: The $C_\alpha$ RMSD values for the different MD simulations of *Sc*Nep1 and *Af*Nep1. The RMSD increase for *Sc*Nep1 md3 (part 2 – top right) was due to movements of a flexible loop at the N-end of the protein chain. RMSD excluding the loop residues is presented for comparison (red), showing a good stability of the protein core.

#### 2.1 Hydrogen bonds analysis

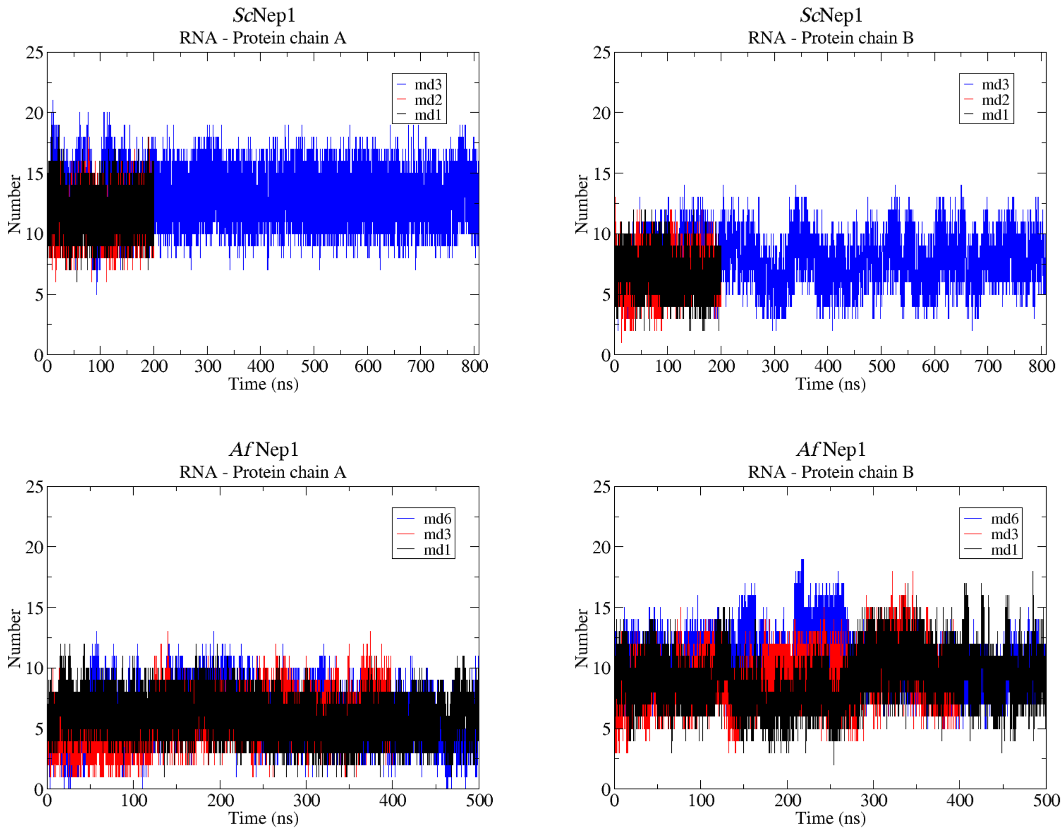


Figure S3: Number of hydrogen bonds formed between the rRNA and the two peptide chains of the protein across selected *Sc*Nep1 and *Af*Nep1 MD simulations, including md1 used for constructing the active site models. The number of hydrogen bonds between the nucleic acid and the protein chains remains approximately constant through the simulations. The number of hydrogen bonds to each of the two peptide chains is different because the rRNA structure is not symmetrical. The data suggests that the two proteins bind rRNA somewhat differently: in *Sc*Nep1 there are more interactions with chain A of the protein, whereas in *Af*Nep1 more hydrogen bonds are found to chain B.

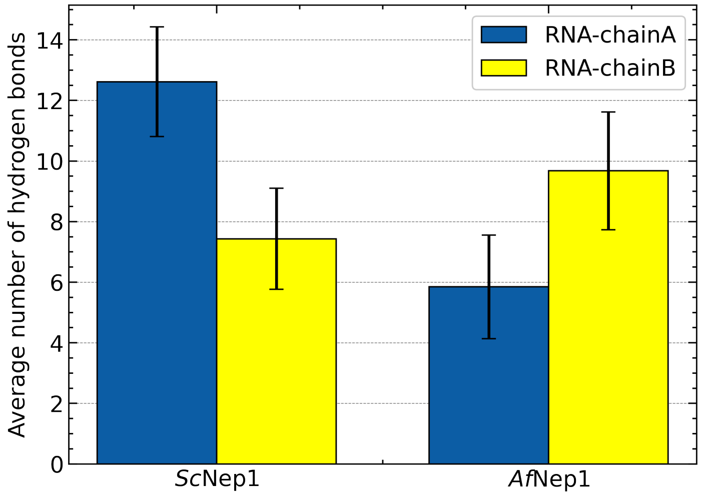


Figure S4: Average number (and standard deviation) of hydrogen bonds between the rRNA and both protein chains in MD simulations. Cutoffs used for H–bond calculations are 3.5Å for donor–acceptor distance and 30° for the angle.

Table S5: Occurrence of particular hydrogen bonds between rRNA and the four conserved arginines in MD simulations (NEF atom names). Results are shown for occupancy >10% in at least one of the organisms. Cutoffs used for H–bond calculations are 3.5Å for donor–acceptor distance and 30° for the angle

| Hydrogen bond | | Occupancy [%] | |
|---|---|---|---|
| Residue | Nucleotide | S. cerevisiae | A. fulgidus |
| R129 (Main–N) | A8 (N1) | 95.44 | 91.43 |
| R129 (Main–O) | A9 (N6) | 93.48 | 86.05 |
| R129 (Side–NH1) | C7 (O2P) | 82.20 | 3.23 |
| R129 (Side–NH1) | A8 (O1P) | 44.39 | 1.78 |
| R129 (Side–NH1) | A8 (O5') | 18.92 | 1.74 |
| R129 (Side–NH1) | C7 (O3') | 14.18 | 0.86 |
| R129 (Side–NH2) | C7 (O1P) | 0.00 | 36.29 |
| R129 (Side–NE) | C7 (O1P) | 0.00 | 33.65 |
| R136 (Side–NE) | C10 (O2) | 94.15 | 25.81 |
| R136 (Side–NH2) | C10 (N3) | 86.73 | 38.95 |
| R136 (Side–NH2) | U5 (O2) | 55.24 | 42.01 |
| R136 (Main–O) | C10 (O2') | 22.32 | 24.34 |
| R136 (Side–NH2) | C10 (O2) | 1.70 | 10.46 |
| R88 (Side–NH2) | Ψ6 (O4) | 87.10 | 76.44 |
| R88 (Side–NH1) | Ψ6 (O4') | 63.70 | 60.33 |
| R88 (Side–NH1) | Ψ6 (O1P) | 56.28 | 0.25 |
| R132 (Main–N) | C7 (O4') | 40.45 | 33.77 |
| R132 (Side–NE) | Ψ6 (O4) | 5.17 | 10.50 |

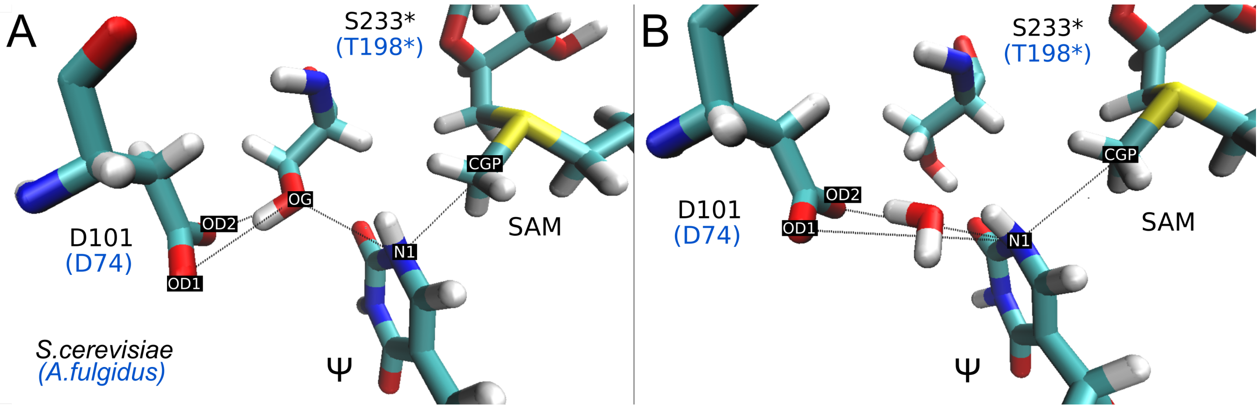


Figure S5: Distances evaluated to identify MD snapshots suitable for the construction of active-site models for the QM calculations (NEF atom names): (A) Serine/threonine–mediated proton transfer: between OG(OG1) of S233*(T198*) and N1 of Ψ (cutoff 4.0 Å); between OD1 or OD2 of D101(D74) and OG(OG1) of S233*(T198*) (smaller of the two; cutoff 4.0 Å); between CGP of SAM and N1 of Ψ (cutoff 4.0 Å). (B) Water–mediated proton transfer: between OD1 or OD2 of D101(D74) and N1 of Ψ (smaller of the two; cutoff 7.0 Å); between CGP of SAM and N1 of Ψ (cutoff 4.0 Å). The variations of these distances during the MD simulations are shown in Figures S6–9.

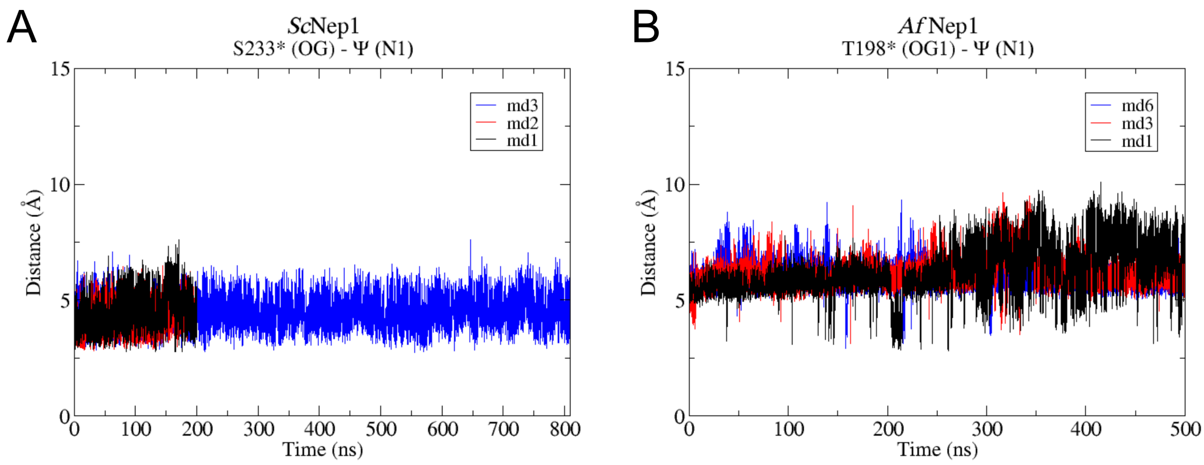


Figure S6: Variation of the distance between OG(OG1) of S233*(T198*) and N1 of Ψ during the MD simulations of (A) *Sc*Nep1 and (B) *Af*Nep1.

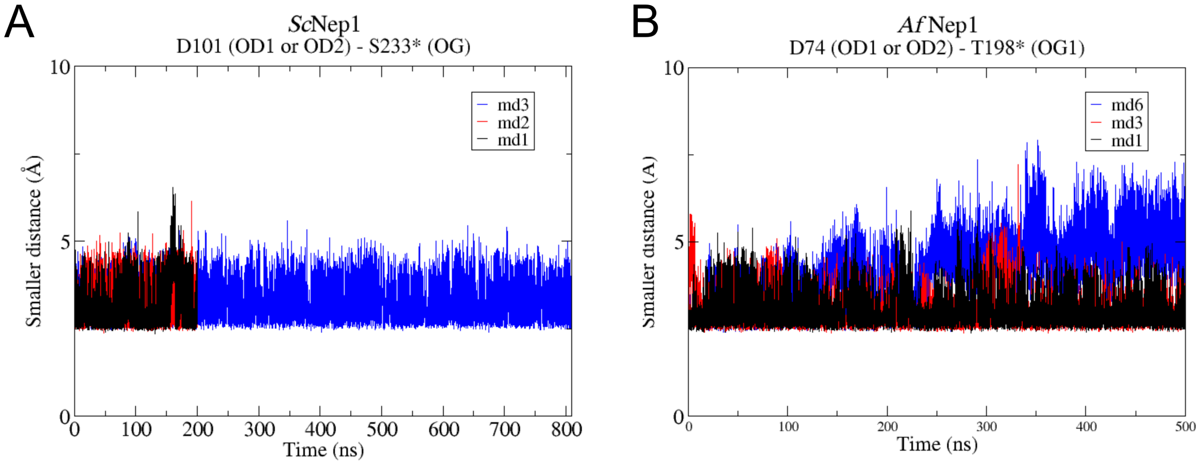


Figure S7: Variation of the distance between OD1 or OD2 of D101(D74) and OG(OG1) of S233*(T198*) (smaller of the two) during the MD simulations of (A) *Sc*Nep1 and (B)*Af*Nep1

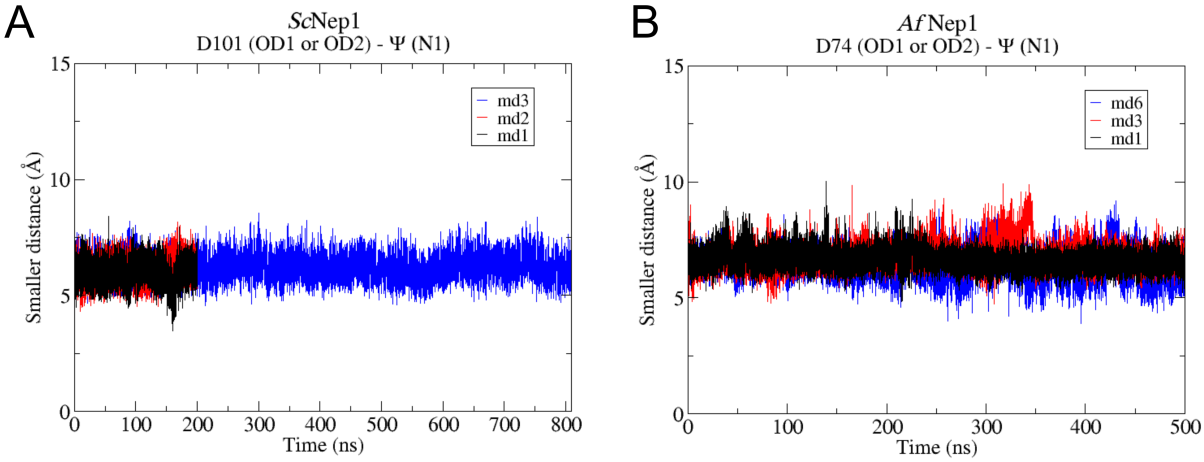


Figure S8: Variation of the distance between OD1 or OD2 of D101(D74) and N1 of Ψ (smaller of the two) during the MD simulations of (A) *Sc*Nep1 and (B) *Af*Nep1.

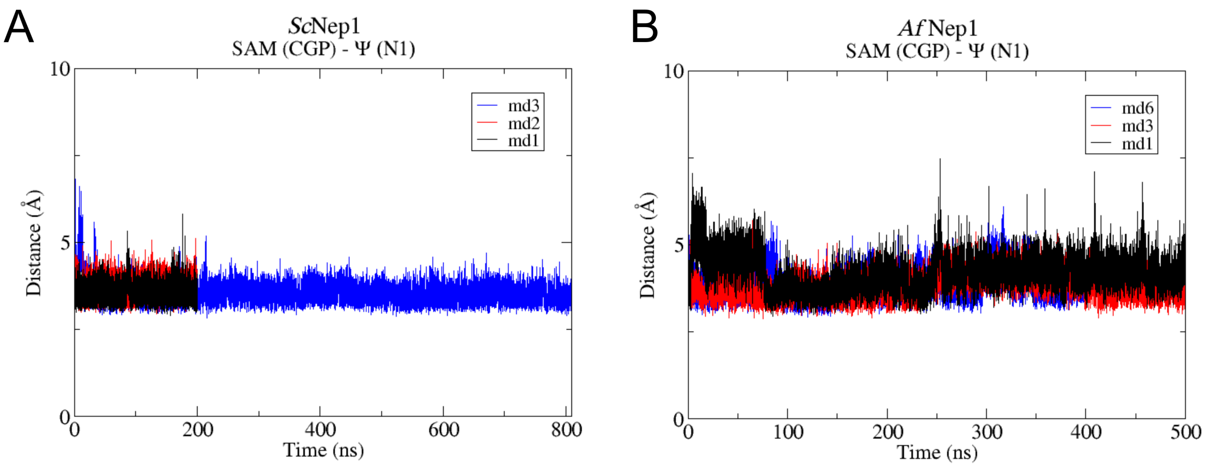


Figure S9: Variation of the distance between CGP of SAM and N1 of Ψ during the MD simulations of (A) *Sc*Nep1 and (B) *Af*Nep1.

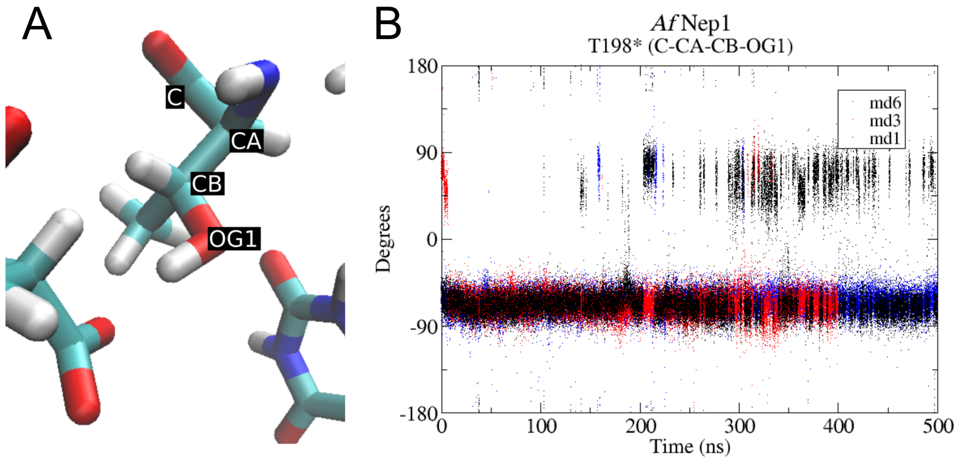


Figure S10: (A) Dihedral angle C-CA-CB-OG1 in T198* evaluated to identify MD snapshots suitable for the construction of an active-site model for the QM calculations (NEF atom names). For the threonine-mediated proton transfer to be possible, the dihedral angle needs to be larger than 0° or smaller than -120 °. (B) Variation of the dihedral angle C-CA-CB-OG1 in T198* during the MD simulations of *Af*Nep1.

### 3 QM calculations

Table S6: Composition of the investigated active–site models. Models were constructed based on MD snapshots, apart from Model 1, which was based on the crystal structure (PDB code: 3oin).

| | residues included | number of water molecules | number of atoms |
|---|---|---|---|
| ***S. cerevisiae*** | | | |
| Model 1 | D101, R132, R88*, D90*, I91*, Q94*, A208*, M209*, A210*, R211*, L232*, S233*, A234*, SAM, Ψ | 3 | 215 |
| Model 2 | D101, R132, R88*, D90*, I91*, A208*, M209*, A210*, R211*,L232*, S233*, A234*, SAM, Ψ | 1 | 194 |
| Model 3 | D101, R132, R88*, D90*, I91*, A208*, M209*, A210*, R211*,L232*, S233*, A234*, SAM, Ψ | 2 | 197 |
| Models 3' and 3" | D101, R132, R88*, D90*, A208*, M209*, A210*, R211*,L232*, S233*, A234*, SAM, Ψ | 3 | 181 |
| ***A. fulgidus*** | | | |
| Model 4 | L70, D74, R102, R61*, D63*, I64*, F171*, P172*, H173*, S196*, Y197*, T198*, S199*, SAM, Ψ | 4 | 240 |
| Models 5 and 5' | L70, L71, L73, D74, R102, R61*, D63*, I64*,F171*, F172*, H173*, S196*,Y197*, T198*, S199*, SAM, Ψ | 7 | 273 |

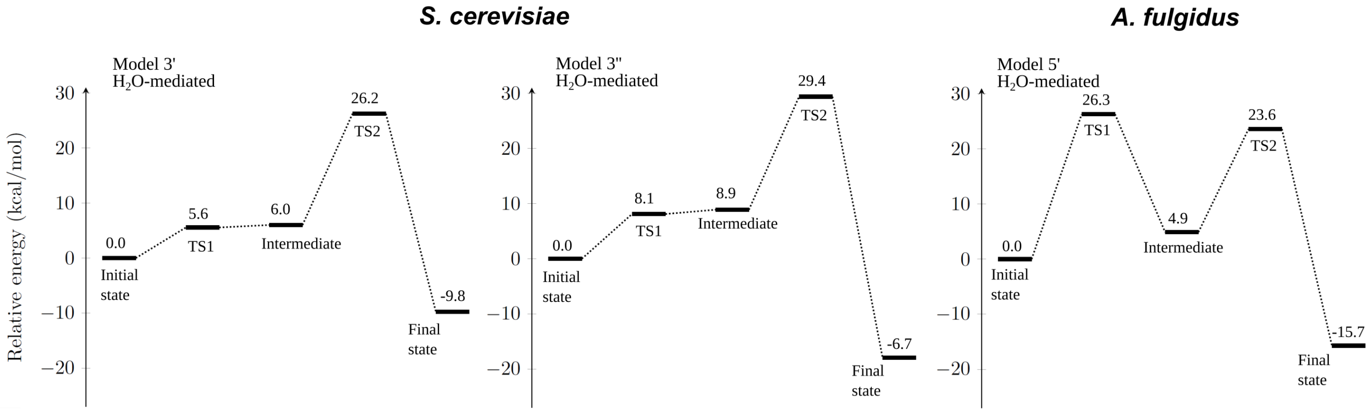


Figure S11: Energy profiles for models 3',3", and 5'.

Table S7: Energies of the stationary points for all the studied models.

**ΔE (kcal/mol)**

| ***Sc*Nep1** | TS1 | Intermediate | TS2 | Final state |
|---|---|---|---|---|
| **S233*-mediated proton transfer** | | | | |
| Model 1 (based on crystal structure) | 2.1 | 1.0 | 29.1 | –17.9 |

| | $TS1_1$ | $Intermediate_1$ | $TS1_2$ | $Intermediate_2$ | TS2 | Final state |
|---|---|---|---|---|---|---|
| Model 2 | –0.5 | 0.9 | 0.4 | –2.1 | 19.8 | –10.4 |

| **H2O-mediated proton transfer** | TS1 | Intermediate | TS2 | Final state |
|---|---|---|---|---|
| Model 3 | 4.0 | 2.1 | 18.5 | –16.4 |
| Model 3’ | 5.6 | 6.0 | 26.2 | –9.8 |
| Model 3” | 8.1 | 8.9 | 29.4 | –6.7 |

| ***Af*Nep1** | TS1 | Intermediate | TS2 | Final state |
|---|---|---|---|---|
| **T198*-mediated proton transfer** | | | | |
| Model 4 | 20.5 | 7.8 | 24.2 | –14.0 |
| **H2O-mediated proton transfer** | | | | |
| Model 5 | 15.8 | 0.9 | 19.1 | –17.9 |
| Model 5’ | 26.3 | 4.9 | 23.6 | –15.7 |

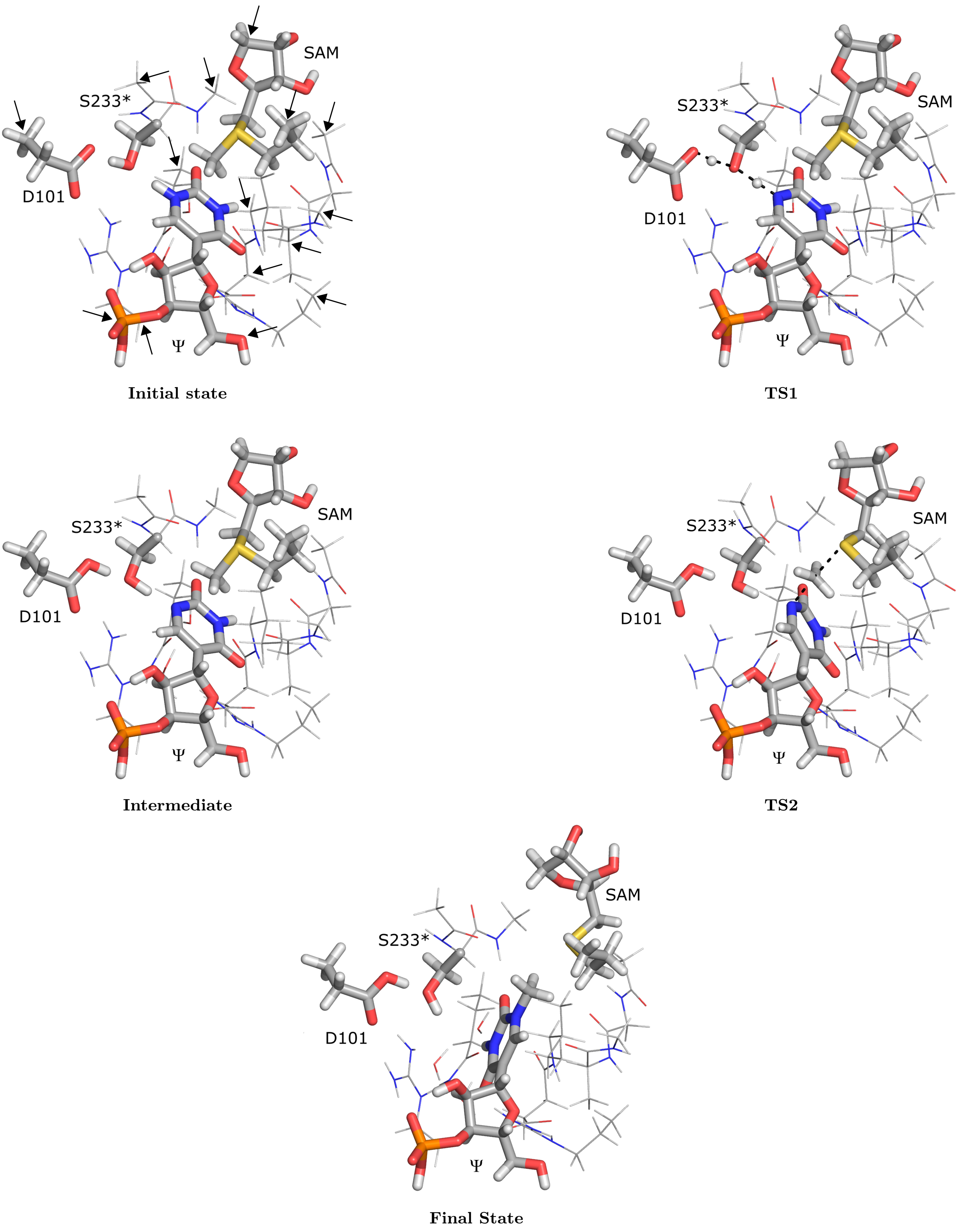


Figure S12: Optimized structures of the stationary points for Model 1 (arrows in the Initial state indicate atoms that were kept fixed during geometry optimizations).

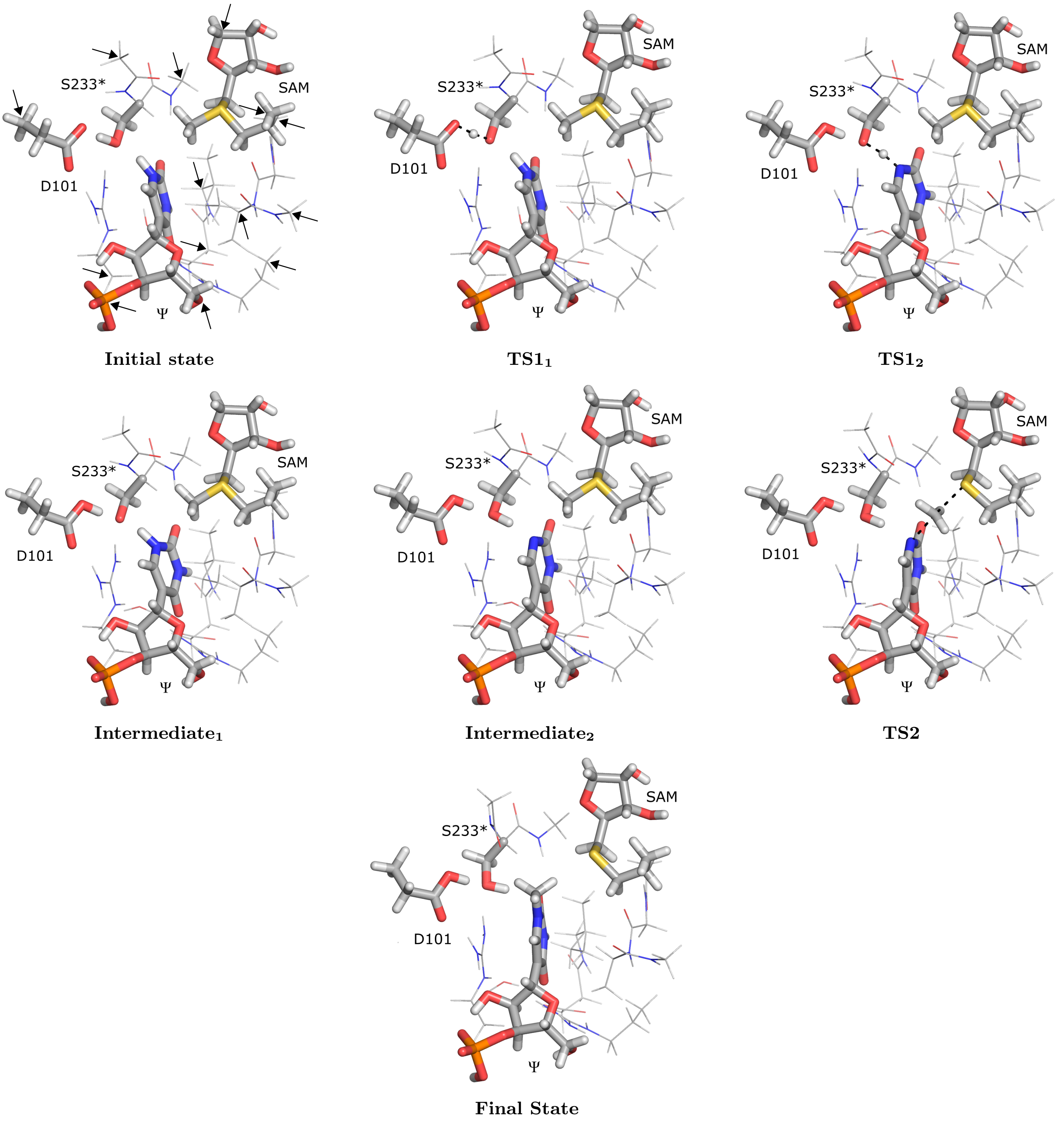


Figure S13: Optimized structures of the stationary points for Model 2 (arrows in the Initial state indicate atoms that were kept fixed during geometry optimizations).

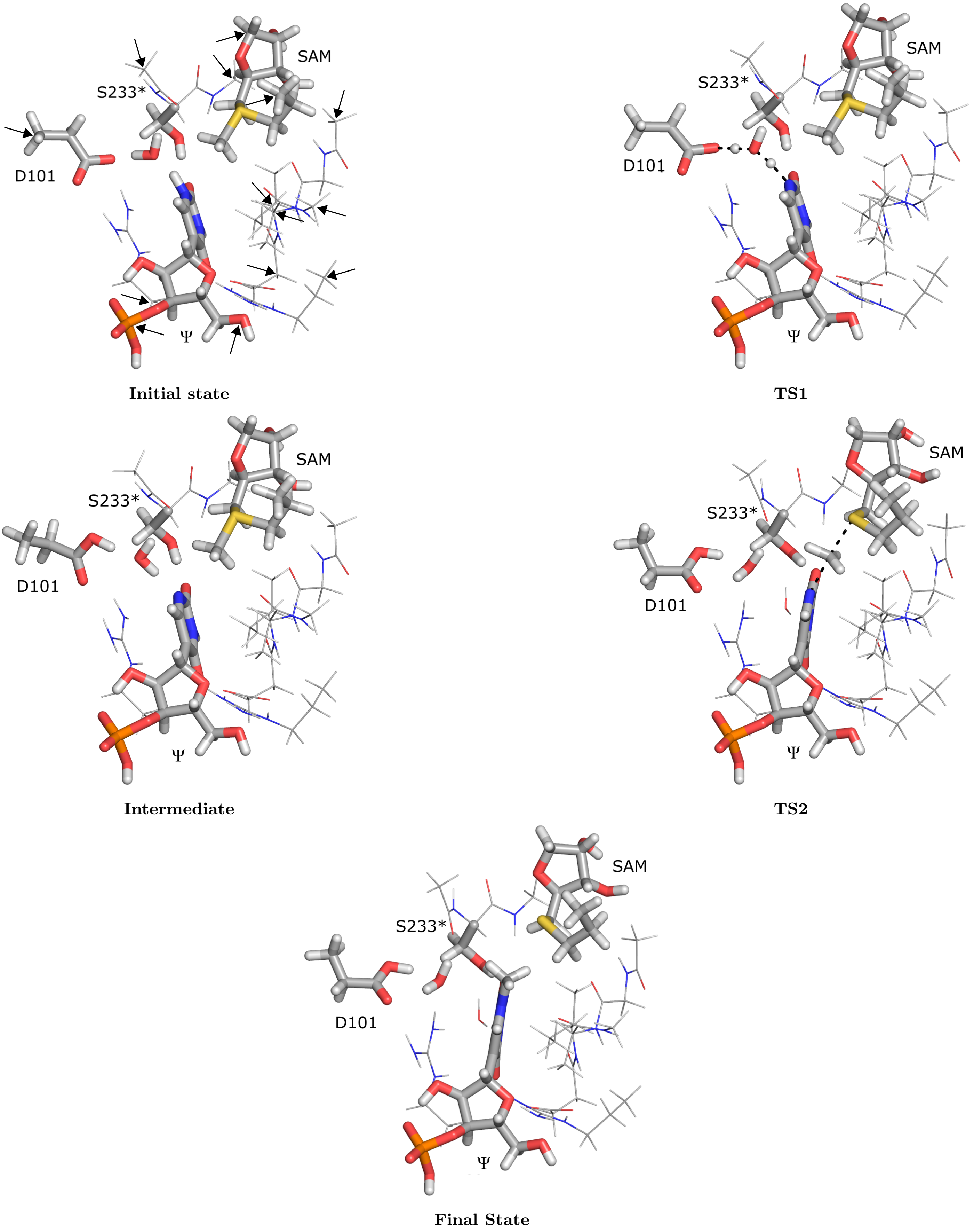


Figure S14: Optimized structures of the stationary points for Model 3 (arrows in the Initial state indicate atoms that were kept fixed during geometry optimizations).

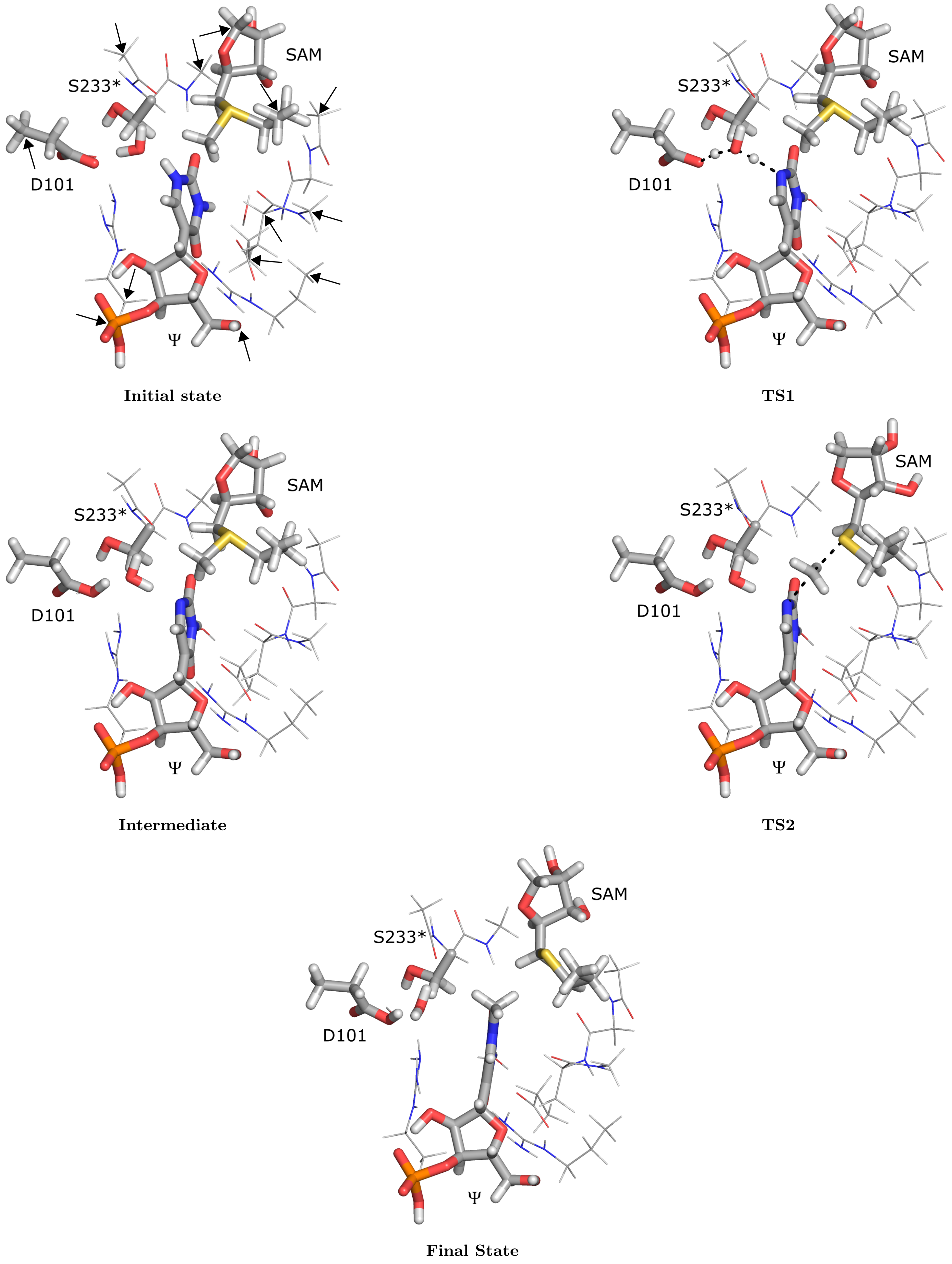


Figure S15: Optimized structures of the stationary points for Model 3' (arrows in the Initial state indicate atoms that were kept fixed during geometry optimizations).

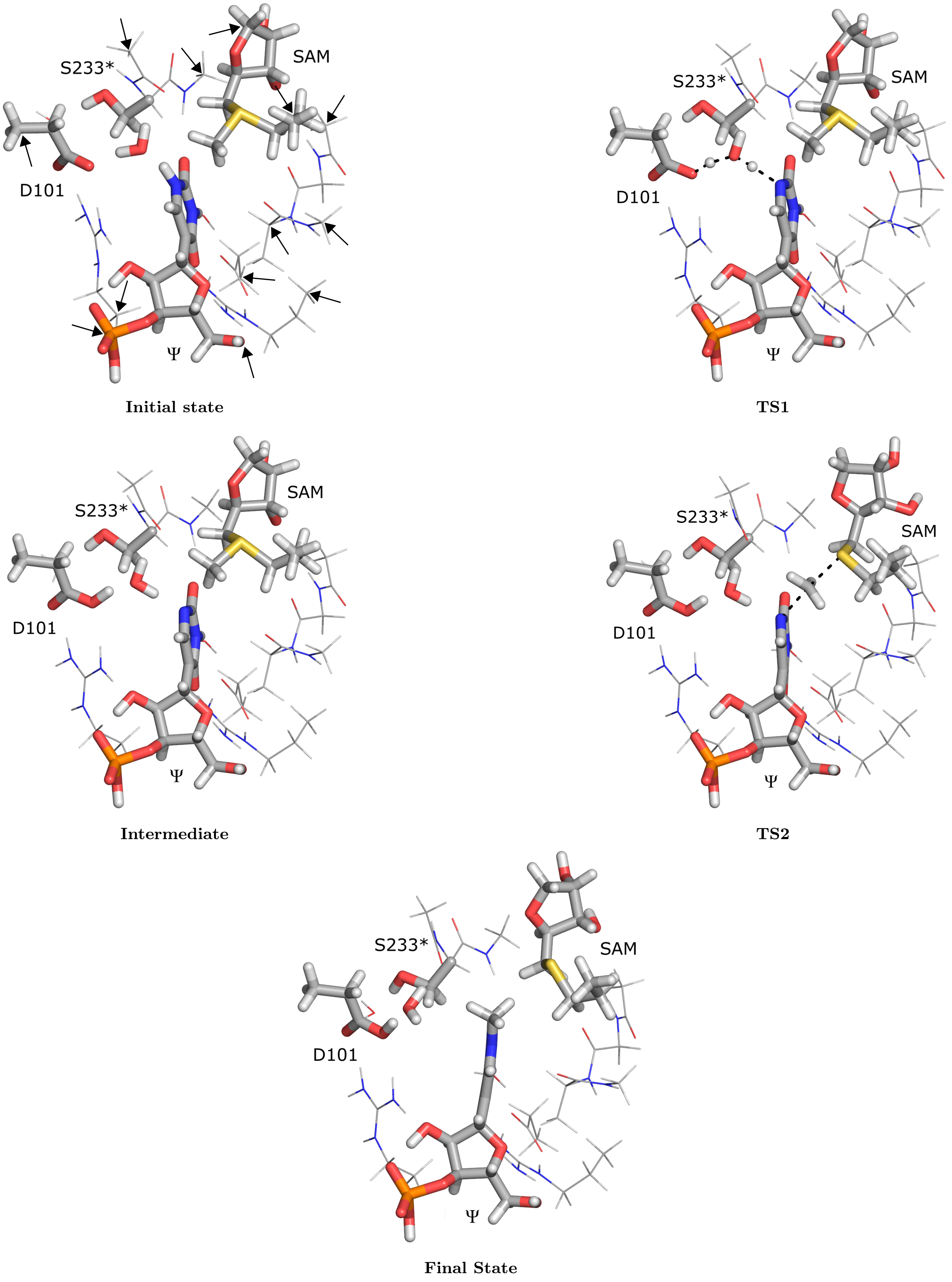


Figure S16: Optimized structures of the stationary points for Model 3" (arrows in the Initial state indicate atoms that were kept fixed during geometry optimizations).

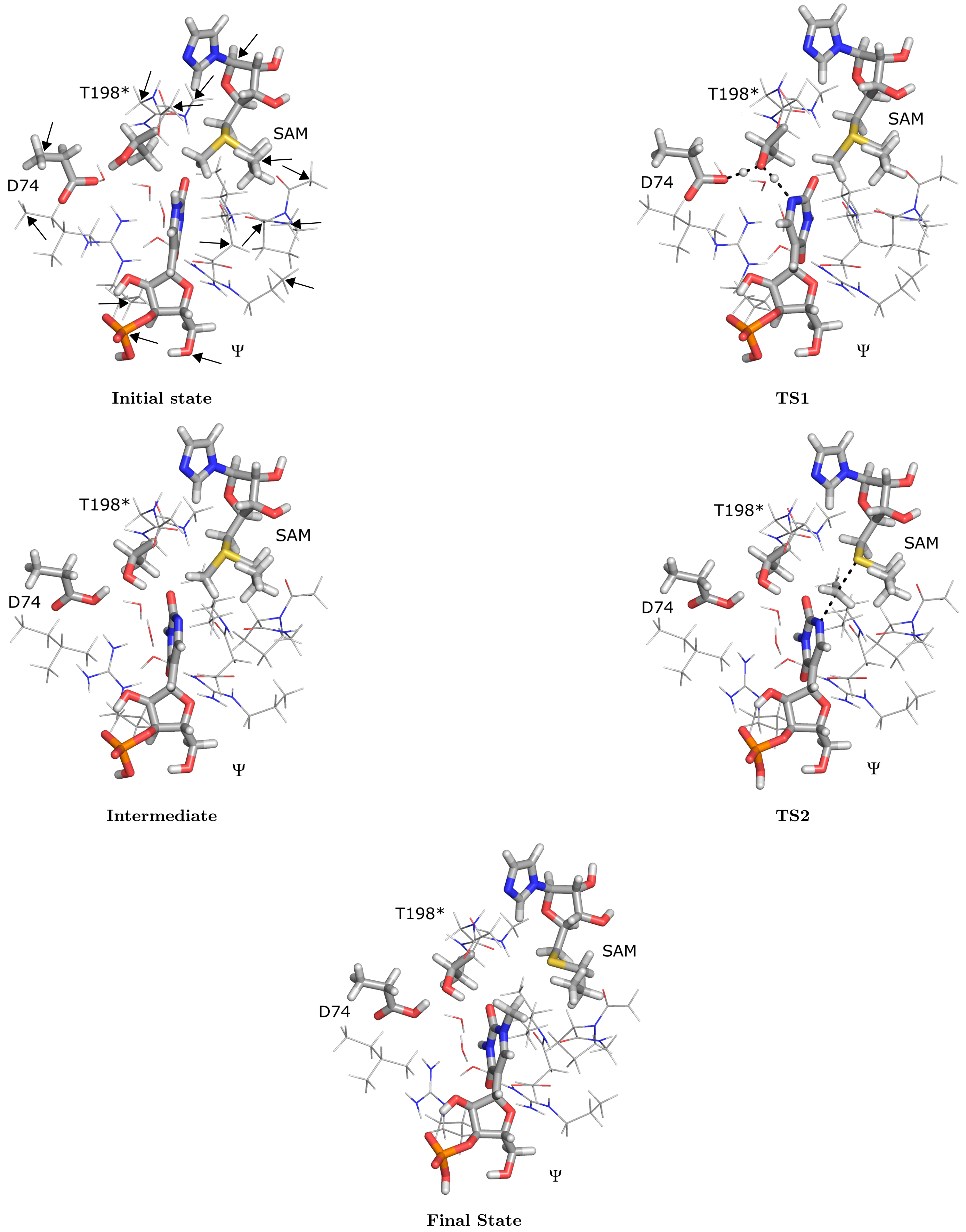


Figure S17: Optimized structures of the stationary points for Model 4 (arrows in the Initial state indicate atoms that were kept fixed during geometry optimizations).

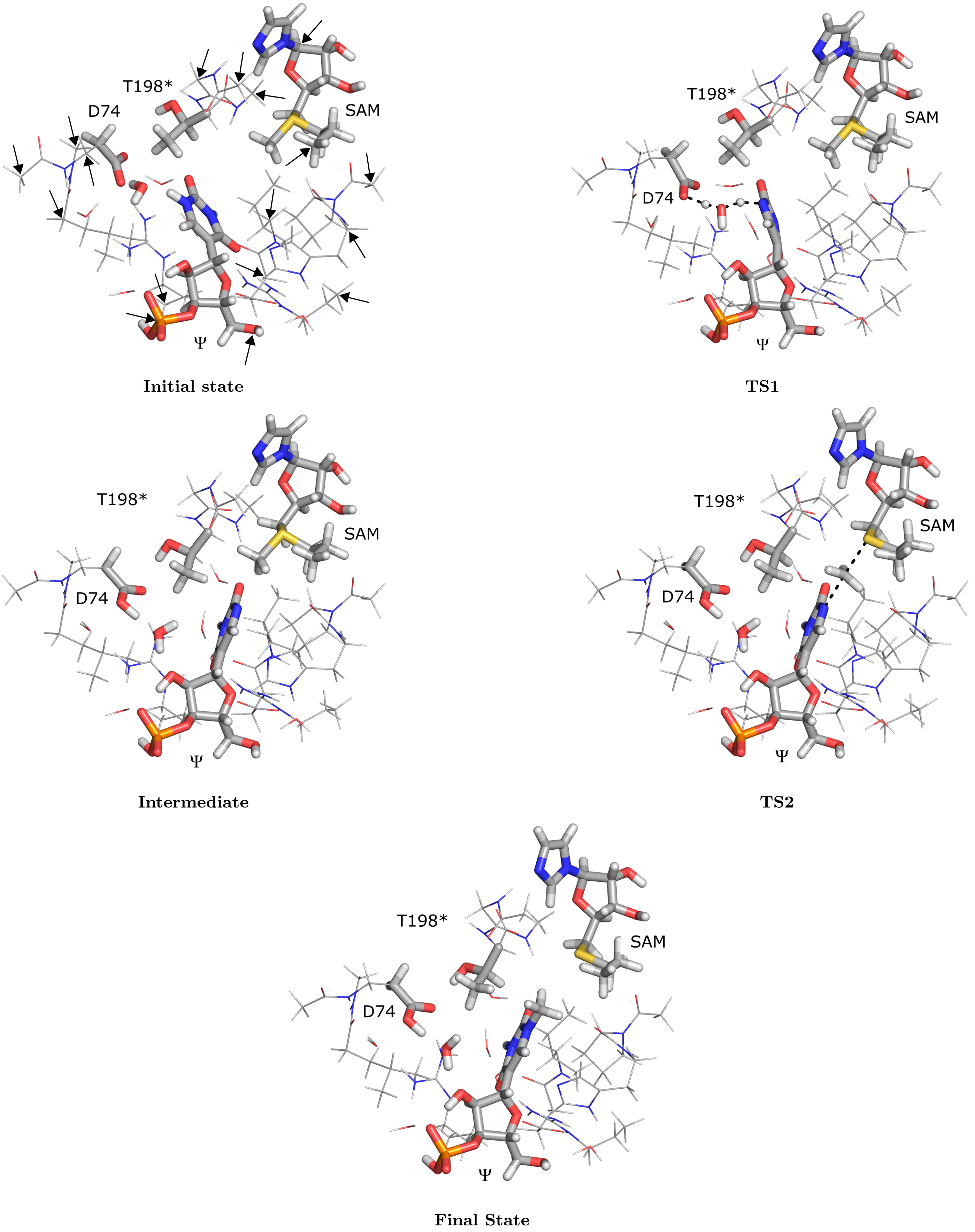


Figure S18: Optimized structures of the stationary points for Model 5 (arrows in the Initial state indicate atoms that were kept fixed during geometry optimizations).

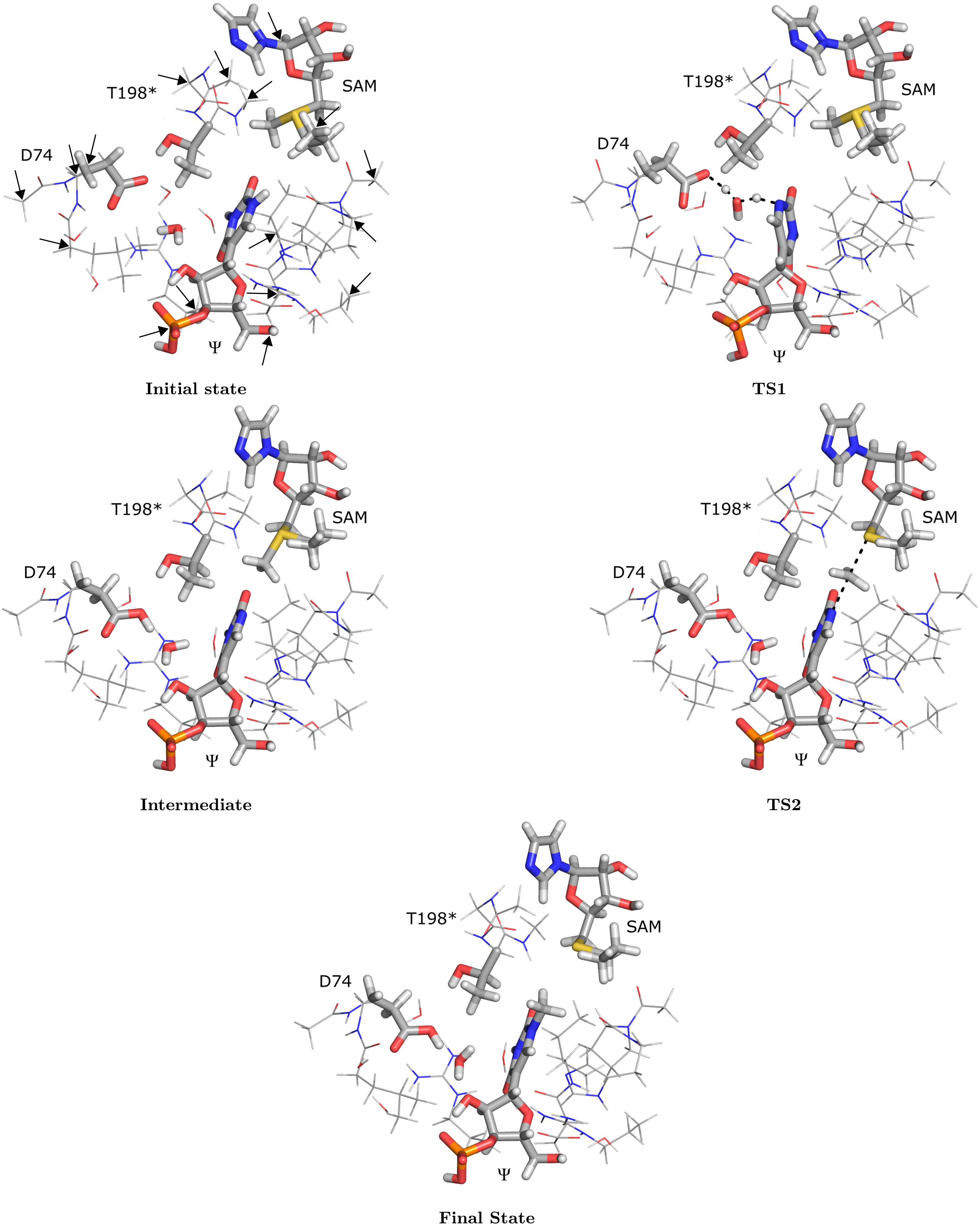


Figure S19: Optimized structures of the stationary points for Model 5' (arrows in the Initial state indicate atoms that were kept fixed during geometry optimizations).

Table S8: Calculated energies and energy corrections for all stationary points (in atomic units).

#### ***Sc*Nep1**

| **Model 1** | **E opt** | **E solv** | **ZPE** | **E bb** |
|---|---|---|---|---|
| Initial state | –5754.632930 | –5754.699131 | 1.821358 | –5761.009002 |
| TS1 | –5754.628208 | –5754.692167 | 1.816188 | –5761.002689 |
| Intermediate | –5754.635210 | –5754.698930 | 1.822151 | –5761.010766 |
| TS2 | –5754.595647 | –5754.656187 | 1.822688 | –5760.969567 |
| Final state | –5754.668626 | –5754.730645 | 1.821305 | –5761.041709 |

| **Model 2** | **E opt** | **E solv** | **ZPE** | **E bb** |
|---|---|---|---|---|
| Initial state | –5314.109674 | –5314.191598 | 1.627493 | –5319.947905 |
| TS1 1 | –5314.109666 | –5314.191305 | 1.626193 | –5319.947679 |
| Intermediate 1 | –5314.111963 | –5314.189405 | 1.627430 | –5319.950954 |
| TS1 2 | –5314.111761 | –5314.188547 | 1.625691 | –5319.950648 |
| Intermediate 2 | –5314.118901 | –5314.195057 | 1.630679 | –5319.960269 |
| TS2 | –5314.090760 | –5314.162613 | 1.629854 | –5319.928838 |
| Final state | –5314.143615 | –5314.214515 | 1.632009 | –5319.980023 |

| **Model 3** | **E opt** | **E solv** | **ZPE** | **E bb** |
|---|---|---|---|---|
| Initial state | –5390.506619 | –5390.588302 | 1.657054 | –5396.448275 |
| TS1 | –5390.497980 | –5390.578092 | 1.651046 | –5396.437461 |
| Intermediate | –5390.508292 | –5390.584394 | 1.657204 | –5396.450681 |
| TS2 | –5390.491623 | –5390.564523 | 1.657599 | –5396.428097 |
| Final state | –5390.556450 | –5390.627843 | 1.660409 | –5396.487970 |

| **Model 3’** | **E opt** | **E solv** | **ZPE** | **E bb** |
|---|---|---|---|---|
| Initial state | –5101.856604 | –5101.923633 | 1.513114 | –5107.475946 |
| TS1 | –5101.839082 | –5101.909323 | 1.506849 | –5107.457626 |
| Intermediate | –5101.841220 | –5101.911263 | 1.511230 | –5107.461479 |
| TS2 | –5101.811983 | –5101.879258 | 1.511305 | –5107.432106 |
| Final state | –5101.877043 | –5101.943153 | 1.513018 | –5107.492321 |

| **Model 3”** | **E opt** | **E solv** | **ZPE** | **E bb** |
|---|---|---|---|---|
| Initial state | –5101.855570 | –5101.925249 | 1.511516 | –5107.478232 |
| TS1 | –5101.837507 | –5101.908599 | 1.506512 | –5107.458900 |
| Intermediate | –5101.837848 | –5101.908942 | 1.509315 | –5107.460466 |
| TS2 | –5101.808945 | –5101.877148 | 1.509842 | –5107.431235 |
| Final state | –5101.872933 | –5101.939404 | 1.511907 | –5107.492488 |

#### ***Af*Nep1**

| **Model 4** | **E opt** | **E solv** | **ZPE** | **E bb** |
|---|---|---|---|---|
| Initial state | –6161.087830 | –6161.172395 | 2.044835 | –6167.934352 |
| TS1 | –6161.050308 | –6161.133888 | 2.037628 | –6167.895496 |
| Intermediate | –6161.077931 | –6161.159864 | 2.043398 | –6167.923171 |
| TS2 | –6161.062734 | –6161.133881 | 2.041087 | –6167.905447 |
| Final state | –6161.126387 | –6161.198620 | 2.043057 | –6167.967141 |

| **Model 5** | **E opt** | **E solv** | **ZPE** | **E bb** |
|---|---|---|---|---|
| Initial state | –7070.244846 | –7070.338342 | 21033 | –7078.176239 |
| TS1 | –7070.209393 | –7070.301701 | 14042 | –7078.145288 |
| Intermediate | –7070.249057 | –7070.332306 | 17142 | –7078.181235 |
| TS2 | –7070.227079 | –7070.307193 | 15900 | –7078.154085 |
| Final state | –7070.295339 | –7070.374109 | 19416 | –7078.217936 |

| **Model 5’** | **E opt** | **E solv** | **ZPE** | **E bb** |
|---|---|---|---|---|
| Initial state | –7070.214077 | –7070.315089 | 17133 | –7078.154759 |
| TS1 | –7070.167220 | –7070.271723 | 2.315476 | –7078.107720 |
| Intermediate | –7070.217629 | –7070.309713 | 2.315955 | –7078.154699 |
| TS2 | –7070.196693 | –7070.283957 | 2.314751 | –7078.128518 |
| Final state | –7070.266015 | –7070.353732 | 2.317698 | –7078.193653 |

### Cartesian coordinates for stationary points

#### Model1 – Initial state

| | | | |
|---|---|---|---|
| C | -2.18043200 | 6.70012000 | 6.40334900 |
| C | -2.13117200 | 6.92903500 | 4.88542100 |
| C | -1.66032900 | 5.68134300 | 4.13813400 |
| O | -0.54676100 | 5.69818500 | 3.56613700 |
| O | -2.43253200 | 4.66931700 | 4.17717900 |
| H | -1.45588400 | 7.76067900 | 4.63741000 |
| H | -1.17924200 | 6.45996400 | 6.79635500 |
| H | -3.14036700 | 7.18475400 | 4.52048300 |
| C | 4.99224900 | 3.58649400 | -0.25518500 |
| C | 4.19257800 | 4.58643200 | 0.57043200 |
| N | 3.07989600 | 3.92865300 | 1.25216800 |
| C | 2.05293400 | 4.55880700 | 1.83534000 |
| N | 1.20821500 | 3.85372100 | 2.60263300 |
| N | 1.84529500 | 5.87040700 | 1.67337600 |
| H | 1.03144500 | 6.24557700 | 2.16063600 |
| H | 4.34731100 | 3.05791400 | -0.97371300 |
| H | 3.81072500 | 5.37998500 | -0.08119100 |
| H | 3.07662800 | 2.90252000 | 1.32172900 |
| H | 1.45505500 | 2.91718000 | 2.89272600 |
| H | 2.12113000 | 6.31556900 | 0.77782800 |
| H | 5.76327100 | 4.12181900 | -0.82587700 |
| H | 4.85107700 | 5.06908400 | 1.31587000 |
| H | 0.42243100 | 4.35917700 | 3.05939400 |
| C | 3.96401000 | -4.36986700 | -3.56316200 |
| C | 4.09793100 | -3.08702100 | -4.37790100 |
| C | 5.23130700 | -2.19093200 | -3.86186100 |
| N | 5.03302500 | -1.75454500 | -2.49555100 |
| C | 4.26919600 | -0.70969200 | -2.13991100 |
| N | 3.74197800 | 0.11144400 | -3.06873700 |
| N | 4.09406300 | -0.48776700 | -0.84307300 |
| H | 3.54053000 | 0.27809800 | -0.45669300 |
| H | 3.81626200 | -4.15461000 | -2.49712100 |
| H | 3.15041600 | -2.52007700 | -4.34459300 |
| H | 5.37504500 | -1.31516100 | -4.51693300 |
| H | 5.29137400 | -2.42416900 | -1.73681100 |
| H | 3.70906100 | -0.18960900 | -4.03286600 |
| H | 4.72020900 | -1.00528400 | -0.19444100 |
| H | 3.11539000 | -4.98116500 | -3.90710500 |
| H | 4.28658700 | -3.32002600 | -5.44047700 |
| H | 6.18067600 | -2.74889700 | -3.89017900 |
| H | 2.98856100 | 0.76323100 | -2.82484400 |
| C | 6.33549000 | -5.31329600 | 1.77172100 |
| C | 4.91937400 | -5.33431400 | 2.34725900 |
| O | 4.70370800 | -5.53112100 | 3.54195000 |
| C | 6.81696100 | -3.86114600 | 1.59101400 |
| C | 6.06707200 | -3.04833300 | 0.52220000 |
| O | 5.61994500 | -3.64732600 | -0.49209200 |
| O | 5.96835100 | -1.80387400 | 0.70452600 |
| H | 7.87836200 | -3.86460500 | 1.28612400 |
| H | 6.98359900 | -5.84757800 | 2.47875400 |
| H | 6.76760300 | -3.30935300 | 2.54201800 |
| N | 3.93149400 | -5.04976600 | 1.46090400 |
| C | 2.57749200 | -4.74574100 | 1.90014900 |
| C | 1.57968800 | -4.64059100 | 0.74465300 |
| C | 1.97741600 | -3.55111700 | -0.26621800 |
| C | 0.18161000 | -4.37736100 | 1.31828300 |
| C | 1.12910700 | -3.56354100 | -1.53572400 |
| H | 0.06834300 | -3.35614800 | -1.32104100 |
| H | 4.23169300 | -4.75557400 | 0.53180800 |
| H | 2.57723000 | -3.79417400 | 2.46351500 |
| H | 1.56090800 | -5.61140300 | 0.21553700 |
| H | 3.03297100 | -3.66491300 | -0.55603200 |
| H | -0.10723700 | -5.15258500 | 2.04603500 |
| H | 1.17195000 | -4.55490600 | -2.01301800 |
| H | 1.93469400 | -2.56953300 | 0.23672200 |
| H | -0.58449800 | -4.35958000 | 0.52934900 |
| H | 1.46527600 | -2.79623200 | -2.24418900 |
| H | 0.15288200 | -3.40266900 | 1.83264300 |
| C | 3.34949700 | -4.87613600 | 7.04338600 |
| C | 3.36745000 | -3.65846100 | 6.12373400 |
| C | 4.78754100 | -3.31308900 | 5.68868500 |
| C | 4.87995500 | -2.13687300 | 4.75027100 |
| N | 6.06296300 | -1.49090100 | 4.70722600 |
| O | 3.93716700 | -1.78743400 | 4.02202800 |
| H | 6.84053800 | -1.78968100 | 5.27938100 |
| H | 3.76648200 | -5.75368700 | 6.52400400 |
| H | 2.90406700 | -2.78953000 | 6.62296200 |
| H | 5.19965100 | -4.17600100 | 5.13467400 |
| H | 6.17774400 | -0.73187100 | 4.02984200 |
| H | 2.77165500 | -3.86014100 | 5.22078600 |
| H | 5.43820800 | -3.13582800 | 6.56248900 |
| C | -2.57775300 | -6.55588400 | -0.14748500 |
| C | -1.79339500 | -6.11860900 | -1.37555100 |
| O | -0.79163600 | -6.70606200 | -1.74638800 |
| H | -1.88012900 | -6.65153000 | 0.69570800 |
| N | -2.30015300 | -5.04098600 | -2.05695300 |
| C | -1.69172600 | -4.55939100 | -3.26743100 |
| C | -1.67463700 | -3.03460300 | -3.33162200 |
| O | -2.40338100 | -2.35353000 | -2.61574100 |
| H | -3.09461300 | -4.52226000 | -1.70190300 |
| H | -0.67884100 | -4.98203700 | -3.32625900 |
| N | -0.87012900 | -2.50403700 | -4.29070200 |
| C | -0.69181300 | -1.05416300 | -4.42154000 |
| C | -1.97435200 | -0.36804200 | -4.88110100 |
| O | -2.31951900 | 0.71571000 | -4.43164400 |
| C | 0.49930100 | -0.72503400 | -5.31391600 |
| H | 0.35294400 | -1.07072900 | -6.35030700 |
| H | -0.15384400 | -3.10443500 | -4.68561600 |
| H | -0.48291400 | -0.64225700 | -3.43005200 |
| H | 1.40307500 | -1.19372900 | -4.89622200 |
| H | 0.67784500 | 0.35888300 | -5.31828100 |
| N | -2.65600700 | -1.04409100 | -5.84076500 |
| C | -3.81945400 | -0.51057500 | -6.49244900 |
| H | -2.29104500 | -1.94522800 | -6.12365800 |
| H | -4.07199500 | 0.44078000 | -6.00671900 |
| C | -6.62502600 | 3.55891000 | 3.07158700 |
| C | -5.54748500 | 2.58644900 | 2.62631300 |
| O | -5.71975200 | 1.80344800 | 1.68862900 |
| H | -6.20664900 | 4.46327700 | 3.53779400 |
| N | -4.40986900 | 2.56990200 | 3.36429900 |
| C | -3.49791600 | 1.45850100 | 3.24147700 |
| C | -4.20066700 | 0.14471500 | 3.61936300 |
| O | -5.18117600 | 0.08419400 | 4.33524300 |
| C | -2.21059400 | 1.66139900 | 4.05873700 |
| O | -1.34090800 | 2.55485300 | 3.41796400 |
| H | -1.72418500 | 3.50240700 | 3.62020400 |
| H | -4.16017200 | 3.34577300 | 3.97654400 |
| H | -3.17032800 | 1.35497700 | 2.19567600 |
| H | -1.70629300 | 0.68927800 | 4.15765200 |
| H | -2.46838100 | 2.01741000 | 5.07254600 |
| N | -3.60780000 | -0.95151700 | 3.05328200 |
| C | -4.06774900 | -2.27344300 | 3.39944400 |
| H | -2.66326400 | -0.85155100 | 2.67064900 |
| H | -5.16685600 | -2.27005400 | 3.42997600 |
| C | -7.21456700 | -0.02762100 | -3.72081100 |
| C | -6.01729900 | -0.94015600 | -3.44818900 |
| C | -4.86924400 | -0.28035400 | -2.69581900 |
| S | -5.31645700 | 0.33574800 | -1.01432400 |
| C | -5.01610600 | -1.12199100 | 0.06841600 |
| C | -6.25230500 | -1.87038200 | 0.53007500 |
| O | -7.00253400 | -1.11121600 | 1.44627400 |
| C | -7.28412900 | -2.31351000 | -0.51479400 |
| O | -6.87268100 | -3.37343200 | -1.32629500 |
| C | -8.48409000 | -2.65866400 | 0.39296200 |
| O | -8.38832000 | -4.02524900 | 0.77464600 |
| C | -8.32250900 | -1.65653900 | 1.54680100 |

| | | | |
|---|---|---|---|
| H | -7.69116200 | 0.31577000 | -2.78776800 |
| H | -5.57809100 | -1.26691600 | -4.40428600 |
| H | -4.50847100 | 0.62180300 | -3.21300800 |
| H | -4.50286400 | -0.71827700 | 0.95436800 |
| H | -5.84868700 | -2.79280400 | 1.00685400 |
| H | -7.54735900 | -1.44834100 | -1.14677800 |
| H | -7.20421700 | -4.17685400 | -0.88573400 |
| H | -9.44373100 | -2.55482300 | -0.13267200 |
| H | -7.86226000 | -4.09960300 | 1.58269900 |
| H | -8.46334500 | -2.15755200 | 2.52261800 |
| H | -6.32553100 | -1.86819600 | -2.94422600 |
| H | -4.00512700 | -0.94913300 | -2.57399800 |
| H | -4.33850600 | -1.78674600 | -0.48622000 |
| O | 1.82625500 | -1.11526100 | 2.39919200 |
| H | 0.90727900 | -1.14887800 | 2.71102500 |
| H | 2.40118700 | -1.46271600 | 3.10957900 |
| O | 5.22574300 | 0.04563600 | 2.43438200 |
| H | 5.66041600 | -0.58487200 | 1.80447000 |
| H | 4.60834700 | -0.55667800 | 2.90944900 |
| O | 3.01296700 | 1.10250500 | 1.27769500 |
| H | 3.90346100 | 0.87569000 | 1.65522900 |
| H | 2.45374000 | 0.38557300 | 1.65890400 |
| C | 3.35781200 | 3.11014800 | -4.74594900 |
| O | 3.06719900 | 2.38023700 | -5.92602300 |
| C | 2.12502100 | 3.32065200 | -3.90095700 |
| O | 1.71654300 | 2.14868400 | -3.16861800 |
| C | 2.35643800 | 4.35681600 | -2.80230800 |
| O | 2.24476800 | 5.66061900 | -3.29641100 |
| C | 1.32740400 | 3.95411300 | -1.70603000 |
| O | 0.30589600 | 4.86797800 | -1.45321700 |
| C | 0.73638900 | 2.64054900 | -2.24558500 |
| C | 0.52877600 | 0.25620200 | -1.18226100 |
| O | 1.06089700 | -0.39103800 | -2.07447800 |
| N | -0.00274000 | -0.42635700 | -0.07349900 |
| C | -0.52934100 | 0.11556700 | 1.07124900 |
| O | -0.91496300 | -0.59183300 | 2.01201300 |
| C | -0.16545400 | 2.23016000 | 0.01514900 |
| H | -0.28450800 | 3.31106900 | 0.12532300 |
| H | 4.13259100 | 2.60899300 | -4.13321600 |
| H | 1.29238200 | 3.65603300 | -4.54579600 |
| H | 3.37118500 | 4.19959400 | -2.40214600 |
| H | 1.89001100 | 3.72049400 | -0.78931900 |
| H | 0.78324100 | 5.70081500 | -1.19635600 |
| H | -0.17504900 | 2.89343700 | -2.82025600 |
| H | 0.13218900 | -1.43478200 | -0.07128900 |
| H | -0.90137300 | 1.95838800 | 1.97154000 |
| H | 3.71993300 | 4.12796200 | -4.98173000 |
| P | 2.81699900 | 6.88958800 | -2.31761700 |
| O | 4.40156700 | 6.42366600 | -2.12883000 |
| O | 2.16810100 | 6.64554100 | -0.95031300 |
| O | 2.74436300 | 8.16716500 | -3.06972500 |
| C | 0.36498000 | 1.69655500 | -1.12521000 |
| N | -0.59439300 | 1.47780400 | 1.07423200 |
| C | -3.81692600 | 1.29450400 | -0.67205600 |
| H | -3.94285100 | 1.64785800 | 0.35755200 |
| H | -3.80187500 | 2.13512200 | -1.37687900 |
| H | -2.93479800 | 0.65726700 | -0.81767800 |
| H | 4.92411800 | 6.97767100 | -2.72566200 |
| H | 3.81306600 | 2.47891700 | -6.52939800 |
| H | 6.36515400 | -5.82123600 | 0.79589900 |
| H | 2.26367100 | -5.52788500 | 2.60842500 |
| H | 2.32506000 | -5.12624000 | 7.36142200 |
| H | 3.94832600 | -4.70483400 | 7.95386200 |
| H | -3.71895000 | -2.99540500 | 2.64508800 |
| H | -3.70858800 | -2.60894900 | 4.38927800 |
| H | -7.25403500 | 3.82965100 | 2.21398500 |
| H | -7.24807900 | 3.04155000 | 3.81862800 |
| H | -9.04194500 | -0.82586900 | 1.48874000 |
| H | -2.54586700 | 7.59512600 | 6.93101300 |
| H | -2.84796600 | 5.85786300 | 6.64145100 |
| H | -2.23547900 | -4.91378100 | -4.16648400 |
| H | -3.38998100 | -5.86554600 | 0.12459300 |
| H | -4.68228200 | -1.19326300 | -6.41702400 |
| H | -3.63512000 | -0.31170700 | -7.56218700 |
| H | -7.98190100 | -0.55270700 | -4.30905000 |
| H | -6.91147700 | 0.87037900 | -4.28328700 |
| H | -3.00161000 | -7.55268700 | -0.34232600 |
| H | 4.87576800 | -4.98347900 | -3.64380200 |
| H | 5.47975100 | 2.82578200 | 0.37634000 |

#### Model1 – TS1

| | | | |
|---|---|---|---|
| C | -2.22092400 | 6.66269500 | 6.41857600 |
| C | -2.15509900 | 6.94751600 | 4.90987200 |
| C | -1.63737600 | 5.76171400 | 4.11437500 |
| O | -0.55258700 | 5.79005100 | 3.53486600 |
| O | -2.42314600 | 4.72123200 | 4.11915300 |
| H | -1.49666700 | 7.80206200 | 4.70051300 |
| H | -1.22230800 | 6.42624900 | 6.81871400 |
| H | -3.16366500 | 7.19115900 | 4.53602200 |
| C | 4.95675300 | 3.62846900 | -0.27126400 |
| C | 4.14639100 | 4.61941500 | 0.54591800 |
| N | 3.04803700 | 3.93482000 | 1.21834700 |
| C | 2.02555500 | 4.54207500 | 1.82574300 |
| N | 1.22785500 | 3.80787000 | 2.61391700 |
| N | 1.80246100 | 5.85274800 | 1.67685600 |
| H | 0.99321000 | 6.22237900 | 2.17173800 |
| H | 4.31901400 | 3.09477200 | -0.99262600 |
| H | 3.75348100 | 5.40633400 | -0.10689600 |
| H | 3.05739000 | 2.90805500 | 1.27642800 |
| H | 1.43243500 | 2.82962900 | 2.76644800 |
| H | 2.06362700 | 6.30080000 | 0.77726600 |
| H | 5.72865600 | 4.16504700 | -0.83975400 |
| H | 4.79257500 | 5.11187300 | 1.29587300 |
| H | 0.38587900 | 4.22532100 | 3.01890900 |
| C | 4.00342400 | -4.33966000 | -3.57337700 |
| C | 4.09928200 | -3.04867400 | -4.38051500 |
| C | 5.21691900 | -2.12957300 | -3.87057100 |
| N | 5.02037100 | -1.70024900 | -2.50179200 |
| C | 4.23597800 | -0.67290300 | -2.13827200 |
| N | 3.68533900 | 0.13952100 | -3.06124300 |
| N | 4.06615900 | -0.45680000 | -0.84000900 |
| H | 3.49561100 | 0.29444600 | -0.45003000 |
| H | 3.86252400 | -4.13487600 | -2.50426200 |
| H | 3.13943800 | -2.50412100 | -4.33302800 |
| H | 5.33665900 | -1.24955600 | -4.52478900 |
| H | 5.29897800 | -2.36603000 | -1.74721100 |
| H | 3.65690800 | -0.15856500 | -4.02644500 |
| H | 4.70098000 | -0.96772600 | -0.19550700 |
| H | 3.16503400 | -4.96760900 | -3.91241000 |
| H | 4.28192200 | -3.27027200 | -5.44662500 |
| H | 6.17790100 | -2.66682100 | -3.90778000 |
| H | 2.91069500 | 0.76395200 | -2.81120100 |
| C | 6.40900300 | -5.25523600 | 1.75108700 |
| C | 4.99577100 | -5.29322900 | 2.33291100 |
| O | 4.78806800 | -5.49074600 | 3.52904200 |
| C | 6.86906000 | -3.79578100 | 1.56859600 |
| C | 6.09821000 | -2.99191500 | 0.50704400 |
| O | 5.66510700 | -3.59375600 | -0.51145300 |
| O | 5.96854200 | -1.75195900 | 0.70014200 |
| H | 7.92786500 | -3.78437900 | 1.25498400 |
| H | 7.06745300 | -5.78068300 | 2.45516800 |
| H | 6.81887500 | -3.24657300 | 2.52104800 |
| N | 4.00074100 | -5.02062000 | 1.45108500 |
| C | 2.64544800 | -4.72995200 | 1.89609000 |
| C | 1.64391100 | -4.62666900 | 0.74319200 |
| C | 2.02737500 | -3.52553500 | -0.26086500 |
| C | 0.24417300 | -4.38192000 | 1.32121600 |
| C | 1.17840600 | -3.53833500 | -1.52969900 |
| H | 0.11504100 | -3.35135500 | -1.31098200 |
| H | 4.29223600 | -4.72502700 | 0.51973900 |

| | | | |
|---|---|---|---|
| H | 2.63937800 | -3.78138900 | 2.46446700 |
| H | 1.63467700 | -5.59439100 | 0.20776300 |
| H | 3.08426900 | -3.62418200 | -0.55178100 |
| H | -0.03357200 | -5.16251300 | 2.04768700 |
| H | 1.24040500 | -4.52232300 | -2.02058700 |
| H | 1.97043000 | -2.54816000 | 0.24812100 |
| H | -0.52357400 | -4.37123200 | 0.53369400 |
| H | 1.49904800 | -2.75488600 | -2.22773100 |
| H | 0.20593300 | -3.40866600 | 1.83750400 |
| C | 3.44166400 | -4.85058800 | 7.03588300 |
| C | 3.42823600 | -3.63033600 | 6.12038100 |
| C | 4.83752500 | -3.26093500 | 5.67169800 |
| C | 4.90082200 | -2.07882200 | 4.73851900 |
| N | 6.07721000 | -1.42119000 | 4.67967000 |
| O | 3.94136200 | -1.73466900 | 4.03021400 |
| H | 6.86861800 | -1.72103000 | 5.23189700 |
| H | 3.86820000 | -5.71900600 | 6.50893800 |
| H | 2.95589300 | -2.77066500 | 6.62715200 |
| H | 5.25775500 | -4.11507100 | 5.11014000 |
| H | 6.17486600 | -0.66104800 | 4.00073300 |
| H | 2.82574800 | -3.83902400 | 5.22348600 |
| H | 5.49460500 | -3.07704700 | 6.53941900 |
| C | -2.49812900 | -6.59840900 | -0.12836600 |
| C | -1.72977000 | -6.17043800 | -1.36997300 |
| O | -0.74590300 | -6.77459000 | -1.76167300 |
| H | -1.79273400 | -6.66655700 | 0.71132800 |
| N | -2.22683500 | -5.07982700 | -2.03621000 |
| C | -1.62256600 | -4.59837300 | -3.24901300 |
| C | -1.63802500 | -3.07427000 | -3.32142400 |
| O | -2.37778600 | -2.40629000 | -2.60470800 |
| H | -3.00317200 | -4.54383800 | -1.66618600 |
| H | -0.60075400 | -4.99983600 | -3.30287100 |
| N | -0.84950100 | -2.52995400 | -4.28605600 |
| C | -0.69313300 | -1.07671400 | -4.41164000 |
| C | -1.98742100 | -0.40705600 | -4.86187000 |
| O | -2.34506600 | 0.67196900 | -4.41014800 |
| C | 0.48674300 | -0.72636000 | -5.31095100 |
| H | 0.33740300 | -1.06551300 | -6.34912900 |
| H | -0.12191300 | -3.11684000 | -4.68069500 |
| H | -0.47987500 | -0.66596900 | -3.41959200 |
| H | 1.39923900 | -1.18755500 | -4.90389200 |
| H | 0.65191100 | 0.35966300 | -5.30671200 |
| N | -2.66530200 | -1.09005000 | -5.81973100 |
| C | -3.83587800 | -0.56877600 | -6.46871500 |
| H | -2.29104700 | -1.98709100 | -6.10367000 |
| H | -4.09058600 | 0.38548700 | -5.98976700 |
| C | -6.64461200 | 3.47091800 | 3.10701000 |
| C | -5.55795000 | 2.54086700 | 2.60094600 |
| O | -5.69416500 | 1.84680300 | 1.59242800 |
| H | -6.23067800 | 4.35512000 | 3.61417800 |
| N | -4.43712100 | 2.49194300 | 3.35983300 |
| C | -3.49748000 | 1.40724500 | 3.21954500 |
| C | -4.15030500 | 0.08570200 | 3.65022500 |
| O | -5.13488300 | 0.02836700 | 4.36578700 |
| C | -2.16500100 | 1.71530600 | 3.93207400 |
| O | -1.43798700 | 2.66374900 | 3.21670600 |
| H | -1.95943600 | 3.87054300 | 3.61009100 |
| H | -4.35853900 | 3.08921300 | 4.17494200 |
| H | -3.24058700 | 1.30389900 | 2.15731300 |
| H | -1.60331000 | 0.77075200 | 4.02869700 |
| H | -2.38060100 | 2.07021000 | 4.96039900 |
| N | -3.54747200 | -1.00980800 | 3.10257800 |
| C | -4.02051600 | -2.33234800 | 3.42442300 |
| H | -2.60955200 | -0.90114800 | 2.70053500 |
| H | -5.11800900 | -2.31021100 | 3.48870200 |
| C | -7.22387800 | -0.12345800 | -3.68213500 |
| C | -6.03074100 | -1.04336900 | -3.41783100 |
| C | -4.86804700 | -0.38694900 | -2.68552200 |
| S | -5.29170300 | 0.29635600 | -1.02400900 |
| C | -4.99292000 | -1.11533700 | 0.11856200 |

| | | | |
|---|---|---|---|
| C | -6.21647000 | -1.88056400 | 0.58529900 |
| O | -7.02206500 | -1.11457800 | 1.44855600 |
| C | -7.20852100 | -2.40664100 | -0.45825700 |
| O | -6.73456400 | -3.46844400 | -1.23231700 |
| C | -8.40443300 | -2.78464400 | 0.44386600 |
| O | -8.26871000 | -4.14238400 | 0.84278600 |
| C | -8.29008300 | -1.76363100 | 1.59059400 |
| H | -7.69078100 | 0.22412100 | -2.74577700 |
| H | -5.60567900 | -1.37975200 | -4.37721800 |
| H | -4.49013000 | 0.49173300 | -3.23034800 |
| H | -4.50241600 | -0.66947900 | 0.99559500 |
| H | -5.79335900 | -2.76538200 | 1.11303100 |
| H | -7.50419500 | -1.57262000 | -1.11711600 |
| H | -7.05528200 | -4.27375200 | -0.78699000 |
| H | -9.36158100 | -2.72042600 | -0.09226100 |
| H | -7.73333200 | -4.19055500 | 1.64690200 |
| H | -8.35974300 | -2.27409300 | 2.57000700 |
| H | -6.33802100 | -1.96594000 | -2.90407000 |
| H | -4.01592100 | -1.06782200 | -2.54818600 |
| H | -4.29120000 | -1.78670300 | -0.39677200 |
| O | 1.81093400 | -1.11190100 | 2.38532500 |
| H | 0.88046000 | -1.11599900 | 2.67059200 |
| H | 2.36416800 | -1.44364300 | 3.11852700 |
| O | 5.19884900 | 0.09501700 | 2.42237300 |
| H | 5.63722400 | -0.53384100 | 1.79340200 |
| H | 4.58944700 | -0.51068700 | 2.90452300 |
| O | 2.96525000 | 1.12048100 | 1.28963300 |
| H | 3.85950200 | 0.90232800 | 1.66346800 |
| H | 2.41814800 | 0.38408100 | 1.65411200 |
| C | 3.28829200 | 3.12304500 | -4.74184900 |
| O | 3.02027400 | 2.39944200 | -5.93334600 |
| C | 2.04581600 | 3.33311600 | -3.90711700 |
| O | 1.63163400 | 2.16622800 | -3.17174200 |
| C | 2.27249100 | 4.37264800 | -2.80904400 |
| O | 2.16947900 | 5.67746600 | -3.30500900 |
| C | 1.23356000 | 3.97884200 | -1.71890200 |
| O | 0.21206200 | 4.89913300 | -1.48659900 |
| C | 0.64626600 | 2.66062400 | -2.25115300 |
| C | 0.48325400 | 0.28945600 | -1.16195900 |
| O | 1.02948500 | -0.36209500 | -2.04772500 |
| N | -0.01871600 | -0.38290900 | -0.03842700 |
| C | -0.56562900 | 0.16991500 | 1.09621400 |
| O | -0.90878200 | -0.55823800 | 2.04657900 |
| C | -0.28495600 | 2.24949400 | 0.00948500 |
| H | -0.43559500 | 3.33010600 | 0.08891300 |
| H | 4.05471400 | 2.61929100 | -4.12109700 |
| H | 1.21843600 | 3.66865700 | -4.55894000 |
| H | 3.28391100 | 4.21179500 | -2.40146400 |
| H | 1.78705300 | 3.75492800 | -0.79441900 |
| H | 0.69016800 | 5.72672800 | -1.21965400 |
| H | -0.26259500 | 2.90972000 | -2.83141800 |
| H | 0.15199200 | -1.38531600 | -0.01251100 |
| H | -1.05460300 | 2.08608000 | 2.07144500 |
| H | 3.65338300 | 4.14205000 | -4.96772300 |
| P | 2.73537800 | 6.90647500 | -2.32460300 |
| O | 4.32415500 | 6.45195300 | -2.14152600 |
| O | 2.09399200 | 6.65756400 | -0.95366700 |
| O | 2.65157900 | 8.18626500 | -3.07235700 |
| C | 0.27728600 | 1.72080200 | -1.12506400 |
| N | -0.69592400 | 1.52181800 | 1.08814300 |
| C | -3.77207600 | 1.24543500 | -0.73826900 |
| H | -3.88372600 | 1.66313800 | 0.26801300 |
| H | -3.74079100 | 2.04021800 | -1.49369000 |
| H | -2.90258600 | 0.58428100 | -0.84722500 |
| H | 4.83943000 | 7.00406100 | -2.74631500 |
| H | 3.77606000 | 2.50844000 | -6.52232700 |
| H | 6.44087300 | -5.76225400 | 0.77484200 |
| H | 2.33961000 | -5.51848900 | 2.60084600 |
| H | 2.42549900 | -5.11999600 | 7.36468500 |
| H | 4.04772400 | -4.67177400 | 7.94017500 |

| | | | |
|---|---|---|---|
| H | -3.70619500 | -3.03992300 | 2.64147100 |
| H | -3.63928000 | -2.70340200 | 4.39328700 |
| H | -7.28156500 | 3.78187700 | 2.26926100 |
| H | -7.25961200 | 2.91078100 | 3.82973800 |
| H | -9.07359200 | -0.99247400 | 1.55645300 |
| H | -2.60650100 | 7.53758300 | 6.96372200 |
| H | -2.88115200 | 5.80678300 | 6.62461100 |
| H | -2.15544300 | -4.97089200 | -4.14734400 |
| H | -3.31925300 | -5.91637800 | 0.13789800 |
| H | -4.69507000 | -1.25491000 | -6.38203700 |
| H | -3.65896600 | -0.37860900 | -7.54126400 |
| H | -7.99937400 | -0.64359800 | -4.26416100 |
| H | -6.91922900 | 0.77205400 | -4.24784300 |
| H | -2.90563900 | -7.60568000 | -0.30179300 |
| H | 4.92786800 | -4.93195300 | -3.66822900 |
| H | 5.44493100 | 2.87138100 | 0.36430300 |

#### Model1 – Intermediate

| | | | |
|---|---|---|---|
| C | -2.21977000 | 6.72022400 | 6.36377600 |
| C | -2.09935900 | 7.09482200 | 4.87973400 |
| C | -1.67046900 | 5.91434600 | 4.03847100 |
| O | -0.61766900 | 5.87308700 | 3.41950300 |
| O | -2.53119200 | 4.91429900 | 4.06304700 |
| H | -1.36494700 | 7.89727600 | 4.72717700 |
| H | -1.25635100 | 6.36233900 | 6.75965900 |
| H | -3.07484800 | 7.44579400 | 4.50328700 |
| C | 4.98567800 | 3.60896600 | -0.26027900 |
| C | 4.16714600 | 4.61228100 | 0.53197700 |
| N | 3.04889800 | 3.94222200 | 1.18609000 |
| C | 2.02663100 | 4.56478700 | 1.77202100 |
| N | 1.20361300 | 3.84067500 | 2.54734500 |
| N | 1.83487400 | 5.88144600 | 1.62715000 |
| H | 1.01701500 | 6.26318100 | 2.09530700 |
| H | 4.35898100 | 3.07636900 | -0.99186400 |
| H | 3.79528800 | 5.39748600 | -0.13435600 |
| H | 3.03636400 | 2.91363500 | 1.23550800 |
| H | 1.36880000 | 2.84907700 | 2.66089200 |
| H | 2.10409400 | 6.31606200 | 0.71994900 |
| H | 5.77464700 | 4.13473100 | -0.81532400 |
| H | 4.79991400 | 5.10447900 | 1.29343400 |
| H | 0.35334100 | 4.26254500 | 2.91815200 |
| C | 4.02613800 | -4.38319400 | -3.50210300 |
| C | 4.12481000 | -3.09954900 | -4.32065700 |
| C | 5.24358900 | -2.17720000 | -3.81862000 |
| N | 5.04472900 | -1.73005100 | -2.45587100 |
| C | 4.25065000 | -0.70359300 | -2.10825200 |
| N | 3.70667000 | 0.10102400 | -3.04233700 |
| N | 4.06773700 | -0.47671300 | -0.81416700 |
| H | 3.48063200 | 0.26998100 | -0.43973500 |
| H | 3.88556900 | -4.16851600 | -2.43485900 |
| H | 3.16595900 | -2.55299500 | -4.27825600 |
| H | 5.36728000 | -1.30576200 | -4.48355000 |
| H | 5.31775700 | -2.38833400 | -1.69380400 |
| H | 3.69133200 | -0.20574800 | -4.00517200 |
| H | 4.69060700 | -0.98519100 | -0.15814000 |
| H | 3.18654800 | -5.01231000 | -3.83599700 |
| H | 4.30773700 | -3.33116200 | -5.38465400 |
| H | 6.20339500 | -2.71711400 | -3.84674100 |
| H | 2.90738900 | 0.69878500 | -2.80259300 |
| C | 6.39897500 | -5.26222800 | 1.84318100 |
| C | 4.98225900 | -5.29524600 | 2.41700900 |
| O | 4.76725300 | -5.48694600 | 3.61292600 |
| C | 6.85718500 | -3.80276500 | 1.65201800 |
| C | 6.09332300 | -3.00825200 | 0.57729900 |
| O | 5.67371700 | -3.61818500 | -0.44175700 |
| O | 5.95676800 | -1.76776300 | 0.76187500 |
| H | 7.91860200 | -3.79124100 | 1.34734200 |
| H | 7.05370400 | -5.78117400 | 2.55547400 |
| H | 6.79726500 | -3.24600800 | 2.59962300 |
| N | 3.99350500 | -5.02280800 | 1.52831800 |
| C | 2.63631300 | -4.72419700 | 1.96265700 |
| C | 1.64408700 | -4.62806600 | 0.80147200 |
| C | 2.03499600 | -3.53211100 | -0.20533500 |
| C | 0.23879700 | -4.38081400 | 1.36512600 |
| C | 1.19805200 | -3.55410200 | -1.48147700 |
| H | 0.13123800 | -3.38090400 | -1.26987300 |
| H | 4.29087300 | -4.73477800 | 0.59642100 |
| H | 2.62998700 | -3.77096900 | 2.52364500 |
| H | 1.64205900 | -5.59973400 | 0.27248000 |
| H | 3.09509800 | -3.62890200 | -0.48525300 |
| H | -0.04107500 | -5.15113000 | 2.10183800 |
| H | 1.27779800 | -4.53594800 | -1.97486000 |
| H | 1.96722400 | -2.55214000 | 0.29681000 |
| H | -0.52236000 | -4.38605300 | 0.57119200 |
| H | 1.51346100 | -2.76330600 | -2.17349600 |
| H | 0.19198800 | -3.39880100 | 1.86346800 |
| C | 3.40333200 | -4.80490100 | 7.10767900 |
| C | 3.36179100 | -3.59405000 | 6.18114400 |
| C | 4.76170000 | -3.20854000 | 5.71830200 |
| C | 4.80273100 | -2.03253300 | 4.77615500 |
| N | 5.96714800 | -1.35305700 | 4.71830500 |
| O | 3.83839300 | -1.71408400 | 4.06263000 |
| H | 6.76322500 | -1.64100200 | 5.27009200 |
| H | 3.84018300 | -5.67077000 | 6.58500700 |
| H | 2.88063300 | -2.73691700 | 6.68412400 |
| H | 5.18956300 | -4.06176000 | 5.16095400 |
| H | 6.05661000 | -0.60357800 | 4.02580300 |
| H | 2.75473500 | -3.81978200 | 5.29145400 |
| H | 5.42226800 | -3.00980700 | 6.58010500 |
| C | -2.50162900 | -6.59348800 | -0.07528400 |
| C | -1.74990600 | -6.21955100 | -1.34472900 |
| O | -0.83336900 | -6.89670700 | -1.77799400 |
| H | -1.77743100 | -6.67112600 | 0.74776900 |
| N | -2.17775300 | -5.08505200 | -1.98232500 |
| C | -1.56696400 | -4.63085800 | -3.20255400 |
| C | -1.59347800 | -3.10849200 | -3.29610600 |
| O | -2.33984100 | -2.43936600 | -2.58608000 |
| H | -2.89552100 | -4.48931200 | -1.58587700 |
| H | -0.54222500 | -5.02615200 | -3.24897300 |
| N | -0.80840900 | -2.56682600 | -4.26345100 |
| C | -0.65550800 | -1.11291300 | -4.39370500 |
| C | -1.94983400 | -0.44826500 | -4.85002000 |
| O | -2.31422900 | 0.63055700 | -4.40274300 |
| C | 0.52576100 | -0.76279000 | -5.29121700 |
| H | 0.37873200 | -1.10079200 | -6.33018900 |
| H | -0.07599000 | -3.15217300 | -4.65146000 |
| H | -0.44118700 | -0.69946600 | -3.40273500 |
| H | 1.43671000 | -1.22448600 | -4.88145700 |
| H | 0.69221600 | 0.32301800 | -5.28348300 |
| N | -2.62196500 | -1.13434600 | -5.81039600 |
| C | -3.78500300 | -0.61224800 | -6.47244000 |
| H | -2.24355400 | -2.03036200 | -6.09214700 |
| H | -4.03257300 | 0.35165600 | -6.00923300 |
| C | -6.63474000 | 3.51480100 | 3.05373600 |
| C | -5.51835200 | 2.61688500 | 2.55317600 |
| O | -5.58294100 | 1.99510700 | 1.49583300 |
| H | -6.24445700 | 4.38935400 | 3.59524300 |
| N | -4.43903100 | 2.52935400 | 3.37788500 |
| C | -3.50083700 | 1.44351500 | 3.23058700 |
| C | -4.15599700 | 0.11764600 | 3.64779300 |
| O | -5.15095300 | 0.07246200 | 4.35287700 |
| C | -2.18124700 | 1.72418100 | 3.95916800 |
| O | -1.44953400 | 2.72251400 | 3.28446700 |
| H | -2.13089900 | 4.10919600 | 3.58970400 |
| H | -4.50705100 | 2.96073300 | 4.29249600 |
| H | -3.23980400 | 1.36091800 | 2.16880100 |
| H | -1.59943200 | 0.79169400 | 4.00763200 |
| H | -2.37351000 | 2.06533800 | 4.99087700 |
| N | -3.54686400 | -0.97398600 | 3.11539000 |
| C | -4.03059900 | -2.29351300 | 3.43367100 |

| | | | |
|---|---|---|---|
| H | -2.61919100 | -0.86358600 | 2.67738100 |
| H | -5.12971400 | -2.27417700 | 3.46806200 |
| C | -7.18719900 | -0.13359700 | -3.70875700 |
| C | -6.02580800 | -1.09305500 | -3.44813200 |
| C | -4.83778400 | -0.47184900 | -2.72729100 |
| S | -5.22595800 | 0.25356400 | -1.07559800 |
| C | -4.97594500 | -1.14500600 | 0.09521900 |
| C | -6.21786500 | -1.87860400 | 0.56544000 |
| O | -7.01537400 | -1.08695600 | 1.41523000 |
| C | -7.21643200 | -2.39810100 | -0.47514800 |
| O | -6.75933500 | -3.47268600 | -1.24032000 |
| C | -8.42056500 | -2.74951000 | 0.42734400 |
| O | -8.31513300 | -4.10790300 | 0.83151500 |
| C | -8.28804400 | -1.72695200 | 1.57120300 |
| H | -7.63629000 | 0.23304200 | -2.77091600 |
| H | -5.61867900 | -1.44759000 | -4.40897900 |
| H | -4.42872900 | 0.38600000 | -3.28307600 |
| H | -4.48046500 | -0.69358900 | 0.96664900 |
| H | -5.81572600 | -2.76417500 | 1.10820100 |
| H | -7.49687400 | -1.56360700 | -1.14004300 |
| H | -7.09835400 | -4.27023200 | -0.79476900 |
| H | -9.37526600 | -2.66775700 | -0.11060600 |
| H | -7.77002400 | -4.16635100 | 1.62843900 |
| H | -8.35064100 | -2.23597300 | 2.55187200 |
| H | -6.36057100 | -2.00237100 | -2.92840000 |
| H | -4.00883000 | -1.17871500 | -2.58142100 |
| H | -4.28225800 | -1.83955100 | -0.39967600 |
| O | 1.71929600 | -1.06160700 | 2.36178700 |
| H | 0.76728600 | -1.05905200 | 2.57876000 |
| H | 2.22304600 | -1.39728700 | 3.12590700 |
| O | 5.10897300 | 0.08759900 | 2.43431900 |
| H | 5.57025200 | -0.54499700 | 1.82534400 |
| H | 4.48897400 | -0.51574800 | 2.90663300 |
| O | 2.91466300 | 1.14256000 | 1.26673200 |
| H | 3.79668900 | 0.90828800 | 1.65863700 |
| H | 2.34622600 | 0.41637900 | 1.62029700 |
| C | 3.33737500 | 3.07196400 | -4.73577000 |
| O | 3.07725200 | 2.33919300 | -5.92298100 |
| C | 2.08318000 | 3.30051800 | -3.92544500 |
| O | 1.63866500 | 2.13886100 | -3.20582000 |
| C | 2.30232800 | 4.33602500 | -2.82195800 |
| O | 2.22099500 | 5.64232600 | -3.32371500 |
| C | 1.23721000 | 3.95110600 | -1.75306200 |
| O | 0.22908900 | 4.89216000 | -1.54181700 |
| C | 0.64079400 | 2.64048300 | -2.30152200 |
| C | 0.48115200 | 0.27597100 | -1.19396600 |
| O | 1.07766100 | -0.38590500 | -2.04719200 |
| N | -0.04462700 | -0.38283600 | -0.07599200 |
| C | -0.64803600 | 0.19353200 | 1.02350300 |
| O | -0.98539700 | -0.54646700 | 1.98502600 |
| C | -0.39658300 | 2.22137500 | -0.08425500 |
| H | -0.56842500 | 3.30245100 | -0.05056800 |
| H | 4.08711600 | 2.56436000 | -4.09822200 |
| H | 1.27518300 | 3.65255700 | -4.59321100 |
| H | 3.30492200 | 4.16682400 | -2.39663200 |
| H | 1.76716000 | 3.71541900 | -0.81780200 |
| H | 0.71468400 | 5.71219100 | -1.27111100 |
| H | -0.25046300 | 2.90936400 | -2.90039800 |
| H | 0.15267900 | -1.37853700 | -0.02022700 |
| H | -1.20856100 | 2.33121700 | 2.36739600 |
| H | 3.71818100 | 4.08436200 | -4.96603300 |
| P | 2.78610300 | 6.87667800 | -2.35299100 |
| O | 4.37107800 | 6.41693500 | -2.14986900 |
| O | 2.13035000 | 6.64695100 | -0.98518500 |
| O | 2.71670200 | 8.15049700 | -3.11277100 |
| C | 0.23632700 | 1.69537700 | -1.19172900 |
| N | -0.83447500 | 1.52721200 | 0.99765000 |
| C | -3.66536600 | 1.14157500 | -0.80985600 |
| H | -3.75299700 | 1.59233600 | 0.18359200 |
| H | -3.59588100 | 1.91288300 | -1.58651500 |
| H | -2.82712800 | 0.43935600 | -0.90069600 |
| H | 4.89355000 | 6.95432000 | -2.76164100 |
| H | 3.84872100 | 2.42005500 | -6.49596200 |
| H | 6.43702500 | -5.77639800 | 0.87088500 |
| H | 2.32384900 | -5.50625000 | 2.67175900 |
| H | 2.39492100 | -5.08883600 | 7.44822300 |
| H | 4.01429600 | -4.60737300 | 8.00480100 |
| H | -3.69261500 | -3.00570100 | 2.66542800 |
| H | -3.67459800 | -2.65570200 | 4.41511800 |
| H | -7.24916500 | 3.84128300 | 2.20551600 |
| H | -7.26385000 | 2.92845200 | 3.74301800 |
| H | -9.06526700 | -0.94945000 | 1.54430500 |
| H | -2.52713500 | 7.59228300 | 6.96031700 |
| H | -2.96433800 | 5.92335000 | 6.50577700 |
| H | -2.09407100 | -5.02292100 | -4.09602000 |
| H | -3.28769600 | -5.87419700 | 0.19857900 |
| H | -4.65176600 | -1.28820300 | -6.37947100 |
| H | -3.60114800 | -0.44058200 | -7.54688100 |
| H | -7.98397900 | -0.62752700 | -4.28492700 |
| H | -6.85472300 | 0.74926500 | -4.27867600 |
| H | -2.95083000 | -7.58765100 | -0.21674500 |
| H | 4.94939700 | -4.97825400 | -3.59152500 |
| H | 5.45288200 | 2.85164200 | 0.39056200 |

#### Model1 – TS2

| | | | |
|---|---|---|---|
| C | -1.79853700 | 6.96206400 | 6.29543100 |
| C | -1.55798300 | 7.33601300 | 4.82554000 |
| C | -1.26632500 | 6.13438200 | 3.95453100 |
| O | -0.22643400 | 5.98487200 | 3.33321200 |
| O | -2.24694000 | 5.24652800 | 3.94228000 |
| H | -0.71164100 | 8.02852000 | 4.72177800 |
| H | -0.91389000 | 6.46923600 | 6.72852900 |
| H | -2.45499700 | 7.83206900 | 4.41760400 |
| C | 5.25910900 | 3.25770500 | -0.18103700 |
| C | 4.52007200 | 4.32368300 | 0.60576600 |
| N | 3.31236900 | 3.75268600 | 1.19069700 |
| C | 2.32074000 | 4.45179600 | 1.73869900 |
| N | 1.37760800 | 3.78482600 | 2.42273300 |
| N | 2.28079500 | 5.78811700 | 1.66278400 |
| H | 1.47171400 | 6.22779400 | 2.09426500 |
| H | 4.61491400 | 2.82023400 | -0.95815200 |
| H | 4.25555100 | 5.15780700 | -0.05413600 |
| H | 3.20884700 | 2.72854800 | 1.20538200 |
| H | 1.45193900 | 2.78328000 | 2.53404100 |
| H | 2.63171200 | 6.21976900 | 0.78533700 |
| H | 6.12859100 | 3.70449700 | -0.68288300 |
| H | 5.16639700 | 4.72658600 | 1.40730700 |
| H | 0.54673300 | 4.27185000 | 2.75305500 |
| C | 3.77037100 | -4.68274800 | -3.34921900 |
| C | 4.05946600 | -3.47653300 | -4.23828700 |
| C | 5.29954800 | -2.69446700 | -3.78230200 |
| N | 5.18123900 | -2.15697400 | -2.44110600 |
| C | 4.46925100 | -1.05832300 | -2.13596000 |
| N | 3.98089600 | -0.24680600 | -3.10089800 |
| N | 4.30881400 | -0.76142600 | -0.85485900 |
| H | 3.71201600 | -0.00195900 | -0.52775900 |
| H | 3.65219800 | -4.38773600 | -2.29844400 |
| H | 3.18801000 | -2.79786400 | -4.24256600 |
| H | 5.53511000 | -1.88156400 | -4.49006500 |
| H | 5.36505200 | -2.80985600 | -1.65044400 |
| H | 3.96542400 | -0.57733500 | -4.05596100 |
| H | 4.86198300 | -1.30728500 | -0.16383400 |
| H | 2.85263100 | -5.20361800 | -3.66324900 |
| H | 4.21165400 | -3.79468300 | -5.28444500 |
| H | 6.17628100 | -3.36079600 | -3.78691400 |
| H | 3.16589200 | 0.32686200 | -2.85741700 |
| C | 5.99416300 | -5.66634200 | 2.04170800 |
| C | 4.57464000 | -5.58342700 | 2.59917300 |
| O | 4.32596900 | -5.75573500 | 3.79098000 |
| C | 6.57361100 | -4.25307200 | 1.84557400 |

| | | | |
|---|---|---|---|
| C | 5.93438600 | -3.42248300 | 0.71964800 |
| O | 5.47309500 | -4.01498300 | -0.29030300 |
| O | 5.94685000 | -2.16678800 | 0.85756700 |
| H | 7.64632400 | -4.32941500 | 1.59460900 |
| H | 6.59626100 | -6.23084400 | 2.76570700 |
| H | 6.51516700 | -3.67091400 | 2.77824300 |
| N | 3.62561300 | -5.23469600 | 1.69531000 |
| C | 2.27913900 | -4.85614400 | 2.09665700 |
| C | 1.35139500 | -4.66116500 | 0.89339000 |
| C | 1.87420700 | -3.57947200 | -0.06928300 |
| C | -0.06266600 | -4.33148900 | 1.38655100 |
| C | 1.05891900 | -3.44515400 | -1.35353300 |
| H | 0.02446300 | -3.12885600 | -1.15184500 |
| H | 3.95709200 | -4.98872700 | 0.76336200 |
| H | 2.31792200 | -3.92315500 | 2.68909500 |
| H | 1.31356100 | -5.62030900 | 0.34227500 |
| H | 2.92035900 | -3.78248300 | -0.34417600 |
| H | -0.41389800 | -5.07217400 | 2.12263900 |
| H | 1.01404400 | -4.41118100 | -1.88169900 |
| H | 1.91513500 | -2.62157600 | 0.47744100 |
| H | -0.78374300 | -4.31705400 | 0.55698400 |
| H | 1.49938400 | -2.69153800 | -2.01926900 |
| H | -0.08352700 | -3.33866300 | 1.86471600 |
| C | 2.96235900 | -4.92983000 | 7.25358200 |
| C | 2.96447200 | -3.75126100 | 6.28402500 |
| C | 4.38173900 | -3.42121200 | 5.83008000 |
| C | 4.48186200 | -2.27961700 | 4.85104600 |
| N | 5.64692000 | -1.60125300 | 4.84568300 |
| O | 3.56761300 | -1.99284900 | 4.05912900 |
| H | 6.40219600 | -1.86621700 | 5.46266200 |
| H | 3.38362400 | -5.82434600 | 6.76781400 |
| H | 2.50038300 | -2.86477300 | 6.75122100 |
| H | 4.78885800 | -4.30497600 | 5.30574900 |
| H | 5.78809900 | -0.88460500 | 4.12659000 |
| H | 2.36389200 | -3.99273800 | 5.39407800 |
| H | 5.03437000 | -3.21885900 | 6.69669600 |
| C | -2.95030100 | -6.37331000 | -0.00035200 |
| C | -2.32993500 | -6.14473500 | -1.36792400 |
| O | -1.83448600 | -7.04624200 | -2.02275600 |
| H | -2.15301900 | -6.65610100 | 0.70320800 |
| N | -2.34586700 | -4.85672700 | -1.82271400 |
| C | -1.77648700 | -4.51451100 | -3.09734400 |
| C | -1.69934100 | -3.00007000 | -3.22923000 |
| O | -2.36336800 | -2.26284200 | -2.50648500 |
| H | -2.75785200 | -4.09302800 | -1.29707300 |
| H | -0.78365200 | -4.97830200 | -3.20939200 |
| N | -0.89399000 | -2.52975100 | -4.21677900 |
| C | -0.64819200 | -1.09339000 | -4.34963600 |
| C | -1.88286000 | -0.33817500 | -4.83634500 |
| O | -2.16211800 | 0.77065800 | -4.40865300 |
| C | 0.57796100 | -0.81444200 | -5.21326400 |
| H | 0.42884100 | -1.13102500 | -6.25836500 |
| H | -0.22542000 | -3.17043300 | -4.63101300 |
| H | -0.45078800 | -0.68707300 | -3.34906700 |
| H | 1.44799600 | -1.34170100 | -4.79236800 |
| H | 0.81595100 | 0.25799900 | -5.18752000 |
| N | -2.58783500 | -0.98865000 | -5.79694700 |
| C | -3.70213700 | -0.39281700 | -6.48210700 |
| H | -2.27525800 | -1.91492000 | -6.06002500 |
| H | -3.95796900 | 0.54029800 | -5.96393600 |
| C | -6.38510400 | 4.04460200 | 2.95244400 |
| C | -5.13900100 | 3.34580600 | 2.46523600 |
| O | -4.88158400 | 3.19600300 | 1.27935100 |
| H | -6.16380400 | 4.72230400 | 3.79100600 |
| N | -4.34140700 | 2.86536400 | 3.47023300 |
| C | -3.45556200 | 1.76023700 | 3.20444700 |
| C | -4.12995600 | 0.46747500 | 3.67781000 |
| O | -5.10349700 | 0.48592800 | 4.41972600 |
| C | -2.03579700 | 1.98220800 | 3.72710400 |
| O | -1.37680100 | 2.94891700 | 2.92228000 |
| H | -1.97624200 | 4.45215400 | 3.39822300 |
| H | -4.77461400 | 2.82154900 | 4.38902800 |
| H | -3.36570800 | 1.69537100 | 2.11746000 |
| H | -1.47723300 | 1.03435400 | 3.71652900 |
| H | -2.05377400 | 2.36971200 | 4.75733300 |
| N | -3.60136200 | -0.65885600 | 3.15519600 |
| C | -4.21495000 | -1.93183200 | 3.43676500 |
| H | -2.71952500 | -0.61035000 | 2.63269400 |
| H | -5.28224500 | -1.90463400 | 3.16802000 |
| C | -7.10076700 | 0.36413600 | -3.77712700 |
| C | -5.98655000 | -0.64705700 | -3.51993400 |
| C | -4.85702000 | -0.10323000 | -2.65276100 |
| S | -5.34978600 | 0.20050100 | -0.90712700 |
| C | -5.26768900 | -1.51080200 | -0.24707400 |
| C | -6.58537500 | -1.99817000 | 0.32368500 |
| O | -6.96391600 | -1.20512600 | 1.43813800 |
| C | -7.79957800 | -1.97017600 | -0.61874800 |
| O | -7.84201000 | -3.02096200 | -1.53706300 |
| C | -8.95408600 | -1.97136700 | 0.39772800 |
| O | -9.22031300 | -3.31070600 | 0.79715100 |
| C | -8.39203800 | -1.08130100 | 1.50218700 |
| H | -7.55333000 | 0.71085400 | -2.83389100 |
| H | -5.53282000 | -0.95079400 | -4.47672500 |
| H | -4.49775300 | 0.86910100 | -3.02287800 |
| H | -4.50195500 | -1.52227100 | 0.54193400 |
| H | -6.42182400 | -3.05400700 | 0.63435500 |
| H | -7.79972100 | -1.01767000 | -1.17216800 |
| H | -8.29118300 | -3.75234000 | -1.07948000 |
| H | -9.89083500 | -1.58456300 | -0.02811100 |
| H | -8.60695000 | -3.56244000 | 1.50102300 |
| H | -8.75115000 | -1.38465800 | 2.49974000 |
| H | -6.39687300 | -1.57188200 | -3.08054400 |
| H | -3.99109000 | -0.77855200 | -2.62700400 |
| H | -4.93713400 | -2.17426300 | -1.05825000 |
| O | 1.54076700 | -1.04461200 | 2.39379000 |
| H | 0.58723400 | -1.02578700 | 2.58718200 |
| H | 2.01673800 | -1.46771500 | 3.13359000 |
| O | 5.00619200 | -0.29017000 | 2.44246000 |
| H | 5.50300900 | -0.94406800 | 1.88386800 |
| H | 4.33258300 | -0.86715300 | 2.87099500 |
| O | 3.00175200 | 0.94699600 | 1.14552500 |
| H | 3.83968300 | 0.64170200 | 1.58370400 |
| H | 2.34741300 | 0.32216200 | 1.53134400 |
| C | 3.65828200 | 2.81522400 | -4.70685100 |
| O | 3.34729400 | 2.06085500 | -5.85844200 |
| C | 2.40032200 | 3.18962500 | -3.97224000 |
| O | 1.75432000 | 2.05809100 | -3.37911600 |
| C | 2.69080300 | 4.14632200 | -2.81285400 |
| O | 2.60599600 | 5.47211300 | -3.26653100 |
| C | 1.65785000 | 3.71787500 | -1.72939400 |
| O | 0.81886800 | 4.70593600 | -1.23071000 |
| C | 0.83909100 | 2.63185600 | -2.45103500 |
| C | 0.74604300 | 0.34647400 | -1.20664000 |
| O | 1.56180600 | -0.28128500 | -1.87470700 |
| N | 0.20560800 | -0.26441900 | -0.07302100 |
| C | -0.76955500 | 0.23617200 | 0.74810200 |
| O | -1.14540600 | -0.43033400 | 1.73491700 |
| C | -0.75712600 | 2.13514400 | -0.64021700 |
| H | -1.17109700 | 3.13313700 | -0.80993100 |
| H | 4.32512000 | 2.25861100 | -4.01966800 |
| H | 1.70107100 | 3.69037000 | -4.66673400 |
| H | 3.70725600 | 3.95495700 | -2.43554100 |
| H | 2.21529200 | 3.21508300 | -0.91911700 |
| H | 1.39846100 | 5.49322700 | -1.04546600 |
| H | 0.02882600 | 3.14416300 | -3.00324100 |
| H | 0.62237600 | -1.15394100 | 0.19480200 |
| H | -1.36092100 | 2.59639100 | 1.99982300 |
| H | 4.15506500 | 3.77077500 | -4.96025300 |
| P | 3.33333200 | 6.65033100 | -2.34357600 |
| O | 4.88380700 | 6.05945300 | -2.23270700 |

| | | | |
|---|---|---|---|
| O | 2.74602000 | 6.46682700 | -0.93706200 |
| O | 3.32366600 | 7.92983700 | -3.09608800 |
| C | 0.23429300 | 1.67664600 | -1.46055200 |
| N | -1.31129300 | 1.44739600 | 0.41490500 |
| C | -3.23406300 | 0.96488800 | -0.19002000 |
| H | -3.26336600 | 0.20694100 | 0.58458800 |
| H | -3.63564100 | 1.95200600 | 0.03379900 |
| H | -2.83317300 | 0.71680300 | -1.16955900 |
| H | 5.41664200 | 6.56246900 | -2.86457600 |
| H | 4.13275900 | 2.02244500 | -6.41682100 |
| H | 6.00273300 | -6.19067500 | 1.07414100 |
| H | 1.88288400 | -5.63594800 | 2.76554800 |
| H | 1.94253700 | -5.17572500 | 7.58933800 |
| H | 3.56615600 | -4.71553800 | 8.15154100 |
| H | -3.70948900 | -2.71073000 | 2.84762600 |
| H | -4.14252000 | -2.20096400 | 4.50441400 |
| H | -6.83884900 | 4.60778700 | 2.12803900 |
| H | -7.09950400 | 3.28448500 | 3.30933600 |
| H | -8.67051900 | -0.02569100 | 1.34330300 |
| H | -2.01000100 | 7.86207400 | 6.89185200 |
| H | -2.65266200 | 6.27542300 | 6.38869500 |
| H | -2.38941100 | -4.90297100 | -3.93453900 |
| H | -3.47521700 | -5.49174800 | 0.39609600 |
| H | -4.58060400 | -1.05788800 | -6.48095800 |
| H | -3.45768700 | -0.14784200 | -7.53053600 |
| H | -7.90247800 | -0.07151800 | -4.39272400 |
| H | -6.71433100 | 1.25380400 | -4.30064300 |
| H | -3.64769600 | -7.22025400 | -0.06763800 |
| H | 4.59880600 | -5.40841700 | -3.38793700 |
| H | 5.60939000 | 2.43499800 | 0.46385500 |

#### Model1 − Final state

| | | | |
|---|---|---|---|
| C | -0.66688300 | 6.99390500 | 6.46244700 |
| C | -0.46229800 | 7.42875800 | 5.00394900 |
| C | -0.32922100 | 6.22177700 | 4.10903600 |
| O | 0.71722600 | 5.89321800 | 3.56837800 |
| O | -1.44703400 | 5.52458700 | 4.01599300 |
| H | 0.44705400 | 8.03428200 | 4.89126500 |
| H | 0.18186800 | 6.38682800 | 6.81474000 |
| H | -1.32561700 | 8.02392500 | 4.66514900 |
| C | 5.64338500 | 2.71505600 | -0.41925600 |
| C | 5.00903900 | 3.82825700 | 0.40110000 |
| N | 3.85292200 | 3.34715700 | 1.15200600 |
| C | 2.91420700 | 4.12174500 | 1.69906600 |
| N | 2.06374100 | 3.54028000 | 2.59038100 |
| N | 2.78618700 | 5.40752700 | 1.39600100 |
| H | 2.05730600 | 5.91755300 | 1.88725500 |
| H | 4.91411400 | 2.28234800 | -1.11757500 |
| H | 4.69996500 | 4.64323800 | -0.25896700 |
| H | 3.81414200 | 2.36829900 | 1.43809200 |
| H | 2.44831400 | 2.71170300 | 3.05177900 |
| H | 3.12724600 | 5.79419100 | 0.48101500 |
| H | 6.48006000 | 3.12701100 | -0.99939700 |
| H | 5.75171700 | 4.25093200 | 1.10332100 |
| H | 1.49899200 | 4.19029700 | 3.14141500 |
| C | 3.16843900 | -4.94667000 | -3.65345300 |
| C | 3.27880700 | -3.60825500 | -4.37624900 |
| C | 4.58235900 | -2.87562900 | -4.03588700 |
| N | 4.67953200 | -2.48605200 | -2.64374700 |
| C | 4.19445000 | -1.33715200 | -2.14819700 |
| N | 3.54718800 | -0.45340400 | -2.93262900 |
| N | 4.37305400 | -1.08652200 | -0.85885500 |
| H | 3.94485600 | -0.26189500 | -0.43969500 |
| H | 3.26039000 | -4.82314800 | -2.56381500 |
| H | 2.42512400 | -2.96181100 | -4.10401300 |
| H | 4.71381600 | -1.98565500 | -4.67338000 |
| H | 4.93245200 | -3.20735600 | -1.93306300 |
| H | 3.34019100 | -0.64906200 | -3.90076200 |
| H | 5.01377000 | -1.70884300 | -0.32449800 |
| H | 2.20595400 | -5.44172600 | -3.85791300 |
| H | 3.23074300 | -3.75386900 | -5.46957800 |
| H | 5.43885400 | -3.53281900 | -4.25625200 |
| H | 3.19516000 | 0.43397200 | -2.57172700 |
| C | 5.52500900 | -6.28409200 | 1.60279700 |
| C | 4.14552700 | -6.06681900 | 2.21893700 |
| O | 3.90448000 | -6.31961600 | 3.39830400 |
| C | 6.27859700 | -4.94945200 | 1.44055500 |
| C | 5.74914500 | -3.97238900 | 0.37523200 |
| O | 5.07232400 | -4.41631800 | -0.59087700 |
| O | 6.06847800 | -2.75758100 | 0.51619000 |
| H | 7.32697300 | -5.15357000 | 1.15984700 |
| H | 6.07796500 | -6.95179400 | 2.27589000 |
| H | 6.32637900 | -4.40657900 | 2.39749600 |
| N | 3.22669900 | -5.52579300 | 1.38442300 |
| C | 1.92454900 | -5.08840300 | 1.85469000 |
| C | 1.05963300 | -4.53868800 | 0.70798400 |
| C | 1.69131400 | -3.29062100 | 0.05901600 |
| C | -0.35753900 | -4.25366600 | 1.21689000 |
| C | 0.94933400 | -2.76318100 | -1.17100800 |
| H | -0.08047600 | -2.44843400 | -0.94107100 |
| H | 3.55579100 | -5.24794200 | 0.45937100 |
| H | 2.05473600 | -4.31396500 | 2.63183200 |
| H | 0.98871200 | -5.33081500 | -0.06195600 |
| H | 2.73122700 | -3.50373900 | -0.23088700 |
| H | -0.78274400 | -5.13431600 | 1.72290000 |
| H | 0.89140200 | -3.53799200 | -1.95386800 |
| H | 1.75430100 | -2.49831300 | 0.82233000 |
| H | -1.03686600 | -3.98177700 | 0.39621300 |
| H | 1.47005600 | -1.89612600 | -1.60584400 |
| H | -0.35905600 | -3.41516400 | 1.93294300 |
| C | 2.83726500 | -5.35367200 | 6.96964500 |
| C | 2.93118800 | -4.21249300 | 5.96352300 |
| C | 4.35921900 | -4.06274700 | 5.45040100 |
| C | 4.55076900 | -2.95020400 | 4.45638700 |
| N | 5.74329900 | -2.33068600 | 4.46936000 |
| O | 3.67276500 | -2.63592900 | 3.62881200 |
| H | 6.46556500 | -2.60952900 | 5.11886200 |
| H | 3.13899700 | -6.30329800 | 6.49985500 |
| H | 2.58976400 | -3.26573800 | 6.41836200 |
| H | 4.63009400 | -4.99338800 | 4.91945600 |
| H | 5.91935100 | -1.61535700 | 3.75503800 |
| H | 2.27357600 | -4.41251200 | 5.10439400 |
| H | 5.06467100 | -3.93529100 | 6.28877400 |
| C | -3.52963100 | -5.99484700 | -0.00974000 |
| C | -2.67802400 | -5.79906100 | -1.25202300 |
| O | -1.96857600 | -6.68114600 | -1.70734300 |
| H | -2.87621100 | -6.31143400 | 0.81574300 |
| N | -2.76476000 | -4.57280400 | -1.85061400 |
| C | -1.99006300 | -4.27181100 | -3.02352700 |
| C | -1.93738800 | -2.76429300 | -3.22396500 |
| O | -2.54068800 | -1.99893900 | -2.48524300 |
| H | -3.26595900 | -3.79351900 | -1.43372600 |
| H | -0.97103900 | -4.68273700 | -2.91776800 |
| N | -1.19545700 | -2.31392200 | -4.27586000 |
| C | -0.88384500 | -0.88829700 | -4.37037100 |
| C | -2.09711300 | -0.03214200 | -4.73047900 |
| O | -2.29273200 | 1.04960400 | -4.19405300 |
| C | 0.27574100 | -0.64352400 | -5.33570200 |
| H | -0.00328300 | -0.86851700 | -6.37710000 |
| H | -0.57739700 | -2.96916000 | -4.74336400 |
| H | -0.58102300 | -0.53633200 | -3.37234900 |
| H | 1.13361600 | -1.27608800 | -5.05665300 |
| H | 0.61033700 | 0.40105600 | -5.27139400 |
| N | -2.85331800 | -0.54108200 | -5.73047900 |
| C | -3.94224200 | 0.18076400 | -6.33719100 |
| H | -2.63255600 | -1.47599900 | -6.05152500 |
| H | -4.01543400 | 1.15672100 | -5.84027100 |
| C | -5.69048300 | 4.65638400 | 3.29664700 |
| C | -4.77102400 | 3.64649600 | 2.62673000 |
| O | -4.99472300 | 3.15737500 | 1.53362000 |

| | | | |
|---|---|---|---|
| H | -5.11912800 | 5.44459800 | 3.81018000 |
| N | -3.65041300 | 3.31245200 | 3.36195100 |
| C | -2.97491500 | 2.07049400 | 3.06807000 |
| C | -3.83380800 | 0.88273500 | 3.57039200 |
| O | -4.83570000 | 1.07388100 | 4.24359500 |
| C | -1.56914400 | 2.04672500 | 3.67526400 |
| O | -0.76775600 | 3.12273400 | 3.21125100 |
| H | -1.26803000 | 4.66765200 | 3.53635800 |
| H | -3.69402200 | 3.55230100 | 4.34753100 |
| H | -2.89158000 | 1.99561400 | 1.97487600 |
| H | -1.08208700 | 1.08453600 | 3.46368600 |
| H | -1.64186400 | 2.14651100 | 4.77170100 |
| N | -3.38867300 | -0.34342100 | 3.22080100 |
| C | -4.15035200 | -1.52571900 | 3.56546900 |
| H | -2.56843400 | -0.44078400 | 2.62290000 |
| H | -5.07648300 | -1.59989500 | 2.96809500 |
| C | -7.10991900 | 1.22969400 | -3.45510200 |
| C | -6.77259700 | -0.25761600 | -3.54311000 |
| C | -5.55928600 | -0.66034900 | -2.70457100 |
| S | -5.82335900 | -0.60563400 | -0.88751700 |
| C | -6.80991500 | -2.13295300 | -0.64525300 |
| C | -8.00079100 | -1.90092400 | 0.27166000 |
| O | -7.58111700 | -1.39667100 | 1.54152000 |
| C | -9.05407100 | -0.90648300 | -0.26906100 |
| O | -10.05371000 | -1.50295000 | -1.04480100 |
| C | -9.57276100 | -0.27367800 | 1.03343400 |
| O | -10.55815400 | -1.13131200 | 1.60254000 |
| C | -8.29883800 | -0.19356500 | 1.85401100 |
| H | -7.28945600 | 1.52843100 | -2.40999600 |
| H | -6.56256300 | -0.52827900 | -4.59440200 |
| H | -4.71785200 | 0.02702400 | -2.88730600 |
| H | -6.16143400 | -2.91867000 | -0.22686400 |
| H | -8.50275900 | -2.87843800 | 0.41896400 |
| H | -8.54860000 | -0.14078300 | -0.87786600 |
| H | -10.72354400 | -1.80562800 | -0.40779400 |
| H | -10.05339500 | 0.70213100 | 0.87345500 |
| H | -10.10079700 | -1.83408900 | 2.08608200 |
| H | -8.48519600 | -0.16346100 | 2.93826400 |
| H | -7.64806200 | -0.86313600 | -3.24941200 |
| H | -5.19556900 | -1.66622400 | -2.96062600 |
| H | -7.18362600 | -2.48091000 | -1.62029100 |
| O | 1.62840800 | -1.04223200 | 2.88854300 |
| H | 0.68899900 | -1.28556400 | 2.84080700 |
| H | 2.16024400 | -1.79672000 | 3.22342800 |
| O | 5.17966400 | -0.88057100 | 2.15706000 |
| H | 5.71275900 | -1.46430500 | 1.56371200 |
| H | 4.47634200 | -1.50006200 | 2.45612100 |
| O | 3.41360000 | 1.06741700 | 2.93753300 |
| H | 4.21887800 | 0.56115400 | 2.70390900 |
| H | 2.73181700 | 0.37732000 | 3.05378000 |
| C | 3.79623600 | 2.55235500 | -4.86890600 |
| O | 3.35017500 | 1.86007700 | -6.01629900 |
| C | 2.61878200 | 3.01664300 | -4.05748100 |
| O | 1.93680200 | 1.93767100 | -3.40707200 |
| C | 3.01773200 | 3.93876000 | -2.91314300 |
| O | 3.23924100 | 5.22995200 | -3.40812500 |
| C | 1.84221000 | 3.78632500 | -1.89526200 |
| O | 1.00228400 | 4.89200200 | -1.81377900 |
| C | 1.05788700 | 2.57825300 | -2.46997500 |
| C | 1.47184300 | 1.02486900 | -0.47839400 |
| O | 2.69912200 | 1.11153000 | -0.51789200 |
| N | 0.86039900 | 0.30760900 | 0.54339200 |
| C | -0.48093200 | 0.04367700 | 0.68594300 |
| O | -0.90973500 | -0.60197900 | 1.64838800 |
| C | -0.76530700 | 1.32245800 | -1.32241100 |
| H | -1.48919800 | 1.66732100 | -2.06519000 |
| H | 4.44950800 | 1.92054600 | -4.23354000 |
| H | 1.91382100 | 3.55622100 | -4.71646000 |
| H | 3.93810800 | 3.54373900 | -2.45228300 |
| H | 2.26622400 | 3.52685200 | -0.91397500 |
| H | 1.59287300 | 5.61964700 | -1.51145800 |
| H | 0.19470100 | 2.99877600 | -3.01471900 |
| H | 1.46230600 | -0.10983400 | 1.26963500 |
| H | -0.16694000 | 2.87642700 | 2.49122800 |
| H | 4.36117100 | 3.46636200 | -5.12857600 |
| P | 3.99098700 | 6.34199500 | -2.42202600 |
| O | 5.45732100 | 5.60050700 | -2.17131300 |
| O | 3.24865100 | 6.23411900 | -1.08527100 |
| O | 4.18195200 | 7.60466800 | -3.17729600 |
| C | 0.55785700 | 1.61766100 | -1.43765800 |
| N | -1.28782500 | 0.55477700 | -0.30842000 |
| C | -2.73130300 | 0.28016200 | -0.30472900 |
| H | -2.93999400 | -0.53898100 | 0.38490200 |
| H | -3.30497200 | 1.16752500 | 0.00211000 |
| H | -3.01871200 | -0.03753200 | -1.30978000 |
| H | 6.09300400 | 6.04740500 | -2.74785600 |
| H | 4.08320300 | 1.79802300 | -6.64024100 |
| H | 5.43245900 | -6.76333900 | 0.61592400 |
| H | 1.41479900 | -5.93729100 | 2.33906300 |
| H | 1.81041600 | -5.47631100 | 7.34857600 |
| H | 3.49453900 | -5.18150300 | 7.83847200 |
| H | -3.53330400 | -2.41673600 | 3.38207600 |
| H | -4.43936900 | -1.49845600 | 4.62635400 |
| H | -6.35144600 | 5.09912600 | 2.54181400 |
| H | -6.30168100 | 4.12408500 | 4.04366300 |
| H | -7.70036800 | 0.69305700 | 1.57658400 |
| H | -0.75088600 | 7.87130000 | 7.12088800 |
| H | -1.58384000 | 6.39542000 | 6.56554300 |
| H | -2.41164700 | -4.74352400 | -3.93211200 |
| H | -4.08408300 | -5.09346700 | 0.29010700 |
| H | -4.90142800 | -0.34844100 | -6.21307000 |
| H | -3.76695500 | 0.34282600 | -7.41356600 |
| H | -8.01025700 | 1.47368500 | -4.03918400 |
| H | -6.27772200 | 1.84512500 | -3.83488200 |
| H | -4.24117300 | -6.81256600 | -0.19762500 |
| H | 3.97013500 | -5.63313500 | -3.97071700 |
| H | 6.01961400 | 1.89779900 | 0.21880000 |

#### Model2 – Initial state

| | | | |
|---|---|---|---|
| O | -3.14305800 | -0.91382300 | -3.04676400 |
| H | -2.51128400 | -1.13149200 | -3.74619300 |
| H | -3.77742800 | -1.66750900 | -3.08871000 |
| C | 1.85038200 | 7.57717600 | -5.24252900 |
| C | 1.67293100 | 7.55345100 | -3.71841600 |
| C | 1.34574600 | 6.15368600 | -3.20681700 |
| O | 0.17989900 | 5.86893000 | -2.89942500 |
| O | 2.33575700 | 5.33096600 | -3.15704800 |
| H | 0.92683600 | 7.25240600 | -5.74804800 |
| H | 0.85676700 | 8.22543000 | -3.41628600 |
| H | 2.60307000 | 7.89077700 | -3.23238800 |
| C | -5.57584500 | 2.14142000 | -1.59071500 |
| C | -4.52029300 | 3.11479700 | -2.09635900 |
| N | -3.24785300 | 2.43431800 | -2.38664100 |
| C | -2.07090500 | 3.08343400 | -2.54827100 |
| N | -1.82793600 | 4.23109000 | -1.94042900 |
| N | -1.12093600 | 2.53667100 | -3.33104200 |
| H | -5.22962300 | 1.61474600 | -0.68962500 |
| H | -6.49084300 | 2.69708500 | -1.34012500 |
| H | -4.35054000 | 3.89174800 | -1.34362700 |
| H | -4.87933000 | 3.62232700 | -3.00968300 |
| H | -3.30925000 | 1.48567800 | -2.75518400 |
| H | -2.37519000 | 4.49266200 | -1.09556900 |
| H | -1.01389100 | 4.80326300 | -2.23833000 |
| H | -1.33427600 | 1.73213900 | -3.90332100 |
| H | -0.13617700 | 2.83413300 | -3.23918900 |
| C | -3.65989800 | -5.76270300 | 1.62779100 |
| C | -4.34203200 | -4.76390400 | 2.56141500 |
| C | -5.64696500 | -4.20667300 | 1.97419200 |
| N | -5.45676600 | -3.50377700 | 0.72078700 |
| C | -4.98752400 | -2.25317400 | 0.62847800 |

| | | | |
|---|---|---|---|
| N | -4.85645800 | -1.45982800 | 1.71300000 |
| N | -4.66651900 | -1.77580100 | -0.56887300 |
| H | -3.45272700 | -5.31700700 | 0.64372000 |
| H | -2.70896900 | -6.12552400 | 2.04982300 |
| H | -3.66239500 | -3.91965500 | 2.77949400 |
| H | -4.57012800 | -5.23669100 | 3.53262600 |
| H | -6.15676000 | -3.55141400 | 2.70098300 |
| H | -6.34428400 | -5.03599900 | 1.77744100 |
| H | -5.32425300 | -4.08440200 | -0.14620500 |
| H | -5.06133600 | -1.78405800 | 2.64583300 |
| H | -4.41002100 | -0.54802000 | 1.62268300 |
| H | -4.76999700 | -2.36487500 | -1.42572200 |
| H | -4.18364400 | -0.87955500 | -0.60142600 |
| C | -4.90183000 | -6.69319500 | -3.91822100 |
| C | -3.47001900 | -6.34363200 | -4.41504600 |
| O | -3.15076400 | -6.44884100 | -5.58845400 |
| C | -5.66710900 | -5.35393900 | -3.76739300 |
| C | -5.12278600 | -4.43473200 | -2.65722200 |
| O | -4.91684500 | -3.21563100 | -2.91823800 |
| O | -4.93412200 | -4.96528800 | -1.52412500 |
| H | -5.36473400 | -7.32950400 | -4.68213600 |
| H | -6.71999800 | -5.56267300 | -3.50949200 |
| H | -5.66115000 | -4.80494100 | -4.71985500 |
| N | -2.62718100 | -5.80202400 | -3.47552600 |
| C | -1.36922800 | -5.09721700 | -3.78192200 |
| C | -0.70230800 | -4.61791200 | -2.48287000 |
| C | -1.61043500 | -3.63501800 | -1.72225800 |
| C | 0.66979800 | -4.00349900 | -2.77748900 |
| C | -1.14158900 | -3.26704200 | -0.31824400 |
| H | -3.00197100 | -5.70025400 | -2.53743600 |
| H | -1.57486200 | -4.23951300 | -4.45080100 |
| H | -0.53839600 | -5.50735300 | -1.84792400 |
| H | -2.62746800 | -4.04126300 | -1.63046600 |
| H | -1.72754100 | -2.72269600 | -2.32652500 |
| H | 1.21101900 | -3.77104800 | -1.84762800 |
| H | 0.57223100 | -3.05992400 | -3.34084800 |
| H | 1.29677800 | -4.69294400 | -3.36517400 |
| H | -1.08501300 | -4.17305300 | 0.30564800 |
| H | -1.85402800 | -2.57495600 | 0.15801100 |
| H | -0.15188400 | -2.78394600 | -0.31121500 |
| C | 3.54673700 | -6.07849300 | -0.82111600 |
| C | 2.30673300 | -5.81856800 | 0.02317100 |
| O | 1.26345000 | -6.42169500 | -0.16288100 |
| H | 3.30832700 | -5.84267200 | -1.86884500 |
| N | 2.42884400 | -4.87382800 | 1.01313300 |
| C | 1.35560500 | -4.62080800 | 1.93975600 |
| C | 1.23439100 | -3.14632200 | 2.31807700 |
| O | 2.20307000 | -2.39207600 | 2.28974000 |
| H | 3.29798000 | -4.37483400 | 1.15697300 |
| H | 0.43370000 | -5.01259400 | 1.48978600 |
| N | 0.00659400 | -2.77476900 | 2.77300600 |
| C | -0.30893600 | -1.39853600 | 3.17160500 |
| C | 0.48040600 | -1.00107300 | 4.42747700 |
| O | 1.15275100 | 0.01779700 | 4.47939400 |
| C | -1.81170700 | -1.20873500 | 3.33828400 |
| H | -0.77398600 | -3.38674300 | 2.55937100 |
| H | 0.05505800 | -0.71450400 | 2.38878700 |
| H | -2.04421300 | -0.14779100 | 3.50831900 |
| H | -2.21081700 | -1.80365400 | 4.17560700 |
| H | -2.33450300 | -1.50247100 | 2.41728600 |
| N | 0.32530800 | -1.86377900 | 5.46476100 |
| C | 0.88671000 | -1.63461300 | 6.77211800 |
| H | -0.21326600 | -2.70542900 | 5.30066700 |
| H | 1.34115900 | -0.63531300 | 6.77031200 |
| C | 6.66818800 | 4.79969300 | -2.01929800 |
| C | 5.73218700 | 3.66147300 | -1.64108800 |
| O | 5.97041800 | 2.91071500 | -0.69429400 |
| H | 6.13244100 | 5.64801800 | -2.47102400 |
| N | 4.64913400 | 3.48099000 | -2.44042700 |
| C | 3.91573000 | 2.23660500 | -2.44003500 |

| | | | |
|---|---|---|---|
| C | 4.81808000 | 1.07948400 | -2.91481900 |
| O | 5.88741900 | 1.25325000 | -3.46291400 |
| C | 2.64074400 | 2.32932000 | -3.29986000 |
| O | 1.63871600 | 3.06922700 | -2.66085700 |
| H | 4.36802300 | 4.20595700 | -3.09534000 |
| H | 3.58308900 | 1.99396600 | -1.41829000 |
| H | 2.26630600 | 1.31196300 | -3.49016800 |
| H | 2.90280100 | 2.77650200 | -4.27826800 |
| H | 1.90357400 | 4.15717100 | -2.81288000 |
| N | 4.30318200 | -0.17092600 | -2.67047700 |
| C | 4.94058000 | -1.32352700 | -3.28617900 |
| H | 3.31320000 | -0.24190700 | -2.44520400 |
| H | 6.01620000 | -1.30189900 | -3.06262100 |
| C | -4.45122700 | 1.94062300 | 4.46138100 |
| O | -5.03907100 | 0.78412500 | 3.87947100 |
| C | -3.66147500 | 2.83337100 | 3.51308700 |
| O | -2.66609100 | 2.05335200 | 2.85529200 |
| C | -4.42786800 | 3.61540800 | 2.40956800 |
| O | -4.62564400 | 4.93373600 | 2.83706100 |
| C | -3.49092400 | 3.45665600 | 1.16143300 |
| O | -3.17177700 | 4.62130200 | 0.46561800 |
| C | -2.21787300 | 2.81745400 | 1.75046000 |
| N | 0.45417300 | 1.73764600 | -0.69019900 |
| C | -0.02233400 | 0.58067400 | -1.27808700 |
| O | 0.58289900 | -0.02847800 | -2.14864500 |
| N | -1.25946000 | 0.17565500 | -0.80589200 |
| C | -2.04033300 | 0.81294700 | 0.15695800 |
| O | -3.16728300 | 0.37587400 | 0.39785400 |
| C | -1.44067900 | 1.98400200 | 0.77537700 |
| C | -0.25002600 | 2.40959300 | 0.26968700 |
| H | 1.10927600 | 2.28792700 | -1.31475700 |
| H | -1.72401000 | -0.56112100 | -1.34497100 |
| H | 0.19665800 | 3.35039800 | 0.59953000 |
| H | -1.57873700 | 3.65550500 | 2.09240800 |
| H | -4.03647600 | 5.20995600 | 0.42512600 |
| H | -5.40388900 | 3.14261200 | 2.18802600 |
| H | -3.18126500 | 3.62007900 | 4.12818400 |
| H | -5.20703900 | 2.57172100 | 4.96497200 |
| H | -3.96483500 | 2.70829300 | 0.50840400 |
| H | -3.76314900 | 1.56488200 | 5.23472500 |
| P | -5.68894000 | 5.96362000 | 2.01393200 |
| O | -5.35603300 | 5.78399200 | 0.52125200 |
| O | -5.69048600 | 7.25891200 | 2.72857600 |
| O | -7.09675700 | 5.10833700 | 2.22093200 |
| C | 6.08122100 | 0.03748100 | 5.39061700 |
| C | 5.10850500 | -0.80469300 | 4.55081600 |
| C | 4.21913400 | -0.02794300 | 3.57850300 |
| C | 4.01283000 | 1.80064800 | 1.38394200 |
| S | 5.18869100 | 0.68962400 | 2.18341800 |
| C | 5.27325000 | -0.71274500 | 0.98758800 |
| C | 6.70140400 | -1.07657800 | 0.62970000 |
| O | 7.28780500 | -0.02440500 | -0.09206700 |
| C | 7.68793600 | -1.33380400 | 1.77151800 |
| O | 7.45812400 | -2.59593100 | 2.36149800 |
| C | 9.02308600 | -1.22042400 | 1.00857600 |
| O | 9.25592900 | -2.40976200 | 0.29275900 |
| C | 8.72162200 | -0.08403500 | 0.02737700 |
| H | 6.80536300 | 0.57902400 | 4.75952200 |
| H | 5.65123600 | -1.59401100 | 4.00166300 |
| H | 4.40930500 | -1.33705500 | 5.21703500 |
| H | 3.73447300 | 0.83517800 | 4.05790200 |
| H | 3.42656300 | -0.65316700 | 3.13994800 |
| H | 3.81284200 | 2.61804800 | 2.08916600 |
| H | 4.56037800 | 2.18046300 | 0.49954600 |
| H | 3.09012600 | 1.25915500 | 1.13521700 |
| H | 9.17580600 | -0.29988300 | -0.95135500 |
| H | 8.97580200 | -3.14404900 | 0.86136100 |
| H | 7.61578700 | -0.51813400 | 2.51460000 |
| H | 6.64212100 | -2.00691300 | 0.02711000 |
| H | 4.71950900 | -1.55071700 | 1.43338800 |

| | | | |
|---|---|---|---|
| H | 7.93037700 | -2.64172000 | 3.20360000 |
| H | 9.86930700 | -0.97696100 | 1.68059600 |
| H | 4.75478400 | -0.37433700 | 0.07759000 |
| H | 2.09423000 | 8.58990300 | -5.59959000 |
| H | 2.66179800 | 6.89848900 | -5.54910900 |
| H | 7.38453100 | 4.40873100 | -2.75944200 |
| H | 7.21978500 | 5.12922600 | -1.13001900 |
| H | 9.07582600 | 0.89485900 | 0.38560600 |
| H | 6.65078000 | -0.60124300 | 6.08212100 |
| H | 5.54044600 | 0.78581700 | 5.99109400 |
| H | 1.66519800 | -2.37583000 | 7.02472000 |
| H | 0.10933900 | -1.66855700 | 7.55253300 |
| H | 1.50052200 | -5.17325900 | 2.89067700 |
| H | 3.77910500 | -7.15194800 | -0.77102200 |
| H | 4.42673500 | -5.49759200 | -0.50807300 |
| H | -0.70039300 | -5.77497500 | -4.33572200 |
| H | -4.88216500 | -7.21357700 | -2.94877900 |
| H | -4.30210800 | -6.63998600 | 1.45000900 |
| H | -5.73095200 | 1.08234900 | 3.27244300 |
| H | -7.43170500 | 4.91791700 | 1.33431300 |
| H | -5.83101400 | 1.38185600 | -2.34842200 |
| H | 4.50084800 | -2.24447500 | -2.87760800 |
| H | 4.82548300 | -1.33035600 | -4.38389800 |

#### Model2 − $TS1_1$

| | | | |
|---|---|---|---|
| O | -3.14311700 | -0.91324000 | -3.04959700 |
| H | -2.51093800 | -1.13234800 | -3.74818000 |
| H | -3.77825300 | -1.66609900 | -3.09190200 |
| C | 1.85010500 | 7.57414200 | -5.24695200 |
| C | 1.66669200 | 7.55544000 | -3.72346200 |
| C | 1.33102000 | 6.15983400 | -3.20829700 |
| O | 0.16786400 | 5.88265900 | -2.89024400 |
| O | 2.31807500 | 5.32990400 | -3.16607200 |
| H | 0.92710500 | 7.25252400 | -5.75542200 |
| H | 0.85261800 | 8.23172500 | -3.42563500 |
| H | 2.59653100 | 7.88978100 | -3.23474200 |
| C | -5.57583800 | 2.13988200 | -1.59223300 |
| C | -4.51746200 | 3.10877500 | -2.10045100 |
| N | -3.24539900 | 2.42585400 | -2.38719200 |
| C | -2.06800200 | 3.07459800 | -2.54687100 |
| N | -1.82854700 | 4.22394000 | -1.94054800 |
| N | -1.11476800 | 2.52807600 | -3.32576600 |
| H | -5.23127500 | 1.61499200 | -0.68941200 |
| H | -6.48920400 | 2.69901900 | -1.34341900 |
| H | -4.34728900 | 3.88780600 | -1.35002400 |
| H | -4.87466500 | 3.61394700 | -3.01581400 |
| H | -3.30653800 | 1.47627200 | -2.75361900 |
| H | -2.37767500 | 4.48646900 | -1.09715200 |
| H | -1.01576200 | 4.79759500 | -2.23683400 |
| H | -1.32493800 | 1.71986300 | -3.89409400 |
| H | -0.12922100 | 2.82483900 | -3.22871000 |
| C | -3.65953900 | -5.76279200 | 1.62963900 |
| C | -4.34119700 | -4.76239000 | 2.56194700 |
| C | -5.64547100 | -4.20449100 | 1.97383100 |
| N | -5.45417700 | -3.50233400 | 0.72018100 |
| C | -4.98380000 | -2.25217600 | 0.62755800 |
| N | -4.85362800 | -1.45797400 | 1.71153600 |
| N | -4.66080300 | -1.77600000 | -0.56976800 |
| H | -3.45188300 | -5.31840200 | 0.64507800 |
| H | -2.70894100 | -6.12572700 | 2.05231700 |
| H | -3.66091100 | -3.91850600 | 2.77941200 |
| H | -4.57002200 | -5.23398600 | 3.53357200 |
| H | -6.15508200 | -3.54855100 | 2.70013700 |
| H | -6.34329800 | -5.03340900 | 1.77715700 |
| H | -5.32171500 | -4.08368000 | -0.14641200 |
| H | -5.05904000 | -1.78149700 | 2.64450200 |
| H | -4.40461900 | -0.54742200 | 1.62081700 |
| H | -4.76505200 | -2.36536200 | -1.42630400 |
| H | -4.17648300 | -0.88058600 | -0.60266500 |
| C | -4.90183800 | -6.69577000 | -3.91588000 |
| C | -3.47000600 | -6.34653800 | -4.41297100 |
| O | -3.15081000 | -6.45246200 | -5.58633600 |
| C | -5.66694100 | -5.35631900 | -3.76574300 |
| C | -5.12229300 | -4.43651800 | -2.65619900 |
| O | -4.91645500 | -3.21759300 | -2.91773300 |
| O | -4.93322900 | -4.96669500 | -1.52292500 |
| H | -5.36483500 | -7.33242700 | -4.67944900 |
| H | -6.71982000 | -5.56475200 | -3.50754700 |
| H | -5.66107300 | -4.80790600 | -4.71854600 |
| N | -2.62708800 | -5.80451300 | -3.47376200 |
| C | -1.36923800 | -5.09965400 | -3.78052200 |
| C | -0.70247100 | -4.61930800 | -2.48177400 |
| C | -1.61105100 | -3.63690400 | -1.72101600 |
| C | 0.66921400 | -4.00415300 | -2.77681700 |
| C | -1.14153400 | -3.26857200 | -0.31728700 |
| H | -3.00198900 | -5.70212700 | -2.53578100 |
| H | -1.57502600 | -4.24246300 | -4.45001800 |
| H | -0.53784700 | -5.50840900 | -1.84652700 |
| H | -2.62764500 | -4.04403600 | -1.62842600 |
| H | -1.72969700 | -2.72474700 | -2.32536600 |
| H | 1.21059600 | -3.77130100 | -1.84716000 |
| H | 0.57106400 | -3.06064200 | -3.34018300 |
| H | 1.29639000 | -4.69328000 | -3.36467300 |
| H | -1.08369900 | -4.17465900 | 0.30635600 |
| H | -1.85435100 | -2.57733500 | 0.15963300 |
| H | -0.15226100 | -2.78459000 | -0.31085300 |
| C | 3.54694200 | -6.07955600 | -0.81961800 |
| C | 2.30702400 | -5.81919200 | 0.02466600 |
| O | 1.26384400 | -6.42270900 | -0.16075400 |
| H | 3.30851000 | -5.84394200 | -1.86739300 |
| N | 2.42921800 | -4.87400200 | 1.01416300 |
| C | 1.35604500 | -4.62060200 | 1.94074900 |
| C | 1.23472500 | -3.14593800 | 2.31832800 |
| O | 2.20337900 | -2.39168100 | 2.28970300 |
| H | 3.29818100 | -4.37452500 | 1.15743400 |
| H | 0.43413300 | -5.01268000 | 1.49106300 |
| N | 0.00698200 | -2.77429600 | 2.77331200 |
| C | -0.30852600 | -1.39788800 | 3.17129700 |
| C | 0.48090500 | -0.99975700 | 4.42690200 |
| O | 1.15282400 | 0.01940600 | 4.47845700 |
| C | -1.81123800 | -1.20807600 | 3.33835100 |
| H | -0.77362300 | -3.38631600 | 2.55989600 |
| H | 0.05521200 | -0.71421000 | 2.38805500 |
| H | -2.04368200 | -0.14703900 | 3.50790800 |
| H | -2.21004000 | -1.80259000 | 4.17610800 |
| H | -2.33421400 | -1.50220600 | 2.41762100 |
| N | 0.32636200 | -1.86229200 | 5.46444100 |
| C | 0.88736900 | -1.63235600 | 6.77183100 |
| H | -0.21193800 | -2.70416800 | 5.30060700 |
| H | 1.34202800 | -0.63315100 | 6.76948400 |
| C | 6.66820900 | 4.79828900 | -2.02284500 |
| C | 5.72911700 | 3.66399300 | -1.63946200 |
| O | 5.96109100 | 2.92213600 | -0.68428100 |
| H | 6.13544100 | 5.64369600 | -2.48348300 |
| N | 4.64944000 | 3.47872400 | -2.44213800 |
| C | 3.91425600 | 2.23511200 | -2.43984900 |
| C | 4.81545900 | 1.07731900 | -2.91541100 |
| O | 5.88638400 | 1.25163000 | -3.46077600 |
| C | 2.63568900 | 2.33042900 | -3.29535600 |
| O | 1.63814300 | 3.07339400 | -2.65542100 |
| H | 4.38835800 | 4.18977400 | -3.11930300 |
| H | 3.58498100 | 1.99532100 | -1.41669300 |
| H | 2.26152200 | 1.31255200 | -3.48570900 |
| H | 2.89771300 | 2.77540000 | -4.27540400 |
| H | 1.90463800 | 4.18343800 | -2.82213500 |
| N | 4.30007500 | -0.17320200 | -2.67394000 |
| C | 4.94052600 | -1.32578800 | -3.28693100 |
| H | 3.30980200 | -0.24458700 | -2.44958400 |
| H | 6.01541600 | -1.30312800 | -3.05990600 |
| C | -4.45053300 | 1.94160800 | 4.45996300 |

| | | | |
|---|---|---|---|
| O | -5.03865600 | 0.78500600 | 3.87853000 |
| C | -3.66092600 | 2.83461600 | 3.51170300 |
| O | -2.66385100 | 2.05584100 | 2.85503500 |
| C | -4.42749700 | 3.61551800 | 2.40740400 |
| O | -4.62463500 | 4.93444900 | 2.83355900 |
| C | -3.49059600 | 3.45546900 | 1.15944800 |
| O | -3.17410500 | 4.61935800 | 0.46118000 |
| C | -2.21624700 | 2.81987500 | 1.74968300 |
| N | 0.46145100 | 1.74332800 | -0.68761300 |
| C | -0.00931300 | 0.58096700 | -1.26899900 |
| O | 0.59923500 | -0.03169900 | -2.13495100 |
| N | -1.24583800 | 0.17343500 | -0.79677700 |
| C | -2.03114200 | 0.81214100 | 0.16149900 |
| O | -3.15739000 | 0.37237100 | 0.40125700 |
| C | -1.43590400 | 1.98691000 | 0.77653400 |
| C | -0.24639300 | 2.41513900 | 0.26950600 |
| H | 1.11301700 | 2.29655800 | -1.31822500 |
| H | -1.70543000 | -0.56904300 | -1.33147000 |
| H | 0.19610900 | 3.35900600 | 0.59637100 |
| H | -1.57940600 | 3.65976400 | 2.09141200 |
| H | -4.03943900 | 5.20734900 | 0.42074000 |
| H | -5.40376700 | 3.14287000 | 2.18661900 |
| H | -3.18226200 | 3.62218400 | 4.12690000 |
| H | -5.20626400 | 2.57276400 | 4.96364700 |
| H | -3.96304600 | 2.70462800 | 0.50807900 |
| H | -3.76233600 | 1.56596200 | 5.23324700 |
| P | -5.68872000 | 5.96369200 | 2.01073600 |
| O | -5.35825300 | 5.78149900 | 0.51780300 |
| O | -5.68845600 | 7.26003100 | 2.72349500 |
| O | -7.09683000 | 5.10970200 | 2.22112100 |
| C | 6.08178600 | 0.03919100 | 5.38928400 |
| C | 5.11084300 | -0.80459600 | 4.54904900 |
| C | 4.21899600 | -0.02931400 | 3.57793600 |
| C | 4.00370500 | 1.80290300 | 1.38859500 |
| S | 5.18482100 | 0.69336900 | 2.18273500 |
| C | 5.27033200 | -0.70581900 | 0.98312700 |
| C | 6.69826900 | -1.07163900 | 0.62612500 |
| O | 7.28814300 | -0.01998400 | -0.09369600 |
| C | 7.68338000 | -1.33310600 | 1.76818000 |
| O | 7.44980600 | -2.59537200 | 2.35646000 |
| C | 9.01930400 | -1.22289600 | 1.00609400 |
| O | 9.24910500 | -2.41219400 | 0.28923900 |
| C | 8.72178600 | -0.08464500 | 0.02588600 |
| H | 6.80433300 | 0.58313300 | 4.75841800 |
| H | 5.65500300 | -1.59216900 | 3.99889100 |
| H | 4.41322700 | -1.33921800 | 5.21514300 |
| H | 3.73180500 | 0.83171900 | 4.05851500 |
| H | 3.42802200 | -0.65631300 | 3.13904200 |
| H | 3.80190500 | 2.61710400 | 2.09698200 |
| H | 4.54864300 | 2.18764500 | 0.50486100 |
| H | 3.08253300 | 1.25906600 | 1.13934400 |
| H | 9.17541500 | -0.30106400 | -0.95298500 |
| H | 8.96619900 | -3.14620400 | 0.85682300 |
| H | 7.61312500 | -0.51813300 | 2.51222700 |
| H | 6.63729000 | -2.00090300 | 0.02203500 |
| H | 4.71462100 | -1.54426700 | 1.42560400 |
| H | 7.92175700 | -2.64362000 | 3.19859700 |
| H | 9.86584200 | -0.98261700 | 1.67886500 |
| H | 4.75378600 | -0.36432800 | 0.07317700 |
| H | 2.09983500 | 8.58496900 | -5.60518000 |
| H | 2.65960600 | 6.89117000 | -5.54878900 |
| H | 7.38785000 | 4.40092900 | -2.75635800 |
| H | 7.21572600 | 5.13426200 | -1.13347200 |
| H | 9.07931900 | 0.89263900 | 0.38524500 |
| H | 6.65319500 | -0.59875400 | 6.07998600 |
| H | 5.53935200 | 0.78561500 | 5.99064200 |
| H | 1.66560300 | -2.37357300 | 7.02519000 |
| H | 0.10974400 | -1.66557000 | 7.55203700 |
| H | 1.50107900 | -5.17255400 | 2.89194300 |
| H | 3.77911100 | -7.15304000 | -0.76926400 |
| H | 4.42704400 | -5.49870800 | -0.50677300 |
| H | -0.70018600 | -5.77763300 | -4.33378900 |
| H | -4.88218200 | -7.21559900 | -2.94614800 |
| H | -4.30232100 | -6.63984300 | 1.45278800 |
| H | -5.72907800 | 1.08318200 | 3.26980900 |
| H | -7.43243500 | 4.91692600 | 1.33526200 |
| H | -5.83304400 | 1.37862300 | -2.34753200 |
| H | 4.50006200 | -2.24679500 | -2.87927800 |
| H | 4.82899000 | -1.33328500 | -4.38500200 |

#### Model2 – Intermediate$_1$

| | | | |
|---|---|---|---|
| O | -3.70857200 | -0.74607300 | -3.37827300 |
| H | -4.20359400 | -0.48339600 | -4.16682400 |
| H | -3.99357100 | -1.67834900 | -3.25713100 |
| C | 1.80921100 | 7.46712800 | -5.41131900 |
| C | 1.44491400 | 7.53101100 | -3.92298400 |
| C | 1.15374100 | 6.16549800 | -3.32718700 |
| O | 0.05109100 | 5.87684800 | -2.87914800 |
| O | 2.17541300 | 5.34489800 | -3.34516000 |
| H | 0.98224400 | 7.04259400 | -6.00251000 |
| H | 0.55720800 | 8.15707700 | -3.75827300 |
| H | 2.28332100 | 7.97077100 | -3.35632000 |
| C | -5.60303400 | 2.08279700 | -1.65085200 |
| C | -4.46733800 | 2.93434500 | -2.19126700 |
| N | -3.21146500 | 2.17542500 | -2.30277100 |
| C | -2.01284700 | 2.78318300 | -2.45886300 |
| N | -1.81674600 | 4.00789500 | -1.99430100 |
| N | -1.01036800 | 2.13602200 | -3.07729000 |
| H | -5.37340200 | 1.70952400 | -0.64140100 |
| H | -6.51920900 | 2.68733000 | -1.59137700 |
| H | -4.31554600 | 3.78562300 | -1.52056900 |
| H | -4.73831700 | 3.34048800 | -3.18315300 |
| H | -3.28624900 | 1.19914700 | -2.59774600 |
| H | -2.40455300 | 4.34934800 | -1.20805000 |
| H | -0.99690000 | 4.56098000 | -2.28509100 |
| H | -1.19929500 | 1.27003700 | -3.56555900 |
| H | -0.03646500 | 2.50644800 | -3.01682600 |
| C | -3.66610700 | -5.75686400 | 1.70891000 |
| C | -4.25680600 | -4.64012300 | 2.57059300 |
| C | -5.54715400 | -4.04968200 | 1.98146100 |
| N | -5.34721800 | -3.38853300 | 0.70643200 |
| C | -4.89604900 | -2.13571100 | 0.57340300 |
| N | -4.78206500 | -1.29963300 | 1.62643800 |
| N | -4.58804200 | -1.69840800 | -0.64488300 |
| H | -3.50377200 | -5.42116600 | 0.67364000 |
| H | -2.70540200 | -6.11413800 | 2.11289500 |
| H | -3.52393700 | -3.82160000 | 2.69503800 |
| H | -4.47621800 | -5.01489500 | 3.58564400 |
| H | -6.02069400 | -3.35535500 | 2.69563100 |
| H | -6.27876000 | -4.85682400 | 1.81879100 |
| H | -5.24004000 | -4.00266100 | -0.14370600 |
| H | -4.94543200 | -1.60884600 | 2.57326600 |
| H | -4.24219800 | -0.44012900 | 1.51951500 |
| H | -4.74108700 | -2.31264800 | -1.47724900 |
| H | -4.12155900 | -0.80386000 | -0.74478300 |
| C | -4.91264200 | -6.79118200 | -3.81767500 |
| C | -3.47865500 | -6.44722600 | -4.30783100 |
| O | -3.13838200 | -6.60979300 | -5.46906100 |
| C | -5.69299100 | -5.45768600 | -3.68882100 |
| C | -5.16718100 | -4.50296200 | -2.59886600 |
| O | -5.00888500 | -3.28175000 | -2.88463700 |
| O | -4.94389300 | -5.00174200 | -1.45840600 |
| H | -5.36627900 | -7.44207900 | -4.57466200 |
| H | -6.74367500 | -5.67520600 | -3.42848100 |
| H | -5.69502300 | -4.92628600 | -4.65133900 |
| N | -2.65473900 | -5.85264800 | -3.38466200 |
| C | -1.38315400 | -5.18562100 | -3.71482500 |
| C | -0.75576700 | -4.58181300 | -2.45284000 |
| C | -1.63645900 | -3.47153800 | -1.84709800 |
| C | 0.66194300 | -4.08210500 | -2.74488800 |

| | | | |
|---|---|---|---|
| C | -1.32599800 | -3.17722300 | -0.38121300 |
| H | -3.03309600 | -5.71495100 | -2.45186500 |
| H | -1.56214600 | -4.40233100 | -4.47466900 |
| H | -0.66892300 | -5.39680300 | -1.71186400 |
| H | -2.69797000 | -3.74290400 | -1.91679100 |
| H | -1.53499000 | -2.55134300 | -2.45156200 |
| H | 1.12922400 | -3.64403700 | -1.84896600 |
| H | 0.65253700 | -3.30250500 | -3.52553000 |
| H | 1.30695600 | -4.90455200 | -3.09096000 |
| H | -1.49664700 | -4.07802300 | 0.23069300 |
| H | -1.98405200 | -2.38697200 | 0.01197300 |
| H | -0.27693900 | -2.87312100 | -0.23007500 |
| C | 3.53831300 | -6.10255200 | -0.74255200 |
| C | 2.25465300 | -5.83105500 | 0.02740200 |
| O | 1.21055900 | -6.40404600 | -0.23610700 |
| H | 3.41191300 | -5.73658100 | -1.77292900 |
| N | 2.33561300 | -4.90583100 | 1.03701000 |
| C | 1.20224400 | -4.61786100 | 1.87533800 |
| C | 1.15501500 | -3.15433300 | 2.30669000 |
| O | 2.15421600 | -2.44231200 | 2.30128300 |
| H | 3.19799200 | -4.41299500 | 1.23518100 |
| H | 0.29591100 | -4.91332900 | 1.32843200 |
| N | -0.05704000 | -2.74230100 | 2.76898900 |
| C | -0.32240600 | -1.35832500 | 3.16869700 |
| C | 0.49913400 | -0.96002100 | 4.40355400 |
| O | 1.17692700 | 0.05632500 | 4.43086600 |
| C | -1.81407100 | -1.14023100 | 3.39404300 |
| H | -0.86095700 | -3.32395100 | 2.55615100 |
| H | 0.03020300 | -0.68521300 | 2.37269400 |
| H | -2.02172400 | -0.07300700 | 3.55457100 |
| H | -2.18391400 | -1.71514900 | 4.25814900 |
| H | -2.37488300 | -1.43708600 | 2.49875300 |
| N | 0.34786600 | -1.80170600 | 5.45888900 |
| C | 0.87798300 | -1.52606000 | 6.77140800 |
| H | -0.21073600 | -2.63466600 | 5.31876900 |
| H | 1.36009400 | -0.54044800 | 6.73627800 |
| C | 6.63586900 | 4.75857100 | -2.14311300 |
| C | 5.66133300 | 3.68628700 | -1.67857500 |
| O | 5.80911200 | 3.08433800 | -0.61634900 |
| H | 6.12950300 | 5.56375400 | -2.69639900 |
| N | 4.63360500 | 3.41824100 | -2.52633800 |
| C | 3.89743500 | 2.17542400 | -2.46212100 |
| C | 4.77781100 | 1.01527200 | -2.96475800 |
| O | 5.81188900 | 1.20212400 | -3.58137600 |
| C | 2.57259300 | 2.26712200 | -3.24922600 |
| O | 1.61873700 | 3.04182700 | -2.59433900 |
| H | 4.60741100 | 3.91131200 | -3.41164000 |
| H | 3.63055900 | 1.97457600 | -1.41750600 |
| H | 2.20518900 | 1.23790200 | -3.40997200 |
| H | 2.80011700 | 2.67972100 | -4.25587100 |
| H | 1.89598200 | 4.39853300 | -2.95187100 |
| N | 4.29711400 | -0.23159800 | -2.67777500 |
| C | 4.91941200 | -1.39085000 | -3.29588500 |
| H | 3.35370400 | -0.30903500 | -2.29665300 |
| H | 6.00784900 | -1.34290000 | -3.14987100 |
| C | -4.47941800 | 1.99620400 | 4.40943400 |
| O | -5.05626600 | 0.82720800 | 3.84228100 |
| C | -3.69168300 | 2.89335100 | 3.46355000 |
| O | -2.63511800 | 2.15397900 | 2.85690700 |
| C | -4.45014100 | 3.62838600 | 2.32103200 |
| O | -4.63513700 | 4.96367600 | 2.70457400 |
| C | -3.49750200 | 3.42301700 | 1.09157600 |
| O | -3.25630400 | 4.54629600 | 0.30429800 |
| C | -2.19666300 | 2.92179800 | 1.74601400 |
| N | 0.76962900 | 1.99731600 | -0.42480400 |
| C | 0.61233600 | 0.65701100 | -0.69229200 |
| O | 1.43939700 | -0.01892100 | -1.29746400 |
| N | -0.57419300 | 0.11643700 | -0.22938700 |
| C | -1.61059300 | 0.78739800 | 0.42849900 |
| O | -2.69059100 | 0.21131400 | 0.56997100 |
| C | -1.26809300 | 2.12430800 | 0.87557700 |
| C | -0.12851500 | 2.67060600 | 0.35387400 |
| H | 1.30510400 | 2.53218800 | -1.24305800 |
| H | -0.75576600 | -0.84961200 | -0.49067600 |
| H | 0.11048000 | 3.72270200 | 0.52948100 |
| H | -1.65987300 | 3.82746000 | 2.09174900 |
| H | -4.12911400 | 5.12846800 | 0.27627800 |
| H | -5.42988400 | 3.15833800 | 2.11068600 |
| H | -3.26941700 | 3.71163200 | 4.07935700 |
| H | -5.24450600 | 2.62613700 | 4.90070900 |
| H | -3.90639800 | 2.58485100 | 0.50064600 |
| H | -3.79295800 | 1.63732300 | 5.19220900 |
| P | -5.71899700 | 5.97034200 | 1.88311700 |
| O | -5.42944200 | 5.73158000 | 0.38944900 |
| O | -5.69241100 | 7.28987500 | 2.55146000 |
| O | -7.12533700 | 5.13350500 | 2.16195200 |
| C | 6.06741100 | 0.13114000 | 5.35333800 |
| C | 5.13444300 | -0.76436000 | 4.52282000 |
| C | 4.20437000 | -0.04220800 | 3.54693200 |
| C | 3.87518700 | 1.81083000 | 1.39918500 |
| S | 5.11917500 | 0.73326500 | 2.14457900 |
| C | 5.21774200 | -0.63463200 | 0.90966500 |
| C | 6.64371800 | -1.00914600 | 0.55690700 |
| O | 7.27146900 | 0.05336900 | -0.11539800 |
| C | 7.60942100 | -1.33327100 | 1.69899900 |
| O | 7.33201900 | -2.60206400 | 2.25366900 |
| C | 8.95475200 | -1.24968800 | 0.95019600 |
| O | 9.15531800 | -2.43140600 | 0.21217600 |
| C | 8.70171300 | -0.08296400 | -0.01001800 |
| H | 6.76387900 | 0.70042900 | 4.71564800 |
| H | 5.70925900 | -1.53355800 | 3.97863600 |
| H | 4.46369000 | -1.32298700 | 5.19702300 |
| H | 3.66768900 | 0.79006200 | 4.02494300 |
| H | 3.44662500 | -0.71113100 | 3.11166900 |
| H | 3.61877500 | 2.57166300 | 2.14763600 |
| H | 4.39843400 | 2.27153700 | 0.54094400 |
| H | 2.99229600 | 1.22683900 | 1.11010400 |
| H | 9.13759200 | -0.30736400 | -0.99546000 |
| H | 8.83568700 | -3.16615000 | 0.75900800 |
| H | 7.55818700 | -0.53470800 | 2.46242700 |
| H | 6.57150600 | -1.91338800 | -0.08284300 |
| H | 4.65420800 | -1.48252200 | 1.32319200 |
| H | 7.80053500 | -2.68826600 | 3.09470200 |
| H | 9.80229100 | -1.04937600 | 1.63465300 |
| H | 4.70835700 | -0.26925400 | 0.00532800 |
| H | 2.02393100 | 8.47235800 | -5.80491800 |
| H | 2.69722900 | 6.83656300 | -5.56847000 |
| H | 7.36942900 | 4.28367000 | -2.81443800 |
| H | 7.16241300 | 5.17168800 | -1.27402300 |
| H | 9.10866000 | 0.87040800 | 0.36058800 |
| H | 6.66768100 | -0.47305700 | 6.04988200 |
| H | 5.49148600 | 0.85836900 | 5.94707700 |
| H | 1.62730400 | -2.27523300 | 7.08069200 |
| H | 0.07785400 | -1.50167900 | 7.52923400 |
| H | 1.21571300 | -5.22287100 | 2.80489000 |
| H | 3.68519100 | -7.19023400 | -0.79632400 |
| H | 4.42750400 | -5.63090900 | -0.29906800 |
| H | -0.69791000 | -5.91715300 | -4.17299000 |
| H | -4.89490000 | -7.29831800 | -2.84091000 |
| H | -4.34956300 | -6.61942400 | 1.66021000 |
| H | -5.73807900 | 1.10922500 | 3.21646900 |
| H | -7.47823100 | 4.90076200 | 1.29252500 |
| H | -5.79943000 | 1.20714600 | -2.29174200 |
| H | 4.52771000 | -2.30630300 | -2.82988500 |
| H | 4.72902000 | -1.43708900 | -4.38218000 |

#### Model2 $-$ $TS1_2$

| | | | |
|---|---|---|---|
| O | -3.70085900 | -0.74084800 | -3.38382700 |
| H | -4.19731900 | -0.48316100 | -4.17313900 |
| H | -3.98754100 | -1.67162400 | -3.25547700 |

| | | | |
|---|---|---|---|
| C | 1.80289200 | 7.46206100 | -5.41644600 |
| C | 1.45397100 | 7.55046200 | -3.92491000 |
| C | 1.15837600 | 6.19442400 | -3.31463600 |
| O | 0.04311700 | 5.88123000 | -2.92592200 |
| O | 2.20460500 | 5.39774100 | -3.25853800 |
| H | 0.96842500 | 7.03064500 | -5.99165400 |
| H | 0.56943300 | 8.18190000 | -3.76422900 |
| H | 2.29959900 | 7.99290700 | -3.37233000 |
| C | -5.60629300 | 2.08364600 | -1.64150900 |
| C | -4.47944800 | 2.94194400 | -2.18508800 |
| N | -3.22590700 | 2.18303000 | -2.30818400 |
| C | -2.02879600 | 2.78858600 | -2.46249000 |
| N | -1.83168500 | 4.01452100 | -2.00153700 |
| N | -1.02846100 | 2.13984700 | -3.08407700 |
| H | -5.36650200 | 1.70431800 | -0.63677600 |
| H | -6.52511300 | 2.68298300 | -1.57055700 |
| H | -4.32426000 | 3.79037800 | -1.51152300 |
| H | -4.75711400 | 3.35211400 | -3.17345600 |
| H | -3.30191700 | 1.20433600 | -2.59507900 |
| H | -2.41141300 | 4.35434600 | -1.20676400 |
| H | -1.01031700 | 4.56112700 | -2.29261600 |
| H | -1.21221500 | 1.26192400 | -3.55239800 |
| H | -0.05933600 | 2.50319200 | -3.00846900 |
| C | -3.66877300 | -5.75673700 | 1.71622600 |
| C | -4.24810900 | -4.63092700 | 2.57406100 |
| C | -5.53850000 | -4.03558600 | 1.98949200 |
| N | -5.34209300 | -3.37848800 | 0.71183200 |
| C | -4.88530400 | -2.12801300 | 0.57379700 |
| N | -4.76293500 | -1.28878000 | 1.62314500 |
| N | -4.58020900 | -1.69421800 | -0.64655000 |
| H | -3.51113800 | -5.42871900 | 0.67772400 |
| H | -2.70737900 | -6.11624200 | 2.11654700 |
| H | -3.50976300 | -3.81601900 | 2.68916300 |
| H | -4.46372200 | -4.99813600 | 3.59271000 |
| H | -6.00444400 | -3.33684400 | 2.70431800 |
| H | -6.27515500 | -4.83942200 | 1.83315200 |
| H | -5.23963000 | -3.99457300 | -0.13705000 |
| H | -4.92383200 | -1.59235400 | 2.57222000 |
| H | -4.21810100 | -0.43281400 | 1.50784800 |
| H | -4.73551000 | -2.30999100 | -1.47673900 |
| H | -4.09690000 | -0.80788300 | -0.74471200 |
| C | -4.92606100 | -6.79122500 | -3.80789100 |
| C | -3.49149600 | -6.45110400 | -4.29961900 |
| O | -3.15200300 | -6.61782100 | -5.46055700 |
| C | -5.70318700 | -5.45543700 | -3.68172200 |
| C | -5.17197400 | -4.49840500 | -2.59606800 |
| O | -5.00905200 | -3.27909200 | -2.88661600 |
| O | -4.94851100 | -4.99459100 | -1.45440900 |
| H | -5.38144000 | -7.44299700 | -4.56308400 |
| H | -6.75375000 | -5.66967500 | -3.41822700 |
| H | -5.70602900 | -4.92724500 | -4.64601500 |
| N | -2.66640600 | -5.85462700 | -3.37870000 |
| C | -1.39526600 | -5.18812300 | -3.71175200 |
| C | -0.76334000 | -4.58620000 | -2.45109300 |
| C | -1.64008900 | -3.47449700 | -1.84219700 |
| C | 0.65437800 | -4.08874800 | -2.74714000 |
| C | -1.32474900 | -3.18083900 | -0.37720600 |
| H | -3.04504900 | -5.71210900 | -2.44669200 |
| H | -1.57573200 | -4.40352300 | -4.46999300 |
| H | -0.67580000 | -5.40170300 | -1.71077900 |
| H | -2.70231300 | -3.74383700 | -1.90889300 |
| H | -1.53855500 | -2.55445500 | -2.44699900 |
| H | 1.12417100 | -3.64974800 | -1.85300900 |
| H | 0.64393500 | -3.31017200 | -3.52884700 |
| H | 1.29752500 | -4.91244400 | -3.09380100 |
| H | -1.49884100 | -4.08070600 | 0.23516500 |
| H | -1.97685100 | -2.38615700 | 0.01660000 |
| H | -0.27411100 | -2.88118300 | -0.22880500 |
| C | 3.53094200 | -6.10794700 | -0.74824000 |
| C | 2.24828900 | -5.83781900 | 0.02448300 |
| O | 1.20516300 | -6.41406300 | -0.23583200 |
| H | 3.40024300 | -5.74627100 | -1.77957000 |
| N | 2.32909900 | -4.91080500 | 1.03227700 |
| C | 1.19660000 | -4.62323600 | 1.87230400 |
| C | 1.15336500 | -3.16055400 | 2.30756400 |
| O | 2.15520000 | -2.45211400 | 2.30365200 |
| H | 3.19024100 | -4.41463300 | 1.22767500 |
| H | 0.28900200 | -4.91519500 | 1.32575700 |
| N | -0.05717800 | -2.74576100 | 2.77113800 |
| C | -0.31934300 | -1.36027100 | 3.16910500 |
| C | 0.50330700 | -0.96238700 | 4.40306900 |
| O | 1.18248100 | 0.05311400 | 4.42966800 |
| C | -1.81023100 | -1.13675800 | 3.39178500 |
| H | -0.86282100 | -3.32403900 | 2.55556000 |
| H | 0.03505200 | -0.68942900 | 2.37185600 |
| H | -2.01385200 | -0.06851300 | 3.55068400 |
| H | -2.18453300 | -1.71001100 | 4.25512900 |
| H | -2.36947200 | -1.43125100 | 2.49478700 |
| N | 0.35242700 | -1.80330100 | 5.45912800 |
| C | 0.88746500 | -1.52816700 | 6.76966400 |
| H | -0.20716500 | -2.63581800 | 5.32033100 |
| H | 1.36807000 | -0.54186600 | 6.73341600 |
| C | 6.63357100 | 4.75073400 | -2.15648800 |
| C | 5.65287300 | 3.69310500 | -1.67424500 |
| O | 5.78111300 | 3.12196100 | -0.59381500 |
| H | 6.12850600 | 5.55196500 | -2.71670000 |
| N | 4.63666700 | 3.40711000 | -2.53218500 |
| C | 3.89796500 | 2.16758900 | -2.45053400 |
| C | 4.76486800 | 1.00472600 | -2.97092300 |
| O | 5.79279400 | 1.19366000 | -3.59823200 |
| C | 2.55941900 | 2.26305500 | -3.20772100 |
| O | 1.61939200 | 3.04547400 | -2.53100900 |
| H | 4.65543600 | 3.85342200 | -3.44215400 |
| H | 3.65466500 | 1.97169500 | -1.39986400 |
| H | 2.18060400 | 1.23830400 | -3.36176000 |
| H | 2.75658200 | 2.68493000 | -4.21470100 |
| H | 1.92776900 | 4.47502400 | -2.87442000 |
| N | 4.28423500 | -0.24050700 | -2.68718200 |
| C | 4.91071100 | -1.39769800 | -3.30497200 |
| H | 3.34995000 | -0.32123300 | -2.28305500 |
| H | 5.99523300 | -1.37071900 | -3.12528700 |
| C | -4.46436100 | 1.99824100 | 4.40793800 |
| O | -5.05043800 | 0.82871200 | 3.85086200 |
| C | -3.67747200 | 2.88925100 | 3.45485300 |
| O | -2.62740400 | 2.14618100 | 2.84291600 |
| C | -4.43880200 | 3.62707600 | 2.31616500 |
| O | -4.62672200 | 4.96082900 | 2.70544700 |
| C | -3.48704300 | 3.42513400 | 1.08498400 |
| O | -3.24808900 | 4.55314300 | 0.30196500 |
| C | -2.18517800 | 2.91650200 | 1.73319100 |
| N | 0.80368400 | 1.99524400 | -0.41966700 |
| C | 0.63579400 | 0.65971200 | -0.68655300 |
| O | 1.46587900 | -0.02766100 | -1.28244700 |
| N | -0.56316100 | 0.11988000 | -0.24956200 |
| C | -1.60093600 | 0.78793300 | 0.40698400 |
| O | -2.68491900 | 0.21186100 | 0.53909500 |
| C | -1.25620800 | 2.11847700 | 0.86224200 |
| C | -0.10135400 | 2.65695000 | 0.35636300 |
| H | 1.33704600 | 2.56229800 | -1.33673300 |
| H | -0.74478300 | -0.84365400 | -0.51887300 |
| H | 0.13750900 | 3.70690700 | 0.54984600 |
| H | -1.64546300 | 3.81874100 | 2.08356900 |
| H | -4.11950200 | 5.13684100 | 0.28065000 |
| H | -5.41793800 | 3.15579900 | 2.10546700 |
| H | -3.24901500 | 3.70672100 | 4.06773500 |
| H | -5.22376700 | 2.63424400 | 4.90036500 |
| H | -3.89845800 | 2.59090600 | 0.49062100 |
| H | -3.77542400 | 1.63989000 | 5.18878300 |
| P | -5.71311400 | 5.97194000 | 1.89192500 |
| O | -5.42045200 | 5.74486900 | 0.39695200 |

| | | | |
|---|---|---|---|
| O | -5.68993500 | 7.28627900 | 2.57068100 |
| O | -7.11868500 | 5.13132300 | 2.16265100 |
| C | 6.07547200 | 0.12512100 | 5.34186300 |
| C | 5.14352700 | -0.77568900 | 4.51603300 |
| C | 4.20457100 | -0.05905600 | 3.54493300 |
| C | 3.84778400 | 1.79893700 | 1.40820700 |
| S | 5.10717700 | 0.73011600 | 2.14226300 |
| C | 5.21204700 | -0.63090700 | 0.90003700 |
| C | 6.63768400 | -1.00589900 | 0.54562300 |
| O | 7.26965200 | 0.05572400 | -0.12513700 |
| C | 7.60373700 | -1.33624500 | 1.68553000 |
| O | 7.32162400 | -2.60463800 | 2.23873600 |
| C | 8.94827900 | -1.25936200 | 0.93428100 |
| O | 9.14185700 | -2.44231100 | 0.19639300 |
| C | 8.69988500 | -0.09184600 | -0.02622300 |
| H | 6.76600700 | 0.69784600 | 4.70080900 |
| H | 5.71959900 | -1.54180200 | 3.96904300 |
| H | 4.47924100 | -1.33796000 | 5.19364200 |
| H | 3.66140700 | 0.76673800 | 4.02678400 |
| H | 3.45127500 | -0.73292700 | 3.10952700 |
| H | 3.57522700 | 2.54278800 | 2.16784800 |
| H | 4.36581100 | 2.28214500 | 0.56033700 |
| H | 2.97548400 | 1.20667800 | 1.10451600 |
| H | 9.12948700 | -0.32116700 | -1.01331200 |
| H | 8.81876500 | -3.17512900 | 0.74382700 |
| H | 7.55752900 | -0.53842600 | 2.45002200 |
| H | 6.56270000 | -1.90830700 | -0.09640700 |
| H | 4.64844800 | -1.48146400 | 1.30802600 |
| H | 7.78933100 | -2.69350700 | 3.07993800 |
| H | 9.79788800 | -1.06307700 | 1.61730200 |
| H | 4.70190000 | -0.26191100 | -0.00215000 |
| H | 2.01536800 | 8.46087800 | -5.82681200 |
| H | 2.68871600 | 6.82824600 | -5.57382000 |
| H | 7.36075200 | 4.26352200 | -2.82602000 |
| H | 7.16658000 | 5.17178200 | -1.29527400 |
| H | 9.11571800 | 0.85884200 | 0.34129500 |
| H | 6.68208800 | -0.47543200 | 6.03608100 |
| H | 5.49835900 | 0.84963500 | 5.93777400 |
| H | 1.63911700 | -2.27654500 | 7.07542000 |
| H | 0.09039900 | -1.50558500 | 7.53076400 |
| H | 1.20994500 | -5.23085700 | 2.80014100 |
| H | 3.68120200 | -7.19535300 | -0.79827800 |
| H | 4.41985400 | -5.63174800 | -0.30903900 |
| H | -0.71236900 | -5.91985200 | -4.17303900 |
| H | -4.90872800 | -7.29590300 | -2.82985500 |
| H | -4.35709600 | -6.61597100 | 1.67786600 |
| H | -5.72806000 | 1.11058200 | 3.22042200 |
| H | -7.46538900 | 4.89729700 | 1.29109600 |
| H | -5.80434800 | 1.21116600 | -2.28625700 |
| H | 4.49019500 | -2.31437100 | -2.86788800 |
| H | 4.75402000 | -1.42088400 | -4.39706500 |

#### Model2 – Intermediate$_2$

| | | | |
|---|---|---|---|
| O | -3.61801600 | -0.75836400 | -3.42532700 |
| H | -4.07040700 | -0.52424700 | -4.24781200 |
| H | -3.93260500 | -1.67635700 | -3.27200200 |
| C | 1.81120900 | 7.41700000 | -5.46813900 |
| C | 1.39314500 | 7.57185600 | -4.00036600 |
| C | 1.17702800 | 6.23532400 | -3.32777200 |
| O | 0.08912100 | 5.84175700 | -2.95127900 |
| O | 2.29161000 | 5.52306700 | -3.21014300 |
| H | 1.03913600 | 6.88265500 | -6.04393000 |
| H | 0.45646100 | 8.13853100 | -3.91169500 |
| H | 2.17809500 | 8.11054800 | -3.44406100 |
| C | -5.61022600 | 2.07584000 | -1.65905400 |
| C | -4.48738700 | 2.94501100 | -2.18401800 |
| N | -3.24840700 | 2.17188000 | -2.33104200 |
| C | -2.04313000 | 2.75216000 | -2.46025000 |
| N | -1.82561700 | 3.97306500 | -1.99570100 |
| N | -1.05483500 | 2.08179900 | -3.08455000 |
| H | -5.36189000 | 1.66665200 | -0.66823600 |
| H | -6.52911200 | 2.67191700 | -1.56425400 |
| H | -4.32191100 | 3.77592000 | -1.49108100 |
| H | -4.76382500 | 3.38152100 | -3.16137100 |
| H | -3.33714700 | 1.18946800 | -2.60181400 |
| H | -2.39584600 | 4.31977700 | -1.19128500 |
| H | -0.98458300 | 4.49166900 | -2.26476200 |
| H | -1.24055100 | 1.17655200 | -3.49686700 |
| H | -0.09298800 | 2.41861600 | -2.98736700 |
| C | -3.68736600 | -5.74691300 | 1.74712400 |
| C | -4.23081200 | -4.58457200 | 2.58024300 |
| C | -5.51406800 | -3.97109200 | 1.99729000 |
| N | -5.31783200 | -3.33683700 | 0.70820300 |
| C | -4.84725200 | -2.09393900 | 0.54636300 |
| N | -4.70547400 | -1.23624000 | 1.57793100 |
| N | -4.54610700 | -1.68039400 | -0.68230800 |
| H | -3.53308100 | -5.45005100 | 0.69857800 |
| H | -2.73013600 | -6.11764900 | 2.14704000 |
| H | -3.47226900 | -3.78482900 | 2.66572100 |
| H | -4.44282800 | -4.91989900 | 3.61064300 |
| H | -5.95687100 | -3.25150400 | 2.70595400 |
| H | -6.26933800 | -4.76153300 | 1.86196800 |
| H | -5.22782000 | -3.96509000 | -0.13214600 |
| H | -4.86071200 | -1.51571200 | 2.53543200 |
| H | -4.14940300 | -0.38989300 | 1.43156600 |
| H | -4.70604200 | -2.30769800 | -1.50101900 |
| H | -4.01889100 | -0.81723900 | -0.78102400 |
| C | -4.94668200 | -6.81337500 | -3.77051200 |
| C | -3.50738600 | -6.48772200 | -4.26033900 |
| O | -3.16268100 | -6.67580600 | -5.41665600 |
| C | -5.71429000 | -5.46965000 | -3.66591100 |
| C | -5.17033100 | -4.49747100 | -2.59905100 |
| O | -4.98980100 | -3.28718600 | -2.91358200 |
| O | -4.95223300 | -4.97649900 | -1.44861400 |
| H | -5.40287700 | -7.47248700 | -4.51879500 |
| H | -6.76551200 | -5.67139600 | -3.39543600 |
| H | -5.71562200 | -4.95878900 | -4.63953700 |
| N | -2.68449400 | -5.87886100 | -3.34535200 |
| C | -1.41267000 | -5.21658100 | -3.68423400 |
| C | -0.77353400 | -4.61170100 | -2.42806200 |
| C | -1.63983700 | -3.48877300 | -1.82479600 |
| C | 0.64733900 | -4.12752200 | -2.73152600 |
| C | -1.30857700 | -3.17066100 | -0.36814400 |
| H | -3.06977700 | -5.71500400 | -2.41941100 |
| H | -1.59284100 | -4.43316000 | -4.44407000 |
| H | -0.69044400 | -5.42294600 | -1.68253500 |
| H | -2.70311100 | -3.75778800 | -1.87509800 |
| H | -1.54450400 | -2.57947900 | -2.44701800 |
| H | 1.12299500 | -3.68651500 | -1.84157000 |
| H | 0.64020200 | -3.35404700 | -3.51857600 |
| H | 1.28279300 | -4.95843900 | -3.07534800 |
| H | -1.48688400 | -4.05878000 | 0.26024300 |
| H | -1.94827300 | -2.36137800 | 0.01525300 |
| H | -0.25582000 | -2.87352900 | -0.23267900 |
| C | 3.51169000 | -6.12768500 | -0.71498800 |
| C | 2.22798300 | -5.85757500 | 0.05747100 |
| O | 1.19019500 | -6.44780100 | -0.19318800 |
| H | 3.36997200 | -5.79496900 | -1.75446200 |
| N | 2.30163200 | -4.91661900 | 1.05208200 |
| C | 1.16701000 | -4.62676500 | 1.88952800 |
| C | 1.13340400 | -3.16474800 | 2.32920500 |
| O | 2.14229100 | -2.46637700 | 2.33032000 |
| H | 3.15675700 | -4.40668400 | 1.23930200 |
| H | 0.25869000 | -4.91113500 | 1.34052100 |
| N | -0.07486600 | -2.74018300 | 2.78987200 |
| C | -0.32921500 | -1.34813200 | 3.17283000 |
| C | 0.48936400 | -0.94364500 | 4.40649700 |
| O | 1.17075000 | 0.07056300 | 4.42922100 |
| C | -1.81901100 | -1.10731900 | 3.37716700 |
| H | -0.88433500 | -3.30959100 | 2.56501700 |

| | | | |
|---|---|---|---|
| H | 0.03571600 | -0.68872200 | 2.37034700 |
| H | -2.01389100 | -0.03609500 | 3.52812200 |
| H | -2.21189900 | -1.67321600 | 4.23725900 |
| H | -2.36795800 | -1.40053400 | 2.47339800 |
| N | 0.33448000 | -1.77668600 | 5.46834000 |
| C | 0.87718300 | -1.49591500 | 6.77416100 |
| H | -0.22371200 | -2.61085000 | 5.33386200 |
| H | 1.35037800 | -0.50625500 | 6.73271900 |
| C | 6.63585700 | 4.71609200 | -2.19076300 |
| C | 5.62526900 | 3.70609200 | -1.68240800 |
| O | 5.67939700 | 3.22535200 | -0.55668200 |
| H | 6.15415000 | 5.50183500 | -2.79209900 |
| N | 4.65632500 | 3.36328400 | -2.58233800 |
| C | 3.91632800 | 2.13083600 | -2.44946700 |
| C | 4.75747100 | 0.95608100 | -2.98171900 |
| O | 5.77192800 | 1.14218500 | -3.63392600 |
| C | 2.56647500 | 2.20653300 | -3.16740500 |
| O | 1.65986200 | 3.06502600 | -2.49981800 |
| H | 4.81118000 | 3.66789000 | -3.53761000 |
| H | 3.70787200 | 1.96309000 | -1.38625900 |
| H | 2.14140300 | 1.19540400 | -3.24005900 |
| H | 2.70450100 | 2.59766500 | -4.18959100 |
| H | 2.06185500 | 4.63258200 | -2.81461200 |
| N | 4.27043500 | -0.27815400 | -2.69283700 |
| C | 4.90088000 | -1.43654900 | -3.30107300 |
| H | 3.36623100 | -0.35840500 | -2.22041700 |
| H | 5.96829500 | -1.47323300 | -3.03444800 |
| C | -4.49142300 | 2.03403700 | 4.39609500 |
| O | -5.05603100 | 0.85445700 | 3.84070800 |
| C | -3.68656800 | 2.90652400 | 3.44250000 |
| O | -2.64740800 | 2.14121700 | 2.84393700 |
| C | -4.43314400 | 3.63922100 | 2.29131600 |
| O | -4.62986800 | 4.97471300 | 2.67402000 |
| C | -3.46516800 | 3.42546100 | 1.07374400 |
| O | -3.21407700 | 4.55445600 | 0.29088700 |
| C | -2.17437600 | 2.90353200 | 1.73738700 |
| N | 0.96628900 | 1.95754400 | -0.18144500 |
| C | 0.72657600 | 0.68182400 | -0.59305700 |
| O | 1.55426200 | -0.01487700 | -1.20379900 |
| N | -0.53855500 | 0.16922700 | -0.31687200 |
| C | -1.58505300 | 0.81751600 | 0.34267400 |
| O | -2.69294800 | 0.25873700 | 0.39250900 |
| C | -1.22603300 | 2.09800700 | 0.89539300 |
| C | 0.02097000 | 2.58390900 | 0.55696300 |
| H | 1.53506900 | 2.73958600 | -1.54314400 |
| H | -0.74094700 | -0.75659900 | -0.68369600 |
| H | 0.28834600 | 3.59285300 | 0.89287100 |
| H | -1.63458300 | 3.80123900 | 2.09993000 |
| H | -4.07957700 | 5.14350900 | 0.26421100 |
| H | -5.40925900 | 3.16572200 | 2.07085200 |
| H | -3.25067500 | 3.72605600 | 4.04773000 |
| H | -5.26515400 | 2.67513700 | 4.85920200 |
| H | -3.87725800 | 2.59559200 | 0.47434200 |
| H | -3.82049300 | 1.69052500 | 5.19896600 |
| P | -5.70867200 | 5.98671700 | 1.84962700 |
| O | -5.38581600 | 5.76671300 | 0.35952100 |
| O | -5.70268500 | 7.29768900 | 2.53557700 |
| O | -7.11769900 | 5.14324900 | 2.09070300 |
| C | 6.06842600 | 0.13818400 | 5.33620800 |
| C | 5.14010300 | -0.77900500 | 4.52553100 |
| C | 4.19094900 | -0.07511100 | 3.55673600 |
| C | 3.82268700 | 1.84538300 | 1.48542500 |
| S | 5.08681300 | 0.74878900 | 2.17039600 |
| C | 5.17912900 | -0.57518100 | 0.88814700 |
| C | 6.59719300 | -0.96850200 | 0.52079900 |
| O | 7.26706200 | 0.09476000 | -0.11282000 |
| C | 7.55021200 | -1.36686800 | 1.64951400 |
| O | 7.22314000 | -2.64030900 | 2.16465700 |
| C | 8.89731100 | -1.31788500 | 0.89980900 |
| O | 9.04925300 | -2.48724500 | 0.13122000 |
| C | 8.69248600 | -0.11740000 | -0.03036300 |
| H | 6.74980500 | 0.71036400 | 4.68513200 |
| H | 5.71862800 | -1.54327700 | 3.97871600 |
| H | 4.48591000 | -1.34282200 | 5.21166500 |
| H | 3.62698500 | 0.73381700 | 4.04310300 |
| H | 3.45287600 | -0.75857400 | 3.11021900 |
| H | 3.55818800 | 2.55676300 | 2.27833500 |
| H | 4.32834300 | 2.36399100 | 0.65419400 |
| H | 2.94208700 | 1.27864300 | 1.15824200 |
| H | 9.10379700 | -0.34185500 | -1.02638700 |
| H | 8.69784100 | -3.22151500 | 0.65908300 |
| H | 7.53115400 | -0.59063200 | 2.43695600 |
| H | 6.49920400 | -1.84532400 | -0.15274600 |
| H | 4.60647200 | -1.43118900 | 1.27161200 |
| H | 7.68042500 | -2.76818600 | 3.00650800 |
| H | 9.75295500 | -1.17019300 | 1.58743300 |
| H | 4.66571700 | -0.17724300 | 0.00141000 |
| H | 1.95908100 | 8.40200700 | -5.93576900 |
| H | 2.75162100 | 6.85170600 | -5.55005900 |
| H | 7.36396800 | 4.18866400 | -2.82837200 |
| H | 7.16268900 | 5.16410300 | -1.33970100 |
| H | 9.15159100 | 0.80527200 | 0.35629800 |
| H | 6.68482900 | -0.44858800 | 6.03363700 |
| H | 5.48767400 | 0.86364900 | 5.92750300 |
| H | 1.63674800 | -2.23816300 | 7.07604400 |
| H | 0.08569000 | -1.47802700 | 7.54115900 |
| H | 1.17477600 | -5.23810700 | 2.81494800 |
| H | 3.67848600 | -7.21366400 | -0.73763400 |
| H | 4.39532800 | -5.62689400 | -0.29266100 |
| H | -0.73450200 | -5.95233500 | -4.14589900 |
| H | -4.93634800 | -7.30422600 | -2.78539700 |
| H | -4.39606200 | -6.59033300 | 1.73967000 |
| H | -5.70135200 | 1.12335800 | 3.17187200 |
| H | -7.43462800 | 4.89190200 | 1.21271300 |
| H | -5.81418700 | 1.22239200 | -2.32714900 |
| H | 4.40397900 | -2.34818500 | -2.94176100 |
| H | 4.83636000 | -1.40400600 | -4.40121500 |

#### Model2 – TS2

| | | | |
|---|---|---|---|
| O | -3.67265900 | -0.78279400 | -3.43045100 |
| H | -4.16348400 | -0.55536900 | -4.23265800 |
| H | -3.95553200 | -1.71058100 | -3.27458700 |
| C | 1.69473400 | 7.41905500 | -5.50310100 |
| C | 1.24072500 | 7.57325800 | -4.04513100 |
| C | 1.06142100 | 6.24078500 | -3.35459500 |
| O | -0.02069700 | 5.79832300 | -3.01748400 |
| O | 2.20487800 | 5.58901600 | -3.17131000 |
| H | 0.95543300 | 6.85029200 | -6.08879600 |
| H | 0.28066900 | 8.10331100 | -3.98390100 |
| H | 1.99210100 | 8.14937600 | -3.48047400 |
| C | -5.63503600 | 2.00972300 | -1.61399000 |
| C | -4.54168900 | 2.91847200 | -2.13774000 |
| N | -3.29696600 | 2.17032800 | -2.35039100 |
| C | -2.09382600 | 2.75686400 | -2.45215700 |
| N | -1.86256500 | 3.94798600 | -1.92442500 |
| N | -1.10420600 | 2.11486900 | -3.10738900 |
| H | -5.35818500 | 1.58361800 | -0.63788100 |
| H | -6.56581400 | 2.58105300 | -1.48870500 |
| H | -4.36964700 | 3.72939700 | -1.42220800 |
| H | -4.85318700 | 3.38462900 | -3.09017200 |
| H | -3.37861300 | 1.19058500 | -2.63182400 |
| H | -2.43198300 | 4.28440000 | -1.11181400 |
| H | -1.04413200 | 4.49004100 | -2.21823000 |
| H | -1.29259700 | 1.25282100 | -3.60204300 |
| H | -0.14616100 | 2.44758200 | -2.98445700 |
| C | -3.60227100 | -5.79263800 | 1.77511700 |
| C | -4.11385500 | -4.61133800 | 2.60159000 |
| C | -5.40182600 | -3.99165600 | 2.03701000 |
| N | -5.22419100 | -3.37132100 | 0.73843600 |
| C | -4.79764800 | -2.11785400 | 0.55035500 |

| | | | |
|---|---|---|---|
| N | -4.65378000 | -1.24378600 | 1.56908500 |
| N | -4.54130400 | -1.71145000 | -0.69079100 |
| H | -3.46217900 | -5.51268400 | 0.71982300 |
| H | -2.64349100 | -6.17184100 | 2.16295100 |
| H | -3.34491600 | -3.81888200 | 2.65875600 |
| H | -4.30807100 | -4.92875700 | 3.64103200 |
| H | -5.82504000 | -3.26142600 | 2.74641200 |
| H | -6.16632000 | -4.77690500 | 1.92416500 |
| H | -5.14812200 | -4.00609100 | -0.10005300 |
| H | -4.77431900 | -1.51275100 | 2.53478800 |
| H | -4.15149700 | -0.37149300 | 1.39923700 |
| H | -4.71050200 | -2.35130300 | -1.50040000 |
| H | -4.06402800 | -0.82534600 | -0.82257600 |
| C | -4.90929300 | -6.87388700 | -3.72854400 |
| C | -3.47863900 | -6.53310200 | -4.23257800 |
| O | -3.14269400 | -6.71927700 | -5.39160800 |
| C | -5.69069400 | -5.53811500 | -3.61957100 |
| C | -5.14706000 | -4.55256700 | -2.56510900 |
| O | -4.99050600 | -3.34057400 | -2.88852300 |
| O | -4.90622500 | -5.01993000 | -1.41451600 |
| H | -5.36579500 | -7.53902800 | -4.47118800 |
| H | -6.73623600 | -5.75045400 | -3.33544400 |
| H | -5.71008900 | -5.03204800 | -4.59547800 |
| N | -2.65423200 | -5.91205300 | -3.32700100 |
| C | -1.39001700 | -5.24361500 | -3.68010800 |
| C | -0.74771900 | -4.61725500 | -2.43517600 |
| C | -1.61628300 | -3.49075100 | -1.84102200 |
| C | 0.66923900 | -4.13082800 | -2.75299300 |
| C | -1.26858600 | -3.13935600 | -0.39491500 |
| H | -3.02986200 | -5.75308500 | -2.39634100 |
| H | -1.58059200 | -4.47207900 | -4.44945800 |
| H | -0.65570800 | -5.41778400 | -1.67918200 |
| H | -2.67725800 | -3.77206500 | -1.87144100 |
| H | -1.53892100 | -2.59404700 | -2.48479600 |
| H | 1.15024800 | -3.68190300 | -1.86996800 |
| H | 0.65413700 | -3.36487400 | -3.54744100 |
| H | 1.30453100 | -4.96288700 | -3.09363000 |
| H | -1.42232600 | -4.01859400 | 0.25170400 |
| H | -1.91969600 | -2.33739500 | -0.01484100 |
| H | -0.22097000 | -2.81743800 | -0.27816600 |
| C | 3.57379300 | -6.10631100 | -0.76250500 |
| C | 2.28810500 | -5.84484000 | 0.00967900 |
| O | 1.25455800 | -6.44435500 | -0.23826600 |
| H | 3.41590100 | -5.81714200 | -1.81238700 |
| N | 2.35639600 | -4.90490700 | 1.00525200 |
| C | 1.22250500 | -4.62332300 | 1.84562000 |
| C | 1.17253600 | -3.15802000 | 2.27108300 |
| O | 2.15190700 | -2.42596100 | 2.19769700 |
| H | 3.20249900 | -4.37471100 | 1.17713600 |
| H | 0.31445700 | -4.92089700 | 1.30206600 |
| N | -0.01793900 | -2.75787400 | 2.80309000 |
| C | -0.27103800 | -1.36133100 | 3.16366400 |
| C | 0.53043000 | -0.92090100 | 4.39383900 |
| O | 1.12024300 | 0.14906800 | 4.42397800 |
| C | -1.76198900 | -1.11412900 | 3.35745500 |
| H | -0.82673500 | -3.34735600 | 2.63429600 |
| H | 0.09789900 | -0.71859200 | 2.34939400 |
| H | -1.95511600 | -0.04303200 | 3.51082300 |
| H | -2.15860200 | -1.67773600 | 4.21731600 |
| H | -2.30910700 | -1.41175400 | 2.45332300 |
| N | 0.45054400 | -1.78360000 | 5.43913300 |
| C | 0.97430900 | -1.49588000 | 6.75200700 |
| H | -0.02360700 | -2.66407000 | 5.27863600 |
| H | 1.38214300 | -0.47670700 | 6.73314800 |
| C | 6.57903800 | 4.76587300 | -2.27566500 |
| C | 5.41772900 | 3.91553000 | -1.77923700 |
| O | 5.18673200 | 3.70587200 | -0.59860400 |
| H | 6.25339500 | 5.45604300 | -3.06861200 |
| N | 4.65225400 | 3.37768100 | -2.78824300 |
| C | 3.89431900 | 2.17046800 | -2.56262800 |
| C | 4.67532100 | 0.98761500 | -3.15771200 |
| O | 5.62994000 | 1.17054000 | -3.89892700 |
| C | 2.47736800 | 2.22880300 | -3.13658000 |
| O | 1.61606400 | 3.07571500 | -2.38126600 |
| H | 5.08300500 | 3.39598100 | -3.70985500 |
| H | 3.82018500 | 2.02381900 | -1.47735400 |
| H | 2.05311400 | 1.21505200 | -3.17671100 |
| H | 2.50641400 | 2.63915700 | -4.15775400 |
| H | 2.01072100 | 4.70962800 | -2.75272500 |
| N | 4.22359600 | -0.24045000 | -2.81805500 |
| C | 4.89110300 | -1.40355200 | -3.36502100 |
| H | 3.38906200 | -0.33549000 | -2.23884400 |
| H | 5.91874200 | -1.48372700 | -2.97398100 |
| C | -4.47335900 | 1.98208900 | 4.44877900 |
| O | -5.01155800 | 0.79646100 | 3.88011800 |
| C | -3.67887200 | 2.87209300 | 3.50352600 |
| O | -2.62788700 | 2.11749200 | 2.90764600 |
| C | -4.42382700 | 3.59762300 | 2.34825400 |
| O | -4.64678700 | 4.92788800 | 2.73127500 |
| C | -3.44091300 | 3.40566300 | 1.13694500 |
| O | -3.19811000 | 4.54259800 | 0.36521000 |
| C | -2.15173300 | 2.88987000 | 1.81280900 |
| N | 1.07615200 | 1.77619900 | 0.13460600 |
| C | 0.72925300 | 0.61813000 | -0.51317300 |
| O | 1.53086500 | -0.06261000 | -1.16414100 |
| N | -0.59803400 | 0.22940300 | -0.40763200 |
| C | -1.61786600 | 0.87387500 | 0.29523700 |
| O | -2.75276500 | 0.38666100 | 0.27651800 |
| C | -1.19560700 | 2.07816500 | 0.98246900 |
| C | 0.12531200 | 2.43096500 | 0.86365600 |
| H | 1.58021800 | 2.75808800 | -1.44657700 |
| H | -0.85516200 | -0.62545500 | -0.89527200 |
| H | 0.47995100 | 3.32706200 | 1.38415800 |
| H | -1.61443200 | 3.78644000 | 2.18121400 |
| H | -4.07868000 | 5.11646700 | 0.32886400 |
| H | -5.38858000 | 3.10815900 | 2.11433600 |
| H | -3.25267200 | 3.69319100 | 4.11286200 |
| H | -5.26115000 | 2.60486200 | 4.91227400 |
| H | -3.84203400 | 2.57800200 | 0.52852600 |
| H | -3.79942300 | 1.64603200 | 5.25204000 |
| P | -5.73358100 | 5.92114800 | 1.89396500 |
| O | -5.38954400 | 5.70316700 | 0.40713900 |
| O | -5.75601000 | 7.23392900 | 2.57471800 |
| O | -7.12791200 | 5.05121100 | 2.12129700 |
| C | 6.13413500 | 0.18683800 | 5.25859300 |
| C | 5.19737500 | -0.73830000 | 4.47322200 |
| C | 4.22793400 | -0.04202200 | 3.51872800 |
| C | 2.96653300 | 1.44960900 | 1.07333300 |
| S | 4.99688400 | 0.75982600 | 2.04597800 |
| C | 5.27283400 | -0.69037200 | 0.94346000 |
| C | 6.72026500 | -1.01737900 | 0.61526900 |
| O | 7.26780300 | -0.02320700 | -0.22636400 |
| C | 7.72753400 | -1.11885700 | 1.76044600 |
| O | 7.57637800 | -2.33905300 | 2.46192000 |
| C | 9.05618500 | -1.03238200 | 0.98171400 |
| O | 9.35541900 | -2.29163700 | 0.42669200 |
| C | 8.70393800 | -0.04668800 | -0.13491000 |
| H | 6.77642900 | 0.77786900 | 4.58475900 |
| H | 5.77547200 | -1.49558100 | 3.91842700 |
| H | 4.56717000 | -1.30459600 | 5.18140600 |
| H | 3.68751700 | 0.77141900 | 4.02552100 |
| H | 3.46833200 | -0.73926800 | 3.13756000 |
| H | 2.46020400 | 1.46045100 | 2.03650400 |
| H | 3.41682600 | 2.35592300 | 0.67008900 |
| H | 2.96196200 | 0.54012500 | 0.48160200 |
| H | 9.14529300 | -0.38735400 | -1.08469500 |
| H | 9.05964000 | -2.95348900 | 1.07251900 |
| H | 7.61109900 | -0.24708100 | 2.42716900 |
| H | 6.70798500 | -2.00329100 | 0.10474700 |
| H | 4.76725600 | -1.55102900 | 1.40273400 |

| | | | |
|---|---|---|---|
| H | 7.98583900 | -2.25177600 | 3.33263900 |
| H | 9.88742400 | -0.66274200 | 1.61373700 |
| H | 4.76423500 | -0.46495500 | -0.00538900 |
| H | 1.81414100 | 8.40503100 | -5.97639800 |
| H | 2.65799700 | 6.89021600 | -5.55990600 |
| H | 7.35108300 | 4.10039400 | -2.69494400 |
| H | 7.00213500 | 5.33186900 | -1.43720500 |
| H | 9.05107100 | 0.97628000 | 0.08089400 |
| H | 6.78768400 | -0.38954800 | 5.93178700 |
| H | 5.56302100 | 0.89979500 | 5.87460200 |
| H | 1.78146000 | -2.19282200 | 7.03612900 |
| H | 0.18341000 | -1.54713200 | 7.51816200 |
| H | 1.24009500 | -5.22864900 | 2.77465200 |
| H | 3.77139300 | -7.18748300 | -0.74648300 |
| H | 4.44609100 | -5.56711200 | -0.36513500 |
| H | -0.70710300 | -5.97893300 | -4.13543300 |
| H | -4.88394500 | -7.36320700 | -2.74292700 |
| H | -4.32315500 | -6.62545500 | 1.79270000 |
| H | -5.69322200 | 1.05751600 | 3.24522300 |
| H | -7.44820700 | 4.81551000 | 1.24019100 |
| H | -5.83088100 | 1.16854700 | -2.29951300 |
| H | 4.32858100 | -2.30619300 | -3.09002700 |
| H | 4.95746900 | -1.33922200 | -4.46179300 |

#### Model2 – Final state

| | | | |
|---|---|---|---|
| O | -3.81152200 | -0.83405600 | -3.69560700 |
| H | -4.49596900 | -0.64105000 | -4.35200800 |
| H | -3.95665300 | -1.79216500 | -3.53077400 |
| C | 1.83821000 | 7.36833600 | -5.39304100 |
| C | 1.03279700 | 7.36021500 | -4.08878900 |
| C | 0.95753000 | 6.00647800 | -3.42053800 |
| O | -0.08086400 | 5.50210600 | -3.03101900 |
| O | 2.13997300 | 5.41829400 | -3.28789200 |
| H | 1.38907000 | 6.69786800 | -6.14285100 |
| H | -0.00043800 | 7.70037000 | -4.24227500 |
| H | 1.49847100 | 8.04737500 | -3.36028100 |
| C | -5.58061600 | 1.89717800 | -1.76852700 |
| C | -4.42508800 | 2.74189700 | -2.25313000 |
| N | -3.22795600 | 1.92528600 | -2.49081900 |
| C | -1.99277500 | 2.44565500 | -2.51758700 |
| N | -1.67683500 | 3.52333800 | -1.80801800 |
| N | -1.04544300 | 1.82598100 | -3.25450100 |
| H | -5.34030900 | 1.41886800 | -0.80606000 |
| H | -6.46359800 | 2.53427700 | -1.61660300 |
| H | -4.20539200 | 3.50508000 | -1.50014400 |
| H | -4.69703100 | 3.26429600 | -3.18812200 |
| H | -3.36288200 | 0.99700800 | -2.90548300 |
| H | -2.31741300 | 3.88425600 | -1.06800900 |
| H | -0.91187100 | 4.13010300 | -2.12773100 |
| H | -1.33489500 | 1.17326500 | -3.97287500 |
| H | -0.09566600 | 2.20782500 | -3.28888300 |
| C | -3.60144300 | -5.87389700 | 1.72253500 |
| C | -4.07349800 | -4.63530300 | 2.48716100 |
| C | -5.31606800 | -3.97312100 | 1.87197300 |
| N | -5.07644800 | -3.38015100 | 0.56945600 |
| C | -4.69313100 | -2.11865800 | 0.36061400 |
| N | -4.62072200 | -1.20564700 | 1.35563700 |
| N | -4.41040300 | -1.74100300 | -0.88509100 |
| H | -3.41090300 | -5.64319500 | 0.66286300 |
| H | -2.67819500 | -6.28822300 | 2.15718600 |
| H | -3.26881200 | -3.87857600 | 2.53389900 |
| H | -4.30817800 | -4.90160800 | 3.53249300 |
| H | -5.72965900 | -3.21583000 | 2.55737900 |
| H | -6.10843300 | -4.72898600 | 1.74864700 |
| H | -4.99520500 | -4.03156300 | -0.25863000 |
| H | -4.73417200 | -1.46721800 | 2.32550800 |
| H | -4.06700700 | -0.36299200 | 1.19574100 |
| H | -4.57021200 | -2.39834400 | -1.68516600 |
| H | -4.04214500 | -0.81412800 | -1.05689700 |
| C | -4.68599400 | -6.98495800 | -3.82332900 |
| C | -3.23437500 | -6.63953800 | -4.25840200 |
| O | -2.82651700 | -6.87293500 | -5.38538000 |
| C | -5.48170500 | -5.65246600 | -3.76195000 |
| C | -4.99655000 | -4.64172000 | -2.70148400 |
| O | -4.85923800 | -3.42829200 | -3.03429300 |
| O | -4.78366100 | -5.08513000 | -1.53710200 |
| H | -5.10170100 | -7.65820000 | -4.58214100 |
| H | -6.53676200 | -5.87001400 | -3.51952900 |
| H | -5.46579700 | -5.15973600 | -4.74481800 |
| N | -2.47492500 | -5.96138900 | -3.33705200 |
| C | -1.18773500 | -5.31919700 | -3.64594400 |
| C | -0.59674000 | -4.64809500 | -2.39906300 |
| C | -1.43757000 | -3.43829700 | -1.94740300 |
| C | 0.86526700 | -4.26627000 | -2.64863900 |
| C | -1.12990300 | -2.94580000 | -0.53297100 |
| H | -2.89375200 | -5.78123700 | -2.42876900 |
| H | -1.32616800 | -4.57843700 | -4.45570800 |
| H | -0.60061300 | -5.40051400 | -1.58958500 |
| H | -2.50689200 | -3.68685400 | -1.99765200 |
| H | -1.29694800 | -2.61518600 | -2.67368700 |
| H | 1.32039400 | -3.80601400 | -1.75789700 |
| H | 0.94925600 | -3.54752100 | -3.48248000 |
| H | 1.46292800 | -5.15463800 | -2.90166600 |
| H | -1.36354300 | -3.73180100 | 0.20378100 |
| H | -1.74356100 | -2.06959600 | -0.27497000 |
| H | -0.06884400 | -2.67648200 | -0.39761600 |
| C | 3.66925400 | -6.12287400 | -0.53779100 |
| C | 2.33619300 | -5.88949400 | 0.15707400 |
| O | 1.33155500 | -6.51542700 | -0.14287400 |
| H | 3.58549900 | -5.78398000 | -1.58171500 |
| N | 2.32589500 | -4.93610200 | 1.13945600 |
| C | 1.13382100 | -4.65222700 | 1.89244100 |
| C | 1.08019300 | -3.18953200 | 2.32658400 |
| O | 2.03980500 | -2.44062200 | 2.21604400 |
| H | 3.14191600 | -4.36831600 | 1.33946500 |
| H | 0.26220000 | -4.91838900 | 1.27530500 |
| N | -0.10234500 | -2.80373800 | 2.89101000 |
| C | -0.37009900 | -1.40501700 | 3.22217300 |
| C | 0.33065200 | -0.92883800 | 4.49942700 |
| O | 0.76474000 | 0.20744900 | 4.58925800 |
| C | -1.87133600 | -1.15978600 | 3.32993400 |
| H | -0.90177200 | -3.41759900 | 2.77185600 |
| H | 0.05167800 | -0.77396900 | 2.42525000 |
| H | -2.08011500 | -0.09388100 | 3.49835500 |
| H | -2.30896100 | -1.73745600 | 4.16045600 |
| H | -2.36241700 | -1.45003600 | 2.39292400 |
| N | 0.31723300 | -1.83332300 | 5.51309500 |
| C | 0.73829400 | -1.51607000 | 6.85602900 |
| H | -0.01409100 | -2.76619000 | 5.30005100 |
| H | 1.01911900 | -0.45478800 | 6.87381400 |
| C | 6.62164900 | 4.77440300 | -1.97216600 |
| C | 5.24753000 | 4.15169900 | -2.02301800 |
| O | 4.41788800 | 4.29388900 | -1.13707500 |
| H | 7.00978800 | 5.01590900 | -2.97276600 |
| N | 5.02769500 | 3.34815200 | -3.10819100 |
| C | 4.11369800 | 2.23933500 | -2.99962400 |
| C | 4.89096200 | 0.97792800 | -3.36966500 |
| O | 5.89595300 | 1.02430700 | -4.06376900 |
| C | 2.80931000 | 2.35062100 | -3.81574900 |
| O | 1.73016200 | 2.78003400 | -2.98169300 |
| H | 5.81960800 | 3.13197300 | -3.71348800 |
| H | 3.81759800 | 2.15575300 | -1.94366800 |
| H | 2.55550600 | 1.37826000 | -4.27051600 |
| H | 2.92481800 | 3.09448900 | -4.61552900 |
| H | 2.00174100 | 4.51555900 | -2.90628800 |
| N | 4.35880100 | -0.16271100 | -2.88200300 |
| C | 5.03817100 | -1.41498900 | -3.10413400 |
| H | 3.59578500 | -0.10637000 | -2.21221200 |
| H | 5.90471300 | -1.52944800 | -2.42932800 |
| C | -4.80628100 | 1.89513300 | 4.43690700 |

| | | | |
|---|---|---|---|
| O | -5.15582200 | 0.70701000 | 3.74951500 |
| C | -3.96701700 | 2.85616000 | 3.61672700 |
| O | -2.80863700 | 2.16120900 | 3.15614600 |
| C | -4.61755600 | 3.53394300 | 2.37357900 |
| O | -4.81001500 | 4.89089000 | 2.66704500 |
| C | -3.56754700 | 3.25700200 | 1.24631000 |
| O | -3.31397300 | 4.28109000 | 0.33500500 |
| C | -2.31587200 | 2.91793200 | 2.07223600 |
| N | 1.10996800 | 2.24664300 | 0.68777400 |
| C | 0.94058700 | 1.22072200 | -0.20815500 |
| O | 1.81842600 | 0.88024400 | -1.01071600 |
| N | -0.29436700 | 0.62198300 | -0.18119700 |
| C | -1.43847200 | 1.03771800 | 0.51850100 |
| O | -2.48563600 | 0.41655000 | 0.35404500 |
| C | -1.21512800 | 2.20588800 | 1.35062500 |
| C | 0.04509200 | 2.71350200 | 1.41157300 |
| H | 1.76376500 | 2.19291000 | -2.19340100 |
| H | -0.42482400 | -0.14663700 | -0.83524500 |
| H | 0.26243700 | 3.57350900 | 2.04553600 |
| H | -1.91568800 | 3.88814200 | 2.42930300 |
| H | -4.19163300 | 4.86442600 | 0.23823100 |
| H | -5.58479800 | 3.06725300 | 2.10931000 |
| H | -3.66583300 | 3.69758000 | 4.26914000 |
| H | -5.69664200 | 2.43493700 | 4.80977300 |
| H | -3.88887500 | 2.33057800 | 0.73683200 |
| H | -4.21680700 | 1.57704500 | 5.31053300 |
| P | -5.85234900 | 5.81806500 | 1.71966100 |
| O | -5.48597100 | 5.45293300 | 0.26533500 |
| O | -5.85881600 | 7.19273300 | 2.26477300 |
| O | -7.26717400 | 4.99864200 | 1.99829700 |
| C | 5.93444800 | 0.21404200 | 5.55516800 |
| C | 5.01421700 | -0.68554500 | 4.72313200 |
| C | 4.15944400 | 0.05513900 | 3.69193300 |
| C | 2.43126500 | 2.88230400 | 0.78929000 |
| S | 5.09840900 | 0.84518200 | 2.31708000 |
| C | 5.21499500 | -0.54531900 | 1.11918000 |
| C | 6.63180400 | -0.96489600 | 0.75963300 |
| O | 7.29331200 | 0.06864500 | 0.04735700 |
| C | 7.57279000 | -1.27849100 | 1.92100000 |
| O | 7.31281600 | -2.57105300 | 2.44165400 |
| C | 8.94554700 | -1.16826900 | 1.23392700 |
| O | 9.20916500 | -2.35349500 | 0.51836400 |
| C | 8.71084900 | -0.00987100 | 0.26470900 |
| H | 6.62420700 | 0.77797900 | 4.90644600 |
| H | 5.60060600 | -1.46577200 | 4.20673900 |
| H | 4.32095000 | -1.22397700 | 5.39489100 |
| H | 3.59769500 | 0.87309100 | 4.16909400 |
| H | 3.40993700 | -0.61284700 | 3.24360400 |
| H | 2.37529900 | 3.65511500 | 1.56360300 |
| H | 2.73446700 | 3.34825600 | -0.15587600 |
| H | 3.18620100 | 2.14101500 | 1.08786800 |
| H | 9.23488900 | -0.20081800 | -0.68541700 |
| H | 8.81671800 | -3.07237100 | 1.04046100 |
| H | 7.45717300 | -0.49832100 | 2.69142100 |
| H | 6.56099500 | -1.87959100 | 0.13417800 |
| H | 4.66766400 | -1.40278400 | 1.53599900 |
| H | 7.63369500 | -2.60730400 | 3.35168600 |
| H | 9.75917200 | -0.94461400 | 1.95228900 |
| H | 4.69763200 | -0.23967300 | 0.19707300 |
| H | 1.86505300 | 8.38145500 | -5.82092100 |
| H | 2.87229600 | 7.03860400 | -5.21708000 |
| H | 7.30742700 | 4.04853100 | -1.50452300 |
| H | 6.59701700 | 5.67873700 | -1.35126500 |
| H | 9.05734800 | 0.95064600 | 0.68230600 |
| H | 6.53548700 | -0.37255600 | 6.26790600 |
| H | 5.34953900 | 0.94852800 | 6.13269800 |
| H | 1.61149100 | -2.11478100 | 7.16694500 |
| H | -0.07538200 | -1.68183600 | 7.58150700 |
| H | 1.06491700 | -5.27566600 | 2.80652300 |
| H | 3.86314000 | -7.20432000 | -0.55897400 |
| H | 4.51268200 | -5.60554800 | -0.05789900 |
| H | -0.49300600 | -6.08041500 | -4.03629200 |
| H | -4.70428600 | -7.46757700 | -2.83414000 |
| H | -4.36788300 | -6.66476500 | 1.74686900 |
| H | -5.78436200 | 0.94163000 | 3.05281700 |
| H | -7.60707300 | 4.73249800 | 1.13340000 |
| H | -5.84091500 | 1.09870400 | -2.48267100 |
| H | 4.34265400 | -2.25020900 | -2.93958400 |
| H | 5.41198200 | -1.45417000 | -4.13591100 |

#### Model3 – Initial state

| | | | |
|---|---|---|---|
| O | 2.64943800 | 1.55916800 | -3.07855500 |
| H | 1.90298600 | 1.56860100 | -3.69333000 |
| H | 3.04280900 | 2.45962700 | -3.16824600 |
| O | -1.25721000 | -3.89215900 | -1.08010000 |
| H | -2.16265100 | -4.12793500 | -0.83143800 |
| H | -1.01829700 | -4.31162000 | -1.96635200 |
| C | -0.70176000 | -8.19325800 | -4.97946300 |
| C | -1.07895200 | -6.76354500 | -4.54159300 |
| C | -0.08132200 | -5.93868900 | -3.68913600 |
| O | 0.99474700 | -6.41036300 | -3.31385400 |
| O | -0.47998600 | -4.73087300 | -3.42162500 |
| H | -0.50403600 | -8.83464500 | -4.10689600 |
| H | -2.02242100 | -6.77316800 | -3.96496400 |
| H | -1.30323600 | -6.13800800 | -5.42345600 |
| C | 5.83904800 | -1.18336800 | -1.60928800 |
| C | 5.13475200 | -2.45885400 | -2.08996900 |
| N | 3.69144000 | -2.23786300 | -2.33273800 |
| C | 2.75667000 | -3.21399900 | -2.49550200 |
| N | 2.85467500 | -4.37579600 | -1.85940900 |
| N | 1.69591000 | -2.99034900 | -3.28312800 |
| H | 5.39215100 | -0.78664000 | -0.69085900 |
| H | 6.89357600 | -1.42355200 | -1.40991500 |
| H | 5.27104700 | -3.26202000 | -1.35472900 |
| H | 5.60477400 | -2.81184800 | -3.02501700 |
| H | 3.46728700 | -1.34970500 | -2.77282300 |
| H | 3.36871600 | -4.42434000 | -0.96043300 |
| H | 2.22749200 | -5.13759700 | -2.15970100 |
| H | 1.70338100 | -2.21340300 | -3.93073000 |
| H | 0.88620400 | -3.66369200 | -3.31567500 |
| C | 2.38491400 | 6.30253300 | 1.62406600 |
| C | 3.38222300 | 5.50689900 | 2.46095500 |
| C | 4.72330900 | 5.32063900 | 1.73749800 |
| N | 4.60922700 | 4.60060400 | 0.48514700 |
| C | 4.48533500 | 3.26948200 | 0.39863600 |
| N | 4.57927700 | 2.47492600 | 1.48868700 |
| N | 4.28824000 | 2.71488800 | -0.78902700 |
| H | 2.23939600 | 5.84215100 | 0.63508000 |
| H | 1.40356500 | 6.37653200 | 2.11996000 |
| H | 2.96465700 | 4.51294400 | 2.70486000 |
| H | 3.56844400 | 6.01093300 | 3.42512800 |
| H | 5.45642000 | 4.81776300 | 2.39163200 |
| H | 5.15433600 | 6.30693300 | 1.50544900 |
| H | 4.30987200 | 5.13170300 | -0.36995100 |
| H | 4.67830300 | 2.85919700 | 2.41518400 |
| H | 4.37968300 | 1.47796400 | 1.38626000 |
| H | 4.15117700 | 3.30349800 | -1.64292700 |
| H | 4.08652700 | 1.71508500 | -0.80603900 |
| C | 2.90356600 | 7.49021300 | -4.10187900 |
| C | 1.44727600 | 6.91364200 | -4.26390600 |
| O | 0.84634900 | 7.01448700 | -5.32153300 |
| C | 3.89778100 | 6.30981100 | -4.08840600 |
| C | 3.76571800 | 5.37095600 | -2.87662600 |
| O | 3.63214200 | 5.90117200 | -1.73911800 |
| O | 3.82670500 | 4.12333700 | -3.08381600 |
| H | 3.06859600 | 8.14525700 | -4.96528700 |
| H | 4.93004600 | 6.70165100 | -4.06358100 |
| H | 3.79921200 | 5.71253800 | -5.00620200 |
| N | 0.91242300 | 6.21962700 | -3.19753000 |
| C | -0.36290000 | 5.45104900 | -3.22411600 |

| | | | |
|---|---|---|---|
| C | -0.55774200 | 4.69598000 | -1.90938900 |
| C | 0.52489200 | 3.62131700 | -1.72280300 |
| C | -1.97150600 | 4.11165700 | -1.83475700 |
| C | 0.66961700 | 3.12124200 | -0.29111200 |
| H | 1.47601200 | 6.17295900 | -2.35236300 |
| H | -0.34534600 | 4.75283700 | -4.07967300 |
| H | -0.45813100 | 5.43892800 | -1.09440100 |
| H | 1.49758700 | 4.00662500 | -2.05655700 |
| H | 0.29670800 | 2.76850500 | -2.38463200 |
| H | -2.13515200 | 3.38232700 | -2.64602400 |
| H | -2.73086800 | 4.90319200 | -1.92491000 |
| H | -2.14833700 | 3.59209800 | -0.88083000 |
| H | -0.21268300 | 2.55361300 | 0.04084500 |
| H | 0.81322900 | 3.97117100 | 0.39835300 |
| H | 1.54717200 | 2.46976900 | -0.18855600 |
| C | -5.30654400 | 5.50685800 | 0.08107500 |
| C | -3.94706600 | 5.55133300 | 0.74338700 |
| O | -3.21550700 | 6.52467200 | 0.68329600 |
| H | -5.20030000 | 5.82622200 | -0.96541400 |
| N | -3.59380000 | 4.43272100 | 1.44348500 |
| C | -2.30477400 | 4.36320500 | 2.06857200 |
| C | -1.94934700 | 2.91970000 | 2.37330800 |
| O | -2.78140500 | 2.01732000 | 2.35647400 |
| H | -4.13194300 | 3.57211200 | 1.41405300 |
| H | -1.54510400 | 4.82633900 | 1.41641000 |
| N | -0.64652300 | 2.71872800 | 2.69602100 |
| C | -0.11802000 | 1.37434700 | 2.85852800 |
| C | -0.70484100 | 0.70330400 | 4.12077000 |
| O | -1.17075300 | -0.42987600 | 4.09207500 |
| C | 1.40962000 | 1.40525700 | 2.91893300 |
| H | 0.01317900 | 3.45771800 | 2.47713300 |
| H | -0.44797100 | 0.74487900 | 2.01739200 |
| H | 1.83098600 | 0.40112000 | 3.06837400 |
| H | 1.75992700 | 2.03997100 | 3.74909800 |
| H | 1.82566800 | 1.80016300 | 1.98205700 |
| N | -0.60116200 | 1.45260200 | 5.24689000 |
| C | -0.93817800 | 0.99139000 | 6.57985700 |
| H | -0.24476000 | 2.39488600 | 5.13935200 |
| H | -1.14903100 | -0.08442500 | 6.52349500 |
| C | -5.28903100 | -5.66707100 | -1.33229000 |
| C | -4.48385900 | -4.39235500 | -1.30653900 |
| O | -4.02234900 | -3.94293000 | -0.24872800 |
| H | -5.30251300 | -6.14552800 | -2.32164500 |
| N | -4.28345500 | -3.77509800 | -2.49232500 |
| C | -3.85541400 | -2.39527400 | -2.56411400 |
| C | -5.11190300 | -1.49374500 | -2.46582400 |
| O | -6.18360900 | -1.97037000 | -2.10172700 |
| C | -2.93318000 | -2.14332800 | -3.77182700 |
| O | -2.43449200 | -0.82494400 | -3.76441900 |
| H | -4.79056900 | -4.13600800 | -3.29260400 |
| H | -3.24975600 | -2.20734500 | -1.66643300 |
| H | -3.50182600 | -2.28440700 | -4.70824100 |
| H | -2.13250800 | -2.90310100 | -3.74809900 |
| H | -1.62108400 | -0.77633500 | -3.21515500 |
| N | -4.91487200 | -0.18363400 | -2.72795600 |
| C | -5.98529200 | 0.79745500 | -2.66457900 |
| H | -3.99195600 | 0.05812300 | -3.10294100 |
| H | -6.92670200 | 0.28053200 | -2.43901300 |
| C | 5.86315300 | -0.70217500 | 3.68809500 |
| O | 5.41636200 | -0.03776100 | 4.85585400 |
| C | 4.89746100 | -1.76663100 | 3.21943300 |
| O | 3.68875100 | -1.20076200 | 2.70160100 |
| C | 5.50884800 | -2.59641500 | 2.07439500 |
| O | 6.13227300 | -3.73706700 | 2.59330900 |
| C | 4.30343100 | -2.83248700 | 1.10875600 |
| O | 4.03233700 | -4.15243400 | 0.74708100 |
| C | 3.11516100 | -2.17591500 | 1.84052200 |
| N | 0.18780100 | -1.81956300 | -0.51173400 |
| C | 0.50198700 | -0.77011900 | -1.33145300 |
| O | -0.16428200 | -0.43345500 | -2.30965800 |
| N | 1.65017100 | -0.09658900 | -0.97670400 |
| C | 2.52234400 | -0.43298800 | 0.05179400 |
| O | 3.55587600 | 0.22853900 | 0.18872700 |
| C | 2.14072800 | -1.57313600 | 0.86820200 |
| C | 0.99902900 | -2.22334700 | 0.51077200 |
| H | -0.54493400 | -2.51999100 | -0.81984000 |
| H | 1.96919000 | 0.63946300 | -1.63529400 |
| H | 0.68081700 | -3.14427000 | 1.00379800 |
| H | 2.60908200 | -2.96778600 | 2.42665500 |
| H | 4.95896500 | -4.57317700 | 0.56665100 |
| H | 6.25298500 | -1.98070000 | 1.53836100 |
| H | 4.66491500 | -2.45434700 | 4.05346100 |
| H | 6.02134300 | 0.01144600 | 2.85237800 |
| H | 4.49779100 | -2.21719200 | 0.22516600 |
| H | 6.82128700 | -1.22682000 | 3.85848400 |
| P | 7.20650300 | -4.57327200 | 1.58906500 |
| O | 6.41523500 | -4.78806900 | 0.29802400 |
| O | 7.85646000 | -5.63732900 | 2.39720600 |
| O | 8.29089700 | -3.36689300 | 1.24263600 |
| C | -5.97053700 | -2.02998900 | 5.56407500 |
| C | -5.67650900 | -0.73144800 | 4.80841600 |
| C | -4.44935500 | -0.76157300 | 3.89745900 |
| C | -2.83507200 | -2.48727400 | 2.32811700 |
| S | -4.57914800 | -2.04120300 | 2.57217200 |
| C | -4.88263700 | -1.11932300 | 1.01803000 |
| C | -6.33615800 | -0.81250400 | 0.72549500 |
| O | -7.16828400 | -1.89986200 | 1.06485200 |
| C | -7.03599900 | 0.36803400 | 1.39593700 |
| O | -6.54622700 | 1.61002900 | 0.92020800 |
| C | -8.49975300 | 0.08009400 | 0.96641100 |
| O | -8.73569800 | 0.63020300 | -0.30687900 |
| C | -8.52577700 | -1.46315600 | 0.88493500 |
| H | -6.21060100 | -2.85303900 | 4.87252900 |
| H | -6.56699900 | -0.43258000 | 4.23219700 |
| H | -5.49764600 | 0.08815200 | 5.52557300 |
| H | -3.53089500 | -1.02753400 | 4.44273300 |
| H | -4.24639500 | 0.19705100 | 3.39635400 |
| H | -2.80491100 | -3.05734400 | 1.39121000 |
| H | -2.20965700 | -1.58495100 | 2.32718000 |
| H | -2.55718600 | -3.12414400 | 3.17745300 |
| H | -8.89827400 | -1.75755900 | -0.10889100 |
| H | -8.19772000 | 1.43585800 | -0.36812300 |
| H | -6.93985700 | 0.29936000 | 2.49124500 |
| H | -6.37149700 | -0.62867700 | -0.36102700 |
| H | -4.25959900 | -0.21264000 | 1.01483400 |
| H | -6.77463700 | 2.29963500 | 1.55711800 |
| H | -9.22737700 | 0.46753000 | 1.70540900 |
| H | -4.52002000 | -1.84910100 | 0.27908000 |
| H | -9.13579800 | -1.93357700 | 1.66923200 |
| H | -6.83035200 | -1.90011700 | 6.23748700 |
| H | -5.10748400 | -2.34114600 | 6.17471200 |
| H | -4.88888400 | -6.36315600 | -0.58385700 |
| H | -6.32298700 | -5.41280200 | -1.04770100 |
| H | -5.79559100 | 1.54113700 | -1.87258900 |
| H | -6.08721600 | 1.32751900 | -3.62493900 |
| H | 0.21366400 | -8.18799800 | -5.59073500 |
| H | -1.51503800 | -8.64241000 | -5.57036500 |
| H | 5.80214000 | -0.38984700 | -2.37484700 |
| H | 9.05985900 | -3.51803700 | 1.81000600 |
| H | 6.19281800 | 0.29524000 | 5.31989200 |
| H | 2.74518300 | 7.32842600 | 1.44696700 |
| H | 3.00006800 | 8.06007700 | -3.16507400 |
| H | -1.20143200 | 6.14497500 | -3.40153700 |
| H | -5.96826100 | 6.23406400 | 0.57644500 |
| H | -5.77444800 | 4.51104600 | 0.10919600 |
| H | -0.09615400 | 1.15127400 | 7.27105500 |
| H | -1.82729000 | 1.50734800 | 6.98081300 |
| H | -2.27433100 | 4.93522800 | 3.01682300 |

#### Model3 − TS1

| | | | |
|---|---|---|---|
| O | 2.54015300 | 1.49965900 | -3.12499200 |
| H | 1.78208400 | 1.53592000 | -3.72419800 |
| H | 2.94371400 | 2.39297600 | -3.21197200 |
| O | -1.07821900 | -4.07871200 | -1.09092700 |
| H | -2.02315600 | -4.11788600 | -0.84020100 |
| H | -0.86061100 | -4.41209800 | -2.15090000 |
| C | -0.63591100 | -8.16724100 | -4.98776900 |
| C | -1.00758600 | -6.70667000 | -4.67603700 |
| C | -0.16233900 | -5.92897800 | -3.64992800 |
| O | 0.85817600 | -6.38235800 | -3.14737500 |
| O | -0.63027600 | -4.72509000 | -3.38954600 |
| H | -0.70749500 | -8.79596400 | -4.08690400 |
| H | -2.05167400 | -6.63217000 | -4.32391500 |
| H | -0.98608700 | -6.09674700 | -5.59721200 |
| C | 5.86418200 | -1.11687400 | -1.62386800 |
| C | 5.15887800 | -2.39084600 | -2.09876300 |
| N | 3.72103700 | -2.16262300 | -2.36251100 |
| C | 2.77839800 | -3.12768300 | -2.48422000 |
| N | 2.86792800 | -4.26919300 | -1.81231900 |
| N | 1.72499700 | -2.91753500 | -3.29234500 |
| H | 5.38400300 | -0.69327300 | -0.73381500 |
| H | 6.90678400 | -1.36680400 | -1.37892500 |
| H | 5.27979700 | -3.18768400 | -1.35515600 |
| H | 5.63471800 | -2.75749000 | -3.02558500 |
| H | 3.49533600 | -1.26727000 | -2.78787800 |
| H | 3.39186200 | -4.29555400 | -0.90837100 |
| H | 2.21325700 | -5.02211300 | -2.05135000 |
| H | 1.69674900 | -2.10817500 | -3.89720800 |
| H | 0.90889100 | -3.54873500 | -3.29379400 |
| C | 2.36596200 | 6.33864400 | 1.63219100 |
| C | 3.34346600 | 5.48438600 | 2.43529100 |
| C | 4.67057200 | 5.26579200 | 1.69416000 |
| N | 4.52416500 | 4.56041300 | 0.43763800 |
| C | 4.39446400 | 3.23044700 | 0.33676100 |
| N | 4.49886500 | 2.42103400 | 1.41484400 |
| N | 4.18492100 | 2.68985700 | -0.85571400 |
| H | 2.19183700 | 5.90961700 | 0.63341200 |
| H | 1.39393100 | 6.43528300 | 2.14227900 |
| H | 2.89265300 | 4.50000500 | 2.65876700 |
| H | 3.55786000 | 5.95620700 | 3.41000500 |
| H | 5.39448700 | 4.73549400 | 2.33657200 |
| H | 5.12750000 | 6.24202500 | 1.46806500 |
| H | 4.22746700 | 5.10407200 | -0.41108500 |
| H | 4.57210900 | 2.79567400 | 2.34778400 |
| H | 4.27537100 | 1.42811300 | 1.29730000 |
| H | 4.05444300 | 3.29034800 | -1.70066300 |
| H | 3.94854800 | 1.69564600 | -0.87621300 |
| C | 2.85998700 | 7.53980100 | -4.09308600 |
| C | 1.39643400 | 6.97433500 | -4.24048900 |
| O | 0.77227100 | 7.11124600 | -5.28077400 |
| C | 3.84308300 | 6.34792900 | -4.10768700 |
| C | 3.70398300 | 5.38361800 | -2.91458100 |
| O | 3.57516800 | 5.89780200 | -1.76735500 |
| O | 3.74785600 | 4.14083100 | -3.14386000 |
| H | 3.01775000 | 8.20418900 | -4.95063600 |
| H | 4.87960300 | 6.72813500 | -4.07911200 |
| H | 3.73423500 | 5.77077000 | -5.03723700 |
| N | 0.88303900 | 6.24858600 | -3.18460200 |
| C | -0.38933000 | 5.47596400 | -3.20974800 |
| C | -0.57904700 | 4.71222300 | -1.89930500 |
| C | 0.49137100 | 3.62208600 | -1.73280500 |
| C | -1.99931000 | 4.14465700 | -1.81710200 |
| C | 0.63922300 | 3.10039100 | -0.30928800 |
| H | 1.46855700 | 6.17023700 | -2.35633900 |
| H | -0.37558500 | 4.78125200 | -4.06867500 |
| H | -0.46201400 | 5.44664800 | -1.07894500 |
| H | 1.46669600 | 3.99887300 | -2.06909900 |
| H | 0.24734600 | 2.78111900 | -2.40433500 |
| H | -2.18094000 | 3.42996200 | -2.63761600 |
| H | -2.74932000 | 4.94709600 | -1.88738900 |
| H | -2.17219100 | 3.61242800 | -0.86965400 |
| H | -0.25687100 | 2.55870800 | 0.02891600 |
| H | 0.82213600 | 3.93744800 | 0.38691900 |
| H | 1.49453700 | 2.41756700 | -0.22710400 |
| C | -5.32393700 | 5.49082300 | 0.10939600 |
| C | -3.96833700 | 5.55288000 | 0.77876600 |
| O | -3.25750100 | 6.54229700 | 0.73702900 |
| H | -5.21140200 | 5.78902200 | -0.94291400 |
| N | -3.59221100 | 4.42944800 | 1.45788200 |
| C | -2.29894700 | 4.37047500 | 2.07565900 |
| C | -1.93804600 | 2.92918600 | 2.38684700 |
| O | -2.76942400 | 2.02549300 | 2.37765900 |
| H | -4.11472600 | 3.55977600 | 1.41730800 |
| H | -1.54315400 | 4.83106600 | 1.41697200 |
| N | -0.63389800 | 2.73306600 | 2.70400500 |
| C | -0.09831900 | 1.39062000 | 2.86506500 |
| C | -0.68292400 | 0.71725900 | 4.12721100 |
| O | -1.15971900 | -0.41160900 | 4.09717600 |
| C | 1.42845200 | 1.42739000 | 2.92031300 |
| H | 0.02229700 | 3.47322700 | 2.47838000 |
| H | -0.42590000 | 0.76010600 | 2.02388500 |
| H | 1.85361300 | 0.42436900 | 3.06662500 |
| H | 1.78045800 | 2.06499000 | 3.74780200 |
| H | 1.83741400 | 1.81998200 | 1.97968200 |
| N | -0.56846700 | 1.46050700 | 5.25664700 |
| C | -0.90523000 | 0.99543100 | 6.58802200 |
| H | -0.20165400 | 2.39919300 | 5.15303000 |
| H | -1.11605800 | -0.08024100 | 6.52855900 |
| C | -5.23079100 | -5.68020200 | -1.32340100 |
| C | -4.36801700 | -4.44151400 | -1.35682500 |
| O | -3.76955400 | -4.04266700 | -0.34506400 |
| H | -5.38982900 | -6.11710800 | -2.31905700 |
| N | -4.28142200 | -3.78806100 | -2.53491900 |
| C | -3.84366700 | -2.40888300 | -2.60801300 |
| C | -5.09512100 | -1.50742000 | -2.46337900 |
| O | -6.15371800 | -1.98903100 | -2.06575100 |
| C | -2.96074600 | -2.13660900 | -3.84101700 |
| O | -2.45074400 | -0.82600800 | -3.81742200 |
| H | -4.90118200 | -4.09932900 | -3.27482200 |
| H | -3.20968400 | -2.22869900 | -1.72789000 |
| H | -3.56062100 | -2.25283400 | -4.76109700 |
| H | -2.16647700 | -2.90308700 | -3.85957000 |
| H | -1.67448000 | -0.79538200 | -3.20769900 |
| N | -4.90899700 | -0.19996400 | -2.73620200 |
| C | -5.97679700 | 0.78140300 | -2.64257200 |
| H | -3.99352300 | 0.04184500 | -3.13176400 |
| H | -6.91600300 | 0.26421300 | -2.40854300 |
| C | 5.89045400 | -0.63156500 | 3.66226600 |
| O | 5.45156300 | 0.01453400 | 4.84435200 |
| C | 4.93561700 | -1.70937000 | 3.20042000 |
| O | 3.71398100 | -1.16327000 | 2.69372500 |
| C | 5.54952900 | -2.53432800 | 2.05037400 |
| O | 6.17977100 | -3.67589600 | 2.56447300 |
| C | 4.33880200 | -2.77007600 | 1.09354600 |
| O | 4.09903100 | -4.08851100 | 0.69476600 |
| C | 3.14222200 | -2.15954200 | 1.84936200 |
| N | 0.09413500 | -1.99118100 | -0.39084700 |
| C | 0.35553000 | -0.94573700 | -1.22108000 |
| O | -0.34411400 | -0.63008200 | -2.19411600 |
| N | 1.49864500 | -0.21092800 | -0.94240000 |
| C | 2.42795800 | -0.48438900 | 0.04955900 |
| O | 3.45033800 | 0.21745100 | 0.11542800 |
| C | 2.11354200 | -1.60239000 | 0.90631400 |
| C | 0.96614600 | -2.29322700 | 0.60417800 |
| H | -0.64431700 | -3.08429600 | -0.81744100 |
| H | 1.76000700 | 0.52181800 | -1.61894100 |
| H | 0.72193200 | -3.19930100 | 1.16937300 |
| H | 2.68565300 | -2.96578400 | 2.45679100 |
| H | 5.03575000 | -4.49713100 | 0.52175300 |
| H | 6.29058700 | -1.91567900 | 1.51388400 |

| | | | |
|---|---|---|---|
| H | 4.72189000 | -2.40119000 | 4.03638200 |
| H | 6.02205800 | 0.09154000 | 2.83020100 |
| H | 4.50869400 | -2.12383700 | 0.22699000 |
| H | 6.86048800 | -1.14039600 | 3.81409700 |
| P | 7.26483000 | -4.50240200 | 1.56476500 |
| O | 6.48312000 | -4.71709600 | 0.26709300 |
| O | 7.91491900 | -5.56766300 | 2.37145800 |
| O | 8.34611400 | -3.29088700 | 1.22761600 |
| C | -5.91877200 | -2.05999900 | 5.58110900 |
| C | -5.62512400 | -0.75582500 | 4.83355600 |
| C | -4.40256200 | -0.77459200 | 3.91505000 |
| C | -2.77665600 | -2.49400400 | 2.33721000 |
| S | -4.52219800 | -2.06235100 | 2.59708400 |
| C | -4.84510400 | -1.15777600 | 1.03555200 |
| C | -6.30047700 | -0.85372300 | 0.74936700 |
| O | -7.13240700 | -1.94310100 | 1.08401900 |
| C | -6.99771800 | 0.32408700 | 1.42682800 |
| O | -6.50886300 | 1.56710800 | 0.95407300 |
| C | -8.46302300 | 0.03906700 | 1.00019300 |
| O | -8.70193700 | 0.59592400 | -0.26949400 |
| C | -8.49113900 | -1.50335200 | 0.91013700 |
| H | -6.16911100 | -2.87610900 | 4.88505700 |
| H | -6.51889800 | -0.45049200 | 4.26592900 |
| H | -5.44107600 | 0.05746100 | 5.55654400 |
| H | -3.47590800 | -1.02579500 | 4.45359300 |
| H | -4.21418100 | 0.18443600 | 3.40876500 |
| H | -2.75979800 | -3.10997500 | 1.42960200 |
| H | -2.16526500 | -1.58511500 | 2.26910800 |
| H | -2.46752000 | -3.07969200 | 3.21192000 |
| H | -8.86630900 | -1.79211700 | -0.08427600 |
| H | -8.17207200 | 1.40757200 | -0.32316700 |
| H | -6.89898700 | 0.25078500 | 2.52139200 |
| H | -6.33821900 | -0.66460300 | -0.33609200 |
| H | -4.22505700 | -0.24897000 | 1.01966400 |
| H | -6.70975300 | 2.25105400 | 1.60601900 |
| H | -9.18801300 | 0.42308200 | 1.74346500 |
| H | -4.47961500 | -1.88471100 | 0.29536000 |
| H | -9.09909600 | -1.97761300 | 1.69365100 |
| H | -6.77207200 | -1.93055200 | 6.26278500 |
| H | -5.05224600 | -2.38044200 | 6.18193500 |
| H | -4.77615000 | -6.42156100 | -0.65366800 |
| H | -6.20799500 | -5.38905700 | -0.90474900 |
| H | -5.77320700 | 1.51479700 | -1.84449900 |
| H | -6.09102500 | 1.32346200 | -3.59458300 |
| H | 0.39986000 | -8.23945500 | -5.35237700 |
| H | -1.30848600 | -8.57811000 | -5.75590400 |
| H | 5.86704000 | -0.34103500 | -2.40795900 |
| H | 9.10961700 | -3.43576900 | 1.80389500 |
| H | 6.22606000 | 0.39455400 | 5.27449000 |
| H | 2.76253300 | 7.35473900 | 1.47754900 |
| H | 2.97363500 | 8.09586300 | -3.15001200 |
| H | -1.22917400 | 6.16974600 | -3.38093900 |
| H | -5.99061100 | 6.22639900 | 0.58486100 |
| H | -5.78926700 | 4.49434800 | 0.15299100 |
| H | -0.06296600 | 1.15314500 | 7.27940900 |
| H | -1.79422800 | 1.50996500 | 6.99150000 |
| H | -2.26660000 | 4.94937800 | 3.01936900 |

#### Model3 − Intermediate

| | | | |
|---|---|---|---|
| O | 2.57122100 | 1.37876600 | -3.15156800 |
| H | 1.75280200 | 1.47621800 | -3.65756500 |
| H | 3.00411100 | 2.25601100 | -3.25330000 |
| O | -1.34467200 | -4.18201600 | -1.08960400 |
| H | -2.16780700 | -4.15638400 | -0.57321000 |
| H | -1.48774400 | -5.11132800 | -2.35879400 |
| C | -0.78608200 | -8.18843300 | -4.95020400 |
| C | -0.86012600 | -6.66691400 | -5.18046600 |
| C | -0.64359500 | -5.80834500 | -3.94372900 |
| O | 0.40995200 | -5.22197900 | -3.72478900 |
| O | -1.69061600 | -5.72508500 | -3.14892200 |
| H | -1.55110000 | -8.51743700 | -4.23031100 |
| H | -1.85049500 | -6.40834900 | -5.58931000 |
| H | -0.09639500 | -6.36069600 | -5.90937600 |
| C | 5.86725500 | -1.25858700 | -1.63601700 |
| C | 5.10429300 | -2.47025500 | -2.13894400 |
| N | 3.70889200 | -2.11904100 | -2.44090000 |
| C | 2.68722800 | -2.98936700 | -2.47476900 |
| N | 2.72167700 | -4.11910400 | -1.77444700 |
| N | 1.60312300 | -2.70300200 | -3.21728400 |
| H | 5.36682700 | -0.79556100 | -0.77721300 |
| H | 6.87746500 | -1.57059500 | -1.33209700 |
| H | 5.13593000 | -3.27339300 | -1.39508500 |
| H | 5.57821100 | -2.87241600 | -3.05205000 |
| H | 3.55128900 | -1.20987500 | -2.86951800 |
| H | 3.29069500 | -4.19057500 | -0.89051500 |
| H | 1.94077600 | -4.75639400 | -1.90000600 |
| H | 1.64526000 | -1.96788300 | -3.90996100 |
| H | 0.89589300 | -3.43369400 | -3.32783300 |
| C | 2.53211300 | 6.27647600 | 1.60809600 |
| C | 3.48828000 | 5.38983300 | 2.40207800 |
| C | 4.80095400 | 5.12641400 | 1.64970100 |
| N | 4.62102400 | 4.42042400 | 0.39805100 |
| C | 4.45841300 | 3.09369800 | 0.30382100 |
| N | 4.55656500 | 2.28419200 | 1.38221500 |
| N | 4.22238900 | 2.55306400 | -0.88412100 |
| H | 2.33329200 | 5.85289500 | 0.61157100 |
| H | 1.56912700 | 6.40598500 | 2.12787600 |
| H | 3.00699300 | 4.42101700 | 2.63003800 |
| H | 3.72698000 | 5.85432200 | 3.37470900 |
| H | 5.51324200 | 4.57601000 | 2.28812000 |
| H | 5.28621800 | 6.08697300 | 1.41530800 |
| H | 4.33410000 | 4.96940400 | -0.45103000 |
| H | 4.63027500 | 2.66003000 | 2.31472100 |
| H | 4.30068300 | 1.29760300 | 1.26496900 |
| H | 4.10649100 | 3.15391800 | -1.73003700 |
| H | 3.95796400 | 1.56475800 | -0.89441100 |
| C | 3.02729300 | 7.44824600 | -4.12316100 |
| C | 1.54858500 | 6.91912500 | -4.25802200 |
| O | 0.91628300 | 7.07968200 | -5.29017200 |
| C | 3.97877300 | 6.22974000 | -4.14407900 |
| C | 3.82209500 | 5.26617300 | -2.95109800 |
| O | 3.70920800 | 5.78393400 | -1.80335300 |
| O | 3.83588400 | 4.02314500 | -3.18041700 |
| H | 3.19517100 | 8.10744300 | -4.98275000 |
| H | 5.02523800 | 6.58221200 | -4.12144800 |
| H | 3.84918900 | 5.65702300 | -5.07380200 |
| N | 1.03039800 | 6.19657800 | -3.20238600 |
| C | -0.25916600 | 5.45295400 | -3.22002300 |
| C | -0.46426800 | 4.69915300 | -1.90676800 |
| C | 0.57260900 | 3.57659800 | -1.74068900 |
| C | -1.90052800 | 4.17517500 | -1.81390800 |
| C | 0.71331300 | 3.06610200 | -0.31248800 |
| H | 1.62319500 | 6.09811400 | -2.38128000 |
| H | -0.26483400 | 4.75427700 | -4.07604400 |
| H | -0.31966600 | 5.43199700 | -1.08950400 |
| H | 1.55624100 | 3.91785300 | -2.09113900 |
| H | 0.28986400 | 2.73581800 | -2.39782600 |
| H | -2.10988700 | 3.46597900 | -2.63251300 |
| H | -2.62628300 | 5.00004700 | -1.87861900 |
| H | -2.08171700 | 3.64786600 | -0.86542500 |
| H | -0.19244600 | 2.54370100 | 0.02919300 |
| H | 0.91275600 | 3.90637500 | 0.37516100 |
| H | 1.55477300 | 2.36768000 | -0.22370100 |
| C | -5.17922800 | 5.57786700 | 0.11834800 |
| C | -3.82355900 | 5.61390600 | 0.78883800 |
| O | -3.09507000 | 6.59068600 | 0.75074000 |
| H | -5.06716300 | 5.91556300 | -0.92177100 |
| N | -3.47039300 | 4.48269900 | 1.46663700 |
| C | -2.18032100 | 4.39831300 | 2.08763100 |
| C | -1.84160100 | 2.94941400 | 2.38792100 |

| | | | |
|---|---|---|---|
| O | -2.68712400 | 2.05884200 | 2.37522600 |
| H | -4.01060300 | 3.62394600 | 1.42464700 |
| H | -1.41647000 | 4.85361900 | 1.43473300 |
| N | -0.54053200 | 2.73275500 | 2.70217900 |
| C | -0.02586300 | 1.38300900 | 2.86760100 |
| C | -0.60950700 | 0.72978600 | 4.14064600 |
| O | -1.09966400 | -0.39269800 | 4.12599600 |
| C | 1.50147700 | 1.39369600 | 2.91081400 |
| H | 0.12650200 | 3.46358700 | 2.47842000 |
| H | -0.36932700 | 0.75246300 | 2.03307200 |
| H | 1.90868800 | 0.38329400 | 3.05805200 |
| H | 1.87068100 | 2.02905300 | 3.73265900 |
| H | 1.90890600 | 1.77415900 | 1.96481900 |
| N | -0.48169200 | 1.47924200 | 5.26484800 |
| C | -0.82568400 | 1.01657400 | 6.59503000 |
| H | -0.10476500 | 2.41343900 | 5.15849200 |
| H | -1.04756900 | -0.05657900 | 6.53277300 |
| C | -5.31557000 | -5.59752700 | -1.27640200 |
| C | -4.49923100 | -4.32827100 | -1.20462300 |
| O | -4.04970000 | -3.89117700 | -0.13776000 |
| H | -5.04473300 | -6.19949700 | -2.15644500 |
| N | -4.27878000 | -3.70791800 | -2.38571800 |
| C | -3.84462200 | -2.33311500 | -2.45916400 |
| C | -5.09511300 | -1.42398500 | -2.38898300 |
| O | -6.17822900 | -1.89109600 | -2.04564100 |
| C | -2.91627200 | -2.09916800 | -3.66552100 |
| O | -2.42295300 | -0.78910400 | -3.70679400 |
| H | -4.78049900 | -4.06587800 | -3.19064600 |
| H | -3.23994800 | -2.13492900 | -1.56245500 |
| H | -3.46908800 | -2.29198900 | -4.60280300 |
| H | -2.10900500 | -2.84889600 | -3.59138300 |
| H | -1.68732900 | -0.71057600 | -3.04347900 |
| N | -4.87880400 | -0.11872900 | -2.65284300 |
| C | -5.93724200 | 0.87327100 | -2.61483100 |
| H | -3.95169700 | 0.10543500 | -3.03033100 |
| H | -6.88867800 | 0.36966600 | -2.40078900 |
| C | 5.90296600 | -0.75480400 | 3.63909700 |
| O | 5.50318600 | -0.09738100 | 4.82986800 |
| C | 4.93162700 | -1.83088900 | 3.20069600 |
| O | 3.70294800 | -1.28424100 | 2.71323800 |
| C | 5.52965200 | -2.66335500 | 2.04456900 |
| O | 6.12474500 | -3.82919100 | 2.55062100 |
| C | 4.31867800 | -2.85109200 | 1.07998900 |
| O | 4.07689900 | -4.15077200 | 0.61600600 |
| C | 3.12121600 | -2.26568100 | 1.85253300 |
| N | -0.01971000 | -1.97423000 | -0.27609200 |
| C | 0.25478100 | -0.91449500 | -1.07688100 |
| O | -0.48410500 | -0.49827100 | -1.98634000 |
| N | 1.45799400 | -0.25167200 | -0.86375800 |
| C | 2.41310800 | -0.57133800 | 0.08370000 |
| O | 3.46799900 | 0.09262700 | 0.11313800 |
| C | 2.08288000 | -1.68252400 | 0.93464000 |
| C | 0.88524700 | -2.31777100 | 0.66788100 |
| H | -0.87961200 | -3.31382400 | -0.87276300 |
| H | 1.72038000 | 0.47199200 | -1.54375100 |
| H | 0.63058900 | -3.21358900 | 1.24765700 |
| H | 2.67032900 | -3.08386600 | 2.44831700 |
| H | 5.01580500 | -4.57169700 | 0.45424400 |
| H | 6.29250000 | -2.06281500 | 1.51881200 |
| H | 4.73196400 | -2.51916500 | 4.04298600 |
| H | 6.01956600 | -0.03680500 | 2.80047600 |
| H | 4.49696500 | -2.16690600 | 0.24457700 |
| H | 6.87323900 | -1.26933700 | 3.76947300 |
| P | 7.21250600 | -4.66074500 | 1.55875300 |
| O | 6.45086000 | -4.83197600 | 0.24151200 |
| O | 7.82105900 | -5.75813200 | 2.35483800 |
| O | 8.32864100 | -3.47042300 | 1.26558100 |
| C | -5.90340400 | -1.94110000 | 5.61825300 |
| C | -5.69112400 | -0.68972100 | 4.76191800 |
| C | -4.42027500 | -0.67747800 | 3.90991100 |
| C | -2.70565900 | -2.23903000 | 2.26971500 |
| S | -4.47201600 | -1.93239700 | 2.55929900 |
| C | -4.84060700 | -0.98867000 | 1.03378800 |
| C | -6.30291800 | -0.69883300 | 0.77025200 |
| O | -7.12680600 | -1.77398700 | 1.16583200 |
| C | -6.99329000 | 0.50516800 | 1.40892000 |
| O | -6.51655300 | 1.72880500 | 0.87426000 |
| C | -8.46514600 | 0.19808200 | 1.01304800 |
| O | -8.73265900 | 0.72860000 | -0.26213700 |
| C | -8.48269100 | -1.34869300 | 0.95550600 |
| H | -5.99915900 | -2.84356700 | 4.99339200 |
| H | -6.57180200 | -0.53791800 | 4.11763500 |
| H | -5.63350900 | 0.20148700 | 5.41018800 |
| H | -3.51931000 | -0.93284600 | 4.48905900 |
| H | -4.21725400 | 0.29190900 | 3.43027500 |
| H | -2.65247700 | -2.77948300 | 1.31884700 |
| H | -2.15338500 | -1.29249500 | 2.26315300 |
| H | -2.35124500 | -2.86338700 | 3.09866100 |
| H | -8.83463800 | -1.65746700 | -0.04157200 |
| H | -8.18292800 | 1.52364900 | -0.35400000 |
| H | -6.87684800 | 0.48573600 | 2.50374900 |
| H | -6.36790700 | -0.55394600 | -0.32055700 |
| H | -4.22851800 | -0.07453200 | 1.02896300 |
| H | -6.72115900 | 2.44093400 | 1.49413700 |
| H | -9.17962400 | 0.59145200 | 1.76156400 |
| H | -4.47991800 | -1.69997400 | 0.27842100 |
| H | -9.10378200 | -1.81210000 | 1.73486800 |
| H | -6.82265500 | -1.85080200 | 6.21536500 |
| H | -5.06186600 | -2.09767700 | 6.31206800 |
| H | -5.16056100 | -6.18181900 | -0.36164200 |
| H | -6.38019900 | -5.32228300 | -1.34955300 |
| H | -5.75238900 | 1.62088000 | -1.82588500 |
| H | -6.01801900 | 1.39670900 | -3.58098500 |
| H | 0.20076800 | -8.48147600 | -4.56016100 |
| H | -0.95155400 | -8.72108600 | -5.89816100 |
| H | 5.96637300 | -0.48965900 | -2.42111800 |
| H | 9.07543200 | -3.64162900 | 1.85637500 |
| H | 6.29364700 | 0.27778700 | 5.23433700 |
| H | 2.96145700 | 7.27842400 | 1.44864900 |
| H | 3.16255300 | 8.00147400 | -3.18130300 |
| H | -1.08255200 | 6.16579300 | -3.39252100 |
| H | -5.84620100 | 6.29465900 | 0.62183400 |
| H | -5.64437700 | 4.58036700 | 0.12478300 |
| H | 0.01647600 | 1.16502800 | 7.28872000 |
| H | -1.71046300 | 1.53935400 | 6.99739200 |
| H | -2.14185100 | 4.96797900 | 3.03678400 |

#### Model3 – TS2

| | | | |
|---|---|---|---|
| O | 0.73592800 | 0.15903500 | -3.41114700 |
| H | -0.19158100 | -0.11331700 | -3.60962200 |
| H | 0.90202000 | 0.94521100 | -3.94977400 |
| O | -0.32735300 | -4.02206800 | -1.75279700 |
| H | -1.18925000 | -4.10039700 | -1.27171900 |
| H | -0.40350300 | -4.99236400 | -3.13107700 |
| C | 0.43402900 | -8.02001700 | -5.26874400 |
| C | 1.47494600 | -7.30219900 | -4.40794800 |
| C | 0.93765300 | -6.27501500 | -3.43763300 |
| O | 1.49997000 | -5.99348600 | -2.38602000 |
| O | -0.17775100 | -5.67581300 | -3.82693300 |
| H | -0.14615400 | -7.30370700 | -5.86778400 |
| H | 2.18914000 | -6.75487300 | -5.05071700 |
| H | 2.08231100 | -8.00338400 | -3.81710100 |
| C | 6.08742600 | -0.46425500 | -1.49782700 |
| C | 5.54483200 | -1.80160800 | -1.99929600 |
| N | 4.10561800 | -1.69841600 | -2.28394000 |
| C | 3.18551300 | -2.67258300 | -2.25383500 |
| N | 3.44773700 | -3.84785700 | -1.68578600 |
| N | 1.98952300 | -2.45432700 | -2.84179100 |
| H | 5.56757200 | -0.13954900 | -0.58748100 |
| H | 7.15681800 | -0.57353500 | -1.26721300 |

| | | | |
|---|---|---|---|
| H | 5.74237400 | -2.59082300 | -1.26164200 |
| H | 6.06954000 | -2.10049200 | -2.92347400 |
| H | 3.75403500 | -0.76422700 | -2.45973000 |
| H | 3.98607300 | -3.85970100 | -0.78205000 |
| H | 2.78490500 | -4.60825900 | -1.83338700 |
| H | 1.77557900 | -1.54388200 | -3.25279700 |
| H | 1.18323400 | -2.96647700 | -2.48118800 |
| C | 1.76321100 | 6.56435600 | 1.71541900 |
| C | 2.66042800 | 5.53567000 | 2.40591800 |
| C | 3.97741500 | 5.27782600 | 1.65646800 |
| N | 3.80656000 | 4.64124600 | 0.36689800 |
| C | 3.85618000 | 3.32412500 | 0.15160600 |
| N | 4.04472900 | 2.44129500 | 1.16350100 |
| N | 3.74380500 | 2.86611000 | -1.09201600 |
| H | 1.54429400 | 6.27001700 | 0.67692700 |
| H | 0.80794000 | 6.69208500 | 2.24924100 |
| H | 2.12653200 | 4.57379800 | 2.51820500 |
| H | 2.90484100 | 5.87372100 | 3.42831800 |
| H | 4.66406400 | 4.68068000 | 2.27941000 |
| H | 4.48612000 | 6.24042500 | 1.48332800 |
| H | 3.52867300 | 5.25992300 | -0.45426800 |
| H | 3.94985600 | 2.73209300 | 2.12518400 |
| H | 3.92358400 | 1.44427500 | 0.96810300 |
| H | 3.73364800 | 3.55186100 | -1.90388400 |
| H | 3.60910500 | 1.86490800 | -1.20100900 |
| C | 2.39308800 | 7.92288000 | -3.96109000 |
| C | 1.01713300 | 7.18791700 | -4.18841300 |
| O | 0.42202400 | 7.28887600 | -5.25100900 |
| C | 3.54624100 | 6.90501200 | -3.91596700 |
| C | 3.45314900 | 5.86493700 | -2.78651900 |
| O | 3.06279400 | 6.26708900 | -1.64406300 |
| O | 3.77299100 | 4.68480800 | -3.06094400 |
| H | 2.50185700 | 8.61645300 | -4.80356700 |
| H | 4.50049400 | 7.44313300 | -3.77430700 |
| H | 3.62193700 | 6.35967700 | -4.86768900 |
| N | 0.53522700 | 6.39351500 | -3.16878400 |
| C | -0.68288300 | 5.54744000 | -3.26069000 |
| C | -0.85144700 | 4.66924800 | -2.02058300 |
| C | 0.27549100 | 3.63076700 | -1.88708300 |
| C | -2.23765000 | 4.01551800 | -2.03339000 |
| C | 0.35975400 | 2.99255200 | -0.50392000 |
| H | 1.10728300 | 6.33703000 | -2.32665700 |
| H | -0.62072500 | 4.92622700 | -4.17241000 |
| H | -0.80764600 | 5.34076800 | -1.14148000 |
| H | 1.24386200 | 4.09477500 | -2.12558200 |
| H | 0.12329500 | 2.83856000 | -2.64360600 |
| H | -2.34465900 | 3.34947000 | -2.90709000 |
| H | -3.02625200 | 4.78022000 | -2.07945400 |
| H | -2.41468300 | 3.41195100 | -1.12988400 |
| H | -0.54248700 | 2.41259700 | -0.25345700 |
| H | 0.49549200 | 3.77355300 | 0.26338700 |
| H | 1.22197600 | 2.31766000 | -0.43186600 |
| C | -5.73831400 | 5.03396700 | -0.17062600 |
| C | -4.37413100 | 5.18863600 | 0.47171400 |
| O | -3.63909900 | 6.13207000 | 0.23262500 |
| H | -5.60184100 | 4.66433000 | -1.19990300 |
| N | -3.99886900 | 4.18233500 | 1.31891600 |
| C | -2.69516600 | 4.18363400 | 1.91783200 |
| C | -2.24163300 | 2.76946000 | 2.24909300 |
| O | -3.00216100 | 1.80805700 | 2.23784900 |
| H | -4.57999300 | 3.36704800 | 1.48118200 |
| H | -1.98558400 | 4.67156300 | 1.22992100 |
| N | -0.93201300 | 2.67486100 | 2.60377100 |
| C | -0.27620100 | 1.38358600 | 2.75407000 |
| C | -0.75518900 | 0.61934100 | 4.00504500 |
| O | -0.88956100 | -0.59804900 | 3.99189300 |
| C | 1.23733900 | 1.57108900 | 2.81852500 |
| H | -0.33646600 | 3.46975500 | 2.39559800 |
| H | -0.54354200 | 0.73514400 | 1.90704600 |
| H | 1.75582300 | 0.61700500 | 2.98171600 |
| H | 1.50973600 | 2.24734000 | 3.64539600 |
| H | 1.60787700 | 1.99467500 | 1.87686500 |
| N | -0.92054300 | 1.38164600 | 5.11477400 |
| C | -1.20520200 | 0.83912800 | 6.42922200 |
| H | -0.82878000 | 2.38456400 | 5.00302500 |
| H | -1.13763300 | -0.25435800 | 6.36239000 |
| C | -4.53081700 | -6.04791400 | -1.77827000 |
| C | -3.59789300 | -4.85081300 | -1.75896800 |
| O | -2.77907800 | -4.66629500 | -0.85214100 |
| H | -4.72841500 | -6.39478000 | -2.80265300 |
| N | -3.75688700 | -3.98225600 | -2.78886000 |
| C | -3.41424600 | -2.57775600 | -2.67181800 |
| C | -4.72802600 | -1.80871100 | -2.90583300 |
| O | -5.75402100 | -2.39849100 | -3.22041100 |
| C | -2.28288300 | -2.12761500 | -3.62656500 |
| O | -1.86781300 | -0.80136200 | -3.34751000 |
| H | -4.60365400 | -4.10763300 | -3.34576900 |
| H | -3.05546600 | -2.39004800 | -1.64770500 |
| H | -2.63127100 | -2.16301700 | -4.67124700 |
| H | -1.45055300 | -2.84133600 | -3.52249000 |
| H | -1.60948700 | -0.75714800 | -2.37741100 |
| N | -4.66123600 | -0.47010000 | -2.73213000 |
| C | -5.82347900 | 0.34146700 | -3.02536300 |
| H | -3.72900300 | -0.05916000 | -2.75991900 |
| H | -6.70355900 | -0.09009100 | -2.52807800 |
| C | 5.87266800 | -0.02028600 | 3.78055600 |
| O | 5.28621400 | 0.49151400 | 4.96228800 |
| C | 5.06391500 | -1.17676400 | 3.25457900 |
| O | 3.80714200 | -0.74673300 | 2.72506800 |
| C | 5.75717100 | -1.94352600 | 2.11350400 |
| O | 6.49227700 | -3.00953000 | 2.65307900 |
| C | 4.55803600 | -2.33018500 | 1.18105100 |
| O | 4.49233100 | -3.67738400 | 0.81084400 |
| C | 3.32531800 | -1.86951000 | 1.98238900 |
| N | -0.27254000 | -1.89050400 | 0.82671300 |
| C | -0.28053800 | -1.04608600 | -0.21984600 |
| O | -1.32692900 | -0.77076700 | -0.85608000 |
| N | 0.90611600 | -0.43190700 | -0.55732500 |
| C | 2.11405900 | -0.55827800 | 0.11935500 |
| O | 3.07417900 | 0.14543300 | -0.24551100 |
| C | 2.09636800 | -1.51734800 | 1.19689600 |
| C | 0.87950100 | -2.10127200 | 1.49022000 |
| H | 0.26882300 | -4.69015900 | -1.36614400 |
| H | 0.89953200 | 0.13301900 | -1.41386100 |
| H | 0.81774900 | -2.79416400 | 2.33607300 |
| H | 3.06655500 | -2.69275300 | 2.67677700 |
| H | 5.47439600 | -3.98316300 | 0.65772600 |
| H | 6.43275300 | -1.26721600 | 1.56039300 |
| H | 4.89085400 | -1.89661700 | 4.07600300 |
| H | 5.94496700 | 0.75216200 | 2.98599200 |
| H | 4.61515100 | -1.67624300 | 0.30325700 |
| H | 6.89160900 | -0.41199400 | 3.96071100 |
| P | 7.65900000 | -3.76861300 | 1.69629500 |
| O | 6.91759300 | -4.08257000 | 0.39335100 |
| O | 8.37903000 | -4.75610400 | 2.54056700 |
| O | 8.63870200 | -2.48180300 | 1.33532700 |
| C | -5.85981200 | -2.64958700 | 5.14524700 |
| C | -5.31252200 | -1.26658100 | 4.77843300 |
| C | -4.12551300 | -1.28490000 | 3.81881900 |
| C | -2.30050800 | -2.15881900 | 1.54596600 |
| S | -4.42708400 | -2.14609300 | 2.21930100 |
| C | -4.99724200 | -0.79757800 | 1.10867500 |
| C | -6.48071200 | -0.79689700 | 0.79651200 |
| O | -6.86249900 | -1.99728500 | 0.15470900 |
| C | -7.46161300 | -0.68157600 | 1.96097000 |
| O | -7.48672200 | 0.63696200 | 2.46725900 |
| C | -8.76781100 | -1.11599400 | 1.27453100 |
| O | -9.26658800 | -0.05791900 | 0.49015700 |
| C | -8.26497200 | -2.24009100 | 0.37112400 |
| H | -6.23816600 | -3.18178800 | 4.25811100 |

| | | | |
|---|---|---|---|
| H | -6.10606700 | -0.62431500 | 4.36420400 |
| H | -4.96512300 | -0.75137700 | 5.69123000 |
| H | -3.27950100 | -1.83375700 | 4.25824700 |
| H | -3.75971100 | -0.27704600 | 3.57518400 |
| H | -2.32868500 | -3.15048500 | 1.10044500 |
| H | -2.45361000 | -1.30934300 | 0.88915700 |
| H | -1.87159100 | -1.98952200 | 2.53112400 |
| H | -8.80382800 | -2.23034700 | -0.58878100 |
| H | -9.09308200 | 0.75925800 | 0.98400800 |
| H | -7.18954500 | -1.41478600 | 2.73941200 |
| H | -6.65443800 | 0.07686600 | 0.13507700 |
| H | -4.67984800 | 0.16094800 | 1.54374600 |
| H | -7.92449500 | 0.63521500 | 3.32870400 |
| H | -9.52562200 | -1.47478500 | 1.99878400 |
| H | -4.44488200 | -0.93459000 | 0.16741600 |
| H | -8.38120000 | -3.23042200 | 0.84183300 |
| H | -6.68326800 | -2.57041400 | 5.87120100 |
| H | -5.07382400 | -3.27739100 | 5.59518700 |
| H | -4.10010000 | -6.86003800 | -1.17989300 |
| H | -5.48976300 | -5.74357300 | -1.32750700 |
| H | -5.65838600 | 1.36071500 | -2.64910900 |
| H | -6.03985100 | 0.39113700 | -4.10654900 |
| H | -0.27401300 | -8.58985000 | -4.64712700 |
| H | 0.92758400 | -8.72558600 | -5.95324700 |
| H | 5.97575500 | 0.32960600 | -2.25549800 |
| H | 9.39849700 | -2.53346000 | 1.93210400 |
| H | 5.94123400 | 1.04547500 | 5.40293500 |
| H | 2.25663700 | 7.54803900 | 1.66990400 |
| H | 2.36475600 | 8.48324200 | -3.01346900 |
| H | -1.56577600 | 6.19644400 | -3.38786500 |
| H | -6.21312300 | 6.02191100 | -0.23437000 |
| H | -6.39679000 | 4.33696300 | 0.36886000 |
| H | -0.47164600 | 1.19744900 | 7.16843600 |
| H | -2.21687500 | 1.11219500 | 6.77517000 |
| H | -2.66556400 | 4.77757000 | 2.85341500 |

#### Model3 – Final state

| | | | |
|---|---|---|---|
| O | 0.58817000 | 0.18482400 | -3.19050000 |
| H | -0.32871100 | -0.11582200 | -3.40779100 |
| H | 0.69410900 | 1.05311000 | -3.60538200 |
| O | -0.22122300 | -3.97603800 | -1.67698000 |
| H | -1.12945500 | -4.17312000 | -1.31131600 |
| H | -0.29037200 | -4.90803300 | -3.13472400 |
| C | 0.38153800 | -8.01201900 | -5.22256600 |
| C | 1.45858000 | -7.33314100 | -4.37314500 |
| C | 0.97800600 | -6.26407900 | -3.41926600 |
| O | 1.54575700 | -6.00353500 | -2.36485500 |
| O | -0.09479600 | -5.59959500 | -3.82609900 |
| H | -0.16217900 | -7.27779000 | -5.83396900 |
| H | 2.19845600 | -6.83339100 | -5.02568000 |
| H | 2.03029300 | -8.05432000 | -3.77105500 |
| C | 6.03451700 | -0.35699000 | -1.65685700 |
| C | 5.50179600 | -1.70612000 | -2.13186000 |
| N | 4.05614800 | -1.62360900 | -2.38362400 |
| C | 3.14722600 | -2.60645000 | -2.31776000 |
| N | 3.42595100 | -3.76209700 | -1.72020900 |
| N | 1.94147000 | -2.41378000 | -2.89400000 |
| H | 5.53052200 | -0.02911800 | -0.73799500 |
| H | 7.11004900 | -0.44587300 | -1.44684200 |
| H | 5.72392000 | -2.48548300 | -1.39116100 |
| H | 6.01108600 | -2.00775200 | -3.06371000 |
| H | 3.70264500 | -0.71180600 | -2.64875800 |
| H | 4.00260100 | -3.75711300 | -0.83962800 |
| H | 2.77568500 | -4.53777200 | -1.84306700 |
| H | 1.70686900 | -1.50272900 | -3.28885100 |
| H | 1.14896100 | -2.93393800 | -2.51173100 |
| C | 1.70655600 | 6.61544400 | 1.67164000 |
| C | 2.64303600 | 5.60415500 | 2.33517300 |
| C | 3.93020100 | 5.34998100 | 1.53520600 |
| N | 3.71140800 | 4.70095400 | 0.25796900 |
| C | 3.78345300 | 3.38674400 | 0.04395300 |
| N | 4.03505000 | 2.51279100 | 1.05163500 |
| N | 3.63456200 | 2.92374800 | -1.19518200 |
| H | 1.43960600 | 6.30286300 | 0.64986400 |
| H | 0.77735100 | 6.74538900 | 2.24878900 |
| H | 2.12412100 | 4.63887900 | 2.48231100 |
| H | 2.92745500 | 5.95755800 | 3.34176100 |
| H | 4.64714200 | 4.76402400 | 2.13405200 |
| H | 4.42312800 | 6.31475500 | 1.33171900 |
| H | 3.40063600 | 5.31639800 | -0.55714300 |
| H | 3.95431800 | 2.80592600 | 2.01433600 |
| H | 3.92439200 | 1.51383900 | 0.87465400 |
| H | 3.58315600 | 3.61104100 | -2.00879000 |
| H | 3.52775600 | 1.92166000 | -1.30672000 |
| C | 2.13401700 | 7.96344800 | -4.02608900 |
| C | 0.76258800 | 7.20799700 | -4.20765800 |
| O | 0.12871600 | 7.30112400 | -5.24779000 |
| C | 3.29956000 | 6.95661400 | -4.01025600 |
| C | 3.25034900 | 5.91802300 | -2.87670400 |
| O | 2.90625900 | 6.32137000 | -1.72054500 |
| O | 3.55828900 | 4.73535000 | -3.15989200 |
| H | 2.20993300 | 8.65241500 | -4.87571600 |
| H | 4.25163200 | 7.50498100 | -3.89630700 |
| H | 3.35356200 | 6.41015600 | -4.96274500 |
| N | 0.33115300 | 6.40123400 | -3.17456100 |
| C | -0.88096900 | 5.54360200 | -3.22075600 |
| C | -0.99786900 | 4.67351500 | -1.96884300 |
| C | 0.11557600 | 3.61464800 | -1.89825300 |
| C | -2.39435900 | 4.04663700 | -1.89979500 |
| C | 0.27034000 | 2.97073300 | -0.52375000 |
| H | 0.92420100 | 6.36243900 | -2.34649300 |
| H | -0.84501300 | 4.91651700 | -4.12974600 |
| H | -0.88996400 | 5.34500200 | -1.09538300 |
| H | 1.07644700 | 4.05944100 | -2.19809100 |
| H | -0.10463500 | 2.82842600 | -2.64515700 |
| H | -2.56914000 | 3.38998600 | -2.76982700 |
| H | -3.16832100 | 4.82716000 | -1.89360500 |
| H | -2.53041600 | 3.44219300 | -0.99006700 |
| H | -0.63438900 | 2.42571300 | -0.21122700 |
| H | 0.48214300 | 3.74303000 | 0.23416000 |
| H | 1.11230700 | 2.26631400 | -0.50837200 |
| C | -5.82665200 | 4.96315100 | 0.03071100 |
| C | -4.46776300 | 5.13091500 | 0.67549400 |
| O | -3.77234100 | 6.11732300 | 0.49398000 |
| H | -5.69040500 | 4.88569300 | -1.05917500 |
| N | -4.05586700 | 4.09801300 | 1.47053300 |
| C | -2.75811300 | 4.13301800 | 2.07991700 |
| C | -2.23270600 | 2.72769200 | 2.32359600 |
| O | -2.93783600 | 1.73233700 | 2.24358400 |
| H | -4.59056100 | 3.23989200 | 1.56524300 |
| H | -2.07401700 | 4.70482400 | 1.43139100 |
| N | -0.91837300 | 2.67718500 | 2.68308400 |
| C | -0.21822600 | 1.40756300 | 2.79026000 |
| C | -0.57992800 | 0.61655700 | 4.06346800 |
| O | -0.54296500 | -0.60626400 | 4.06447500 |
| C | 1.29069600 | 1.64150600 | 2.76656000 |
| H | -0.35799300 | 3.50884000 | 2.52879900 |
| H | -0.52257400 | 0.75900800 | 1.95605300 |
| H | 1.84486900 | 0.70652500 | 2.92026000 |
| H | 1.58279700 | 2.33363700 | 3.57343600 |
| H | 1.59862000 | 2.06807900 | 1.80435700 |
| N | -0.82462900 | 1.38262900 | 5.15607500 |
| C | -1.02095200 | 0.86172800 | 6.49507400 |
| H | -0.87703500 | 2.38316900 | 5.00480500 |
| H | -0.87894500 | -0.22624900 | 6.45511600 |
| C | -4.49948800 | -6.10433700 | -1.58133500 |
| C | -3.54429100 | -4.94092300 | -1.69682300 |
| O | -2.59073800 | -4.79917700 | -0.92137600 |
| H | -4.93718700 | -6.38013500 | -2.55110100 |
| N | -3.84131100 | -4.04361400 | -2.66430600 |

| | | | |
|---|---|---|---|
| C | -3.49408100 | -2.64345400 | -2.54110200 |
| C | -4.80775500 | -1.88084200 | -2.79424200 |
| O | -5.81397600 | -2.47335600 | -3.16167700 |
| C | -2.35822100 | -2.17539000 | -3.47590900 |
| O | -1.92630400 | -0.86740600 | -3.12264400 |
| H | -4.75184500 | -4.14376100 | -3.12206300 |
| H | -3.15382200 | -2.45892000 | -1.50951800 |
| H | -2.70227100 | -2.14725300 | -4.52120500 |
| H | -1.53158600 | -2.89942800 | -3.40842300 |
| H | -1.70596100 | -0.87992900 | -2.15353400 |
| N | -4.76021000 | -0.55308900 | -2.57138600 |
| C | -5.92992100 | 0.25988200 | -2.80562300 |
| H | -3.84056500 | -0.12534400 | -2.50915900 |
| H | -6.81453600 | -0.24235000 | -2.39032900 |
| C | 6.02035100 | 0.12984600 | 3.63516500 |
| O | 5.42493500 | 0.60891200 | 4.82255900 |
| C | 5.23059900 | -1.04718700 | 3.13720200 |
| O | 3.93938100 | -0.64647000 | 2.66753300 |
| C | 5.87924200 | -1.81301400 | 1.97292700 |
| O | 6.62975200 | -2.87473600 | 2.49466800 |
| C | 4.64338800 | -2.21922000 | 1.09782700 |
| O | 4.57968600 | -3.56030700 | 0.71683300 |
| C | 3.45876400 | -1.79555800 | 1.98171300 |
| N | -0.19141100 | -1.97550200 | 1.10861400 |
| C | -0.33184800 | -1.19048700 | -0.00896300 |
| O | -1.42151900 | -1.05433100 | -0.57163000 |
| N | 0.80266100 | -0.54924000 | -0.42920700 |
| C | 2.06005400 | -0.58336000 | 0.17293500 |
| O | 2.95041600 | 0.14885100 | -0.26760600 |
| C | 2.16545000 | -1.51023500 | 1.28482700 |
| C | 1.02584700 | -2.12549800 | 1.70724300 |
| H | 0.35877600 | -4.69576800 | -1.36257400 |
| H | 0.72608600 | -0.02819900 | -1.31780100 |
| H | 1.03500000 | -2.77924000 | 2.57912000 |
| H | 3.28607800 | -2.62001200 | 2.70081400 |
| H | 5.56718200 | -3.85407500 | 0.52092800 |
| H | 6.53159300 | -1.14219600 | 1.38683300 |
| H | 5.10653000 | -1.76520800 | 3.96841900 |
| H | 6.05332000 | 0.90540300 | 2.84084500 |
| H | 4.63573800 | -1.55098900 | 0.22749400 |
| H | 7.05448400 | -0.22838200 | 3.79833000 |
| P | 7.75795600 | -3.62550000 | 1.49541200 |
| O | 6.97248800 | -3.92718900 | 0.21317600 |
| O | 8.50325500 | -4.62403500 | 2.30222400 |
| O | 8.72389200 | -2.33794100 | 1.10734300 |
| C | -5.65957500 | -2.70268100 | 5.37074200 |
| C | -5.10875500 | -1.36412200 | 4.87941200 |
| C | -4.08074300 | -1.50978300 | 3.75952300 |
| C | -1.39574200 | -2.61591400 | 1.63748500 |
| S | -4.72409300 | -2.41440400 | 2.28635900 |
| C | -4.92194700 | -1.06943300 | 1.05552500 |
| C | -6.36524200 | -0.91348400 | 0.60879000 |
| O | -6.89070300 | -2.16179000 | 0.17898600 |
| C | -7.33256200 | -0.40377200 | 1.67590800 |
| O | -7.30917300 | 1.01682400 | 1.70834500 |
| C | -8.66967100 | -0.98248200 | 1.18270800 |
| O | -9.15177900 | -0.20013500 | 0.11413400 |
| C | -8.22172700 | -2.34701500 | 0.67475500 |
| H | -6.16812400 | -3.23898100 | 4.55507600 |
| H | -5.93622400 | -0.71777600 | 4.53635100 |
| H | -4.62736500 | -0.82530400 | 5.71611200 |
| H | -3.20958200 | -2.07791500 | 4.12159400 |
| H | -3.69948400 | -0.53457400 | 3.42383200 |
| H | -1.77476000 | -3.37366100 | 0.93916300 |
| H | -2.17099500 | -1.86088300 | 1.78260300 |
| H | -1.15282800 | -3.06571600 | 2.60458400 |
| H | -8.87099400 | -2.70109700 | -0.13988600 |
| H | -8.88152100 | 0.71197100 | 0.30885400 |
| H | -7.06686600 | -0.83642400 | 2.65480300 |
| H | -6.39053400 | -0.18630000 | -0.22276000 |
| H | -4.54978600 | -0.12296300 | 1.47424100 |
| H | -7.65860400 | 1.30927400 | 2.55988500 |
| H | -9.42131000 | -1.06873400 | 1.99295500 |
| H | -4.29185600 | -1.30588500 | 0.18730900 |
| H | -8.21654100 | -3.09174500 | 1.49147800 |
| H | -6.38069600 | -2.56600600 | 6.19185300 |
| H | -4.84733200 | -3.34983400 | 5.74180600 |
| H | -3.98695400 | -6.96591900 | -1.13559200 |
| H | -5.31788900 | -5.79790200 | -0.90863200 |
| H | -5.80213000 | 1.23211200 | -2.30960000 |
| H | -6.11315700 | 0.43231700 | -3.88068000 |
| H | -0.35338400 | -8.53574500 | -4.59182500 |
| H | 0.83987500 | -8.75212700 | -5.89473700 |
| H | 5.89617900 | 0.42728700 | -2.42020700 |
| H | 9.50965600 | -2.39578000 | 1.66906500 |
| H | 6.04882800 | 1.20412300 | 5.25488900 |
| H | 2.18999300 | 7.60137500 | 1.58642300 |
| H | 2.12733900 | 8.52937700 | -3.08141000 |
| H | -1.77336100 | 6.18399600 | -3.32216800 |
| H | -6.42213100 | 5.86763600 | 0.21958100 |
| H | -6.37415700 | 4.07712900 | 0.38566200 |
| H | -0.28932400 | 1.29206300 | 7.19750800 |
| H | -2.03733200 | 1.07196600 | 6.86786300 |
| H | -2.76730000 | 4.66615000 | 3.05199600 |

#### Model3' – Initial state

| | | | |
|---|---|---|---|
| O | 1.69716100 | -3.71251100 | -0.57917200 |
| H | 1.39289700 | -4.34650300 | 0.12301600 |
| H | 2.65453300 | -3.83412100 | -0.71163700 |
| O | 0.71054600 | -3.59728800 | 4.79176200 |
| H | 0.63391100 | -3.79759800 | 5.73399500 |
| H | 0.62592800 | -4.49622100 | 4.33789500 |
| O | -0.91785100 | 3.23738900 | 1.58690400 |
| H | -1.48051400 | 3.54679900 | 2.35086000 |
| H | -0.02122600 | 3.42979200 | 1.89707500 |
| C | 1.65739900 | -8.40668500 | 2.88077300 |
| C | 1.44864900 | -7.31140600 | 1.82641000 |
| C | 0.86276700 | -5.99621600 | 2.35429400 |
| O | 0.60206200 | -5.07465500 | 1.49758100 |
| O | 0.67791200 | -5.88609500 | 3.58055100 |
| H | 0.70152500 | -8.68503700 | 3.34897700 |
| H | 0.79408500 | -7.65834300 | 1.00857800 |
| H | 2.40399600 | -7.04809200 | 1.33629200 |
| C | -5.87802300 | -2.14465000 | 1.93531100 |
| C | -4.83215800 | -3.26685500 | 1.90046700 |
| N | -3.47202300 | -2.79159600 | 2.23354900 |
| C | -2.30981000 | -3.40428000 | 1.91210600 |
| N | -2.22254000 | -4.23870300 | 0.87900300 |
| N | -1.18580700 | -3.15133300 | 2.60707700 |
| H | -5.56975200 | -1.29991200 | 1.29917600 |
| H | -6.82612700 | -2.53706200 | 1.54038400 |
| H | -4.83365100 | -3.73774000 | 0.90927600 |
| H | -5.11595200 | -4.06151100 | 2.61327700 |
| H | -3.40862400 | -2.11718600 | 2.98861700 |
| H | -2.80065800 | -4.10193300 | 0.01875700 |
| H | -1.31972100 | -4.70808400 | 0.77796300 |
| H | -1.19231100 | -2.90499600 | 3.59263500 |
| H | -0.36772000 | -3.71941700 | 2.31856900 |
| C | -3.36156700 | 6.45735200 | 0.64955100 |
| C | -4.51871300 | 5.85295100 | -0.14433500 |
| C | -5.59821500 | 5.23049400 | 0.75459600 |
| N | -5.07903400 | 4.25703500 | 1.69554700 |
| C | -4.66347700 | 3.03674000 | 1.36070200 |
| N | -4.95416800 | 2.49027400 | 0.15853400 |
| N | -3.97254300 | 2.31693200 | 2.25142200 |
| H | -2.81938200 | 5.68240600 | 1.21390500 |
| H | -2.64185600 | 6.95972400 | -0.01617100 |
| H | -4.13127800 | 5.08576900 | -0.83853600 |
| H | -5.00092200 | 6.62125800 | -0.77325500 |
| H | -6.40609100 | 4.78800200 | 0.14425800 |

| | | | |
|---|---|---|---|
| H | -6.07349300 | 6.02055400 | 1.35584400 |
| H | -4.74130100 | 4.62969000 | 2.63217700 |
| H | -5.46198900 | 3.01292900 | -0.53886700 |
| H | -4.47212100 | 1.63474200 | -0.11937300 |
| H | -3.43306800 | 2.86364500 | 2.95765200 |
| H | -3.52274800 | 1.48502300 | 1.86993400 |
| C | -3.40908200 | 6.00786600 | 6.65838000 |
| C | -2.41853900 | 5.39381800 | 5.66360300 |
| C | -3.03185600 | 4.83378400 | 4.37174900 |
| O | -4.13952300 | 5.27637400 | 3.99436500 |
| O | -2.35580200 | 3.94172200 | 3.75826700 |
| H | -2.88301300 | 6.44835800 | 7.51998200 |
| H | -1.82925800 | 4.58416600 | 6.12267500 |
| H | -1.67944000 | 6.15267200 | 5.34571400 |
| C | 4.22709300 | 6.39566000 | 1.86592700 |
| C | 2.84076500 | 6.41607200 | 1.26380600 |
| O | 1.97404700 | 7.18601300 | 1.64489000 |
| H | 4.88117100 | 5.62131200 | 1.43777900 |
| N | 2.61151000 | 5.51522700 | 0.26553400 |
| C | 1.32507800 | 5.48725100 | -0.37083700 |
| C | 1.15661800 | 4.22249400 | -1.19264100 |
| O | 2.09993300 | 3.61864300 | -1.68811300 |
| H | 3.32782200 | 4.89172500 | -0.09549500 |
| H | 0.53375100 | 5.57432400 | 0.39049700 |
| N | -0.14571400 | 3.87926000 | -1.37632500 |
| C | -0.52188900 | 2.62143500 | -1.99307900 |
| C | 0.06352100 | 2.42375300 | -3.41077200 |
| O | 0.61549200 | 1.36906000 | -3.71815300 |
| C | -2.04848900 | 2.54798100 | -2.03988600 |
| H | -0.78522900 | 4.18893300 | -0.64550400 |
| H | -0.11674400 | 1.77623500 | -1.41853000 |
| H | -2.45798500 | 3.38467900 | -2.62786900 |
| H | -2.45319400 | 2.61177400 | -1.02091300 |
| H | -2.40704300 | 1.60905400 | -2.48020000 |
| N | -0.16824700 | 3.42815200 | -4.29374000 |
| C | 0.12697400 | 3.36200100 | -5.72029400 |
| H | -0.59933400 | 4.26907400 | -3.92596400 |
| H | 0.40608400 | 2.32719000 | -5.95727400 |
| C | 6.06106100 | -5.11904800 | -0.20372200 |
| C | 4.99088800 | -4.06541200 | -0.00621300 |
| O | 4.49043500 | -3.47568300 | -0.97879300 |
| H | 6.16587800 | -5.78829200 | 0.66161000 |
| N | 4.62025100 | -3.79232600 | 1.25998600 |
| C | 3.95405700 | -2.55633200 | 1.62996900 |
| C | 5.03302200 | -1.59375800 | 2.16478700 |
| O | 6.21661200 | -1.90409500 | 2.18206500 |
| C | 2.79288500 | -2.78469800 | 2.58791100 |
| O | 3.26988300 | -3.34607000 | 3.77921400 |
| H | 5.11243400 | -4.24710300 | 2.02153100 |
| H | 3.52965800 | -2.12049700 | 0.71886300 |
| H | 2.07592700 | -3.44637900 | 2.07586900 |
| H | 2.27530600 | -1.81978500 | 2.74776300 |
| H | 2.46789200 | -3.48856700 | 4.32822900 |
| N | 4.59533200 | -0.37640400 | 2.56777900 |
| C | 5.51021600 | 0.53493400 | 3.23239500 |
| H | 3.60089500 | -0.17126100 | 2.53687700 |
| H | 6.42330100 | 0.66480400 | 2.63212300 |
| C | -6.05597000 | 0.24047300 | -3.03614700 |
| O | -5.91166100 | 1.18541500 | -4.07507000 |
| C | -4.93542000 | -0.77407200 | -2.91114800 |
| O | -3.73716700 | -0.21258300 | -2.37627500 |
| C | -5.30176800 | -1.93865400 | -1.97626700 |
| O | -5.87737900 | -2.96083700 | -2.74090600 |
| C | -3.94776500 | -2.28635900 | -1.28231100 |
| O | -3.50524900 | -3.60351600 | -1.42158600 |
| C | -2.95733300 | -1.32665300 | -1.95743400 |
| N | 0.43402900 | -1.39721900 | -0.33151800 |
| C | 0.48287100 | -0.32617500 | 0.53568400 |
| O | 1.49365400 | -0.00181000 | 1.14593400 |
| N | -0.72360100 | 0.35079200 | 0.63655700 |
| C | -1.92029700 | 0.00036300 | 0.00238000 |
| O | -2.98041000 | 0.49684600 | 0.38525800 |
| C | -1.79560200 | -0.94765400 | -1.08839900 |
| C | -0.63481600 | -1.65592200 | -1.13748100 |
| H | 1.15945100 | -2.15272000 | -0.30517900 |
| H | -0.75401900 | 1.20303800 | 1.21845200 |
| H | -0.49911700 | -2.50276000 | -1.81359600 |
| H | -2.56770200 | -1.86435500 | -2.84324300 |
| H | -4.34289500 | -4.20993800 | -1.29545600 |
| H | -6.01823100 | -1.59473700 | -1.21012300 |
| H | -4.72632000 | -1.22183900 | -3.90404900 |
| H | -6.12616500 | 0.80670700 | -2.09239200 |
| H | -4.02654900 | -1.99226900 | -0.22343100 |
| H | -6.99273200 | -0.34150100 | -3.13899000 |
| P | -6.65907800 | -4.21885400 | -1.95359900 |
| O | -5.63518500 | -4.75763700 | -0.95114100 |
| O | -7.36411000 | -5.03309500 | -2.97547200 |
| O | -7.74409400 | -3.37049400 | -1.02607700 |
| C | 4.40589300 | 0.84266700 | -6.05950400 |
| C | 4.60837300 | 1.71086600 | -4.81010200 |
| C | 3.77978400 | 1.29406600 | -3.59540600 |
| C | 2.66567300 | -1.26425900 | -3.04444300 |
| S | 4.26213600 | -0.39988400 | -3.02559300 |
| C | 4.53948200 | -0.30913900 | -1.21842600 |
| C | 5.91689400 | 0.16011500 | -0.80470000 |
| O | 6.92940500 | -0.54957000 | -1.49158900 |
| C | 6.30953200 | 1.63212400 | -1.01711800 |
| O | 5.63385700 | 2.54873300 | -0.21063500 |
| C | 7.82683900 | 1.53633000 | -0.74568400 |
| O | 7.94434500 | 1.67373200 | 0.66180400 |
| C | 8.17384300 | 0.12082800 | -1.25918300 |
| H | 4.71698100 | -0.20023100 | -5.88181500 |
| H | 5.67894900 | 1.74381800 | -4.54190500 |
| H | 4.31981400 | 2.75310200 | -5.02858900 |
| H | 2.70110300 | 1.24228100 | -3.81566000 |
| H | 3.89305200 | 1.96895600 | -2.73518700 |
| H | 2.40679800 | -1.41895800 | -4.09935800 |
| H | 2.81222300 | -2.22425200 | -2.53311600 |
| H | 1.90269600 | -0.64221100 | -2.55979100 |
| H | 8.76564500 | -0.42049500 | -0.49984300 |
| H | 8.87020300 | 1.62745300 | 0.92986600 |
| H | 6.14898200 | 1.90991900 | -2.07162500 |
| H | 5.96335600 | -0.03707700 | 0.28279400 |
| H | 3.74745900 | 0.29461000 | -0.75089800 |
| H | 6.08450400 | 2.52927000 | 0.65016500 |
| H | 8.39780200 | 2.32632400 | -1.26332000 |
| H | 4.42872900 | -1.37145200 | -0.94827600 |
| H | 2.09953900 | -9.30781900 | 2.42769900 |
| H | 2.32036600 | -8.05240300 | 3.68377300 |
| H | 7.01678800 | -4.59310000 | -0.35884300 |
| H | 5.83769300 | -5.69843700 | -1.10898500 |
| H | 5.81030900 | 0.15653800 | 4.22385400 |
| H | 5.01829000 | 1.50953600 | 3.35800200 |
| H | 8.73520200 | 0.13860000 | -2.20611700 |
| H | 5.00147300 | 1.22380100 | -6.90189400 |
| H | 3.34735000 | 0.83347200 | -6.36560000 |
| H | 4.69100000 | 7.38275100 | 1.71895900 |
| H | 4.13839500 | 6.24031700 | 2.95145700 |
| H | 1.18064100 | 6.35177800 | -1.04958700 |
| H | 0.96139300 | 4.02850700 | -5.99690500 |
| H | -0.75990900 | 3.63988300 | -6.31021000 |
| H | -3.71906800 | 7.19609800 | 1.38416100 |
| H | -4.00730300 | 6.79048400 | 6.16925200 |
| H | -4.11225800 | 5.24904200 | 7.03696100 |
| H | -5.90270100 | 0.70098500 | -4.91096000 |
| H | -8.59635500 | -3.40539500 | -1.48302400 |
| H | -6.04185400 | -1.76305500 | 2.95669100 |

#### Model3' – TS1

| | | | |
|---|---|---|---|
| O | 1.37257100 | -4.06259200 | -0.59245400 |

| | | | |
|---|---|---|---|
| H | 0.94121300 | -4.65017000 | 0.38286700 |
| H | 2.35409300 | -4.03505900 | -0.71308500 |
| O | 0.31708300 | -3.75667100 | 4.74200800 |
| H | 0.30572400 | -3.98294400 | 5.68152500 |
| H | 0.07344700 | -4.60066300 | 4.27473100 |
| O | -0.57470300 | 1.75535200 | 2.94553900 |
| H | -1.01685900 | 2.55997500 | 3.30123600 |
| H | -0.53466900 | 1.19037800 | 3.72877300 |
| C | 1.23325900 | -8.40847700 | 2.68813700 |
| C | 1.34786300 | -7.18261800 | 1.76634300 |
| C | 0.50760400 | -5.99084800 | 2.20605000 |
| O | 0.34027000 | -4.99669100 | 1.36833000 |
| O | 0.01177900 | -5.96835200 | 3.33048500 |
| H | 0.20170400 | -8.79027600 | 2.70508400 |
| H | 1.08697600 | -7.42758100 | 0.72320200 |
| H | 2.38901400 | -6.80931600 | 1.73831000 |
| C | -6.01708200 | -1.80146700 | 1.86952000 |
| C | -5.02171500 | -2.95750700 | 1.81067200 |
| N | -3.66668000 | -2.52806500 | 2.19843900 |
| C | -2.51147700 | -3.12423000 | 1.85794200 |
| N | -2.42259800 | -3.92969300 | 0.80679300 |
| N | -1.39751200 | -2.87908500 | 2.58244400 |
| H | -5.66203500 | -0.94774100 | 1.27085400 |
| H | -6.97553400 | -2.13505600 | 1.44617500 |
| H | -5.01320500 | -3.38815000 | 0.80214200 |
| H | -5.34551900 | -3.76981000 | 2.48548100 |
| H | -3.60131500 | -1.80396700 | 2.90523700 |
| H | -2.99933600 | -3.77595600 | -0.06408700 |
| H | -1.53856100 | -4.42529900 | 0.70877100 |
| H | -1.44422100 | -2.71222800 | 3.58251900 |
| H | -0.57301100 | -3.41154000 | 2.29982300 |
| C | -3.12158900 | 6.70660800 | 0.77006600 |
| C | -4.18234600 | 5.92172900 | -0.00284600 |
| C | -5.21435400 | 5.23740300 | 0.90921300 |
| N | -4.63642600 | 4.26889900 | 1.81823100 |
| C | -4.26713400 | 3.03368700 | 1.47025500 |
| N | -4.60660800 | 2.49851600 | 0.27257900 |
| N | -3.58433600 | 2.28645900 | 2.33696100 |
| H | -2.60390000 | 6.06486200 | 1.49979100 |
| H | -2.37051000 | 7.14418700 | 0.09256000 |
| H | -3.69825900 | 5.15331200 | -0.63191800 |
| H | -4.72266400 | 6.58954400 | -0.69638800 |
| H | -6.00952300 | 4.76710100 | 0.30402500 |
| H | -5.71642100 | 5.99756600 | 1.52803500 |
| H | -4.26142900 | 4.62754000 | 2.74575900 |
| H | -5.01404000 | 3.07712500 | -0.44648600 |
| H | -4.13356800 | 1.63503800 | -0.01160400 |
| H | -3.11908400 | 2.76681800 | 3.14103200 |
| H | -3.11073000 | 1.47131900 | 1.93924500 |
| C | -3.20012900 | 6.13191000 | 6.76795700 |
| C | -2.21697900 | 5.21034300 | 6.03962000 |
| C | -2.71850100 | 4.62784200 | 4.70773600 |
| O | -3.63207100 | 5.24137900 | 4.10499500 |
| O | -2.16178400 | 3.56133700 | 4.30146500 |
| H | -2.74523900 | 6.56292400 | 7.67410900 |
| H | -1.90751100 | 4.36295200 | 6.67223600 |
| H | -1.28579800 | 5.75857000 | 5.80433300 |
| C | 4.45476500 | 6.28443400 | 1.99199500 |
| C | 3.22661500 | 6.46208500 | 1.11815600 |
| O | 2.48178200 | 7.42056600 | 1.24415600 |
| H | 5.10229900 | 5.45271400 | 1.67573800 |
| N | 2.98245800 | 5.48230400 | 0.19569500 |
| C | 1.81148600 | 5.57650900 | -0.63771600 |
| C | 1.43337100 | 4.22303000 | -1.22070400 |
| O | 2.27362500 | 3.39922400 | -1.55872300 |
| H | 3.60981300 | 4.69829400 | 0.03931100 |
| H | 0.99405800 | 6.02078700 | -0.04978400 |
| N | 0.09835200 | 4.04479900 | -1.40999400 |
| C | -0.44872800 | 2.80638100 | -1.95128600 |
| C | 0.04485300 | 2.55884100 | -3.40134500 |
| O | 0.43731200 | 1.45640000 | -3.76235200 |
| C | -1.97103500 | 2.84307300 | -1.89246800 |
| H | -0.54070200 | 4.67659000 | -0.93934600 |
| H | -0.06553400 | 1.94854500 | -1.37992700 |
| H | -2.36722400 | 3.70869800 | -2.44884400 |
| H | -2.30156000 | 2.90904900 | -0.84739700 |
| H | -2.41150500 | 1.93068300 | -2.31732000 |
| N | -0.05497100 | 3.63021400 | -4.23626900 |
| C | 0.23907800 | 3.59672600 | -5.66119900 |
| H | -0.34735000 | 4.50778500 | -3.82240300 |
| H | 0.41500900 | 2.54952000 | -5.94021800 |
| C | 5.78335100 | -5.25336800 | -0.32138500 |
| C | 4.68979100 | -4.25354200 | -0.02790100 |
| O | 4.01764400 | -3.76217600 | -0.95671200 |
| H | 6.17343500 | -5.74429600 | 0.58036300 |
| N | 4.49941000 | -3.91169800 | 1.25311800 |
| C | 3.81709500 | -2.69712100 | 1.67136300 |
| C | 4.87417300 | -1.84548000 | 2.38677300 |
| O | 5.94234600 | -2.31707200 | 2.75463400 |
| C | 2.56816800 | -2.94854800 | 2.51477300 |
| O | 2.85480900 | -3.82465100 | 3.57041700 |
| H | 5.15332900 | -4.25810800 | 1.95121500 |
| H | 3.46397300 | -2.15864800 | 0.78415900 |
| H | 1.80530800 | -3.35904300 | 1.83851400 |
| H | 2.19106600 | -1.96939500 | 2.85808100 |
| H | 2.04705800 | -3.84722300 | 4.12432900 |
| N | 4.55030800 | -0.54174800 | 2.55193600 |
| C | 5.47572400 | 0.34507800 | 3.23418500 |
| H | 3.65569400 | -0.20412000 | 2.19996800 |
| H | 6.38649600 | 0.51380100 | 2.63484900 |
| C | -6.03720400 | 0.71464900 | -3.01879800 |
| O | -5.89268900 | 1.65402100 | -4.06845600 |
| C | -4.97076700 | -0.36564300 | -2.93022700 |
| O | -3.72794400 | 0.10412100 | -2.40598500 |
| C | -5.39017800 | -1.53926300 | -2.01909300 |
| O | -6.01753400 | -2.51890000 | -2.80503700 |
| C | -4.04484500 | -1.95913600 | -1.34795400 |
| O | -3.70320500 | -3.31299200 | -1.45217900 |
| C | -3.00397800 | -1.08329300 | -2.06301300 |
| N | 0.54146200 | -1.57741600 | -0.85232300 |
| C | 0.68147100 | -0.67711000 | 0.17097400 |
| O | 1.75414600 | -0.49876600 | 0.76834000 |
| N | -0.46024100 | 0.04025500 | 0.52936900 |
| C | -1.71142800 | -0.07566600 | -0.06965800 |
| O | -2.69917900 | 0.46645600 | 0.45588400 |
| C | -1.74328600 | -0.84928800 | -1.28418400 |
| C | -0.61602900 | -1.59224800 | -1.55159300 |
| H | 1.08522500 | -3.06558400 | -0.66453900 |
| H | -0.40699600 | 0.66024000 | 1.35505200 |
| H | -0.64487700 | -2.30349000 | -2.38600000 |
| H | -2.73471600 | -1.62335000 | -2.99151800 |
| H | -4.58140000 | -3.86372300 | -1.35544400 |
| H | -6.08769000 | -1.18336600 | -1.24113800 |
| H | -4.80664100 | -0.80211500 | -3.93686300 |
| H | -6.04207100 | 1.28471100 | -2.07496300 |
| H | -4.07882600 | -1.63015000 | -0.29659300 |
| H | -7.00725800 | 0.18397700 | -3.08464100 |
| P | -6.88154300 | -3.75599600 | -2.06350300 |
| O | -5.90534200 | -4.37117500 | -1.05764100 |
| O | -7.60789400 | -4.50626900 | -3.11935000 |
| O | -7.93890700 | -2.87576200 | -1.13380000 |
| C | 4.40352600 | 0.89892200 | -6.05011300 |
| C | 4.37914500 | 1.71469000 | -4.74809600 |
| C | 3.60080900 | 1.10562500 | -3.57909100 |
| C | 2.91336000 | -1.65574900 | -3.22476200 |
| S | 4.32321700 | -0.52252900 | -3.07957100 |
| C | 4.49249400 | -0.49229200 | -1.25414300 |
| C | 5.85194700 | -0.03647300 | -0.76893500 |
| O | 6.88320700 | -0.79596600 | -1.37791300 |
| C | 6.27160900 | 1.42476400 | -1.00827200 |

| | | | |
|---|---|---|---|
| O | 5.61657400 | 2.36371600 | -0.21105500 |
| C | 7.78816800 | 1.31763600 | -0.74132300 |
| O | 7.90982300 | 1.43917400 | 0.66831700 |
| C | 8.12696300 | -0.09041400 | -1.26275500 |
| H | 4.90026600 | -0.07531800 | -5.90847600 |
| H | 5.41238200 | 1.93549800 | -4.42683100 |
| H | 3.90846900 | 2.69554700 | -4.93162200 |
| H | 2.54046000 | 0.92123300 | -3.81687800 |
| H | 3.60857600 | 1.75153100 | -2.68843100 |
| H | 2.59081000 | -1.62838800 | -4.27298500 |
| H | 3.29141800 | -2.64804900 | -2.94486000 |
| H | 2.10910200 | -1.34615100 | -2.53777600 |
| H | 8.79995900 | -0.61024600 | -0.55900600 |
| H | 8.83972900 | 1.47329400 | 0.92387100 |
| H | 6.10779100 | 1.69049800 | -2.06643200 |
| H | 5.82830700 | -0.19579400 | 0.32623700 |
| H | 3.66813600 | 0.08843800 | -0.81324200 |
| H | 6.07450300 | 2.34750600 | 0.64602700 |
| H | 8.36180300 | 2.10905100 | -1.25337700 |
| H | 4.34399400 | -1.55295000 | -1.00372900 |
| H | 1.89782800 | -9.21428700 | 2.34207800 |
| H | 1.50494100 | -8.14140200 | 3.71891400 |
| H | 6.60425300 | -4.71053900 | -0.81690700 |
| H | 5.40774600 | -6.00536700 | -1.02919200 |
| H | 5.78577700 | -0.08775400 | 4.19697000 |
| H | 4.98159500 | 1.30941400 | 3.41548100 |
| H | 8.60400300 | -0.06273700 | -2.25599800 |
| H | 4.95266700 | 1.43840800 | -6.83561200 |
| H | 3.38195100 | 0.70903000 | -6.41736000 |
| H | 5.02842700 | 7.22200700 | 1.98505100 |
| H | 4.12430300 | 6.10958200 | 3.02742200 |
| H | 1.97259200 | 6.26197200 | -1.49524400 |
| H | 1.13824700 | 4.18720700 | -5.90781300 |
| H | -0.60995900 | 3.98493400 | -6.24531700 |
| H | -3.57608900 | 7.53082900 | 1.34271400 |
| H | -3.51793900 | 6.95017900 | 6.10547900 |
| H | -4.10836200 | 5.58478800 | 7.06715000 |
| H | -5.91292900 | 1.16278100 | -4.90029100 |
| H | -8.77789000 | -2.83911000 | -1.61444000 |
| H | -6.18236100 | -1.45041600 | 2.90180500 |

#### Model3' – Intermediate

| | | | |
|---|---|---|---|
| O | 1.38918000 | -4.09301500 | -0.62420000 |
| H | 0.82239800 | -4.83640500 | 0.60018200 |
| H | 2.35176400 | -4.04654500 | -0.80611200 |
| O | 0.20162500 | -3.77129700 | 4.73966000 |
| H | 0.25967300 | -3.98530700 | 5.68052000 |
| H | -0.09193900 | -4.61036100 | 4.30560500 |
| O | -0.60607400 | 1.65163600 | 3.02021100 |
| H | -1.02249400 | 2.46808100 | 3.37935800 |
| H | -0.65307800 | 1.05553200 | 3.77941800 |
| C | 1.14296800 | -8.43282200 | 2.67769000 |
| C | 1.30275500 | -7.16828900 | 1.81951500 |
| C | 0.41020200 | -6.02627400 | 2.26016700 |
| O | 0.24414300 | -4.99384900 | 1.43970300 |
| O | -0.14785600 | -6.02652500 | 3.34422900 |
| H | 0.12114000 | -8.83245900 | 2.60035100 |
| H | 1.12118100 | -7.36934100 | 0.75056200 |
| H | 2.33184000 | -6.76789800 | 1.88606500 |
| C | -6.04556600 | -1.76250300 | 1.82799200 |
| C | -5.03970000 | -2.90659600 | 1.81002100 |
| N | -3.71284100 | -2.45600700 | 2.25621700 |
| C | -2.52701500 | -2.96711900 | 1.89588200 |
| N | -2.38271200 | -3.72121900 | 0.81746300 |
| N | -1.43963200 | -2.67772700 | 2.65171100 |
| H | -5.67295800 | -0.90517900 | 1.24571600 |
| H | -6.98446300 | -2.10215400 | 1.36678300 |
| H | -4.98157100 | -3.33372800 | 0.80260200 |
| H | -5.38096100 | -3.72399100 | 2.46966800 |
| H | -3.68803000 | -1.72415900 | 2.95755500 |
| H | -2.98738200 | -3.62245400 | -0.04978300 |
| H | -1.49204200 | -4.20046500 | 0.72379400 |
| H | -1.51927600 | -2.59819400 | 3.66024100 |
| H | -0.56827200 | -3.11184000 | 2.36202700 |
| C | -3.07088600 | 6.72267700 | 0.76354100 |
| C | -4.10249400 | 5.90773000 | -0.01814200 |
| C | -5.14283400 | 5.22114800 | 0.88283400 |
| N | -4.57287000 | 4.25538000 | 1.79978000 |
| C | -4.22340900 | 3.01052400 | 1.46573000 |
| N | -4.55916500 | 2.47096100 | 0.26851900 |
| N | -3.56612800 | 2.25615800 | 2.34577500 |
| H | -2.57212100 | 6.10633700 | 1.52763000 |
| H | -2.30208900 | 7.14656700 | 0.09731700 |
| H | -3.59408400 | 5.13622800 | -0.62359700 |
| H | -4.63786700 | 6.55546000 | -0.73426100 |
| H | -5.92808000 | 4.74597500 | 0.26893900 |
| H | -5.65588500 | 5.98083700 | 1.49340500 |
| H | -4.20276400 | 4.61438200 | 2.72888000 |
| H | -4.92762600 | 3.05551800 | -0.46678500 |
| H | -4.09518400 | 1.59665100 | 0.00056100 |
| H | -3.11183700 | 2.72970300 | 3.15977600 |
| H | -3.09816400 | 1.43211800 | 1.95782700 |
| C | -3.19121400 | 6.13685200 | 6.75966800 |
| C | -2.21876200 | 5.17672100 | 6.06715000 |
| C | -2.70425000 | 4.59052500 | 4.73046700 |
| O | -3.58057400 | 5.22525600 | 4.09469800 |
| O | -2.17262500 | 3.50060200 | 4.35481600 |
| H | -2.74491400 | 6.56753800 | 7.67024300 |
| H | -1.94956700 | 4.32954300 | 6.71819100 |
| H | -1.26656600 | 5.69456300 | 5.84764900 |
| C | 4.49393400 | 6.23337800 | 2.03105700 |
| C | 3.26504200 | 6.42408800 | 1.16037600 |
| O | 2.52329700 | 7.38343300 | 1.29823600 |
| H | 5.13931500 | 5.40404500 | 1.70429900 |
| N | 3.01680300 | 5.45544100 | 0.22756800 |
| C | 1.84514000 | 5.56116200 | -0.60369400 |
| C | 1.46621500 | 4.21415100 | -1.20195400 |
| O | 2.30603300 | 3.38985000 | -1.53856100 |
| H | 3.64110400 | 4.67082600 | 0.06170300 |
| H | 1.02809000 | 5.99858800 | -0.00999000 |
| N | 0.13189200 | 4.04250700 | -1.40449000 |
| C | -0.41472200 | 2.80512900 | -1.94925700 |
| C | 0.07790300 | 2.55733100 | -3.39925400 |
| O | 0.44856000 | 1.44950200 | -3.76546700 |
| C | -1.93664100 | 2.83989600 | -1.89085200 |
| H | -0.50829000 | 4.67346800 | -0.93424600 |
| H | -0.03209000 | 1.94638500 | -1.37889900 |
| H | -2.33378400 | 3.70837200 | -2.44210700 |
| H | -2.26742200 | 2.89963600 | -0.84535300 |
| H | -2.37568100 | 1.92973300 | -2.32152700 |
| N | 0.00132900 | 3.63414300 | -4.23002200 |
| C | 0.30261600 | 3.59688000 | -5.65327400 |
| H | -0.27139800 | 4.51672800 | -3.81347600 |
| H | 0.45350600 | 2.54615100 | -5.93361800 |
| C | 5.73826000 | -5.31078500 | -0.29747900 |
| C | 4.68144000 | -4.25969700 | -0.04690800 |
| O | 4.09584500 | -3.71548000 | -0.99921100 |
| H | 6.04846000 | -5.84067600 | 0.61341600 |
| N | 4.42834300 | -3.94287500 | 1.23327900 |
| C | 3.76663800 | -2.71659400 | 1.64603500 |
| C | 4.81660300 | -1.89979700 | 2.41292400 |
| O | 5.84722100 | -2.40864100 | 2.83377800 |
| C | 2.47935100 | -2.95418800 | 2.43857100 |
| O | 2.67849500 | -3.93417000 | 3.42555800 |
| H | 5.02876100 | -4.33142800 | 1.95612700 |
| H | 3.45673800 | -2.15724600 | 0.75606300 |
| H | 1.71692400 | -3.25867300 | 1.70815500 |
| H | 2.15368100 | -1.98673000 | 2.85748600 |
| H | 1.89233500 | -3.90288500 | 4.00737300 |
| N | 4.52843500 | -0.58462500 | 2.56211100 |

| | | | |
|---|---|---|---|
| C | 5.45652100 | 0.28307600 | 3.26745800 |
| H | 3.66482500 | -0.21939200 | 2.16202100 |
| H | 6.35614700 | 0.49088800 | 2.66249200 |
| C | -6.01847600 | 0.76309200 | -3.05647100 |
| O | -5.85518800 | 1.70351200 | -4.10211300 |
| C | -4.96094400 | -0.32495600 | -2.96025500 |
| O | -3.71777000 | 0.13700500 | -2.43191800 |
| C | -5.39461200 | -1.49464400 | -2.05043600 |
| O | -6.00658900 | -2.47952600 | -2.84366300 |
| C | -4.05942400 | -1.90568200 | -1.35408600 |
| O | -3.73473300 | -3.26765800 | -1.40987800 |
| C | -3.00154600 | -1.05535400 | -2.07764500 |
| N | 0.60327700 | -1.43385600 | -0.98819700 |
| C | 0.71039100 | -0.62072600 | 0.10605400 |
| O | 1.77560300 | -0.45342000 | 0.72360800 |
| N | -0.45574200 | 0.02458000 | 0.52728900 |
| C | -1.70963900 | -0.10058000 | -0.06228900 |
| O | -2.70806100 | 0.38697800 | 0.50199000 |
| C | -1.73438000 | -0.81345700 | -1.31126600 |
| C | -0.56560100 | -1.45864500 | -1.66069000 |
| H | 1.10861100 | -3.13910500 | -0.73491100 |
| H | -0.41142300 | 0.59894700 | 1.38464200 |
| H | -0.57358000 | -2.09306300 | -2.55602600 |
| H | -2.73915800 | -1.60695000 | -3.00144900 |
| H | -4.62340000 | -3.80779500 | -1.32561700 |
| H | -6.10705400 | -1.13749700 | -1.28686400 |
| H | -4.79540200 | -0.76474700 | -3.96525200 |
| H | -6.02874100 | 1.33107800 | -2.11136400 |
| H | -4.10493600 | -1.54535800 | -0.31366800 |
| H | -6.99208100 | 0.24018700 | -3.13342500 |
| P | -6.90247400 | -3.70182400 | -2.11420200 |
| O | -5.95950900 | -4.30255500 | -1.06761100 |
| O | -7.59633700 | -4.46620100 | -3.18172100 |
| O | -7.98625900 | -2.80465200 | -1.23329100 |
| C | 4.44621000 | 0.86436600 | -6.02210400 |
| C | 4.38797900 | 1.70105800 | -4.73509800 |
| C | 3.61725600 | 1.08690300 | -3.56401800 |
| C | 2.97415800 | -1.68866000 | -3.25308700 |
| S | 4.35922800 | -0.53143200 | -3.06805500 |
| C | 4.49573900 | -0.50879200 | -1.24219600 |
| C | 5.85216600 | -0.06622500 | -0.73854400 |
| O | 6.87960200 | -0.85564700 | -1.31729200 |
| C | 6.30227400 | 1.38297500 | -0.99478000 |
| O | 5.65734600 | 2.34671200 | -0.21859700 |
| C | 7.81333700 | 1.25238200 | -0.71041800 |
| O | 7.92215000 | 1.38580600 | 0.69975800 |
| C | 8.13155600 | -0.16618300 | -1.21400600 |
| H | 4.96967400 | -0.09236700 | -5.85981100 |
| H | 5.41237600 | 1.95303900 | -4.40848800 |
| H | 3.89384500 | 2.66626500 | -4.93867200 |
| H | 2.55818400 | 0.89264200 | -3.79867300 |
| H | 3.62264200 | 1.73405900 | -2.67436300 |
| H | 2.65644600 | -1.63528100 | -4.30183100 |
| H | 3.37418300 | -2.67947100 | -3.00150000 |
| H | 2.16326800 | -1.41493300 | -2.55404100 |
| H | 8.80027900 | -0.68628500 | -0.50614900 |
| H | 8.85000500 | 1.40421900 | 0.96409700 |
| H | 6.15440400 | 1.63610200 | -2.05847900 |
| H | 5.80775000 | -0.20789700 | 0.35831700 |
| H | 3.66643300 | 0.07257400 | -0.81178100 |
| H | 6.10471900 | 2.33254400 | 0.64392200 |
| H | 8.40678900 | 2.02837700 | -1.22360600 |
| H | 4.33984600 | -1.57004400 | -0.99900600 |
| H | 1.84736600 | -9.21038000 | 2.34769700 |
| H | 1.33425200 | -8.20719400 | 3.73589500 |
| H | 6.61419800 | -4.80400100 | -0.73295100 |
| H | 5.36599800 | -6.02732600 | -1.04293900 |
| H | 5.78546400 | -0.19437300 | 4.20169200 |
| H | 4.95581100 | 1.23228500 | 3.50281500 |
| H | 8.60784300 | -0.15533300 | -2.20830000 |
| H | 4.98547300 | 1.40552900 | -6.81338800 |
| H | 3.43354900 | 0.63844100 | -6.39390700 |
| H | 5.06959500 | 7.16973500 | 2.03440300 |
| H | 4.16403700 | 6.04706400 | 3.06466400 |
| H | 2.00632000 | 6.25709300 | -1.45267400 |
| H | 1.21815900 | 4.16459700 | -5.89347200 |
| H | -0.53269900 | 4.00830200 | -6.24130700 |
| H | -3.54899300 | 7.55986900 | 1.29691000 |
| H | -3.46940200 | 6.95427300 | 6.07856300 |
| H | -4.12180300 | 5.62068200 | 7.04499200 |
| H | -5.85912400 | 1.21306900 | -4.93466700 |
| H | -8.80875800 | -2.77159200 | -1.74184000 |
| H | -6.25588100 | -1.41417400 | 2.85306800 |

#### Model3' − TS2

| | | | |
|---|---|---|---|
| O | -0.80150800 | 4.26582900 | -0.70928400 |
| H | -0.17505700 | 5.00244600 | 0.47636800 |
| H | -1.77808000 | 4.35649100 | -0.85440600 |
| O | 0.41829100 | 4.07499100 | 4.58810900 |
| H | 0.36770600 | 4.30020800 | 5.52670600 |
| H | 0.75158600 | 4.89533100 | 4.14705400 |
| O | 0.98094400 | -1.28771000 | 3.34067800 |
| H | 1.34941000 | -2.12650400 | 3.71158200 |
| H | 0.94846000 | -0.70680300 | 4.11145000 |
| C | -0.11240000 | 8.73790000 | 2.22812000 |
| C | -0.44376400 | 7.42156900 | 1.51582300 |
| C | 0.34076800 | 6.24625600 | 2.05367900 |
| O | 0.41184400 | 5.14521800 | 1.31183200 |
| O | 0.89337000 | 6.27523000 | 3.13951400 |
| H | 0.94014800 | 9.01611900 | 2.06997500 |
| H | -0.28984000 | 7.48966800 | 0.42632000 |
| H | -1.50469300 | 7.14357600 | 1.66167900 |
| C | 6.28118000 | 1.29883600 | 1.40420800 |
| C | 5.45619900 | 2.57736900 | 1.38944000 |
| N | 4.12461300 | 2.34766700 | 1.96509200 |
| C | 2.98384800 | 2.96417500 | 1.63089600 |
| N | 2.85519100 | 3.63582400 | 0.49402100 |
| N | 1.92636600 | 2.87263600 | 2.46875400 |
| H | 5.73784400 | 0.47784800 | 0.91184400 |
| H | 7.21868100 | 1.46725300 | 0.85405300 |
| H | 5.37303100 | 2.95830500 | 0.36502700 |
| H | 5.96528600 | 3.36822500 | 1.96813400 |
| H | 4.06886300 | 1.69686300 | 2.74049800 |
| H | 3.37544500 | 3.39165600 | -0.39548500 |
| H | 2.02035400 | 4.20488300 | 0.40368700 |
| H | 2.05923400 | 2.80126500 | 3.47179100 |
| H | 1.10487400 | 3.41576000 | 2.21494300 |
| C | 2.38353700 | -6.86505700 | 1.02792300 |
| C | 3.42384300 | -6.22363300 | 0.10960600 |
| C | 4.63532200 | -5.66405600 | 0.87282500 |
| N | 4.30595400 | -4.58852000 | 1.78589300 |
| C | 4.13607400 | -3.31416900 | 1.42191900 |
| N | 4.40046600 | -2.89268700 | 0.16047500 |
| N | 3.74061700 | -2.42301900 | 2.32742900 |
| H | 2.06958700 | -6.17048900 | 1.82254500 |
| H | 1.48995300 | -7.18575000 | 0.46762900 |
| H | 2.96348800 | -5.40553200 | -0.47290800 |
| H | 3.78754000 | -6.95912100 | -0.62906400 |
| H | 5.41587300 | -5.33259400 | 0.16607900 |
| H | 5.09236200 | -6.46778500 | 1.47137200 |
| H | 3.97374800 | -4.84341100 | 2.75956000 |
| H | 4.57184800 | -3.55935400 | -0.57717200 |
| H | 4.08079700 | -1.95797500 | -0.10601500 |
| H | 3.31691600 | -2.76905500 | 3.22040900 |
| H | 3.40147900 | -1.53172900 | 1.95766600 |
| C | 2.92530000 | -5.96235600 | 6.96098100 |
| C | 2.06787400 | -4.93626100 | 6.21688400 |
| C | 2.64021900 | -4.46886000 | 4.87083500 |
| O | 3.32439600 | -5.28097700 | 4.20425000 |
| O | 2.36855900 | -3.27993400 | 4.51381300 |

| | | | |
|---|---|---|---|
| H | 2.42158100 | -6.31733500 | 7.87406400 |
| H | 1.88006300 | -4.04059200 | 6.82910200 |
| H | 1.07419300 | -5.36870900 | 5.99603800 |
| C | -4.99936200 | -5.49854200 | 2.66986800 |
| C | -3.78755200 | -5.88620800 | 1.84216000 |
| O | -3.10738700 | -6.86494700 | 2.09861200 |
| H | -5.53671000 | -4.61964800 | 2.28443200 |
| N | -3.49601700 | -5.06396400 | 0.78961300 |
| C | -2.34344600 | -5.30661400 | -0.03287100 |
| C | -1.91305300 | -4.02917400 | -0.74042600 |
| O | -2.69902800 | -3.11678100 | -0.97205600 |
| H | -4.05158000 | -4.24529000 | 0.56859500 |
| H | -1.53635300 | -5.72528900 | 0.58939700 |
| N | -0.61782100 | -4.01170600 | -1.14794000 |
| C | -0.01084600 | -2.84229400 | -1.77461600 |
| C | -0.48317800 | -2.61439600 | -3.22802400 |
| O | -0.57378400 | -1.47786500 | -3.67427400 |
| C | 1.50891000 | -2.97602700 | -1.73607700 |
| H | 0.00730100 | -4.71228700 | -0.76244400 |
| H | -0.32364800 | -1.93804000 | -1.23208900 |
| H | 1.82903700 | -3.90436700 | -2.23805600 |
| H | 1.86141700 | -2.99457600 | -0.69529000 |
| H | 2.00076600 | -2.13417500 | -2.24015600 |
| N | -0.70635300 | -3.73956100 | -3.95758300 |
| C | -1.02841000 | -3.75800500 | -5.37767000 |
| H | -0.63427700 | -4.62184400 | -3.46421900 |
| H | -0.97088700 | -2.72585600 | -5.74729700 |
| C | -5.17742300 | 5.96483700 | -0.29598300 |
| C | -4.11784400 | 4.91071900 | -0.04089500 |
| O | -3.45995200 | 4.42400900 | -0.97142600 |
| H | -5.49468600 | 6.48365200 | 0.61892500 |
| N | -3.97461300 | 4.53137400 | 1.24083800 |
| C | -3.39507900 | 3.26928800 | 1.65893800 |
| C | -4.46513400 | 2.60471900 | 2.53845800 |
| O | -5.43356100 | 3.23083600 | 2.95101000 |
| C | -2.03780800 | 3.40729400 | 2.35616000 |
| O | -2.07455100 | 4.43820700 | 3.31012100 |
| H | -4.64168800 | 4.88258300 | 1.92655300 |
| H | -3.23230600 | 2.64250400 | 0.77069400 |
| H | -1.30044300 | 3.60881600 | 1.56748200 |
| H | -1.77071400 | 2.43263700 | 2.79521700 |
| H | -1.27824100 | 4.32801800 | 3.86671800 |
| N | -4.27369200 | 1.29380200 | 2.80277500 |
| C | -5.26325700 | 0.58024900 | 3.58692500 |
| H | -3.45102500 | 0.82284600 | 2.43109400 |
| H | -6.19587400 | 0.43762200 | 3.01433900 |
| C | 5.73452200 | -1.47286900 | -3.32401700 |
| O | 5.37504400 | -2.45110300 | -4.28148200 |
| C | 4.79026100 | -0.28977600 | -3.19319800 |
| O | 3.57623000 | -0.60859100 | -2.50983800 |
| C | 5.40680600 | 0.88141500 | -2.40360800 |
| O | 6.05170100 | 1.74619400 | -3.29919900 |
| C | 4.18262800 | 1.47815100 | -1.64102000 |
| O | 3.95454200 | 2.84995900 | -1.80308800 |
| C | 3.00040600 | 0.66231000 | -2.18377900 |
| N | -0.42039300 | 1.32712800 | -0.72700900 |
| C | -0.40370800 | 0.67055500 | 0.48399100 |
| O | -1.39306000 | 0.61716100 | 1.22231400 |
| N | 0.78448400 | 0.03991200 | 0.83267600 |
| C | 1.94081200 | 0.01607700 | 0.06877800 |
| O | 2.98197200 | -0.47415800 | 0.53616500 |
| C | 1.82776100 | 0.58350200 | -1.25628400 |
| C | 0.64810900 | 1.21317300 | -1.55580400 |
| H | -0.66114500 | 3.29344900 | -0.66100600 |
| H | 0.82144800 | -0.43776500 | 1.75576900 |
| H | 0.53534200 | 1.69594400 | -2.53288300 |
| H | 2.67157000 | 1.16309000 | -3.11447400 |
| H | 4.89007100 | 3.31235300 | -1.80264100 |
| H | 6.13420900 | 0.49542300 | -1.66863700 |
| H | 4.55302600 | 0.10498100 | -4.20218200 |
| H | 5.79918500 | -1.98460400 | -2.34934100 |
| H | 4.30067300 | 1.21455500 | -0.57982700 |
| H | 6.73483300 | -1.04694800 | -3.53577800 |
| P | 7.09531000 | 2.91240300 | -2.69084000 |
| O | 6.27796600 | 3.66631800 | -1.63769100 |
| O | 7.80089800 | 3.54224500 | -3.83540000 |
| O | 8.13345400 | 1.96368000 | -1.80849500 |
| C | -4.87997200 | -0.62314800 | -5.68956800 |
| C | -4.66455300 | -1.62662200 | -4.55634000 |
| C | -3.68541700 | -1.13668400 | -3.49557700 |
| C | -2.29631700 | 0.94081900 | -1.64034800 |
| S | -4.29552200 | 0.27322800 | -2.47777800 |
| C | -4.91533400 | -0.58460900 | -0.97387900 |
| C | -6.41092200 | -0.48326700 | -0.76747400 |
| O | -6.76772500 | 0.84826700 | -0.43972100 |
| C | -7.31176600 | -0.82424100 | -1.96438600 |
| O | -7.39995000 | -2.18209600 | -2.27423900 |
| C | -8.63229600 | -0.18397800 | -1.51443800 |
| O | -9.23881800 | -1.15283800 | -0.66866500 |
| C | -8.14277500 | 1.06316400 | -0.76253900 |
| H | -5.30008800 | 0.32289600 | -5.30994900 |
| H | -5.61968900 | -1.92287900 | -4.09220600 |
| H | -4.23252600 | -2.55664100 | -4.96772300 |
| H | -2.75931300 | -0.77196200 | -3.96161900 |
| H | -3.39340800 | -1.92289700 | -2.78730500 |
| H | -1.74084400 | 0.38381800 | -2.39144400 |
| H | -2.50040400 | 2.00004400 | -1.77398400 |
| H | -2.51147800 | 0.49098000 | -0.67601600 |
| H | -8.73491200 | 1.21629100 | 0.15689900 |
| H | -10.14822400 | -0.89451600 | -0.47662300 |
| H | -6.93086700 | -0.28827600 | -2.84869700 |
| H | -6.67125400 | -1.17395500 | 0.06227000 |
| H | -4.59202900 | -1.63164500 | -1.03381300 |
| H | -8.08465400 | -2.54389900 | -1.68743700 |
| H | -9.29187600 | 0.06467300 | -2.36343200 |
| H | -4.41020800 | -0.11934600 | -0.11427100 |
| H | -0.74713800 | 9.55129300 | 1.84709700 |
| H | -0.27333400 | 8.64098100 | 3.31063600 |
| H | -6.05296100 | 5.46556100 | -0.74120800 |
| H | -4.80114400 | 6.69016200 | -1.03046000 |
| H | -5.51355900 | 1.14774100 | 4.49492900 |
| H | -4.86307100 | -0.40235100 | 3.87191800 |
| H | -8.22019300 | 1.97152700 | -1.38338600 |
| H | -5.57513400 | -1.01598400 | -6.44726200 |
| H | -3.92839900 | -0.38506500 | -6.19220800 |
| H | -5.68600600 | -6.35668300 | 2.71134600 |
| H | -4.66869100 | -5.29713600 | 3.69996300 |
| H | -2.54528600 | -6.06485900 | -0.81719700 |
| H | -2.04546500 | -4.14473400 | -5.55711500 |
| H | -0.30858800 | -4.37818000 | -5.93447400 |
| H | 2.79562600 | -7.75427000 | 1.53127900 |
| H | 3.13224100 | -6.82489300 | 6.31077800 |
| H | 3.89638900 | -5.53102800 | 7.25300800 |
| H | 5.34916500 | -2.01679200 | -5.14425200 |
| H | 8.92263400 | 1.82474200 | -2.35089100 |
| H | 6.52413700 | 0.97811400 | 2.43118600 |

#### Model3' – Final state

| | | | |
|---|---|---|---|
| O | -0.60449400 | 4.64199800 | -0.77157000 |
| H | -0.09437700 | 5.11232300 | 0.64512800 |
| H | -1.60298300 | 4.61497100 | -0.88158500 |
| O | 0.06057300 | 3.84636300 | 4.60361900 |
| H | -0.18824600 | 3.96637200 | 5.53005500 |
| H | 0.34124200 | 4.74590400 | 4.29567100 |
| O | 0.71171200 | -1.26482600 | 3.15867500 |
| H | 1.00312700 | -2.12179200 | 3.55141300 |
| H | 0.86841700 | -0.64098400 | 3.87965300 |
| C | -0.40601600 | 8.68867400 | 2.25660500 |
| C | -0.69625300 | 7.32055700 | 1.63909300 |
| C | 0.06162400 | 6.19197100 | 2.30423700 |

| | | | |
|---|---|---|---|
| O | 0.30210000 | 5.10937900 | 1.57354300 |
| O | 0.42140100 | 6.23660500 | 3.46702900 |
| H | 0.64469500 | 8.97756000 | 2.10234800 |
| H | -0.50265800 | 7.30755200 | 0.55462400 |
| H | -1.76058100 | 7.04566200 | 1.76381500 |
| C | 6.17545400 | 1.38677700 | 1.74275300 |
| C | 5.30587900 | 2.64013100 | 1.70194500 |
| N | 3.98845500 | 2.41219500 | 2.31006400 |
| C | 2.78767900 | 2.78947100 | 1.85455900 |
| N | 2.60147100 | 3.28284900 | 0.63886300 |
| N | 1.71064500 | 2.62745400 | 2.66539500 |
| H | 5.65734000 | 0.53431200 | 1.27731100 |
| H | 7.10310700 | 1.57810200 | 1.18263100 |
| H | 5.19013600 | 2.98972200 | 0.67089400 |
| H | 5.80386400 | 3.46444200 | 2.24050000 |
| H | 3.98410800 | 1.91651200 | 3.19458300 |
| H | 3.26447000 | 3.19847400 | -0.18795000 |
| H | 1.67629000 | 3.63637400 | 0.42746500 |
| H | 1.82157700 | 2.70524000 | 3.67213100 |
| H | 0.86365300 | 3.10350600 | 2.36762000 |
| C | 2.47478500 | -6.84342300 | 1.00033100 |
| C | 3.56034100 | -6.14307000 | 0.18231400 |
| C | 4.66883600 | -5.52649500 | 1.05062400 |
| N | 4.20403600 | -4.47491500 | 1.93298200 |
| C | 4.03227800 | -3.20253700 | 1.57366200 |
| N | 4.41013200 | -2.74945600 | 0.35093600 |
| N | 3.52215300 | -2.33945100 | 2.45077300 |
| H | 2.04523200 | -6.16745500 | 1.75607600 |
| H | 1.65914700 | -7.21337900 | 0.35830400 |
| H | 3.11256200 | -5.34721800 | -0.43928700 |
| H | 4.02743100 | -6.85464100 | -0.52070600 |
| H | 5.49092500 | -5.15132800 | 0.41651900 |
| H | 5.11152900 | -6.30888600 | 1.68708800 |
| H | 3.81468800 | -4.75144400 | 2.88406100 |
| H | 4.64810700 | -3.40076600 | -0.38250600 |
| H | 4.09476600 | -1.82190800 | 0.06533700 |
| H | 3.02679000 | -2.72019700 | 3.29326300 |
| H | 3.18278400 | -1.45455600 | 2.07170100 |
| C | 2.61551800 | -6.02178200 | 6.96812800 |
| C | 1.81674300 | -4.92024900 | 6.26088600 |
| C | 2.37828100 | -4.44230900 | 4.91126400 |
| O | 3.15005900 | -5.20898900 | 4.28668300 |
| O | 2.01088900 | -3.29269500 | 4.50991700 |
| H | 2.10743300 | -6.34944600 | 7.88882600 |
| H | 1.70072100 | -4.02952900 | 6.89919800 |
| H | 0.78787500 | -5.27008200 | 6.05627900 |
| C | -5.02440900 | -5.64378400 | 2.18088400 |
| C | -3.75539500 | -5.95263500 | 1.41063800 |
| O | -3.00967900 | -6.86686900 | 1.72190200 |
| H | -5.61708300 | -4.82802200 | 1.74034200 |
| N | -3.49541300 | -5.13873200 | 0.34206300 |
| C | -2.30224000 | -5.31826400 | -0.43636000 |
| C | -1.78911900 | -3.99425200 | -0.98430500 |
| O | -2.51907300 | -3.03047700 | -1.16293000 |
| H | -4.10139300 | -4.36180100 | 0.09871200 |
| H | -1.55435800 | -5.82703000 | 0.19182800 |
| N | -0.47165800 | -3.99974700 | -1.33646400 |
| C | 0.18538700 | -2.82461200 | -1.89405500 |
| C | -0.17588300 | -2.57460200 | -3.37902400 |
| O | -0.20334000 | -1.43493100 | -3.82326800 |
| C | 1.69698900 | -2.96482500 | -1.74384700 |
| H | 0.11333000 | -4.74806800 | -0.97987700 |
| H | -0.17672600 | -1.93239300 | -1.36685300 |
| H | 2.04727300 | -3.89387900 | -2.22394100 |
| H | 1.97106300 | -2.99273300 | -0.68004700 |
| H | 2.22978100 | -2.12813100 | -2.21245000 |
| N | -0.36746900 | -3.69376600 | -4.12769000 |
| C | -0.57973400 | -3.70414900 | -5.56792000 |
| H | -0.37106700 | -4.57546400 | -3.62841100 |
| H | -0.53941100 | -2.66399500 | -5.91702600 |
| C | -5.24149900 | 5.85831200 | -0.62836300 |
| C | -4.09272700 | 4.93348600 | -0.28706600 |
| O | -3.19426700 | 4.68128800 | -1.10690800 |
| H | -5.73046800 | 6.26974100 | 0.26539800 |
| N | -4.16317400 | 4.37272000 | 0.92575600 |
| C | -3.55475500 | 3.11302000 | 1.30147100 |
| C | -4.62534800 | 2.43378200 | 2.17165200 |
| O | -5.58689800 | 3.06168300 | 2.60037100 |
| C | -2.20298400 | 3.23851800 | 2.01068800 |
| O | -2.28044600 | 4.18369200 | 3.04349200 |
| H | -4.96704100 | 4.57483000 | 1.52499000 |
| H | -3.39216000 | 2.50149000 | 0.40089700 |
| H | -1.45545500 | 3.51868600 | 1.24938200 |
| H | -1.91455400 | 2.23691400 | 2.37346200 |
| H | -1.52581600 | 4.02859000 | 3.64548200 |
| N | -4.45460000 | 1.11952700 | 2.40751300 |
| C | -5.46875300 | 0.41405300 | 3.16457200 |
| H | -3.66593000 | 0.62841500 | 1.99403000 |
| H | -6.45155900 | 0.49302200 | 2.67202300 |
| C | 6.04577600 | -1.33697300 | -3.06096300 |
| O | 5.71240600 | -2.29200700 | -4.04192300 |
| C | 5.07737100 | -0.17972800 | -2.97173100 |
| O | 3.82231000 | -0.55642000 | -2.40370100 |
| C | 5.58480100 | 0.99275100 | -2.11804400 |
| O | 6.24551300 | 1.89967600 | -2.95908300 |
| C | 4.28003100 | 1.51313900 | -1.43573800 |
| O | 4.02992500 | 2.88643100 | -1.53546300 |
| C | 3.18122100 | 0.68839500 | -2.12579000 |
| N | -0.47831300 | 0.89352500 | -1.28358700 |
| C | -0.58944800 | 0.44090200 | 0.03044100 |
| O | -1.65038500 | 0.45604400 | 0.63659400 |
| N | 0.59576600 | -0.01831900 | 0.57734000 |
| C | 1.85753900 | 0.03671700 | -0.00662700 |
| O | 2.84950600 | -0.32804600 | 0.63388900 |
| C | 1.89824500 | 0.53139900 | -1.36838000 |
| C | 0.72024700 | 0.90875900 | -1.93428800 |
| H | -0.26592300 | 5.03964500 | -1.58487900 |
| H | 0.55164900 | -0.43601400 | 1.53016700 |
| H | 0.68276800 | 1.25414400 | -2.96699900 |
| H | 2.95207200 | 1.20393500 | -3.07857700 |
| H | 4.96165500 | 3.37666200 | -1.46954700 |
| H | 6.27872000 | 0.62818200 | -1.34167500 |
| H | 4.90823700 | 0.23134400 | -3.98771300 |
| H | 6.07254800 | -1.86136800 | -2.09100300 |
| H | 4.31581800 | 1.18927000 | -0.38490400 |
| H | 7.05109500 | -0.90157500 | -3.22607800 |
| P | 7.21874100 | 3.07836600 | -2.26796700 |
| O | 6.30880900 | 3.76991000 | -1.24462000 |
| O | 7.96237900 | 3.76661500 | -3.35202200 |
| O | 8.23552400 | 2.14320300 | -1.35069400 |
| C | -4.46543600 | -0.63912500 | -6.08381600 |
| C | -4.36757700 | -1.75342300 | -5.04065900 |
| C | -3.81470300 | -1.30292800 | -3.68939100 |
| C | -1.68659700 | 1.36882700 | -1.95509700 |
| S | -4.85916600 | 0.03084600 | -2.95979500 |
| C | -4.78471800 | -0.34516500 | -1.16405500 |
| C | -6.15672000 | -0.16516500 | -0.53848200 |
| O | -6.72449300 | 1.08764800 | -0.92629200 |
| C | -7.18361100 | -1.24550100 | -0.92464500 |
| O | -7.13657700 | -2.38886600 | -0.11223500 |
| C | -8.49874900 | -0.45968300 | -0.82954600 |
| O | -8.81501200 | -0.45178600 | 0.55786200 |
| C | -8.07162700 | 0.90856700 | -1.35462200 |
| H | -5.15781500 | 0.15266200 | -5.75605400 |
| H | -5.36136600 | -2.20944100 | -4.88984100 |
| H | -3.70768000 | -2.55533100 | -5.41898700 |
| H | -2.78094100 | -0.93820700 | -3.80057000 |
| H | -3.78369700 | -2.13844800 | -2.97848200 |
| H | -1.45426400 | 1.52545300 | -3.01345500 |
| H | -2.03686500 | 2.31186200 | -1.51424500 |

| | | | |
|---|---|---|---|
| H | -2.47839400 | 0.62074700 | -1.88692600 |
| H | -8.69443600 | 1.72453800 | -0.95071600 |
| H | -9.63740200 | 0.03134400 | 0.70198900 |
| H | -7.01144100 | -1.55375000 | -1.96774400 |
| H | -6.04519300 | -0.18994400 | 0.55988900 |
| H | -4.43437200 | -1.37678900 | -1.01340200 |
| H | -7.63207600 | -2.15512300 | 0.68964000 |
| H | -9.31445200 | -0.91128100 | -1.42232500 |
| H | -4.05762500 | 0.31400800 | -0.66976800 |
| H | -1.04547600 | 9.46061400 | 1.80415100 |
| H | -0.59194700 | 8.66643200 | 3.33936300 |
| H | -5.98460800 | 5.27259200 | -1.19397800 |
| H | -4.88883800 | 6.67195000 | -1.27614200 |
| H | -5.56768800 | 0.83390100 | 4.17746300 |
| H | -5.18961200 | -0.64528200 | 3.23989300 |
| H | -8.13159400 | 0.93174400 | -2.45926900 |
| H | -4.83138000 | -1.02433500 | -7.04853500 |
| H | -3.48177200 | -0.17150500 | -6.25699200 |
| H | -5.63647200 | -6.55636300 | 2.22749600 |
| H | -4.75182200 | -5.38206400 | 3.21457100 |
| H | -2.47146000 | -5.98181900 | -1.30974100 |
| H | -1.56435200 | -4.12711100 | -5.82604600 |
| H | 0.20407100 | -4.28591400 | -6.07906200 |
| H | 2.88684300 | -7.70754300 | 1.54563800 |
| H | 2.74184200 | -6.88917300 | 6.30387800 |
| H | 3.62352800 | -5.66910100 | 7.23856900 |
| H | 5.71636100 | -1.84707100 | -4.89946000 |
| H | 9.06431300 | 2.05146100 | -1.84183100 |
| H | 6.43729400 | 1.10410800 | 2.77592000 |

#### Model3” – Initial state

| | | | |
|---|---|---|---|
| O | -1.73997200 | 3.67455400 | -1.04675600 |
| H | -1.41056000 | 4.39922200 | -0.45551600 |
| H | -2.71253800 | 3.70563300 | -1.08066800 |
| O | -2.03214200 | 5.69011500 | 4.74239400 |
| H | -2.37596800 | 6.56871400 | 4.94829100 |
| H | -1.28822600 | 5.85059800 | 4.09991200 |
| O | 0.84257900 | -1.67683100 | 3.07971000 |
| H | 1.35362800 | -2.43981900 | 3.44647800 |
| H | 0.74063900 | -1.10320400 | 3.85078900 |
| C | -1.57370900 | 8.47522700 | 2.22804200 |
| C | -1.89471800 | 7.10608500 | 1.59457900 |
| C | -0.78241400 | 6.04949200 | 1.70215500 |
| O | -0.51040700 | 5.33049600 | 0.70593000 |
| O | -0.21895500 | 5.92452400 | 2.83103000 |
| H | -0.72338200 | 8.95728200 | 1.72128500 |
| H | -2.16627200 | 7.21539200 | 0.53412800 |
| H | -2.76504000 | 6.65754900 | 2.10887300 |
| C | 5.93368300 | 2.12796800 | 1.73403500 |
| C | 5.04930000 | 3.36773100 | 1.60392800 |
| N | 3.80075600 | 3.24694200 | 2.37015200 |
| C | 2.59638000 | 3.79728900 | 2.08013400 |
| N | 2.21393000 | 4.07650200 | 0.84220400 |
| N | 1.73746500 | 4.01755100 | 3.09583800 |
| H | 5.39723900 | 1.21999900 | 1.41610800 |
| H | 6.81880600 | 2.26081700 | 1.09360100 |
| H | 4.82213000 | 3.57610300 | 0.55563900 |
| H | 5.60236100 | 4.25699500 | 1.95422000 |
| H | 3.92964600 | 3.02451200 | 3.35112600 |
| H | 2.76419400 | 3.81311100 | -0.00254800 |
| H | 1.28694800 | 4.51173700 | 0.71782600 |
| H | 2.13031500 | 4.05936000 | 4.02843800 |
| H | 0.92572900 | 4.66367800 | 2.93942200 |
| C | 3.37893800 | -6.53197900 | 1.05711800 |
| C | 4.49345600 | -5.90336700 | 0.22180000 |
| C | 5.56123500 | -5.19905400 | 1.07460100 |
| N | 5.03210200 | -4.16688700 | 1.94286100 |
| C | 4.63581500 | -2.96123000 | 1.52710100 |
| N | 4.90658300 | -2.51707600 | 0.27630500 |
| N | 3.99483900 | -2.15924500 | 2.37584100 |
| H | 2.85314700 | -5.77195700 | 1.65576600 |
| H | 2.64065200 | -7.04542700 | 0.41996600 |
| H | 4.06151600 | -5.17848700 | -0.49158900 |
| H | 4.99646900 | -6.67242800 | -0.38982500 |
| H | 6.35859900 | -4.78732500 | 0.43024200 |
| H | 6.05223800 | -5.93807300 | 1.72641100 |
| H | 4.69035600 | -4.46563200 | 2.90160300 |
| H | 5.33099200 | -3.12646400 | -0.40625600 |
| H | 4.42094900 | -1.68070200 | -0.05298500 |
| H | 3.51920300 | -2.60901300 | 3.19466300 |
| H | 3.51287800 | -1.36578100 | 1.95126700 |
| C | 3.42516400 | -5.65933300 | 7.01933400 |
| C | 2.43525600 | -4.97427400 | 6.07624200 |
| C | 3.06068900 | -4.41945700 | 4.79007700 |
| O | 4.04714400 | -5.01891700 | 4.30373000 |
| O | 2.52437800 | -3.38109500 | 4.28943100 |
| H | 2.91012800 | -6.10830800 | 7.88347200 |
| H | 1.90044800 | -4.14990900 | 6.57291200 |
| H | 1.65895800 | -5.69486400 | 5.75827800 |
| C | -4.21014800 | -6.35039600 | 2.25952400 |
| C | -2.99672900 | -6.51459100 | 1.36668400 |
| O | -2.21735100 | -7.44426800 | 1.50002300 |
| H | -4.88711500 | -5.54591700 | 1.93523800 |
| N | -2.80808800 | -5.55288200 | 0.41214300 |
| C | -1.65631400 | -5.63513100 | -0.44658400 |
| C | -1.29575800 | -4.28201800 | -1.03544100 |
| O | -2.14326000 | -3.46249000 | -1.36815900 |
| H | -3.46315200 | -4.79087200 | 0.26034600 |
| H | -0.82592800 | -6.07828000 | 0.12349000 |
| N | 0.03495600 | -4.09600200 | -1.24484200 |
| C | 0.55815700 | -2.87974200 | -1.85247600 |
| C | 0.03488200 | -2.72113000 | -3.30528700 |
| O | -0.45120700 | -1.66723900 | -3.70037400 |
| C | 2.08320300 | -2.89361800 | -1.81002300 |
| H | 0.68887700 | -4.71830000 | -0.78226500 |
| H | 0.16762800 | -2.00076300 | -1.31906700 |
| H | 2.48496500 | -3.77670300 | -2.33393700 |
| H | 2.43020400 | -2.91266100 | -0.76764500 |
| H | 2.50837800 | -1.99538500 | -2.27887700 |
| N | 0.20720100 | -3.81564100 | -4.09500600 |
| C | -0.09193100 | -3.87884100 | -5.51863600 |
| H | 0.57667300 | -4.64629400 | -3.64726700 |
| H | -0.36882200 | -2.86868900 | -5.84795500 |
| C | -5.99084100 | 4.99757600 | -0.62021300 |
| C | -4.96587100 | 3.93407800 | -0.30024600 |
| O | -4.50664700 | 3.19046800 | -1.18393900 |
| H | -6.00634200 | 5.80708000 | 0.12288300 |
| N | -4.58140800 | 3.84823500 | 0.98116200 |
| C | -3.91217100 | 2.70478800 | 1.55910400 |
| C | -5.00062100 | 1.69550700 | 2.00653300 |
| O | -6.18134600 | 1.88500700 | 1.74192300 |
| C | -2.99519100 | 3.21243900 | 2.67573800 |
| O | -3.75572100 | 4.05233100 | 3.50038300 |
| H | -4.99928300 | 4.45481200 | 1.68075600 |
| H | -3.28726500 | 2.21110100 | 0.80434700 |
| H | -2.15674700 | 3.74286700 | 2.18892200 |
| H | -2.55883900 | 2.35237300 | 3.21818400 |
| H | -3.14805000 | 4.63448500 | 4.02204500 |
| N | -4.55550800 | 0.59015200 | 2.65698100 |
| C | -5.46730100 | -0.40212700 | 3.20726700 |
| H | -3.55238800 | 0.44444900 | 2.72886700 |
| H | -6.49070900 | -0.05872500 | 3.01404700 |
| C | 6.09299100 | -0.62519100 | -3.02864300 |
| O | 5.95563000 | -1.61868100 | -4.02610700 |
| C | 4.98056800 | 0.40658100 | -2.96047200 |
| O | 3.76953200 | -0.10731800 | -2.39987200 |
| C | 5.36169800 | 1.63248400 | -2.10279300 |
| O | 5.89577900 | 2.61250500 | -2.95133100 |
| C | 4.02248600 | 1.98843100 | -1.38099500 |
| O | 3.58526700 | 3.31171700 | -1.45409900 |

| | | | |
|---|---|---|---|
| C | 3.01073900 | 1.04967700 | -2.05742800 |
| N | -0.40028100 | 1.45645600 | -0.55065400 |
| C | -0.45658200 | 0.61885900 | 0.54453800 |
| O | -1.44610000 | 0.53617700 | 1.26175800 |
| N | 0.70389000 | -0.10975200 | 0.74907000 |
| C | 1.88138300 | 0.00327000 | 0.01326800 |
| O | 2.91153100 | -0.54111700 | 0.42119600 |
| C | 1.80491800 | 0.78497900 | -1.20796200 |
| C | 0.67525100 | 1.52534500 | -1.38186800 |
| H | -1.10127600 | 2.23711400 | -0.64193200 |
| H | 0.74045200 | -0.72202600 | 1.59455800 |
| H | 0.58081400 | 2.25236200 | -2.19160500 |
| H | 2.67872000 | 1.56249700 | -2.98094900 |
| H | 4.42352800 | 3.92818000 | -1.46245300 |
| H | 6.11080600 | 1.34714400 | -1.34463400 |
| H | 4.77772200 | 0.79709100 | -3.97827700 |
| H | 6.15291200 | -1.14753800 | -2.05938700 |
| H | 4.12123400 | 1.65535500 | -0.33353900 |
| H | 7.03493400 | -0.05564900 | -3.15118700 |
| P | 6.72640300 | 3.91879000 | -2.29102000 |
| O | 5.73461900 | 4.55379700 | -1.31668200 |
| O | 7.42291500 | 4.62749600 | -3.39460700 |
| O | 7.82179600 | 3.12315300 | -1.32679500 |
| C | -4.35942100 | -1.37055800 | -6.03884900 |
| C | -4.55744700 | -2.11964100 | -4.71360900 |
| C | -3.71800100 | -1.60984600 | -3.54177300 |
| C | -2.64432300 | 0.98058500 | -3.11639200 |
| S | -4.22785800 | 0.09859900 | -3.04042300 |
| C | -4.49022100 | 0.07314600 | -1.22411600 |
| C | -5.88670700 | -0.33822900 | -0.80518300 |
| O | -6.86339000 | 0.30456700 | -1.60302400 |
| C | -6.29193300 | -1.82015200 | -0.88526000 |
| O | -5.68226700 | -2.65801500 | 0.05376800 |
| C | -7.82319500 | -1.68743000 | -0.70141600 |
| O | -8.02149800 | -1.74778600 | 0.70287700 |
| C | -8.12658400 | -0.29445400 | -1.30468200 |
| H | -4.65578600 | -0.31188500 | -5.95177000 |
| H | -5.62574500 | -2.12040700 | -4.43467300 |
| H | -4.28024000 | -3.17997400 | -4.84279300 |
| H | -2.64585900 | -1.54233700 | -3.78622700 |
| H | -3.79355200 | -2.24082800 | -2.64550800 |
| H | -2.38190700 | 1.05724700 | -4.17909300 |
| H | -2.80405300 | 1.97501500 | -2.68148000 |
| H | -1.87559600 | 0.41072200 | -2.58001900 |
| H | -8.67671300 | 0.31303100 | -0.56434300 |
| H | -8.96390500 | -1.77484100 | 0.90719300 |
| H | -6.07877200 | -2.20724000 | -1.89565000 |
| H | -5.98033100 | -0.02408600 | 0.25127700 |
| H | -3.71738900 | -0.54869500 | -0.74891700 |
| H | -6.24922600 | -2.61642300 | 0.84359900 |
| H | -8.37526400 | -2.49801400 | -1.20727000 |
| H | -4.34306600 | 1.13850100 | -0.98151300 |
| H | -2.44102800 | 9.14977400 | 2.16020100 |
| H | -1.30291100 | 8.35729700 | 3.28790300 |
| H | -6.98042000 | 4.51283600 | -0.63013200 |
| H | -5.79733700 | 5.40410600 | -1.62143200 |
| H | -5.31570500 | -0.52321700 | 4.29174100 |
| H | -5.32533100 | -1.38019000 | 2.71758900 |
| H | -8.70781600 | -0.34541800 | -2.23776700 |
| H | -4.96957300 | -1.81882400 | -6.83660100 |
| H | -3.30441200 | -1.40157700 | -6.35541500 |
| H | -4.75632500 | -7.30392400 | 2.28835100 |
| H | -3.86537400 | -6.13960600 | 3.28356700 |
| H | -1.83007500 | -6.31714100 | -1.30488900 |
| H | -0.92962200 | -4.56501700 | -5.72884500 |
| H | 0.79015400 | -4.21068300 | -6.08816800 |
| H | 3.77987900 | -7.27036800 | 1.76947200 |
| H | 3.97411400 | -6.44737700 | 6.48361600 |
| H | 4.16999800 | -4.94268700 | 7.40131000 |
| H | 5.94615800 | -1.17069500 | -4.88218100 |
| H | 8.66094500 | 3.10282100 | -1.80816700 |
| H | 6.27296900 | 1.97611300 | 2.77267700 |

#### Model3" – TS1

| | | | |
|---|---|---|---|
| O | 1.29514700 | -3.99600900 | -0.81590000 |
| H | 0.70045700 | -4.71284500 | 0.05988400 |
| H | 2.27377500 | -3.98263300 | -0.93180100 |
| O | 1.17780900 | -5.72985800 | 4.71950900 |
| H | 1.46764500 | -6.53408100 | 5.16949200 |
| H | 0.45931500 | -6.01648200 | 4.11662300 |
| O | -0.51750000 | 1.63675200 | 3.05416900 |
| H | -0.90828400 | 2.45420100 | 3.43792200 |
| H | -0.66768600 | 0.99826800 | 3.76444100 |
| C | 1.00293400 | -8.52163000 | 2.06642000 |
| C | 1.26326400 | -7.14399200 | 1.42100000 |
| C | 0.15622500 | -6.12783600 | 1.66729500 |
| O | -0.01396800 | -5.13406700 | 0.85392900 |
| O | -0.56159700 | -6.24302200 | 2.67203800 |
| H | 0.08057800 | -8.97579500 | 1.67568000 |
| H | 1.43410500 | -7.22862000 | 0.33604200 |
| H | 2.17407500 | -6.68078800 | 1.84695400 |
| C | -6.05023700 | -1.66736100 | 1.66429900 |
| C | -5.65198700 | -3.12059000 | 1.43904500 |
| N | -4.62607300 | -3.57899700 | 2.37536400 |
| C | -3.36098700 | -3.95122800 | 2.12055800 |
| N | -2.80814600 | -3.88073800 | 0.91899100 |
| N | -2.61636600 | -4.39406300 | 3.16141700 |
| H | -5.19707900 | -0.98397200 | 1.52976200 |
| H | -6.83589800 | -1.40507600 | 0.94009200 |
| H | -5.31839700 | -3.30418200 | 0.41364900 |
| H | -6.53089600 | -3.77486100 | 1.55158100 |
| H | -4.89970500 | -3.60995100 | 3.35031900 |
| H | -3.26858500 | -3.51995900 | 0.04655000 |
| H | -1.85045900 | -4.22423000 | 0.82759100 |
| H | -3.10837100 | -4.64590100 | 4.00948200 |
| H | -1.78940700 | -4.97916100 | 2.95778500 |
| C | -2.90603900 | 6.80806000 | 1.15336200 |
| C | -3.99892900 | 6.13756200 | 0.32057700 |
| C | -5.06130300 | 5.42859700 | 1.17708600 |
| N | -4.52362200 | 4.38738600 | 2.02763400 |
| C | -4.16956500 | 3.17165100 | 1.60034000 |
| N | -4.50721300 | 2.72443400 | 0.36647100 |
| N | -3.50747400 | 2.35905600 | 2.42076800 |
| H | -2.40638500 | 6.08224600 | 1.81345600 |
| H | -2.14440400 | 7.28095400 | 0.51230200 |
| H | -3.54684800 | 5.40300500 | -0.36980900 |
| H | -4.50821700 | 6.88272400 | -0.31532200 |
| H | -5.86412000 | 5.02347300 | 0.53563600 |
| H | -5.54618900 | 6.16335500 | 1.83884300 |
| H | -4.14550100 | 4.67536600 | 2.97728800 |
| H | -4.89341400 | 3.35888500 | -0.31613100 |
| H | -4.04367800 | 1.87472100 | 0.03007600 |
| H | -3.02743700 | 2.78198600 | 3.24851800 |
| H | -3.04752700 | 1.55962600 | 1.97687500 |
| C | -3.03820100 | 5.83366000 | 7.09847100 |
| C | -2.04772800 | 5.00517000 | 6.27690700 |
| C | -2.57757100 | 4.51690300 | 4.91954500 |
| O | -3.49528100 | 5.17846200 | 4.37637000 |
| O | -2.04005800 | 3.47582600 | 4.43257900 |
| H | -2.57302700 | 6.21241600 | 8.02272900 |
| H | -1.69427400 | 4.12150100 | 6.83161000 |
| H | -1.14287400 | 5.60258200 | 6.05929400 |
| C | 4.64743300 | 6.08556300 | 2.37636800 |
| C | 3.44424600 | 6.36132900 | 1.49385800 |
| O | 2.73918800 | 7.34419500 | 1.65660700 |
| H | 5.27289900 | 5.25401900 | 2.01796800 |
| N | 3.17339400 | 5.43960700 | 0.52065800 |
| C | 2.01951000 | 5.62515500 | -0.32117500 |
| C | 1.59857200 | 4.32228600 | -0.98426800 |
| O | 2.41246400 | 3.49175800 | -1.36788800 |

| | | | |
|---|---|---|---|
| H | 3.77102800 | 4.63943900 | 0.33262400 |
| H | 1.21088600 | 6.06802400 | 0.27981500 |
| N | 0.25955900 | 4.20254200 | -1.18804200 |
| C | -0.32894600 | 3.02403500 | -1.81336700 |
| C | 0.16527600 | 2.85888000 | -3.27525200 |
| O | 0.52554900 | 1.77236100 | -3.70944100 |
| C | -1.84896400 | 3.11343500 | -1.75763700 |
| H | -0.35972500 | 4.82686700 | -0.68235000 |
| H | 0.01908500 | 2.11589100 | -1.29997200 |
| H | -2.20974900 | 4.02662800 | -2.25953100 |
| H | -2.18204300 | 3.12424600 | -0.71145400 |
| H | -2.31874000 | 2.24544100 | -2.24008400 |
| N | 0.10407300 | 3.98780000 | -4.03569200 |
| C | 0.40422700 | 4.04196600 | -5.45857300 |
| H | -0.16341400 | 4.84393000 | -3.56422200 |
| H | 0.54681500 | 3.01091700 | -5.80805800 |
| C | 5.66008300 | -5.30407800 | -0.70396100 |
| C | 4.62580300 | -4.28026100 | -0.31015900 |
| O | 3.95806600 | -3.68061600 | -1.17643900 |
| H | 6.06237000 | -5.86366000 | 0.15104000 |
| N | 4.48473600 | -4.05756000 | 0.99699900 |
| C | 3.77195300 | -2.94413200 | 1.59303800 |
| C | 4.84145000 | -2.10989300 | 2.32098100 |
| O | 5.91381100 | -2.59673100 | 2.65352300 |
| C | 2.65493600 | -3.45655500 | 2.51306300 |
| O | 3.16698900 | -4.49980900 | 3.29786800 |
| H | 5.10281100 | -4.52745800 | 1.65214900 |
| H | 3.29688800 | -2.33603600 | 0.81259100 |
| H | 1.82850400 | -3.78933500 | 1.86145000 |
| H | 2.27529000 | -2.60586000 | 3.11052400 |
| H | 2.45239100 | -4.87825300 | 3.85668500 |
| N | 4.53384400 | -0.80723300 | 2.53164300 |
| C | 5.49042800 | 0.04907800 | 3.21954800 |
| H | 3.63709800 | -0.44802200 | 2.20500400 |
| H | 6.41880700 | 0.15668600 | 2.63465600 |
| C | -5.95739300 | 1.19578400 | -3.01522600 |
| O | -5.79037800 | 2.17482700 | -4.02666200 |
| C | -4.93019700 | 0.07151300 | -2.97307300 |
| O | -3.67099200 | 0.46287300 | -2.42646900 |
| C | -5.40675100 | -1.13308100 | -2.12559200 |
| O | -6.01867800 | -2.05696000 | -2.99014000 |
| C | -4.09713400 | -1.59573100 | -1.40898500 |
| O | -3.79047900 | -2.96156700 | -1.44308100 |
| C | -3.00732700 | -0.76970600 | -2.11473200 |
| N | 0.52663500 | -1.46093200 | -0.96436400 |
| C | 0.68874600 | -0.66587700 | 0.13839600 |
| O | 1.75124400 | -0.61228400 | 0.77825300 |
| N | -0.41793200 | 0.09113100 | 0.52973300 |
| C | -1.66447000 | 0.08393000 | -0.09062900 |
| O | -2.62487800 | 0.66309200 | 0.44636700 |
| C | -1.73031700 | -0.62400900 | -1.34207000 |
| C | -0.62812300 | -1.38586900 | -1.66159400 |
| H | 1.02904300 | -3.00211600 | -0.84797500 |
| H | -0.34568000 | 0.65366300 | 1.39459200 |
| H | -0.67927900 | -2.03341400 | -2.54555200 |
| H | -2.77520300 | -1.30441700 | -3.05669700 |
| H | -4.65668900 | -3.51812200 | -1.57264300 |
| H | -6.13408800 | -0.80194900 | -1.36567400 |
| H | -4.78052100 | -0.32356200 | -3.99867800 |
| H | -5.94040400 | 1.72616400 | -2.04876700 |
| H | -4.15140300 | -1.23283200 | -0.36867800 |
| H | -6.94551500 | 0.70224100 | -3.09946000 |
| P | -6.94587000 | -3.32703000 | -2.39526600 |
| O | -5.97903200 | -4.14913500 | -1.54450400 |
| O | -7.77228400 | -3.86196300 | -3.50736600 |
| O | -7.90293400 | -2.51994600 | -1.29798900 |
| C | 4.49219200 | 1.25706500 | -6.01076800 |
| C | 4.49349600 | 1.95424800 | -4.63999700 |
| C | 3.67811300 | 1.28989700 | -3.52621400 |
| C | 2.90047200 | -1.46451700 | -3.31618500 |
| S | 4.34493500 | -0.38570000 | -3.10736000 |
| C | 4.50827000 | -0.46157800 | -1.27951200 |
| C | 5.88429600 | -0.08775700 | -0.76848900 |
| O | 6.89324600 | -0.82265200 | -1.44284300 |
| C | 6.34264500 | 1.37648300 | -0.89035200 |
| O | 5.71799300 | 2.26363300 | -0.01283000 |
| C | 7.85756600 | 1.20853500 | -0.64611900 |
| O | 7.99119000 | 1.20868300 | 0.76751300 |
| C | 8.15605700 | -0.15945400 | -1.28511900 |
| H | 4.93899800 | 0.25047100 | -5.95326700 |
| H | 5.53309800 | 2.09957000 | -4.29679700 |
| H | 4.07005700 | 2.96821100 | -4.74055100 |
| H | 2.61807200 | 1.14983100 | -3.79332900 |
| H | 3.68392400 | 1.88136700 | -2.59850200 |
| H | 2.58142800 | -1.36883300 | -4.36148700 |
| H | 3.24856000 | -2.48113100 | -3.09004600 |
| H | 2.10444700 | -1.17216900 | -2.61178700 |
| H | 8.82261900 | -0.75207400 | -0.63476400 |
| H | 8.92308400 | 1.18846700 | 1.01707200 |
| H | 6.17824200 | 1.73314800 | -1.92130100 |
| H | 5.85947600 | -0.33625400 | 0.31020400 |
| H | 3.71046200 | 0.13298300 | -0.80972100 |
| H | 6.17653600 | 2.16050400 | 0.83782100 |
| H | 8.44876000 | 2.02483200 | -1.09554500 |
| H | 4.30873100 | -1.52638600 | -1.08374500 |
| H | 1.84270900 | -9.20263700 | 1.86448500 |
| H | 0.89182700 | -8.41655900 | 3.15466000 |
| H | 6.48515700 | -4.77411200 | -1.20625200 |
| H | 5.22287800 | -5.99840800 | -1.43571200 |
| H | 5.75994500 | -0.37457700 | 4.19894200 |
| H | 5.03913300 | 1.03970900 | 3.36743700 |
| H | 8.62378600 | -0.06152500 | -2.27845900 |
| H | 5.07577700 | 1.83753400 | -6.74020800 |
| H | 3.46653100 | 1.15130900 | -6.39956300 |
| H | 5.25168000 | 7.00129300 | 2.44190500 |
| H | 4.29015400 | 5.85412000 | 3.39170200 |
| H | 2.21976900 | 6.35007200 | -1.13700200 |
| H | 1.32384900 | 4.61732000 | -5.66194000 |
| H | -0.42844600 | 4.49729600 | -6.01731700 |
| H | -3.32620200 | 7.58930300 | 1.80679900 |
| H | -3.40188900 | 6.68660100 | 6.50726300 |
| H | -3.91870800 | 5.23391600 | 7.37968400 |
| H | -5.82043200 | 1.71776800 | -4.87758100 |
| H | -8.75562900 | -2.36239800 | -1.72750600 |
| H | -6.45321700 | -1.51146800 | 2.67932100 |

#### Model3" − Intermediate

| | | | |
|---|---|---|---|
| O | 1.20077700 | -4.03101100 | -0.77392700 |
| H | 0.45698600 | -4.81447700 | 0.17805000 |
| H | 2.16820700 | -4.04296100 | -0.93882900 |
| O | 0.93480200 | -5.75028500 | 4.69133500 |
| H | 1.21685700 | -6.53871300 | 5.17330200 |
| H | 0.19054300 | -6.04488000 | 4.12977800 |
| O | -0.47353400 | 1.62465000 | 3.08132200 |
| H | -0.84004500 | 2.45256100 | 3.46633100 |
| H | -0.65939800 | 0.98600100 | 3.78289800 |
| C | 0.76803900 | -8.54662600 | 2.04995500 |
| C | 1.02129200 | -7.15980900 | 1.42313700 |
| C | -0.09741800 | -6.16606800 | 1.68099100 |
| O | -0.25418600 | -5.13099700 | 0.89825100 |
| O | -0.84899200 | -6.31079700 | 2.64834000 |
| H | -0.15024800 | -9.00174900 | 1.65133200 |
| H | 1.19625800 | -7.22692100 | 0.33744600 |
| H | 1.92566900 | -6.69038900 | 1.85632900 |
| C | -6.08706600 | -1.49496100 | 1.63394900 |
| C | -5.86822500 | -2.97780400 | 1.35986300 |
| N | -4.93392600 | -3.60144400 | 2.29739100 |
| C | -3.66054600 | -3.96918700 | 2.08480200 |
| N | -3.04133300 | -3.81348200 | 0.92495400 |
| N | -2.98004500 | -4.50337500 | 3.12766900 |

| | | | | | | | |
|---|---|---|---|---|---|---|---|
| H | -5.15464300 | -0.91680000 | 1.53751000 | H | 3.24975900 | -2.40476700 | 0.79930700 |
| H | -6.81938600 | -1.11022000 | 0.90914300 | H | 1.70539700 | -3.79679100 | 1.79881000 |
| H | -5.53692900 | -3.17353400 | 0.33499500 | H | 2.17659400 | -2.65591500 | 3.08013600 |
| H | -6.82270200 | -3.52108100 | 1.43156300 | H | 2.24101900 | -4.92670900 | 3.81070600 |
| H | -5.26040700 | -3.69756600 | 3.25152100 | N | 4.51293800 | -0.92899400 | 2.54185300 |
| H | -3.45367400 | -3.41752900 | 0.04052700 | C | 5.49021800 | -0.10884400 | 3.24565800 |
| H | -2.08431500 | -4.15893800 | 0.85869900 | H | 3.64175800 | -0.53292700 | 2.18871900 |
| H | -3.51510700 | -4.79720100 | 3.93480200 | H | 6.42168000 | -0.01125000 | 2.66336800 |
| H | -2.14442000 | -5.07344100 | 2.93296500 | C | -5.89333700 | 1.37926200 | -3.03523500 |
| C | -2.70266900 | 6.88954900 | 1.15932500 | O | -5.69284800 | 2.35244300 | -4.04571700 |
| C | -3.79698200 | 6.23385700 | 0.31660700 | C | -4.89806100 | 0.22708700 | -2.98796000 |
| C | -4.88373600 | 5.55139400 | 1.16396300 | O | -3.63225400 | 0.58255200 | -2.43471200 |
| N | -4.37876800 | 4.49836100 | 2.01988300 | C | -5.41228900 | -0.96338900 | -2.14357900 |
| C | -4.06009000 | 3.27056900 | 1.59833300 | O | -6.05186100 | -1.86564700 | -3.01188700 |
| N | -4.40459100 | 2.82964200 | 0.36390400 | C | -4.11851400 | -1.46526400 | -1.42424000 |
| N | -3.42856200 | 2.44032700 | 2.42458800 | O | -3.85357900 | -2.84077700 | -1.45756700 |
| H | -2.23211800 | 6.16133100 | 1.83785600 | C | -3.00122400 | -0.66904400 | -2.12353800 |
| H | -1.91902800 | 7.33577300 | 0.52569500 | N | 0.54651300 | -1.38627300 | -1.02178200 |
| H | -3.35046400 | 5.48525100 | -0.36211500 | C | 0.70738000 | -0.63458900 | 0.10995300 |
| H | -4.28312400 | 6.98395200 | -0.33154600 | O | 1.76838900 | -0.60680600 | 0.75554200 |
| H | -5.68979700 | 5.16430100 | 0.51561800 | N | -0.39811600 | 0.11104600 | 0.53043200 |
| H | -5.35736000 | 6.29842800 | 1.82022300 | C | -1.64679400 | 0.12479500 | -0.08464800 |
| H | -3.99627000 | 4.77770600 | 2.97002400 | O | -2.60314000 | 0.69745700 | 0.46964800 |
| H | -4.75782000 | 3.47560600 | -0.32574700 | C | -1.72078400 | -0.55187600 | -1.35162800 |
| H | -3.96538700 | 1.96372100 | 0.03512700 | C | -0.61173800 | -1.29112200 | -1.70686500 |
| H | -2.94427800 | 2.84813600 | 3.25728000 | H | 0.97332200 | -3.05096200 | -0.84549800 |
| H | -2.99266600 | 1.62418800 | 1.98592500 | H | -0.31843900 | 0.65282600 | 1.40735400 |
| C | -2.88947500 | 5.90451800 | 7.10122100 | H | -0.66413200 | -1.90017500 | -2.61812800 |
| C | -1.91573800 | 5.04787500 | 6.28864700 | H | -2.78077300 | -1.20694400 | -3.06653700 |
| C | -2.45006300 | 4.57094300 | 4.92897300 | H | -4.72590700 | -3.37531200 | -1.62307600 |
| O | -3.33576000 | 5.26419100 | 4.37201400 | H | -6.13119600 | -0.61186700 | -1.38490000 |
| O | -1.94834700 | 3.50670200 | 4.45465700 | H | -4.75462300 | -0.17186500 | -4.01306400 |
| H | -2.42056800 | 6.27461500 | 8.02711700 | H | -5.86654800 | 1.90915700 | -2.06870900 |
| H | -1.58955600 | 4.15666500 | 6.84784200 | H | -4.16578300 | -1.10431400 | -0.38357900 |
| H | -0.99443800 | 5.62127700 | 6.07547500 | H | -6.89453600 | 0.91371900 | -3.12509800 |
| C | 4.82173700 | 5.95110100 | 2.41444900 | P | -7.01056600 | -3.11864200 | -2.43390800 |
| C | 3.62790000 | 6.26380400 | 1.53117600 | O | -6.05690300 | -3.99679200 | -1.62519100 |
| O | 2.94890000 | 7.26412900 | 1.69877100 | O | -7.87632500 | -3.59621300 | -3.54207000 |
| H | 5.42674800 | 5.10658100 | 2.05130200 | O | -7.92028000 | -2.31141000 | -1.29582700 |
| N | 3.33425700 | 5.35543500 | 0.55221500 | C | 4.56870700 | 1.14974200 | -5.98587700 |
| C | 2.18667100 | 5.57589000 | -0.28998900 | C | 4.56537100 | 1.84981300 | -4.61663000 |
| C | 1.73631100 | 4.28861800 | -0.96466000 | C | 3.73394500 | 1.19633500 | -3.50820400 |
| O | 2.53095500 | 3.44055800 | -1.34956700 | C | 2.91328000 | -1.54802500 | -3.32230300 |
| H | 3.91043400 | 4.54057300 | 0.36007500 | S | 4.37073900 | -0.49100900 | -3.09308700 |
| H | 1.38737000 | 6.03268300 | 0.31312400 | C | 4.51583700 | -0.57739600 | -1.26495700 |
| N | 0.39552700 | 4.20415600 | -1.17590900 | C | 5.89710400 | -0.24157300 | -0.74265600 |
| C | -0.21981700 | 3.04166500 | -1.80563400 | O | 6.88981900 | -1.01017900 | -1.40460300 |
| C | 0.27148700 | 2.86695500 | -3.26712900 | C | 6.40021500 | 1.20769400 | -0.86706400 |
| O | 0.59401800 | 1.77061800 | -3.70567500 | O | 5.79753400 | 2.11702900 | 0.00327300 |
| C | -1.73718800 | 3.16634200 | -1.75138100 | C | 7.90785700 | 0.99591800 | -0.61309500 |
| H | -0.20985400 | 4.84016800 | -0.66794400 | O | 8.03293900 | 0.99494300 | 0.80155900 |
| H | 0.10535100 | 2.12409900 | -1.29399900 | C | 8.16948500 | -0.38135100 | -1.24748800 |
| H | -2.07584200 | 4.09078700 | -2.24812100 | H | 5.00425000 | 0.13863500 | -5.92221800 |
| H | -2.07157900 | 3.17874000 | -0.70553800 | H | 5.60363600 | 1.98385100 | -4.26486800 |
| H | -2.22627600 | 2.31262200 | -2.23979900 | H | 4.15443000 | 2.86833500 | -4.72273400 |
| N | 0.25104100 | 4.00041100 | -4.02331500 | H | 2.67245000 | 1.07662400 | -3.77895700 |
| C | 0.55846000 | 4.04753900 | -5.44483000 | H | 3.74789900 | 1.78529700 | -2.57899200 |
| H | 0.01442000 | 4.86398300 | -3.54892800 | H | 2.59781200 | -1.43015000 | -4.36644400 |
| H | 0.66643900 | 3.01288400 | -5.79602700 | H | 3.24973100 | -2.57191500 | -3.11249100 |
| C | 5.52673800 | -5.45490700 | -0.69101000 | H | 2.12094600 | -1.25849100 | -2.60995900 |
| C | 4.52655700 | -4.39070900 | -0.31280300 | H | 8.81950600 | -0.99007800 | -0.59512800 |
| O | 3.90044400 | -3.76345900 | -1.18783100 | H | 8.96236100 | 0.94620300 | 1.05626500 |
| H | 5.88874800 | -6.03483900 | 0.16856400 | H | 6.25261200 | 1.56522800 | -1.90029700 |
| N | 4.37174200 | -4.16804500 | 0.99445600 | H | 5.85559500 | -0.48434100 | 0.33677200 |
| C | 3.68660100 | -3.03452700 | 1.58497300 | H | 3.72818800 | 0.03347700 | -0.79883600 |
| C | 4.76886400 | -2.24492400 | 2.34368300 | H | 6.24565100 | 2.00126700 | 0.85774800 |
| O | 5.80797200 | -2.77794300 | 2.70975500 | H | 8.52573200 | 1.79354800 | -1.06022800 |
| C | 2.52954300 | -3.51182700 | 2.47441900 | H | 4.28783800 | -1.63760400 | -1.07528300 |
| O | 2.97557500 | -4.59222700 | 3.25285700 | H | 1.61337700 | -9.21812600 | 1.84098200 |
| H | 4.96408800 | -4.66052200 | 1.65664000 | H | 0.65506100 | -8.45566700 | 3.13924400 |

| | | | |
|---|---|---|---|
| H | 6.38268000 | -4.95748400 | -1.17446900 |
| H | 5.07580100 | -6.12734800 | -1.43477900 |
| H | 5.74803300 | -0.55984600 | 4.21559500 |
| H | 5.06294300 | 0.88921600 | 3.41359700 |
| H | 8.64128500 | -0.29821900 | -2.24041700 |
| H | 5.16469800 | 1.72241100 | -6.71149800 |
| H | 3.54546500 | 1.05387700 | -6.38361800 |
| H | 5.44878300 | 6.85073600 | 2.48802700 |
| H | 4.45618000 | 5.72127000 | 3.42720800 |
| H | 2.40594000 | 6.30287400 | -1.09901100 |
| H | 1.49844800 | 4.59086800 | -5.64333600 |
| H | -0.25550600 | 4.53297400 | -6.00587000 |
| H | -3.11552000 | 7.68949600 | 1.79458900 |
| H | -3.22716500 | 6.76428800 | 6.50450600 |
| H | -3.78690100 | 5.32869300 | 7.37911600 |
| H | -5.72778200 | 1.89564000 | -4.89658400 |
| H | -8.77523500 | -2.11481900 | -1.70422400 |
| H | -6.48732800 | -1.32654900 | 2.64790400 |

#### Model3” – TS2

| | | | |
|---|---|---|---|
| O | -0.48909200 | 4.19855400 | -0.79783300 |
| H | 0.40487300 | 4.92258200 | 0.07144600 |
| H | -1.45594700 | 4.39504100 | -0.91776400 |
| O | 0.17165200 | 6.03583200 | 4.48699300 |
| H | -0.04030200 | 6.87554100 | 4.91596700 |
| H | 0.93451000 | 6.23386000 | 3.90981800 |
| O | 0.83651000 | -1.23262200 | 3.43235300 |
| H | 1.08193500 | -2.11216200 | 3.80691900 |
| H | 1.17003300 | -0.62415800 | 4.10506800 |
| C | 0.50512500 | 8.73352900 | 1.68678900 |
| C | 0.09523900 | 7.32757500 | 1.20747000 |
| C | 1.15207500 | 6.27035600 | 1.46670600 |
| O | 1.18027200 | 5.18427100 | 0.73728600 |
| O | 1.96914700 | 6.40270900 | 2.38022700 |
| H | 1.40923100 | 9.08332600 | 1.16721500 |
| H | -0.16513600 | 7.32049100 | 0.13714900 |
| H | -0.80395600 | 6.97293700 | 1.74806600 |
| C | 6.30053500 | 0.78972400 | 1.23920200 |
| C | 6.38062100 | 2.26754500 | 0.87487400 |
| N | 5.67285900 | 3.13338500 | 1.81882800 |
| C | 4.45289000 | 3.67687100 | 1.68572200 |
| N | 3.69769800 | 3.51207500 | 0.60938300 |
| N | 3.97258300 | 4.40451600 | 2.72092500 |
| H | 5.26328400 | 0.42089900 | 1.24850800 |
| H | 6.87305000 | 0.21380100 | 0.49774000 |
| H | 6.02631300 | 2.47851900 | -0.14020600 |
| H | 7.43026800 | 2.59591000 | 0.85190700 |
| H | 6.11057600 | 3.26093400 | 2.72331700 |
| H | 3.96299800 | 3.01100700 | -0.27346500 |
| H | 2.80722000 | 4.00901100 | 0.59046500 |
| H | 4.61773200 | 4.66897100 | 3.45390500 |
| H | 3.19035800 | 5.05788100 | 2.56948400 |
| C | 1.78645800 | -7.05795000 | 1.36806600 |
| C | 2.86237900 | -6.56351700 | 0.40158900 |
| C | 4.12706400 | -6.06482100 | 1.11893000 |
| N | 3.89592000 | -4.92759800 | 1.98520800 |
| C | 3.80045100 | -3.66281300 | 1.56231200 |
| N | 4.08126200 | -3.31842600 | 0.28084300 |
| N | 3.46724800 | -2.71043900 | 2.42792000 |
| H | 1.53906200 | -6.29088100 | 2.11796400 |
| H | 0.86313100 | -7.34152900 | 0.83658400 |
| H | 2.46300200 | -5.74506400 | -0.22378100 |
| H | 3.15160300 | -7.36944700 | -0.29541500 |
| H | 4.91774200 | -5.82712200 | 0.38603800 |
| H | 4.53224000 | -6.87341800 | 1.74762800 |
| H | 3.54762900 | -5.11249700 | 2.96875300 |
| H | 4.19391000 | -4.02975000 | -0.42565200 |
| H | 3.81536800 | -2.37949900 | -0.02644800 |
| H | 3.00788900 | -2.99043600 | 3.32707600 |
| H | 3.18218200 | -1.81566100 | 2.02258700 |
| C | 2.47228300 | -5.86375400 | 7.23446100 |
| C | 1.61501600 | -4.90595100 | 6.40525400 |
| C | 2.21497000 | -4.54180400 | 5.04069800 |
| O | 2.86163800 | -5.42526900 | 4.42807900 |
| O | 2.00827400 | -3.36412600 | 4.61449100 |
| H | 1.96040500 | -6.15784300 | 8.16463100 |
| H | 1.40960100 | -3.96900200 | 6.94527100 |
| H | 0.63140600 | -5.36794800 | 6.20063000 |
| C | -5.44874900 | -5.03593500 | 2.99154700 |
| C | -4.30808300 | -5.57166300 | 2.14586900 |
| O | -3.73216200 | -6.61337500 | 2.40979800 |
| H | -5.06265100 | -4.81269300 | 3.99780100 |
| N | -3.95153000 | -4.80667400 | 1.07048000 |
| C | -2.85162100 | -5.19293900 | 0.23069700 |
| C | -2.31710500 | -3.99203700 | -0.53679200 |
| O | -3.02286900 | -3.02656000 | -0.80871400 |
| H | -4.42711500 | -3.94297000 | 0.83482600 |
| H | -2.07297300 | -5.66798900 | 0.84855900 |
| N | -1.02824300 | -4.10633400 | -0.94794800 |
| C | -0.32967600 | -3.02913700 | -1.64077600 |
| C | -0.78990800 | -2.84847400 | -3.10518400 |
| O | -0.78121300 | -1.73835200 | -3.62015400 |
| C | 1.17401700 | -3.28320200 | -1.59408000 |
| H | -0.46272900 | -4.83890300 | -0.53121800 |
| H | -0.56488600 | -2.07228300 | -1.15148700 |
| H | 1.41627700 | -4.26023900 | -2.04472800 |
| H | 1.52851300 | -3.27235300 | -0.55385500 |
| H | 1.72851700 | -2.51112500 | -2.14283000 |
| N | -1.12013100 | -3.99208200 | -3.76320400 |
| C | -1.46080300 | -4.06693100 | -5.17686000 |
| H | -1.12359300 | -4.84532100 | -3.21654000 |
| H | -1.32986300 | -3.06471500 | -5.60550900 |
| C | -4.78948100 | 6.21979600 | -0.62941000 |
| C | -3.79721700 | 5.14215600 | -0.24714300 |
| O | -3.08631800 | 4.59351300 | -1.10428400 |
| H | -5.25759400 | 6.70336300 | 0.23837800 |
| N | -3.76919700 | 4.83161500 | 1.05283600 |
| C | -3.16345000 | 3.65256200 | 1.63599800 |
| C | -4.27697900 | 3.04777200 | 2.51466800 |
| O | -5.22645900 | 3.72828200 | 2.88383300 |
| C | -1.87395700 | 3.98852700 | 2.40047300 |
| O | -2.10232200 | 5.11250800 | 3.20826700 |
| H | -4.44295800 | 5.26130600 | 1.68404100 |
| H | -2.89846000 | 2.93998700 | 0.84319600 |
| H | -1.09295700 | 4.17196600 | 1.64232200 |
| H | -1.56491800 | 3.09400800 | 2.97315200 |
| H | -1.28452400 | 5.33689300 | 3.69939900 |
| N | -4.16761300 | 1.73319400 | 2.80894100 |
| C | -5.23571900 | 1.08732500 | 3.55833500 |
| H | -3.36228200 | 1.20989400 | 2.46905800 |
| H | -6.15510200 | 1.02071200 | 2.95193200 |
| C | 5.46447100 | -2.21947200 | -3.27992400 |
| O | 5.03724600 | -3.19722400 | -4.21280600 |
| C | 4.62353300 | -0.95320900 | -3.18300800 |
| O | 3.39782800 | -1.12961700 | -2.47007400 |
| C | 5.35942100 | 0.19467000 | -2.45630400 |
| O | 6.04501700 | 0.95781300 | -3.41374100 |
| C | 4.21376600 | 0.91061900 | -1.66932200 |
| O | 4.09039000 | 2.29901000 | -1.79785500 |
| C | 2.95134300 | 0.20406400 | -2.18946900 |
| N | -0.37382500 | 1.30640600 | -0.77672600 |
| C | -0.40943900 | 0.74758000 | 0.48248900 |
| O | -1.37901300 | 0.88404200 | 1.23592900 |
| N | 0.69862600 | -0.00355500 | 0.85977600 |
| C | 1.83164400 | -0.21426400 | 0.08648400 |
| O | 2.80965500 | -0.80801600 | 0.56672700 |
| C | 1.77456200 | 0.29079800 | -1.26733500 |
| C | 0.66978300 | 1.03002600 | -1.59985600 |
| H | -0.44275700 | 3.21616400 | -0.72315900 |
| H | 0.69334900 | -0.43773200 | 1.80552600 |

| | | | |
|---|---|---|---|
| H | 0.60211800 | 1.46767700 | -2.60202000 |
| H | 2.67899500 | 0.70310500 | -3.13940400 |
| H | 5.00680900 | 2.71416400 | -2.04402500 |
| H | 6.08027500 | -0.22071200 | -1.73247700 |
| H | 4.40495100 | -0.58043000 | -4.20431100 |
| H | 5.48543600 | -2.70652200 | -2.29085200 |
| H | 4.33066800 | 0.62796000 | -0.61289700 |
| H | 6.49639600 | -1.88265800 | -3.50000800 |
| P | 7.18254500 | 2.10053700 | -2.94891700 |
| O | 6.40036700 | 3.14143600 | -2.14973800 |
| O | 8.04661000 | 2.40292900 | -4.11805200 |
| O | 8.03697600 | 1.23850900 | -1.80747200 |
| C | -5.06605200 | -0.66680900 | -5.63670000 |
| C | -4.84788900 | -1.57493200 | -4.42504200 |
| C | -3.81669600 | -1.04456300 | -3.43349400 |
| C | -2.28612400 | 1.01411500 | -1.67215300 |
| S | -4.33411600 | 0.44802700 | -2.48316700 |
| C | -5.00198300 | -0.30195500 | -0.94159800 |
| C | -6.48831000 | -0.10283200 | -0.73763200 |
| O | -6.76337900 | 1.25996300 | -0.46427900 |
| C | -7.41528800 | -0.43818500 | -1.91605900 |
| O | -7.58831400 | -1.79999200 | -2.16790500 |
| C | -8.69269600 | 0.29823800 | -1.48820200 |
| O | -9.35206900 | -0.59843200 | -0.60321500 |
| C | -8.12585000 | 1.54223200 | -0.78690700 |
| H | -5.43948500 | 0.32450500 | -5.33047500 |
| H | -5.79820600 | -1.79474400 | -3.91152500 |
| H | -4.46321800 | -2.55243100 | -4.76751800 |
| H | -2.89478600 | -0.74587500 | -3.95194400 |
| H | -3.52990700 | -1.79495500 | -2.68523600 |
| H | -1.77777400 | 0.35520200 | -2.37219400 |
| H | -2.39501900 | 2.07445200 | -1.88641400 |
| H | -2.54156600 | 0.66272800 | -0.67704100 |
| H | -8.70130500 | 1.76358200 | 0.12928700 |
| H | -10.24186700 | -0.27638100 | -0.41573600 |
| H | -7.00823900 | 0.03605900 | -2.82375600 |
| H | -6.78504200 | -0.74178500 | 0.12086600 |
| H | -4.74151900 | -1.36824000 | -0.95332100 |
| H | -8.28806400 | -2.09492900 | -1.56177000 |
| H | -9.34047900 | 0.55235900 | -2.34465800 |
| H | -4.46815200 | 0.17272900 | -0.10458200 |
| H | -0.30695000 | 9.45076400 | 1.49925400 |
| H | 0.71938700 | 8.72134700 | 2.76470500 |
| H | -5.57334800 | 5.76026700 | -1.25186400 |
| H | -4.28407400 | 6.97382200 | -1.25014800 |
| H | -5.47269100 | 1.66471900 | 4.46360500 |
| H | -4.91634100 | 0.07616800 | 3.84493300 |
| H | -8.15538900 | 2.42993800 | -1.44085700 |
| H | -5.80059000 | -1.09926900 | -6.33310800 |
| H | -4.12479200 | -0.51029300 | -6.18838900 |
| H | -5.91993000 | -4.13199800 | 2.57838900 |
| H | -6.20593600 | -5.82564700 | 3.10172300 |
| H | -3.14979900 | -5.95478700 | -0.51871400 |
| H | -2.50657200 | -4.38424100 | -5.32456700 |
| H | -0.79773500 | -4.77079700 | -5.70411000 |
| H | 2.13260400 | -7.94277100 | 1.92596200 |
| H | 2.69749500 | -6.76843000 | 6.65108900 |
| H | 3.43412700 | -5.40018400 | 7.50762800 |
| H | 5.05119100 | -2.78768000 | -5.08798500 |
| H | 8.84054600 | 0.91712500 | -2.24037800 |
| H | 6.73793300 | 0.59590600 | 2.23300800 |

#### Model3'' – Final state

| | | | |
|---|---|---|---|
| O | -0.57316100 | 5.37681900 | -1.16860000 |
| H | 0.26983900 | 5.31060200 | 0.08073900 |
| H | -1.53775800 | 5.10859700 | -1.18089200 |
| O | -0.03064200 | 4.47780500 | 4.68784900 |
| H | -0.33594100 | 4.93029800 | 5.48620800 |
| H | 0.50991000 | 5.14156200 | 4.21583100 |
| O | 0.54153900 | -1.46243800 | 3.15183100 |

| | | | |
|---|---|---|---|
| H | 0.80790900 | -2.33965700 | 3.51702400 |
| H | 0.72596400 | -0.87146500 | 3.89423700 |
| C | 0.11922900 | 8.64325000 | 2.14513800 |
| C | -0.27399900 | 7.30741000 | 1.50223400 |
| C | 0.67975500 | 6.17714400 | 1.81816700 |
| O | 0.83784800 | 5.21897600 | 0.92768600 |
| O | 1.29211600 | 6.11251400 | 2.88237600 |
| H | 1.07272200 | 9.01491600 | 1.73976400 |
| H | -0.38820900 | 7.38944200 | 0.41115400 |
| H | -1.24585900 | 6.94563800 | 1.88796500 |
| C | 6.20949500 | 0.92803300 | 1.62038200 |
| C | 6.14672200 | 2.43027500 | 1.37129300 |
| N | 5.26768400 | 3.12658700 | 2.31193100 |
| C | 4.03911200 | 3.61974300 | 2.09660400 |
| N | 3.41770800 | 3.54296400 | 0.92717200 |
| N | 3.40653200 | 4.20241600 | 3.13941900 |
| H | 5.22337600 | 0.45112400 | 1.50829400 |
| H | 6.90533300 | 0.48199900 | 0.89473600 |
| H | 5.85436700 | 2.67967100 | 0.34511900 |
| H | 7.15011300 | 2.87211400 | 1.46768700 |
| H | 5.58937700 | 3.16590000 | 3.27192800 |
| H | 3.79296100 | 3.12020800 | 0.04350800 |
| H | 2.51222700 | 4.00457300 | 0.85547800 |
| H | 3.94261200 | 4.41434500 | 3.97041400 |
| H | 2.58017400 | 4.79429200 | 2.98533500 |
| C | 1.97729000 | -7.05615300 | 1.05406400 |
| C | 3.10531900 | -6.47336600 | 0.20282100 |
| C | 4.27493300 | -5.93467000 | 1.04196200 |
| N | 3.90607500 | -4.84862000 | 1.92700900 |
| C | 3.79353100 | -3.57224400 | 1.55604500 |
| N | 4.16970500 | -3.15457900 | 0.32025800 |
| N | 3.34881400 | -2.67553300 | 2.43396300 |
| H | 1.61153500 | -6.32280900 | 1.78922500 |
| H | 1.12762300 | -7.38125500 | 0.43214600 |
| H | 2.71648100 | -5.65752100 | -0.43246700 |
| H | 3.49872500 | -7.23912100 | -0.48829100 |
| H | 5.10826000 | -5.62359700 | 0.38797200 |
| H | 4.67292100 | -6.74390800 | 1.67433800 |
| H | 3.51099200 | -5.09380800 | 2.88336900 |
| H | 4.34697800 | -3.82800700 | -0.41034000 |
| H | 3.89534300 | -2.21539400 | 0.03005100 |
| H | 2.83113200 | -3.02283500 | 3.27720100 |
| H | 3.05094200 | -1.77700400 | 2.05220000 |
| C | 2.27645800 | -6.17012300 | 7.00691200 |
| C | 1.47225000 | -5.11737600 | 6.23850000 |
| C | 2.07132300 | -4.68760600 | 4.89091900 |
| O | 2.81869100 | -5.49764800 | 4.29186700 |
| O | 1.75899000 | -3.53200800 | 4.46374700 |
| H | 1.75016900 | -6.48241300 | 7.92286800 |
| H | 1.31698300 | -4.20545500 | 6.83652500 |
| H | 0.46029000 | -5.50351200 | 6.01549100 |
| C | -5.40766000 | -5.36255300 | 2.34517300 |
| C | -4.17615700 | -5.75169500 | 1.55021200 |
| O | -3.47120000 | -6.69648400 | 1.86533200 |
| H | -5.96557900 | -4.52221900 | 1.90562500 |
| N | -3.90234300 | -4.97641900 | 0.45632500 |
| C | -2.74212000 | -5.23963000 | -0.34846300 |
| C | -2.14874300 | -3.95821000 | -0.91766800 |
| O | -2.81863500 | -2.95329700 | -1.10453400 |
| H | -4.47695200 | -4.17788400 | 0.20618100 |
| H | -2.01914800 | -5.79662700 | 0.26738500 |
| N | -0.83748500 | -4.05453200 | -1.27870100 |
| C | -0.10140600 | -2.93591600 | -1.85543700 |
| C | -0.44398100 | -2.68270900 | -3.34452300 |
| O | -0.35322400 | -1.55739000 | -3.81352200 |
| C | 1.39667200 | -3.17884400 | -1.70290600 |
| H | -0.30271500 | -4.83722000 | -0.91702400 |
| H | -0.39870600 | -2.01221500 | -1.34087900 |
| H | 1.68102400 | -4.13470700 | -2.17410100 |
| H | 1.66912100 | -3.21457600 | -0.63891800 |

| | | | |
|---|---|---|---|
| H | 1.98325600 | -2.38463700 | -2.18095500 |
| N | -0.75822800 | -3.79264900 | -4.06672400 |
| C | -0.98630300 | -3.81038300 | -5.50401800 |
| H | -0.84596800 | -4.65855500 | -3.54808200 |
| H | -0.86707000 | -2.78267100 | -5.87151300 |
| C | -4.93840700 | 6.09334300 | -0.61796300 |
| C | -3.90739900 | 5.03759200 | -0.30796700 |
| O | -3.06862000 | 4.67353700 | -1.15178100 |
| H | -5.47271200 | 6.44662000 | 0.27405500 |
| N | -4.00231400 | 4.50278300 | 0.90621900 |
| C | -3.37371300 | 3.27791900 | 1.34681100 |
| C | -4.41293500 | 2.68263100 | 2.31298500 |
| O | -5.27307900 | 3.39069800 | 2.82640000 |
| C | -1.98013200 | 3.50730100 | 1.95549600 |
| O | -1.99981700 | 4.62004500 | 2.81016100 |
| H | -4.72912200 | 4.80667900 | 1.55591100 |
| H | -3.25279000 | 2.59902000 | 0.49150100 |
| H | -1.27402200 | 3.65930600 | 1.11708600 |
| H | -1.66582000 | 2.58281400 | 2.46958000 |
| H | -1.34752800 | 4.48064700 | 3.53036900 |
| N | -4.37370200 | 1.34917300 | 2.49898200 |
| C | -5.44585300 | 0.72316300 | 3.25279900 |
| H | -3.65335800 | 0.79678900 | 2.03957400 |
| H | -6.41979800 | 0.90529700 | 2.76987200 |
| C | 5.79554800 | -1.85995500 | -3.11140200 |
| O | 5.40924500 | -2.78671000 | -4.10469300 |
| C | 4.90813700 | -0.63598800 | -2.99960300 |
| O | 3.64271900 | -0.91556100 | -2.40276500 |
| C | 5.51922900 | 0.50234900 | -2.16023600 |
| O | 6.23131300 | 1.34187500 | -3.02983500 |
| C | 4.27092900 | 1.11785000 | -1.43891900 |
| O | 4.09811000 | 2.50556100 | -1.50474000 |
| C | 3.10325400 | 0.37487600 | -2.11558700 |
| N | -0.50243700 | 0.92197500 | -1.19069300 |
| C | -0.63397900 | 0.43647000 | 0.11092600 |
| O | -1.67275900 | 0.54461100 | 0.74541300 |
| N | 0.50702900 | -0.16541500 | 0.61264200 |
| C | 1.76074800 | -0.20287500 | 0.00893400 |
| O | 2.72829700 | -0.67124800 | 0.61615000 |
| C | 1.82311800 | 0.32010600 | -1.34175100 |
| C | 0.67914500 | 0.83094300 | -1.86940000 |
| H | -0.21719000 | 5.18717700 | -2.04590800 |
| H | 0.43902600 | -0.60966700 | 1.55401800 |
| H | 0.65735000 | 1.21027400 | -2.89104300 |
| H | 2.90281600 | 0.90805000 | -3.06529100 |
| H | 5.02182400 | 2.96332400 | -1.64218400 |
| H | 6.20291400 | 0.09096300 | -1.39927400 |
| H | 4.74772700 | -0.21337800 | -4.01228600 |
| H | 5.79131700 | -2.39654000 | -2.14787500 |
| H | 4.30875800 | 0.78009200 | -0.39257200 |
| H | 6.82592200 | -1.48742900 | -3.27528800 |
| P | 7.28748800 | 2.49862300 | -2.43026700 |
| O | 6.40188200 | 3.43392400 | -1.60452800 |
| O | 8.18885900 | 2.92923500 | -3.52786900 |
| O | 8.12446700 | 1.60251000 | -1.30543900 |
| C | -4.67731800 | -0.50799800 | -5.99662800 |
| C | -4.40014300 | -1.51021800 | -4.87356900 |
| C | -3.78010100 | -0.89757900 | -3.61798800 |
| C | -1.63931800 | 1.61292200 | -1.79953400 |
| S | -4.87348700 | 0.38037000 | -2.85666300 |
| C | -4.85820300 | -0.09700200 | -1.08251000 |
| C | -6.22748500 | 0.12930900 | -0.46450500 |
| O | -6.73408900 | 1.42132500 | -0.80379300 |
| C | -7.29956400 | -0.88433800 | -0.90439500 |
| O | -7.30697900 | -2.06602600 | -0.14736900 |
| C | -8.57921100 | -0.04669700 | -0.77322300 |
| O | -8.90161700 | -0.09255300 | 0.61232300 |
| C | -8.09151000 | 1.32662600 | -1.22792500 |
| H | -5.39773500 | 0.25983800 | -5.67135400 |
| H | -5.33485500 | -2.02805900 | -4.59700700 |
| H | -3.70528700 | -2.28802700 | -5.23873500 |
| H | -2.80830500 | -0.43895600 | -3.85944900 |
| H | -3.58399000 | -1.66819500 | -2.86137600 |
| H | -1.47741700 | 1.65792900 | -2.88210200 |
| H | -1.75110500 | 2.63105800 | -1.40061900 |
| H | -2.56382200 | 1.06247900 | -1.61936400 |
| H | -8.67449200 | 2.14655700 | -0.77536300 |
| H | -9.71757100 | 0.39642200 | 0.77207400 |
| H | -7.13572700 | -1.14978900 | -1.96058600 |
| H | -6.13001600 | 0.05208600 | 0.63376000 |
| H | -4.58002300 | -1.15644500 | -0.98381600 |
| H | -7.79705400 | -1.84812800 | 0.66237000 |
| H | -9.41113900 | -0.43232800 | -1.38940600 |
| H | -4.10005500 | 0.48538700 | -0.53980200 |
| H | -0.65432000 | 9.40055600 | 1.95202200 |
| H | 0.23598400 | 8.53193800 | 3.23246300 |
| H | -5.66714900 | 5.65843000 | -1.32083900 |
| H | -4.45925700 | 6.94158600 | -1.12764500 |
| H | -5.49707100 | 1.12757400 | 4.27497500 |
| H | -5.26532100 | -0.35884100 | 3.30189700 |
| H | -8.15685200 | 1.41319200 | -2.32909300 |
| H | -5.09697600 | -1.00604900 | -6.88482000 |
| H | -3.75255800 | 0.00868700 | -6.30348900 |
| H | -6.06705400 | -6.23956800 | 2.42054800 |
| H | -5.09587300 | -5.10077600 | 3.36761400 |
| H | -2.97738100 | -5.89299900 | -1.21421000 |
| H | -2.00399800 | -4.15833900 | -5.74451700 |
| H | -0.25704700 | -4.46052700 | -6.01408000 |
| H | 2.32499000 | -7.93099700 | 1.62648200 |
| H | 2.44676600 | -7.05426100 | 6.37550700 |
| H | 3.26555800 | -5.78032700 | 7.29635200 |
| H | 5.43855200 | -2.33226400 | -4.95688200 |
| H | 8.96858900 | 1.35478400 | -1.70888300 |
| H | 6.58172700 | 0.70246300 | 2.63372400 |

#### Model4 – Initial state

| | | | |
|---|---|---|---|
| O | -1.72577500 | 4.16923200 | -3.90412900 |
| H | -1.45943100 | 4.53884700 | -3.01962700 |
| H | -1.24504200 | 4.71315400 | -4.54185200 |
| O | -1.96417800 | 0.51076300 | -1.92366400 |
| H | -2.86338800 | 0.26315800 | -2.23720900 |
| H | -1.59154600 | 0.87063000 | -2.77999200 |
| O | -4.20086800 | 0.19859100 | -3.45810000 |
| H | -4.95839200 | 0.78978000 | -3.34612900 |
| H | -4.58961000 | -0.68067200 | -3.70978600 |
| O | -1.62122300 | 1.45305000 | -4.34076600 |
| H | -1.66811900 | 2.43428100 | -4.24714400 |
| H | -2.54096600 | 1.15948100 | -4.46740200 |
| C | -5.16953200 | 7.80872400 | -4.09332900 |
| C | -5.07047100 | 6.27892000 | -4.14093200 |
| C | -5.06636500 | 5.55766500 | -2.77863700 |
| C | -6.28076500 | 5.90788100 | -1.91396000 |
| C | -4.97354400 | 4.03895100 | -2.98309800 |
| H | -6.13557300 | 8.14609900 | -3.68655200 |
| H | -4.14614500 | 5.98850800 | -4.67052400 |
| H | -5.90539200 | 5.88026900 | -4.74794700 |
| H | -4.16115400 | 5.87982800 | -2.23184300 |
| H | -6.23782600 | 5.36941600 | -0.95373800 |
| H | -7.22190000 | 5.62455400 | -2.41748000 |
| H | -6.33223300 | 6.98279800 | -1.68400900 |
| H | -4.96601900 | 3.50824900 | -2.01809000 |
| H | -5.84491100 | 3.67252300 | -3.55512000 |
| H | -4.05813500 | 3.76744800 | -3.52794600 |
| C | 0.12615700 | 8.64368100 | -0.83757200 |
| C | 0.15924200 | 7.25002100 | -1.45929400 |
| C | -0.99064500 | 6.29830200 | -1.10809800 |
| O | -1.97043700 | 6.66833200 | -0.45493600 |
| O | -0.82943900 | 5.08964400 | -1.55449600 |
| H | 0.15324300 | 8.58622300 | 0.26148500 |
| H | 1.08687000 | 6.71519000 | -1.19237400 |

| | | | |
|---|---|---|---|
| H | 0.17642000 | 7.31214300 | -2.56266600 |
| C | -3.78535700 | -7.17317300 | -1.54953600 |
| C | -3.91766300 | -6.53309400 | -0.16629300 |
| C | -5.06268500 | -5.51534000 | -0.06670400 |
| N | -4.80693200 | -4.28178200 | -0.78413800 |
| C | -4.33248700 | -3.15424200 | -0.24652200 |
| N | -3.97041100 | -3.07369600 | 1.05352200 |
| N | -4.19743700 | -2.08160100 | -1.02251600 |
| H | -3.70329500 | -6.41126200 | -2.33999800 |
| H | -2.90142800 | -7.82826700 | -1.60302300 |
| H | -2.98200100 | -6.01558500 | 0.10408500 |
| H | -4.07829900 | -7.31569400 | 0.59611900 |
| H | -5.28976800 | -5.29026300 | 0.98803600 |
| H | -5.98461600 | -5.95733800 | -0.47965700 |
| H | -4.86448100 | -4.31118500 | -1.83765900 |
| H | -4.11642400 | -3.85042900 | 1.68015600 |
| H | -3.74903400 | -2.16602200 | 1.46735000 |
| H | -4.53672500 | -2.11436500 | -2.00560600 |
| H | -3.74790000 | -1.25944400 | -0.62363900 |
| C | -4.88511800 | -4.58090900 | -6.43196500 |
| C | -3.53489200 | -3.90584200 | -6.67469400 |
| O | -3.25392400 | -3.36347700 | -7.73156500 |
| C | -5.74882700 | -3.63753500 | -5.55871300 |
| C | -5.21728700 | -3.43211300 | -4.13067100 |
| O | -4.85791000 | -4.46133600 | -3.49370100 |
| O | -5.17929600 | -2.26241000 | -3.65380100 |
| H | -5.35746000 | -4.74607500 | -7.40832100 |
| H | -6.76121200 | -4.06521600 | -5.45540500 |
| H | -5.85407900 | -2.65723600 | -6.04539500 |
| N | -2.71026600 | -3.85221200 | -5.58056000 |
| C | -1.64852600 | -2.86093800 | -5.51760400 |
| C | -0.76169200 | -2.97313000 | -4.27116000 |
| C | -1.56912800 | -2.81441300 | -2.97063900 |
| C | 0.35172700 | -1.92131300 | -4.37463200 |
| C | -0.76182800 | -2.97979300 | -1.68197800 |
| H | -3.07200700 | -4.23192100 | -4.71046300 |
| H | -2.08291600 | -1.84207100 | -5.55161400 |
| H | -0.30146400 | -3.98075800 | -4.27339000 |
| H | -2.40277800 | -3.53415400 | -2.95179000 |
| H | -2.03212300 | -1.81632100 | -2.97005100 |
| H | 0.92039200 | -2.03881800 | -5.31087400 |
| H | 1.06530300 | -1.99678800 | -3.53988400 |
| H | -0.07585200 | -0.90439600 | -4.36603700 |
| H | -1.41909400 | -2.88961300 | -0.80314400 |
| H | 0.02135400 | -2.21302300 | -1.57835200 |
| H | -0.26808900 | -3.96697000 | -1.63626700 |
| C | 3.16354700 | -6.69778000 | -0.80558500 |
| C | 2.70350500 | -5.53614500 | 0.07181400 |
| O | 3.40767700 | -4.55128300 | 0.27189900 |
| H | 2.45303300 | -6.88479100 | -1.62571400 |
| N | 1.46977400 | -5.64825800 | 0.65699200 |
| C | 0.91814100 | -4.52967900 | 1.44135300 |
| C | 1.68933700 | -4.25566400 | 2.72847800 |
| O | 1.79946500 | -3.12407700 | 3.17843100 |
| C | -0.54480100 | -4.95298900 | 1.70040200 |
| C | -0.51728900 | -6.48204000 | 1.57812300 |
| C | 0.48676700 | -6.71794700 | 0.44677300 |
| H | 0.97256500 | -3.59601800 | 0.86371000 |
| H | -1.18815200 | -4.51646100 | 0.92175100 |
| H | -0.91430900 | -4.58908100 | 2.66925100 |
| H | -1.50084900 | -6.91922600 | 1.36399100 |
| H | -0.14887900 | -6.93455500 | 2.51262700 |
| H | -0.00134100 | -6.61815500 | -0.54033900 |
| H | 0.95732900 | -7.71102500 | 0.48897600 |
| N | 2.14754800 | -5.37541100 | 3.34255600 |
| C | 2.78098500 | -5.36902100 | 4.63158200 |
| H | 2.09285800 | -6.24181700 | 2.82316800 |
| H | 2.66444000 | -4.36718600 | 5.06417900 |
| C | 7.71698300 | 3.27378200 | 3.91037400 |
| C | 6.58096000 | 3.48733300 | 2.92302500 |
| O | 5.40854100 | 3.25882000 | 3.20380700 |
| H | 8.66560500 | 3.04209000 | 3.40304300 |
| N | 6.94804800 | 4.04026000 | 1.73224100 |
| C | 5.93945100 | 4.45143500 | 0.79477800 |
| C | 5.16606100 | 3.28957400 | 0.14412400 |
| O | 5.56275500 | 2.11903000 | 0.19049200 |
| H | 7.90326200 | 3.90170600 | 1.38587700 |
| H | 5.21445600 | 5.12211400 | 1.28383300 |
| N | 4.06784500 | 3.64100500 | -0.53324600 |
| C | 3.34562400 | 2.73088600 | -1.40124500 |
| C | 4.24088500 | 2.29333100 | -2.57390700 |
| O | 5.12548500 | 3.00760100 | -3.00923700 |
| C | 2.10255100 | 3.47802300 | -1.93598400 |
| C | 1.15689400 | 2.58734600 | -2.73506700 |
| O | 1.47862100 | 4.03094700 | -0.80379900 |
| H | 3.65512800 | 4.55893300 | -0.40396600 |
| H | 2.99031900 | 1.85976300 | -0.82940300 |
| H | 2.47968300 | 4.28226600 | -2.60532800 |
| H | 0.62364500 | 4.44376900 | -1.09298100 |
| H | 0.28142500 | 3.17098100 | -3.04155800 |
| H | 0.81533200 | 1.73940400 | -2.12238300 |
| H | 1.62213300 | 2.20188400 | -3.65300200 |
| N | 3.90784100 | 1.08305700 | -3.11794100 |
| C | 4.37086300 | 0.68215200 | -4.43625700 |
| H | 3.10390800 | 0.60413600 | -2.72676100 |
| H | 5.13914400 | 1.40120500 | -4.74761800 |
| C | -5.37372100 | -2.15439300 | 4.43217200 |
| O | -6.36238200 | -2.15971200 | 5.42721700 |
| C | -4.26002000 | -1.18276700 | 4.78200100 |
| O | -3.17430500 | -1.26242400 | 3.84401700 |
| C | -4.74760300 | 0.27966400 | 4.78439900 |
| O | -4.64606000 | 0.80600300 | 6.08118200 |
| C | -3.82043400 | 0.95990000 | 3.74893300 |
| O | -3.55949500 | 2.30581600 | 3.92359500 |
| C | -2.59005400 | 0.04165300 | 3.83421200 |
| N | 0.67535500 | 0.69733200 | 2.09421800 |
| C | 0.49356100 | 0.34984600 | 0.76714100 |
| O | 1.40932300 | 0.34163500 | -0.04387100 |
| N | -0.80484100 | 0.00555700 | 0.46777400 |
| C | -1.86496900 | -0.07617900 | 1.35399600 |
| O | -2.99672000 | -0.33222700 | 0.90996000 |
| C | -1.55887300 | 0.15995100 | 2.75582100 |
| C | -0.29759400 | 0.57331300 | 3.04835400 |
| H | 1.58615800 | 1.06698300 | 2.33852200 |
| H | -1.08015500 | -0.02677300 | -0.53960600 |
| H | -0.00240000 | 0.82986500 | 4.06713700 |
| H | -2.09894000 | 0.24812100 | 4.80554200 |
| H | -4.32793100 | 2.69617500 | 4.44924800 |
| H | -5.79084900 | 0.34423200 | 4.43337400 |
| H | -3.89158200 | -1.40587300 | 5.79847800 |
| H | -5.77394900 | -1.89065700 | 3.42746800 |
| H | -4.26027200 | 0.74963400 | 2.75993800 |
| H | -4.97291200 | -3.18138800 | 4.36121100 |
| P | -5.75166300 | 2.01718300 | 6.46562100 |
| O | -5.72651900 | 2.95977700 | 5.25853000 |
| O | -5.54563400 | 2.41361100 | 7.87762200 |
| O | -7.13989600 | 1.08630500 | 6.26664600 |
| C | 6.14303400 | -2.25664400 | 4.61493500 |
| C | 4.99165000 | -2.61642000 | 3.67253800 |
| C | 5.21758700 | -2.42695900 | 2.17469400 |
| C | 4.08917600 | 0.21110800 | 2.04894600 |
| S | 5.61051000 | -0.69524500 | 1.66148300 |
| C | 5.49957500 | -0.84904500 | -0.16993100 |
| C | 6.78412400 | -1.31007200 | -0.87480300 |
| O | 7.27067800 | -0.25687000 | -1.71144300 |
| C | 8.00586900 | -1.71712600 | -0.01969200 |
| O | 7.94238100 | -3.00392300 | 0.51849100 |
| C | 9.13832600 | -1.52412100 | -1.03579200 |
| O | 9.09510700 | -2.66947400 | -1.86252400 |
| C | 8.67580700 | -0.24845700 | -1.76889100 |

| | | | |
|---|---|---|---|
| C | 10.47255000 | 1.44900000 | -1.17740800 |
| C | 10.49982700 | 2.53221000 | -0.32756800 |
| N | 9.26539300 | 2.70705900 | 0.25342600 |
| C | 8.49404200 | 1.74830600 | -0.22646000 |
| N | 9.18027400 | 0.95486600 | -1.10560500 |
| H | 6.36718900 | -1.17684600 | 4.59706800 |
| H | 4.05573000 | -2.11952500 | 3.97128800 |
| H | 4.76297900 | -3.68896000 | 3.76681300 |
| H | 6.09359300 | -2.98148800 | 1.80561600 |
| H | 4.33432700 | -2.74218200 | 1.59873200 |
| H | 3.70270400 | -0.22040500 | 2.98094900 |
| H | 4.37898300 | 1.26271400 | 2.18475400 |
| H | 3.36868700 | 0.08220800 | 1.22830700 |
| H | 7.43041300 | 1.63649400 | -0.01978400 |
| H | 9.01151800 | -0.23692100 | -2.81906700 |
| H | 9.89504100 | -2.72261300 | -2.39993900 |
| H | 8.14130500 | -1.00588200 | 0.80887800 |
| H | 6.52474000 | -2.17497100 | -1.50913600 |
| H | 4.65855300 | -1.52957400 | -0.35742800 |
| H | 8.16878100 | -3.61324500 | -0.20445800 |
| H | 10.12245500 | -1.39062500 | -0.55843500 |
| H | 5.25174500 | 0.17241200 | -0.49722000 |
| C | -7.52709000 | 1.42858900 | -1.16092100 |
| C | -6.59761600 | 0.60948900 | -0.26992400 |
| C | -6.37635000 | 1.19840300 | 1.12837600 |
| C | -5.57541100 | 2.50401100 | 1.19638600 |
| N | -4.27624200 | 2.33252400 | 0.55555700 |
| C | -3.42940100 | 3.31968700 | 0.21421400 |
| N | -3.70347400 | 4.59459000 | 0.50686800 |
| N | -2.31184900 | 3.01628700 | -0.45210600 |
| H | -7.12490400 | 2.43541700 | -1.35777500 |
| H | -5.62897400 | 0.47208400 | -0.77921600 |
| H | -7.00829000 | -0.40731400 | -0.15269600 |
| H | -5.86892200 | 0.44782300 | 1.75433300 |
| H | -7.34824300 | 1.38360700 | 1.61509500 |
| H | -5.45543900 | 2.79618000 | 2.25463700 |
| H | -6.11638300 | 3.31251400 | 0.67664200 |
| H | -3.89530400 | 1.38692700 | 0.53789400 |
| H | -4.37691300 | 4.80226600 | 1.23275700 |
| H | -3.08568800 | 5.35981200 | 0.16783500 |
| H | -2.25097100 | 2.12148700 | -0.95121600 |
| H | -1.72157100 | 3.79735200 | -0.79642300 |
| H | 0.98541400 | 9.24377900 | -1.17490600 |
| H | -0.79889300 | 9.17406200 | -1.10973400 |
| H | -4.37526800 | 8.23912300 | -3.46020800 |
| H | -5.06740000 | 8.24148200 | -5.10120800 |
| H | 3.24478200 | -7.62650100 | -0.21873400 |
| H | 3.85811200 | -5.60392200 | 4.57188500 |
| H | 11.33793000 | 3.19402200 | -0.11498900 |
| H | 7.85382600 | 4.19931100 | 4.49222400 |
| H | 7.44951900 | 2.46761600 | 4.60606700 |
| H | 11.23088700 | 1.00630700 | -1.81849500 |
| H | 6.41990400 | 5.00882300 | -0.02378800 |
| H | 7.06743600 | -2.79180300 | 4.34358500 |
| H | 4.14559800 | -6.44385900 | -1.22061700 |
| H | 2.30965800 | -6.10026000 | 5.30814200 |
| H | -6.60270700 | -1.24521300 | 5.66000000 |
| H | -7.71829400 | 1.57576500 | 5.66598900 |
| H | -8.51904800 | 1.55340200 | -0.69650600 |
| H | -4.76686300 | -5.54270100 | -5.90934900 |
| H | 5.88601400 | -2.52095300 | 5.65114400 |
| H | 3.54814300 | 0.68863900 | -5.17113900 |
| H | -1.03529600 | -2.95546800 | -6.42653100 |
| H | -7.67736300 | 0.93862500 | -2.13670200 |
| H | 4.81715200 | -0.32518000 | -4.41389500 |
| H | -4.67103200 | -7.78636000 | -1.78203100 |

#### Model4 − TS1

| | | | |
|---|---|---|---|
| O | -1.41140600 | 4.42557500 | -3.36524000 |
| H | -1.09070600 | 4.55131800 | -2.44341800 |
| H | -0.89777700 | 5.05837400 | -3.88464600 |
| O | -2.53443600 | 0.63002500 | -2.17600700 |
| H | -3.43644500 | 0.40696300 | -2.51938900 |
| H | -2.10206000 | 1.00906300 | -2.98089800 |
| O | -4.85491000 | 0.15091400 | -3.44271100 |
| H | -5.66877900 | 0.60618600 | -3.19002100 |
| H | -5.13285100 | -0.77373000 | -3.66080600 |
| O | -1.65222000 | 1.88694600 | -4.40045300 |
| H | -1.61065400 | 2.82188100 | -4.09465600 |
| H | -2.46858900 | 1.81856900 | -4.91533800 |
| C | -4.88367300 | 7.76122400 | -4.38411600 |
| C | -4.98348500 | 6.23171000 | -4.40215600 |
| C | -4.66729700 | 5.51971100 | -3.07284100 |
| C | -5.58954800 | 5.95927700 | -1.93136600 |
| C | -4.72393900 | 3.99806100 | -3.25453100 |
| H | -5.63334200 | 8.21444100 | -3.71716600 |
| H | -4.29696700 | 5.83567000 | -5.17207300 |
| H | -5.99891200 | 5.93639600 | -4.72734800 |
| H | -3.62991200 | 5.77693800 | -2.79278700 |
| H | -5.32145800 | 5.44730400 | -0.99422400 |
| H | -6.64316300 | 5.72171800 | -2.16157600 |
| H | -5.52334500 | 7.04097600 | -1.73984100 |
| H | -4.47588600 | 3.47045500 | -2.32122300 |
| H | -5.73243600 | 3.67550800 | -3.56641600 |
| H | -4.00735600 | 3.65311100 | -4.01436100 |
| C | 0.43603300 | 8.58526000 | -1.16488200 |
| C | 0.70118300 | 7.07101500 | -1.06155900 |
| C | -0.47156900 | 6.16862500 | -0.64365400 |
| O | -1.50853400 | 6.60833100 | -0.17765700 |
| O | -0.28978900 | 4.87228500 | -0.86292500 |
| H | 0.05778400 | 8.97986300 | -0.21058100 |
| H | 1.50800300 | 6.87071700 | -0.33139600 |
| H | 1.08105500 | 6.67479700 | -2.01950300 |
| C | -3.84090800 | -7.15186300 | -1.32494400 |
| C | -4.00698900 | -6.49982100 | 0.04841600 |
| C | -5.21817800 | -5.56019000 | 0.12768100 |
| N | -5.06125700 | -4.34863800 | -0.64953200 |
| C | -4.56136800 | -3.19534600 | -0.18541300 |
| N | -4.14778200 | -3.05470000 | 1.09301200 |
| N | -4.45678900 | -2.16393400 | -1.01540800 |
| H | -3.81188800 | -6.39934600 | -2.12786700 |
| H | -2.91680900 | -7.74982700 | -1.37479900 |
| H | -3.10659000 | -5.91535600 | 0.30395000 |
| H | -4.11222100 | -7.27696200 | 0.82622900 |
| H | -5.44219700 | -5.30183500 | 1.17571500 |
| H | -6.11410800 | -6.08392000 | -0.24487200 |
| H | -5.11034300 | -4.42722500 | -1.69662900 |
| H | -4.28090900 | -3.79392900 | 1.76562600 |
| H | -3.89345800 | -2.12135100 | 1.43261600 |
| H | -4.83783800 | -2.23636000 | -1.98013100 |
| H | -3.99973900 | -1.32159500 | -0.65651500 |
| C | -4.88973500 | -4.70523000 | -6.29305900 |
| C | -3.61698200 | -3.88426600 | -6.49426200 |
| O | -3.36993500 | -3.30039800 | -7.53807500 |
| C | -5.89781600 | -3.87057700 | -5.47255300 |
| C | -5.44087200 | -3.60031000 | -4.03173600 |
| O | -4.99875400 | -4.58232100 | -3.37229300 |
| O | -5.53323100 | -2.42773700 | -3.57561800 |
| H | -5.29623500 | -4.93715000 | -7.28588300 |
| H | -6.85255300 | -4.42019500 | -5.40536400 |
| H | -6.10023700 | -2.91233100 | -5.97206300 |
| N | -2.82755200 | -3.77026200 | -5.38046700 |
| C | -1.86192800 | -2.69058900 | -5.26795200 |
| C | -0.92613400 | -2.83018800 | -4.06180100 |
| C | -1.70636200 | -2.81324400 | -2.73656200 |
| C | 0.11909600 | -1.70676100 | -4.11135900 |
| C | -0.88203300 | -3.16841900 | -1.49847400 |
| H | -3.18318000 | -4.18743400 | -4.52449200 |
| H | -2.38859200 | -1.71755900 | -5.19580200 |
| H | -0.40869800 | -3.80534100 | -4.15029900 |

| | | | |
|---|---|---|---|
| H | -2.56166800 | -3.50507500 | -2.78685700 |
| H | -2.14676800 | -1.81048700 | -2.61550600 |
| H | 0.73239800 | -1.77589500 | -5.02469100 |
| H | 0.79303500 | -1.73411300 | -3.24189800 |
| H | -0.37296800 | -0.71985000 | -4.11319300 |
| H | -1.51467000 | -3.13555100 | -0.59779300 |
| H | -0.04829000 | -2.46900600 | -1.33237500 |
| H | -0.45957000 | -4.18584300 | -1.57911300 |
| C | 3.11853900 | -6.81191600 | -0.60540900 |
| C | 2.68144400 | -5.61034200 | 0.22652900 |
| O | 3.41465400 | -4.63926200 | 0.39552800 |
| H | 2.41030600 | -7.01131500 | -1.42461700 |
| N | 1.44503000 | -5.66823400 | 0.80695600 |
| C | 0.92797600 | -4.51609800 | 1.56755500 |
| C | 1.74238500 | -4.21962400 | 2.82284500 |
| O | 1.94575200 | -3.07571000 | 3.20332400 |
| C | -0.53485300 | -4.90703900 | 1.87826100 |
| C | -0.55408900 | -6.43702600 | 1.75981200 |
| C | 0.42866700 | -6.71018900 | 0.61844000 |
| H | 0.98066800 | -3.60134800 | 0.96034600 |
| H | -1.19502300 | -4.45595700 | 1.12368300 |
| H | -0.86220500 | -4.53412900 | 2.85855500 |
| H | -1.55404300 | -6.84158900 | 1.55682100 |
| H | -0.19196100 | -6.90157200 | 2.69130600 |
| H | -0.06840300 | -6.60192300 | -0.36318100 |
| H | 0.87101000 | -7.71591200 | 0.66229700 |
| N | 2.13299300 | -5.32930000 | 3.49929400 |
| C | 2.77692300 | -5.28758900 | 4.78294300 |
| H | 1.99685400 | -6.22246400 | 3.04450300 |
| H | 2.71629700 | -4.26054200 | 5.16394300 |
| C | 7.91108600 | 3.22121700 | 3.74087000 |
| C | 6.78845600 | 3.62264800 | 2.79978600 |
| O | 5.62227300 | 3.67524100 | 3.15839400 |
| H | 8.82827300 | 2.92305600 | 3.21155400 |
| N | 7.16943100 | 3.98304900 | 1.53603100 |
| C | 6.15517400 | 4.33131200 | 0.58828800 |
| C | 5.30806800 | 3.10682600 | 0.13900300 |
| O | 5.59821100 | 1.94569500 | 0.42965400 |
| H | 8.08385300 | 3.73051500 | 1.15094700 |
| H | 5.46653400 | 5.07889700 | 1.01197300 |
| N | 4.23266600 | 3.40727900 | -0.62169600 |
| C | 3.44161800 | 2.42709900 | -1.34899200 |
| C | 4.14715500 | 2.10638200 | -2.70401100 |
| O | 4.81026300 | 2.97081500 | -3.25175400 |
| C | 2.03794400 | 3.01711900 | -1.57877600 |
| C | 1.16915800 | 2.29419800 | -2.59101500 |
| O | 1.37798700 | 3.20844200 | -0.29179900 |
| H | 4.15278000 | 4.36295200 | -0.95193800 |
| H | 3.33965600 | 1.51854200 | -0.74109700 |
| H | 2.18187500 | 4.05116900 | -1.93397400 |
| H | 0.62537900 | 4.02175800 | -0.44499100 |
| H | 0.21893600 | 2.82537200 | -2.70223500 |
| H | 0.97732400 | 1.25561800 | -2.29623800 |
| H | 1.65254300 | 2.31052600 | -3.57790200 |
| N | 3.96810900 | 0.85138300 | -3.20415900 |
| C | 4.49035200 | 0.42083300 | -4.50750200 |
| H | 3.33704200 | 0.23011400 | -2.70742400 |
| H | 4.94346700 | 1.29650500 | -4.98891800 |
| C | -5.32254200 | -1.84749800 | 4.46268100 |
| O | -6.29120600 | -1.85987100 | 5.50325100 |
| C | -4.17020900 | -0.85377600 | 4.69474800 |
| O | -3.14400700 | -0.94476200 | 3.68086400 |
| C | -4.66166400 | 0.60083200 | 4.68769500 |
| O | -4.46506800 | 1.16646500 | 5.95928700 |
| C | -3.83870900 | 1.25460300 | 3.55487400 |
| O | -3.61980100 | 2.62435800 | 3.65798800 |
| C | -2.57566400 | 0.37864400 | 3.52646000 |
| N | 0.37722300 | 1.20193500 | 1.19405600 |
| C | 0.16020100 | 0.30605000 | 0.18950600 |
| O | 1.03172300 | -0.00846600 | -0.63863500 |
| N | -1.12651200 | -0.21436900 | 0.08955200 |
| C | -2.11102800 | -0.07918900 | 1.04205300 |
| O | -3.29257400 | -0.39080200 | 0.74689900 |
| C | -1.68488500 | 0.46350300 | 2.30174100 |
| C | -0.50275200 | 1.19096500 | 2.23885400 |
| H | 1.07798700 | 2.39054700 | 0.32699000 |
| H | -1.45352700 | -0.44603100 | -0.85383600 |
| H | -0.23136400 | 1.82856000 | 3.08787400 |
| H | -1.96783100 | 0.63266400 | 4.41610100 |
| H | -4.34048500 | 2.99459900 | 4.26235900 |
| H | -5.73048300 | 0.64711900 | 4.42366400 |
| H | -3.72752500 | -1.04472000 | 5.68796800 |
| H | -5.77906600 | -1.61532600 | 3.47551200 |
| H | -4.36435000 | 0.99650900 | 2.62391400 |
| H | -4.91686500 | -2.87238700 | 4.39530800 |
| P | -5.58237600 | 2.33525600 | 6.39582200 |
| O | -5.67388900 | 3.25564800 | 5.17245500 |
| O | -5.30341400 | 2.77500200 | 7.78272400 |
| O | -6.95427600 | 1.35959700 | 6.31311800 |
| C | 6.20990200 | -2.25624000 | 4.65500100 |
| C | 5.07376500 | -2.74511100 | 3.74960200 |
| C | 5.26762600 | -2.63039900 | 2.23594300 |
| C | 3.65166800 | -0.30720400 | 1.71894700 |
| S | 5.38176600 | -0.88941800 | 1.64084700 |
| C | 5.58959100 | -1.08524500 | -0.17709000 |
| C | 6.96580400 | -1.55746200 | -0.68726500 |
| O | 7.37175500 | -0.69151500 | -1.75855900 |
| C | 8.17473800 | -1.57313300 | 0.27595900 |
| O | 8.24963800 | -2.69806900 | 1.10056900 |
| C | 9.33615600 | -1.49389200 | -0.73139800 |
| O | 9.49010900 | -2.81041100 | -1.22434300 |
| C | 8.77641200 | -0.52914400 | -1.79726200 |
| C | 10.27146900 | 1.51471000 | -1.86556400 |
| C | 10.27696200 | 2.69376000 | -1.15532200 |
| N | 9.16586700 | 2.76063400 | -0.34687400 |
| C | 8.49269800 | 1.64584200 | -0.55380700 |
| N | 9.11897200 | 0.85180900 | -1.47724400 |
| H | 6.35136300 | -1.16515300 | 4.58233400 |
| H | 4.11091500 | -2.29482500 | 4.03360700 |
| H | 4.92534000 | -3.82427100 | 3.90985400 |
| H | 6.23060900 | -3.05295300 | 1.91596700 |
| H | 4.44688600 | -3.11085800 | 1.68209700 |
| H | 3.15548500 | -0.84850600 | 2.53581800 |
| H | 3.69435700 | 0.77174400 | 1.90439700 |
| H | 3.12768700 | -0.52225200 | 0.77621600 |
| H | 7.53155800 | 1.40807500 | -0.10105500 |
| H | 9.16198000 | -0.75892900 | -2.80304900 |
| H | 10.35556200 | -2.91098800 | -1.63945000 |
| H | 8.16953500 | -0.67952900 | 0.91694200 |
| H | 6.85651200 | -2.58276600 | -1.07958000 |
| H | 4.78519900 | -1.75986100 | -0.50075200 |
| H | 8.62751300 | -3.40771800 | 0.55248800 |
| H | 10.26466100 | -1.10773400 | -0.28186200 |
| H | 5.40210000 | -0.06514300 | -0.54071400 |
| C | -7.38224600 | 1.54244300 | -1.23025600 |
| C | -6.80021300 | 0.66276600 | -0.12867000 |
| C | -6.60238800 | 1.37824400 | 1.21157600 |
| C | -5.69680200 | 2.61292500 | 1.19552000 |
| N | -4.41725300 | 2.31588000 | 0.55375100 |
| C | -3.45072600 | 3.21515800 | 0.33651500 |
| N | -3.53994700 | 4.44832900 | 0.86273000 |
| N | -2.42246500 | 2.89501600 | -0.45235300 |
| H | -6.68991200 | 2.35112500 | -1.51453400 |
| H | -5.84853000 | 0.22998000 | -0.47657500 |
| H | -7.46243300 | -0.20249600 | 0.04179700 |
| H | -6.21039200 | 0.65663600 | 1.94512000 |
| H | -7.57863800 | 1.70602800 | 1.60600000 |
| H | -5.54388400 | 2.94686000 | 2.23481100 |
| H | -6.17407700 | 3.43937500 | 0.64287800 |
| H | -4.15620700 | 1.33241800 | 0.43670100 |

| | | | |
|---|---|---|---|
| H | -3.97312200 | 4.52626900 | 1.77867300 |
| H | -2.85969900 | 5.16523400 | 0.58914200 |
| H | -2.47993400 | 2.06756300 | -1.07228300 |
| H | -1.65717100 | 3.56771600 | -0.54448000 |
| H | 1.36202400 | 9.11801900 | -1.42692900 |
| H | -0.32183600 | 8.80073000 | -1.93338700 |
| H | -3.89046300 | 8.09142700 | -4.03489500 |
| H | -5.04161700 | 8.17996600 | -5.39063000 |
| H | 3.17253700 | -7.72079700 | 0.01477900 |
| H | 3.83965000 | -5.58205600 | 4.73127400 |
| H | 11.01156700 | 3.49625500 | -1.19385600 |
| H | 8.14613700 | 4.07449700 | 4.39671200 |
| H | 7.56006900 | 2.39936300 | 4.37972200 |
| H | 10.95578100 | 1.10306600 | -2.60324100 |
| H | 6.62847600 | 4.75151000 | -0.31294600 |
| H | 7.16794600 | -2.73305100 | 4.39175400 |
| H | 4.10971000 | -6.59686100 | -1.02074800 |
| H | 2.27214500 | -5.95693800 | 5.49821700 |
| H | -6.50850400 | -0.94188400 | 5.74916000 |
| H | -7.58121200 | 1.81358600 | 5.73369500 |
| H | -8.32870100 | 2.01380300 | -0.91739200 |
| H | -4.68088900 | -5.64126400 | -5.75251800 |
| H | 5.98454800 | -2.48896400 | 5.70603400 |
| H | 3.67764900 | 0.02993100 | -5.13832400 |
| H | -1.28303600 | -2.65533300 | -6.20279700 |
| H | -7.60155100 | 0.95210400 | -2.13616300 |
| H | 5.26175100 | -0.35559300 | -4.37916500 |
| H | -4.68626900 | -7.82501100 | -1.54203900 |

#### Model4 – Intermediate

| | | | |
|---|---|---|---|
| O | -1.93954500 | 4.20372100 | -3.71369000 |
| H | -1.79515600 | 4.23437600 | -2.75002600 |
| H | -2.47832300 | 4.98723500 | -3.88995500 |
| O | -2.45252300 | 0.92892000 | -1.88891800 |
| H | -3.27250500 | 0.47488500 | -2.18911800 |
| H | -2.08397400 | 1.19916200 | -2.77161000 |
| O | -4.51720200 | 0.00358000 | -3.44035000 |
| H | -5.37940000 | 0.42721800 | -3.32897900 |
| H | -4.73028700 | -0.94931400 | -3.62757000 |
| O | -2.07084100 | 1.53968400 | -4.44307200 |
| H | -2.13844200 | 2.51704200 | -4.39986400 |
| H | -2.97672500 | 1.19641400 | -4.53773100 |
| C | -5.28413100 | 7.46326100 | -4.26565800 |
| C | -5.34709000 | 5.93098600 | -4.23837900 |
| C | -5.15910800 | 5.27173100 | -2.85712300 |
| C | -6.19927600 | 5.73630800 | -1.83278400 |
| C | -5.17684100 | 3.74207300 | -2.98055100 |
| H | -6.12189900 | 7.91616500 | -3.71346300 |
| H | -4.57593500 | 5.52678400 | -4.92151900 |
| H | -6.31207000 | 5.59441500 | -4.66110700 |
| H | -4.16893200 | 5.57370700 | -2.47001500 |
| H | -6.02943200 | 5.25250800 | -0.85731200 |
| H | -7.22130500 | 5.47945000 | -2.16208500 |
| H | -6.16077200 | 6.82402400 | -1.67119400 |
| H | -5.03074700 | 3.26389200 | -2.00004100 |
| H | -6.14288600 | 3.39323900 | -3.38398500 |
| H | -4.37593400 | 3.37420500 | -3.63775800 |
| C | 0.00430000 | 8.49016800 | -1.05185300 |
| C | 0.21412300 | 6.97467000 | -1.03465500 |
| C | -1.07073800 | 6.18709600 | -1.21383400 |
| O | -2.14063600 | 6.68404400 | -1.47132100 |
| O | -0.97680300 | 4.84739400 | -1.08738600 |
| H | -0.65658300 | 8.80338800 | -0.23051600 |
| H | 0.69402800 | 6.63402600 | -0.10074500 |
| H | 0.89744800 | 6.65767400 | -1.84414900 |
| C | -3.61186900 | -7.41124800 | -1.30041900 |
| C | -3.80290400 | -6.82731200 | 0.09984200 |
| C | -4.74945300 | -5.62314700 | 0.13467500 |
| N | -4.27444500 | -4.47969100 | -0.62456400 |
| C | -3.98490700 | -3.27006300 | -0.12800100 |
| N | -3.76759000 | -3.06145000 | 1.18451700 |
| N | -3.89534200 | -2.24046500 | -0.96938600 |
| H | -3.24012600 | -6.65648800 | -2.00979300 |
| H | -2.90416300 | -8.25497500 | -1.28635300 |
| H | -2.82921200 | -6.52888800 | 0.52511400 |
| H | -4.21193000 | -7.59823300 | 0.77588600 |
| H | -4.94420800 | -5.31795500 | 1.17418000 |
| H | -5.72832100 | -5.92544900 | -0.27897500 |
| H | -4.29957400 | -4.54820300 | -1.67292900 |
| H | -3.69842600 | -3.82479500 | 1.83988600 |
| H | -3.60590700 | -2.10738400 | 1.52359200 |
| H | -4.27208800 | -2.33945300 | -1.93192400 |
| H | -3.57341900 | -1.34934800 | -0.58132500 |
| C | -4.77253400 | -4.98072000 | -6.25046000 |
| C | -3.62562200 | -4.08652800 | -6.72231800 |
| O | -3.61263500 | -3.56309100 | -7.82574100 |
| C | -5.65834100 | -4.15954400 | -5.28087200 |
| C | -4.97139700 | -3.78979000 | -3.95430900 |
| O | -4.34599000 | -4.70697700 | -3.35428000 |
| O | -5.07818000 | -2.60777700 | -3.51897400 |
| H | -5.34439400 | -5.28608700 | -7.13561000 |
| H | -6.55027300 | -4.75301500 | -5.01450900 |
| H | -6.00941900 | -3.24001900 | -5.77111500 |
| N | -2.67189500 | -3.81995900 | -5.77668900 |
| C | -1.78591600 | -2.67723600 | -5.93183000 |
| C | -0.79990200 | -2.50307900 | -4.77031700 |
| C | -1.51528500 | -2.31054200 | -3.42185000 |
| C | 0.13544400 | -1.32690200 | -5.08227800 |
| C | -0.58544100 | -2.22532300 | -2.21269100 |
| H | -2.80550700 | -4.21651500 | -4.85074100 |
| H | -2.38586100 | -1.75252200 | -6.03759500 |
| H | -0.19474500 | -3.42910400 | -4.70603400 |
| H | -2.22536900 | -3.13398600 | -3.25031800 |
| H | -2.12411000 | -1.39353300 | -3.47015600 |
| H | 0.62115100 | -1.45314400 | -6.06324100 |
| H | 0.93127100 | -1.23111300 | -4.32694500 |
| H | -0.42684100 | -0.37831700 | -5.09523500 |
| H | -1.17313300 | -2.19327600 | -1.28436000 |
| H | 0.04847100 | -1.32515500 | -2.22275400 |
| H | 0.08256100 | -3.10406700 | -2.16230000 |
| C | 3.32923300 | -6.78526200 | -0.59255900 |
| C | 3.22000400 | -5.48314600 | 0.19402700 |
| O | 4.19507300 | -4.76428900 | 0.39235800 |
| H | 2.71330400 | -6.74758400 | -1.50523000 |
| N | 1.98665300 | -5.16158600 | 0.70135900 |
| C | 1.78582300 | -3.89130000 | 1.41280300 |
| C | 2.15854200 | -3.91384700 | 2.90538600 |
| O | 2.10131700 | -2.87157000 | 3.54448500 |
| C | 0.29141800 | -3.58100200 | 1.21460800 |
| C | -0.33981800 | -4.97091700 | 1.08125800 |
| C | 0.70652900 | -5.77008900 | 0.29438800 |
| H | 2.41894200 | -3.10967300 | 0.97054600 |
| H | 0.15674500 | -3.01079300 | 0.28605300 |
| H | -0.11691700 | -2.97835200 | 2.03669400 |
| H | -1.31135300 | -4.95015200 | 0.56949800 |
| H | -0.48975000 | -5.42084800 | 2.07609400 |
| H | 0.55523400 | -5.65189500 | -0.79307200 |
| H | 0.68161200 | -6.84707900 | 0.51996500 |
| N | 2.52534500 | -5.11102400 | 3.42627300 |
| C | 2.93612400 | -5.30900000 | 4.80569100 |
| H | 2.53932700 | -5.89661700 | 2.78707600 |
| H | 2.79887800 | -4.35874700 | 5.33759000 |
| C | 7.71065700 | 3.36731700 | 3.85039900 |
| C | 6.55471900 | 3.65349600 | 2.90538200 |
| O | 5.38722000 | 3.62045200 | 3.26545800 |
| H | 8.62397500 | 3.05900700 | 3.31979100 |
| N | 6.91419500 | 4.03115200 | 1.64287700 |
| C | 5.90360900 | 4.42187700 | 0.70762900 |
| C | 5.10575700 | 3.25333100 | 0.09715400 |
| O | 5.38509700 | 2.06833900 | 0.28443200 |

| | | | |
|---|---|---|---|
| H | 7.85190600 | 3.84484200 | 1.27100000 |
| H | 5.18114700 | 5.09604800 | 1.19552800 |
| N | 4.08771400 | 3.64089800 | -0.69571600 |
| C | 3.32881800 | 2.72771000 | -1.52562100 |
| C | 4.15488100 | 2.28364800 | -2.74210600 |
| O | 4.94620300 | 3.03283100 | -3.28788700 |
| C | 2.01162100 | 3.41731600 | -1.95023900 |
| C | 1.21891300 | 2.70075700 | -3.03753200 |
| O | 1.22347700 | 3.59372500 | -0.77307600 |
| H | 3.96958400 | 4.63221100 | -0.86855300 |
| H | 3.05098000 | 1.85350400 | -0.92085700 |
| H | 2.26996700 | 4.43025600 | -2.30992800 |
| H | -0.05036200 | 4.44144600 | -0.92888400 |
| H | 0.31723700 | 3.28207800 | -3.27463600 |
| H | 0.90579000 | 1.70449500 | -2.69330400 |
| H | 1.79719700 | 2.60005800 | -3.96634000 |
| N | 3.88402800 | 1.01540500 | -3.16048300 |
| C | 4.39194400 | 0.47657900 | -4.40801800 |
| H | 3.17371500 | 0.49611500 | -2.65666200 |
| H | 5.11193000 | -0.34013900 | -4.23031200 |
| C | -5.24620700 | -2.27882200 | 4.57616400 |
| O | -6.26445000 | -2.25020000 | 5.53937800 |
| C | -4.14942500 | -1.27760700 | 4.89832000 |
| O | -3.08129000 | -1.37631100 | 3.94734100 |
| C | -4.67879000 | 0.17067600 | 4.84665300 |
| O | -4.58144900 | 0.78274200 | 6.11123900 |
| C | -3.80160800 | 0.80129900 | 3.74656800 |
| O | -3.62954200 | 2.18048300 | 3.80229100 |
| C | -2.52264000 | -0.05647100 | 3.82829000 |
| N | 0.73299000 | 0.51803300 | 1.97427400 |
| C | 0.32498000 | 0.60286100 | 0.69681900 |
| O | 1.06843200 | 0.99195100 | -0.24947700 |
| N | -0.98255800 | 0.25740000 | 0.39777900 |
| C | -1.96220600 | -0.02479600 | 1.32464700 |
| O | -3.12098300 | -0.31565400 | 0.89673400 |
| C | -1.55490200 | 0.04338900 | 2.68777500 |
| C | -0.19638500 | 0.26751500 | 2.90584400 |
| H | 1.08084200 | 2.66680800 | -0.41666600 |
| H | -1.31481300 | 0.34884600 | -0.57549900 |
| H | 0.16468100 | 0.25441000 | 3.94185100 |
| H | -1.98902300 | 0.20887300 | 4.76147300 |
| H | -4.36732900 | 2.54630200 | 4.38203800 |
| H | -5.73114600 | 0.18204100 | 4.51712000 |
| H | -3.77015800 | -1.45642000 | 5.92012000 |
| H | -5.62077700 | -2.06498000 | 3.54999200 |
| H | -4.26676600 | 0.49440200 | 2.79792500 |
| H | -4.83383900 | -3.30301900 | 4.56000800 |
| P | -5.72851700 | 1.96526200 | 6.45569800 |
| O | -5.76458200 | 2.84211400 | 5.19192300 |
| O | -5.51279800 | 2.46552800 | 7.83222100 |
| O | -7.08505300 | 0.98051800 | 6.33963500 |
| C | 6.23892500 | -2.13617500 | 4.68979900 |
| C | 5.13550400 | -2.59324900 | 3.74384400 |
| C | 5.39852900 | -2.42057500 | 2.25280800 |
| C | 3.75655000 | -0.06386700 | 1.89552200 |
| S | 5.47786500 | -0.66967700 | 1.67864300 |
| C | 5.55678000 | -0.92600800 | -0.14059700 |
| C | 6.91100800 | -1.37109100 | -0.72645900 |
| O | 7.31588600 | -0.41226900 | -1.71558300 |
| C | 8.14656100 | -1.51683400 | 0.19511900 |
| O | 8.22482400 | -2.71969900 | 0.89817800 |
| C | 9.28244300 | -1.35235700 | -0.82738100 |
| O | 9.38802500 | -2.60567300 | -1.47291400 |
| C | 8.71640400 | -0.26333400 | -1.75680800 |
| C | 10.27960100 | 1.71753900 | -1.52347200 |
| C | 10.25144400 | 2.84951400 | -0.74080900 |
| N | 9.07319000 | 2.90133700 | -0.03400100 |
| C | 8.39109700 | 1.82300000 | -0.37171000 |
| N | 9.08007800 | 1.06941300 | -1.28221800 |
| H | 6.39635400 | -1.04560900 | 4.63686300 |
| H | 4.16972800 | -2.14619300 | 4.01597200 |
| H | 4.98033200 | -3.67766500 | 3.86068100 |
| H | 6.38086400 | -2.81813700 | 1.96062100 |
| H | 4.64061100 | -2.92647000 | 1.63964500 |
| H | 3.19471900 | -0.79471400 | 2.49154000 |
| H | 3.82654600 | 0.90742200 | 2.39767500 |
| H | 3.28420200 | 0.06829900 | 0.91516700 |
| H | 7.38392800 | 1.58837300 | -0.03007100 |
| H | 9.08176500 | -0.37501000 | -2.79042400 |
| H | 10.23063500 | -2.66936000 | -1.93895800 |
| H | 8.18018400 | -0.69317900 | 0.92363500 |
| H | 6.76188800 | -2.34657100 | -1.21954600 |
| H | 4.76597600 | -1.65154900 | -0.37250900 |
| H | 8.53012400 | -3.38868900 | 0.26177600 |
| H | 10.23157800 | -1.04218000 | -0.36198900 |
| H | 5.29639500 | 0.06374500 | -0.54126400 |
| C | -7.51364900 | 1.12621300 | -1.14494900 |
| C | -6.81893400 | 0.08983500 | -0.26085400 |
| C | -6.73883700 | 0.46570400 | 1.22645200 |
| C | -6.09288000 | 1.81979900 | 1.52260100 |
| N | -4.79451300 | 1.89871200 | 0.86240900 |
| C | -4.06977500 | 3.01026700 | 0.76508900 |
| N | -4.46410700 | 4.11830900 | 1.43129100 |
| N | -3.01197800 | 3.02804400 | -0.04533900 |
| H | -6.95211700 | 2.07189200 | -1.18402300 |
| H | -5.80758000 | -0.10033700 | -0.65598200 |
| H | -7.34381700 | -0.87663800 | -0.34197400 |
| H | -6.18459300 | -0.32071000 | 1.76597100 |
| H | -7.75048300 | 0.48148200 | 1.66499100 |
| H | -6.00014900 | 1.96896700 | 2.61312000 |
| H | -6.71992000 | 2.64128900 | 1.13983300 |
| H | -4.27568800 | 1.01944900 | 0.70918900 |
| H | -4.83420800 | 3.96114400 | 2.36829800 |
| H | -3.91134700 | 4.95969700 | 1.31958100 |
| H | -2.85560200 | 2.26174900 | -0.72287200 |
| H | -2.38532100 | 3.82651400 | -0.10043900 |
| H | 0.96542000 | 9.01379900 | -0.94417100 |
| H | -0.46661900 | 8.80773200 | -1.99304200 |
| H | -4.35011900 | 7.82940100 | -3.80686300 |
| H | -5.32662100 | 7.84148200 | -5.29898400 |
| H | 2.97650500 | -7.64074400 | 0.00419000 |
| H | 3.99460800 | -5.61035100 | 4.87276300 |
| H | 11.00785500 | 3.62831300 | -0.66032700 |
| H | 7.93300100 | 4.28287000 | 4.42161400 |
| H | 7.40656600 | 2.59111200 | 4.56534700 |
| H | 11.01747300 | 1.33103300 | -2.22209800 |
| H | 6.37682500 | 4.96252300 | -0.12772700 |
| H | 7.20127600 | -2.62069300 | 4.45655500 |
| H | 4.37964900 | -6.93291300 | -0.86778000 |
| H | 2.31971100 | -6.08028000 | 5.29401100 |
| H | -6.49515200 | -1.32941500 | 5.75284000 |
| H | -7.70008000 | 1.41884600 | 5.73622800 |
| H | -8.52895200 | 1.35532600 | -0.78065800 |
| H | -4.39904700 | -5.87196800 | -5.72295900 |
| H | 5.97827200 | -2.37643700 | 5.73118100 |
| H | 4.91373700 | 1.28674800 | -4.93396500 |
| H | -1.23443800 | -2.78491400 | -6.87933900 |
| H | -7.61525900 | 0.75863700 | -2.17969700 |
| H | 3.56938200 | 0.10080300 | -5.03703000 |
| H | -4.56807500 | -7.78127900 | -1.70476300 |

#### Model4 − TS2

| | | | |
|---|---|---|---|
| O | -1.91993600 | 4.09640300 | -3.92374000 |
| H | -1.79377300 | 4.21053400 | -2.96200200 |
| H | -2.29212100 | 4.94458700 | -4.20127500 |
| O | -2.64463400 | 1.42824800 | -1.52971100 |
| H | -3.39653900 | 0.87545400 | -1.82017600 |
| H | -2.18326200 | 1.54070500 | -2.40717500 |
| O | -4.34728400 | 0.02569600 | -3.39124200 |
| H | -5.17042100 | 0.51063400 | -3.54189200 |

| | | | |
|---|---|---|---|
| H | -4.57394300 | -0.91629200 | -3.60825700 |
| O | -1.87931700 | 1.38289400 | -4.06956400 |
| H | -1.98249700 | 2.32684700 | -4.31451600 |
| H | -2.74677500 | 0.95599100 | -4.20568100 |
| C | -5.16182400 | 7.43282200 | -4.55825000 |
| C | -5.25926400 | 5.90858100 | -4.40972600 |
| C | -5.11439300 | 5.36455300 | -2.97447300 |
| C | -6.18440600 | 5.91611000 | -2.02701100 |
| C | -5.13353500 | 3.82985000 | -2.96657000 |
| H | -5.99756800 | 7.94665700 | -4.05837900 |
| H | -4.47838700 | 5.43618800 | -5.03443800 |
| H | -6.22074300 | 5.55656100 | -4.82899000 |
| H | -4.13055600 | 5.69219900 | -2.59597900 |
| H | -6.06014400 | 5.50244200 | -1.01271600 |
| H | -7.19773400 | 5.64740200 | -2.37414100 |
| H | -6.13788400 | 7.01240000 | -1.94292200 |
| H | -4.99924900 | 3.43721200 | -1.94560900 |
| H | -6.09559300 | 3.44644000 | -3.34932500 |
| H | -4.32098800 | 3.41425400 | -3.57991900 |
| C | 0.07559800 | 8.48504000 | -1.26996600 |
| C | 0.22408200 | 6.97083200 | -1.10484200 |
| C | -0.98684700 | 6.19151900 | -1.59116300 |
| O | -1.92513400 | 6.69955800 | -2.15858200 |
| O | -0.98444700 | 4.86461600 | -1.36432300 |
| H | -0.78177000 | 8.85908600 | -0.69163100 |
| H | 0.41184300 | 6.68925500 | -0.05431900 |
| H | 1.10004400 | 6.59187700 | -1.66273800 |
| C | -3.63025400 | -7.39759400 | -1.30966500 |
| C | -3.85679100 | -6.78712300 | 0.07380600 |
| C | -4.76779800 | -5.55591900 | 0.06065600 |
| N | -4.22795900 | -4.42377200 | -0.67669200 |
| C | -3.97296600 | -3.21354600 | -0.16519400 |
| N | -3.81525900 | -3.00612400 | 1.15637000 |
| N | -3.86610200 | -2.17809700 | -0.99993600 |
| H | -3.21085300 | -6.66451000 | -2.01532000 |
| H | -2.94796800 | -8.26029800 | -1.25616900 |
| H | -2.89190100 | -6.51341500 | 0.53388400 |
| H | -4.31552000 | -7.53581400 | 0.74282100 |
| H | -5.00174400 | -5.24222100 | 1.08932500 |
| H | -5.73396100 | -5.83029300 | -0.39946100 |
| H | -4.22645500 | -4.49302300 | -1.72790200 |
| H | -3.71845100 | -3.77581600 | 1.80147700 |
| H | -3.64724200 | -2.05668800 | 1.50733200 |
| H | -4.21266500 | -2.27914800 | -1.97374200 |
| H | -3.58341100 | -1.28127400 | -0.60620600 |
| C | -4.68967100 | -5.04622400 | -6.32029100 |
| C | -3.49775200 | -4.21155100 | -6.79245400 |
| O | -3.43726200 | -3.72965800 | -7.91300600 |
| C | -5.54164300 | -4.16328100 | -5.36605000 |
| C | -4.86889100 | -3.78275900 | -4.02954900 |
| O | -4.26526000 | -4.69908600 | -3.40442000 |
| O | -4.97677300 | -2.59564900 | -3.60662600 |
| H | -5.27117800 | -5.33362900 | -7.20472200 |
| H | -6.46346500 | -4.70876300 | -5.09765500 |
| H | -5.84897500 | -3.24094200 | -5.88011900 |
| N | -2.56076300 | -3.94031100 | -5.83079800 |
| C | -1.63181700 | -2.83508400 | -6.00805000 |
| C | -0.69478600 | -2.61071300 | -4.81411300 |
| C | -1.45932900 | -2.33767900 | -3.50600500 |
| C | 0.26739100 | -1.46200300 | -5.14798300 |
| C | -0.56685600 | -2.11545100 | -2.28462600 |
| H | -2.72339800 | -4.30566400 | -4.89656000 |
| H | -2.19905500 | -1.90398700 | -6.20107700 |
| H | -0.10309200 | -3.53732800 | -4.67543200 |
| H | -2.14344500 | -3.17271300 | -3.28978100 |
| H | -2.09955000 | -1.45193100 | -3.64882600 |
| H | 0.74953900 | -1.61625600 | -6.12649600 |
| H | 1.06558700 | -1.37037400 | -4.39515400 |
| H | -0.27304700 | -0.50114700 | -5.17194400 |
| H | -1.17642800 | -2.05257600 | -1.37141100 |
| H | 0.01612100 | -1.18388400 | -2.34851700 |
| H | 0.14811400 | -2.94804700 | -2.15696600 |
| C | 3.30098000 | -6.79877700 | -0.49065300 |
| C | 3.17171100 | -5.48636000 | 0.26814500 |
| O | 4.13498600 | -4.75432900 | 0.45798500 |
| H | 2.70368800 | -6.77981000 | -1.41639600 |
| N | 1.92859600 | -5.17111700 | 0.76256900 |
| C | 1.70772700 | -3.88680300 | 1.44044500 |
| C | 2.03346900 | -3.88102000 | 2.94469200 |
| O | 1.91993600 | -2.83391300 | 3.57267700 |
| C | 0.22098800 | -3.58122500 | 1.18738400 |
| C | -0.40728000 | -4.97433700 | 1.07336400 |
| C | 0.66155500 | -5.79494600 | 0.34100800 |
| H | 2.36074200 | -3.11929000 | 1.00243900 |
| H | 0.11498500 | -3.03706100 | 0.23907300 |
| H | -0.21347700 | -2.95989900 | 1.98118700 |
| H | -1.36272500 | -4.96910900 | 0.53166700 |
| H | -0.58694600 | -5.39577200 | 2.07579100 |
| H | 0.53855700 | -5.71239900 | -0.75329400 |
| H | 0.63091600 | -6.86439800 | 0.59984200 |
| N | 2.42816300 | -5.05647400 | 3.48632500 |
| C | 2.82229100 | -5.22747200 | 4.87379500 |
| H | 2.48985100 | -5.84434000 | 2.85248000 |
| H | 2.67481600 | -4.26892900 | 5.38721600 |
| C | 7.66439000 | 3.40393700 | 3.85417900 |
| C | 6.52951500 | 3.76777900 | 2.91088300 |
| O | 5.37573700 | 3.89102500 | 3.29306100 |
| H | 8.57460000 | 3.08285100 | 3.32662800 |
| N | 6.89939400 | 4.01263200 | 1.62051300 |
| C | 5.91884200 | 4.41439100 | 0.66242800 |
| C | 5.08721200 | 3.25808300 | 0.07122400 |
| O | 5.27794800 | 2.07777100 | 0.35130300 |
| H | 7.82979000 | 3.78171300 | 1.25722400 |
| H | 5.21186800 | 5.12069300 | 1.12702300 |
| N | 4.13027800 | 3.65490300 | -0.80073100 |
| C | 3.35631500 | 2.72985100 | -1.60472500 |
| C | 4.16274400 | 2.26827400 | -2.83026800 |
| O | 4.89720100 | 3.03519900 | -3.43018100 |
| C | 2.03035000 | 3.40356500 | -2.03226200 |
| C | 1.26150100 | 2.67997000 | -3.13144100 |
| O | 1.21943000 | 3.57471700 | -0.86701800 |
| H | 4.13262500 | 4.62433600 | -1.09653700 |
| H | 3.10016500 | 1.86638700 | -0.97622700 |
| H | 2.27735200 | 4.42062900 | -2.38621100 |
| H | -0.10044800 | 4.45559900 | -1.09329500 |
| H | 0.34884100 | 3.24159500 | -3.37190200 |
| H | 0.96527300 | 1.67186300 | -2.80789300 |
| H | 1.85305800 | 2.60126400 | -4.05409800 |
| N | 3.94087600 | 0.97519500 | -3.19344700 |
| C | 4.47350300 | 0.39013100 | -4.41064400 |
| H | 3.29825900 | 0.43098800 | -2.62977400 |
| H | 5.24298800 | -0.36773300 | -4.18597800 |
| C | -5.28910500 | -2.17275600 | 4.50586600 |
| O | -6.37128200 | -2.10453200 | 5.39293900 |
| C | -4.15495700 | -1.23080900 | 4.88571300 |
| O | -3.09672900 | -1.40996800 | 3.94086900 |
| C | -4.55778600 | 0.25931900 | 4.82592000 |
| O | -4.78050000 | 0.79669100 | 6.11128400 |
| C | -3.38583600 | 0.90887300 | 4.01779700 |
| O | -2.88305400 | 2.09835800 | 4.53424600 |
| C | -2.31927100 | -0.21000800 | 3.93971900 |
| N | 0.84212500 | 0.15671700 | 1.84988500 |
| C | 0.34725900 | 0.48036600 | 0.63098200 |
| O | 1.05229400 | 0.91036100 | -0.30979500 |
| N | -1.01202900 | 0.31192100 | 0.44028500 |
| C | -1.92781700 | -0.04025600 | 1.41003500 |
| O | -3.11831200 | -0.24933000 | 1.05629900 |
| C | -1.40370700 | -0.14110400 | 2.74407900 |
| C | -0.01992700 | -0.09957900 | 2.85625800 |
| H | 1.05276300 | 2.65828900 | -0.52818600 |

| | | | |
|---|---|---|---|
| H | -1.43185100 | 0.58146400 | -0.46790800 |
| H | 0.43922200 | -0.28505400 | 3.83302700 |
| H | -1.70672400 | -0.15084700 | 4.86050000 |
| H | -3.67256200 | 2.60940900 | 4.83116900 |
| H | -5.47912300 | 0.35188100 | 4.22551900 |
| H | -3.80273300 | -1.44298100 | 5.91229700 |
| H | -5.58154200 | -1.94013700 | 3.45826700 |
| H | -3.76443700 | 1.03601700 | 2.99495700 |
| H | -4.92360400 | -3.21386700 | 4.51368900 |
| P | -5.82634300 | 2.12296000 | 6.24616900 |
| O | -5.44829200 | 2.98220300 | 5.02699400 |
| O | -5.86420900 | 2.59565100 | 7.65073300 |
| O | -7.25283000 | 1.36820800 | 5.83515500 |
| C | 6.14543400 | -2.07584700 | 4.76170700 |
| C | 5.05511300 | -2.58480500 | 3.82421800 |
| C | 5.33133800 | -2.43425200 | 2.33051900 |
| C | 3.04137500 | -0.18733400 | 1.83689200 |
| S | 5.34081400 | -0.68679100 | 1.74963800 |
| C | 5.56296000 | -0.93898500 | -0.05607900 |
| C | 6.94497900 | -1.40477000 | -0.57726900 |
| O | 7.35116600 | -0.52975700 | -1.65243900 |
| C | 8.16467400 | -1.43502900 | 0.37183400 |
| O | 8.27515100 | -2.58193400 | 1.16190000 |
| C | 9.32237200 | -1.31139600 | -0.63638100 |
| O | 9.49598000 | -2.60838100 | -1.17566700 |
| C | 8.74633900 | -0.32838000 | -1.67142600 |
| C | 10.21520700 | 1.73480000 | -1.58357900 |
| C | 10.15415900 | 2.89835800 | -0.85067600 |
| N | 8.98683800 | 2.93311000 | -0.12530200 |
| C | 8.34345200 | 1.81505800 | -0.40498400 |
| N | 9.04728500 | 1.05116300 | -1.29413700 |
| H | 6.28136300 | -0.98553700 | 4.67078600 |
| H | 4.08370100 | -2.13886300 | 4.08211100 |
| H | 4.92215900 | -3.66744400 | 3.98016800 |
| H | 6.32500600 | -2.82618100 | 2.07390900 |
| H | 4.59650400 | -2.98408400 | 1.72893800 |
| H | 2.80624900 | -1.04553600 | 2.46466700 |
| H | 3.20927700 | 0.79223000 | 2.27749900 |
| H | 2.91958400 | -0.23777900 | 0.75835700 |
| H | 7.35862000 | 1.54801000 | -0.02402400 |
| H | 9.14721400 | -0.50633800 | -2.68225600 |
| H | 10.36012300 | -2.67530100 | -1.59939500 |
| H | 8.14703500 | -0.56461800 | 1.04305900 |
| H | 6.83321100 | -2.42366300 | -0.98525300 |
| H | 4.78795600 | -1.64952400 | -0.37558400 |
| H | 8.61798300 | -3.27891200 | 0.57739200 |
| H | 10.24723700 | -0.92811600 | -0.17627300 |
| H | 5.34259700 | 0.05220700 | -0.47494200 |
| C | -7.48323500 | 1.16316400 | -1.36911100 |
| C | -6.69270900 | 0.09481300 | -0.61206300 |
| C | -6.79860600 | 0.16181000 | 0.91998700 |
| C | -6.37082500 | 1.48283900 | 1.56314600 |
| N | -5.02620600 | 1.82877300 | 1.11892200 |
| C | -4.35732300 | 2.93306100 | 1.47723100 |
| N | -4.80279400 | 3.71939600 | 2.46081600 |
| N | -3.25561800 | 3.23061400 | 0.76790600 |
| H | -7.10539200 | 2.17616000 | -1.16139200 |
| H | -5.63666900 | 0.14639400 | -0.92097200 |
| H | -7.02577300 | -0.90662100 | -0.93217600 |
| H | -6.18976900 | -0.65114900 | 1.35221300 |
| H | -7.83763900 | -0.03456900 | 1.23210100 |
| H | -6.42253900 | 1.39875400 | 2.66202400 |
| H | -7.05187200 | 2.30195900 | 1.27989800 |
| H | -4.42946100 | 1.06543500 | 0.78308200 |
| H | -5.26303700 | 3.33538900 | 3.32459400 |
| H | -4.28744600 | 4.57330300 | 2.64114600 |
| H | -3.03585300 | 2.71242900 | -0.09351300 |
| H | -2.58476600 | 3.90511800 | 1.11032800 |
| H | 0.98256400 | 9.00187900 | -0.92349800 |
| H | -0.09957100 | 8.74700600 | -2.32318000 |

| | | | |
|---|---|---|---|
| H | -4.22559100 | 7.81111800 | -4.11479500 |
| H | -5.18036900 | 7.72821000 | -5.61911200 |
| H | 2.94024800 | -7.64639300 | 0.11252800 |
| H | 3.88135500 | -5.52033800 | 4.95619700 |
| H | 10.87869200 | 3.71021500 | -0.81744200 |
| H | 7.90430100 | 4.28300800 | 4.47310000 |
| H | 7.32271800 | 2.60722400 | 4.52948000 |
| H | 10.95658800 | 1.34614400 | -2.27724000 |
| H | 6.42018200 | 4.92179500 | -0.17787700 |
| H | 7.11721200 | -2.54675500 | 4.54015100 |
| H | 4.35703700 | -6.94583600 | -0.74365200 |
| H | 2.20438200 | -5.99425600 | 5.36779400 |
| H | -6.61221100 | -1.17240600 | 5.52101200 |
| H | -7.76929800 | 1.28003800 | 6.64920000 |
| H | -8.55253800 | 1.14078800 | -1.10112400 |
| H | -4.36274700 | -5.94719600 | -5.77819900 |
| H | 5.89103400 | -2.29000200 | 5.81102300 |
| H | 4.94077300 | 1.19627200 | -4.99120000 |
| H | -1.03708500 | -3.01392800 | -6.91823500 |
| H | -7.41494400 | 1.00593900 | -2.45773900 |
| H | 3.67087300 | -0.07281000 | -5.00633800 |
| H | -4.58083200 | -7.74629000 | -1.74478000 |

#### Model4 − Final state

| | | | |
|---|---|---|---|
| O | -2.12632500 | 3.78430200 | -4.11689200 |
| H | -2.00927500 | 3.95649700 | -3.16323500 |
| H | -2.58068200 | 4.57542200 | -4.43730400 |
| O | -2.71210500 | 1.13785000 | -1.62781600 |
| H | -3.42971200 | 0.52904600 | -1.90027800 |
| H | -2.26267700 | 1.24625400 | -2.51682800 |
| O | -4.36929500 | -0.41758800 | -3.35662600 |
| H | -5.21570900 | 0.01504100 | -3.53574000 |
| H | -4.55075000 | -1.38125100 | -3.51513000 |
| O | -1.96993700 | 1.07492800 | -4.14831800 |
| H | -2.11158400 | 2.00385200 | -4.43154500 |
| H | -2.80680800 | 0.59676500 | -4.29589400 |
| C | -5.44882500 | 6.82495300 | -5.17762300 |
| C | -5.47803800 | 5.32345700 | -4.86305900 |
| C | -5.38788100 | 4.95023100 | -3.36989200 |
| C | -6.53577200 | 5.54491200 | -2.54795900 |
| C | -5.33592600 | 3.42687500 | -3.18886400 |
| H | -6.33657600 | 7.34257700 | -4.78269400 |
| H | -4.64238900 | 4.82829700 | -5.39248000 |
| H | -6.39584500 | 4.87458200 | -5.28726800 |
| H | -4.44469100 | 5.36954500 | -2.97791800 |
| H | -6.45150400 | 5.25286500 | -1.48828200 |
| H | -7.51345100 | 5.18659600 | -2.91556400 |
| H | -6.54510200 | 6.64453600 | -2.58546600 |
| H | -5.24907500 | 3.15884700 | -2.12322300 |
| H | -6.25392200 | 2.95289600 | -3.57776400 |
| H | -4.47027900 | 2.98651000 | -3.70452500 |
| C | -0.27634800 | 8.36343600 | -1.97932900 |
| C | -0.05676400 | 6.87646400 | -1.69911200 |
| C | -1.29381800 | 6.03029600 | -1.94156600 |
| O | -2.31362300 | 6.45232800 | -2.43230700 |
| O | -1.21209600 | 4.73000600 | -1.58966500 |
| H | -1.05577100 | 8.77574700 | -1.32192100 |
| H | 0.28600300 | 6.69744100 | -0.66556600 |
| H | 0.74482500 | 6.46495100 | -2.33952300 |
| C | -3.33353900 | -7.60466500 | -0.68852400 |
| C | -3.45763200 | -6.83375500 | 0.62563600 |
| C | -4.52595000 | -5.73735300 | 0.57669700 |
| N | -4.18340600 | -4.62612100 | -0.29531500 |
| C | -3.95022300 | -3.37359500 | 0.10146100 |
| N | -3.75025500 | -3.05262900 | 1.39545500 |
| N | -3.91696000 | -2.40887700 | -0.82088400 |
| H | -3.14179100 | -6.93386100 | -1.53998800 |
| H | -2.52120700 | -8.34651500 | -0.64097000 |
| H | -2.49125900 | -6.37418700 | 0.89879100 |
| H | -3.71627800 | -7.52725000 | 1.44449400 |

| | | | |
|---|---|---|---|
| H | -4.73740300 | -5.35509300 | 1.58668300 |
| H | -5.47453700 | -6.17298100 | 0.21657500 |
| H | -4.17548400 | -4.79498600 | -1.33593000 |
| H | -3.61839800 | -3.76393600 | 2.09845000 |
| H | -3.62588900 | -2.07639000 | 1.67313800 |
| H | -4.24865400 | -2.61315100 | -1.78618800 |
| H | -3.64658200 | -1.47506500 | -0.52674000 |
| C | -4.46049000 | -5.72821900 | -5.88146000 |
| C | -3.30730100 | -4.87349900 | -6.40780600 |
| O | -3.25800400 | -4.47391200 | -7.56043200 |
| C | -5.37590800 | -4.81041200 | -5.02305200 |
| C | -4.75583700 | -4.28345900 | -3.71201100 |
| O | -4.13400500 | -5.11215300 | -2.99055900 |
| O | -4.92425100 | -3.06833100 | -3.39901200 |
| H | -5.01034800 | -6.12259000 | -6.74460200 |
| H | -6.27584800 | -5.37675300 | -4.72537600 |
| H | -5.71555900 | -3.95266400 | -5.62152700 |
| N | -2.39780900 | -4.48268200 | -5.46144500 |
| C | -1.51904700 | -3.35185200 | -5.71360100 |
| C | -0.62730300 | -2.98632100 | -4.52054600 |
| C | -1.44075400 | -2.64601800 | -3.25889100 |
| C | 0.29444800 | -1.82626700 | -4.92079800 |
| C | -0.58972100 | -2.33872900 | -2.02762600 |
| H | -2.55087900 | -4.78640500 | -4.50428600 |
| H | -2.12521700 | -2.47017000 | -5.99822100 |
| H | -0.00079900 | -3.87095800 | -4.29117200 |
| H | -2.11671100 | -3.47807300 | -3.01002100 |
| H | -2.09076600 | -1.78283200 | -3.47679100 |
| H | 0.81651500 | -2.03894600 | -5.86717400 |
| H | 1.06168700 | -1.63417400 | -4.15497800 |
| H | -0.28496700 | -0.89624300 | -5.04472800 |
| H | -1.22359800 | -2.23448700 | -1.13454200 |
| H | -0.01190500 | -1.40719800 | -2.13246200 |
| H | 0.12807700 | -3.15486500 | -1.83396300 |
| C | 3.56314200 | -6.65579400 | 0.09087000 |
| C | 3.50220000 | -5.27856900 | 0.73251500 |
| O | 4.50364200 | -4.59227000 | 0.87915000 |
| H | 3.01806500 | -6.66789500 | -0.86689700 |
| N | 2.27074800 | -4.85351400 | 1.17203000 |
| C | 2.10099700 | -3.51189500 | 1.74712500 |
| C | 2.24872400 | -3.43900100 | 3.27934600 |
| O | 1.98418800 | -2.39154600 | 3.85958000 |
| C | 0.68573100 | -3.10764700 | 1.31206400 |
| C | -0.06209000 | -4.44605700 | 1.28168300 |
| C | 0.98423200 | -5.43746800 | 0.75394300 |
| H | 2.87879000 | -2.84003100 | 1.35448800 |
| H | 0.71993100 | -2.66072200 | 0.30919000 |
| H | 0.24265600 | -2.38066200 | 2.00247600 |
| H | -0.96325000 | -4.42448600 | 0.65267800 |
| H | -0.37034300 | -4.72621900 | 2.30186600 |
| H | 0.93741600 | -5.50894300 | -0.34653700 |
| H | 0.84696900 | -6.45281400 | 1.15759400 |
| N | 2.68954900 | -4.56289200 | 3.88804400 |
| C | 2.99197900 | -4.66000100 | 5.30287200 |
| H | 2.87891100 | -5.35580900 | 3.28764200 |
| H | 2.74712500 | -3.69488200 | 5.76512400 |
| C | 7.48480600 | 4.04608800 | 3.56950200 |
| C | 6.34736800 | 4.42069000 | 2.62850200 |
| O | 5.23796800 | 4.73883200 | 3.03082100 |
| H | 8.35834500 | 3.63987500 | 3.04104600 |
| N | 6.68600300 | 4.41603700 | 1.30750500 |
| C | 5.71698200 | 4.71226000 | 0.30047300 |
| C | 4.90655600 | 3.49528800 | -0.18769200 |
| O | 4.98799200 | 2.37975600 | 0.30280000 |
| H | 7.59997300 | 4.04180500 | 1.01954400 |
| H | 4.98857800 | 5.43088500 | 0.71072100 |
| N | 4.04899100 | 3.79868600 | -1.20714200 |
| C | 3.25191700 | 2.80130600 | -1.88519600 |
| C | 3.99422400 | 2.25899300 | -3.11197000 |
| O | 4.55865800 | 3.01478800 | -3.88751100 |
| C | 1.89363700 | 3.40198600 | -2.31348600 |
| C | 1.10467000 | 2.55659600 | -3.30600800 |
| O | 1.11976200 | 3.66753200 | -1.13454300 |
| H | 4.19904000 | 4.66624400 | -1.71067000 |
| H | 3.06609500 | 1.99172100 | -1.16649200 |
| H | 2.09591600 | 4.38710000 | -2.76698300 |
| H | -0.28082200 | 4.40398600 | -1.36808000 |
| H | 0.16127700 | 3.05994600 | -3.55594800 |
| H | 0.86977000 | 1.56667400 | -2.88946100 |
| H | 1.66202500 | 2.42168700 | -4.24323000 |
| N | 3.90925600 | 0.91290800 | -3.27440200 |
| C | 4.46375800 | 0.22004300 | -4.41839500 |
| H | 3.45398200 | 0.37576900 | -2.54641600 |
| H | 5.34978100 | -0.37362200 | -4.13555400 |
| C | -5.24025300 | -1.98753300 | 4.66348500 |
| O | -6.32360300 | -1.88353100 | 5.53907300 |
| C | -4.15791500 | -0.96924100 | 4.97566700 |
| O | -3.08181700 | -1.17227000 | 4.05661800 |
| C | -4.62990100 | 0.49015200 | 4.79284700 |
| O | -4.87070300 | 1.12162000 | 6.02757500 |
| C | -3.49046400 | 1.12822100 | 3.93395400 |
| O | -3.03584100 | 2.37436400 | 4.34346500 |
| C | -2.37801700 | 0.05842200 | 3.97082400 |
| N | 0.74913900 | 0.41252500 | 1.86806500 |
| C | 0.25455600 | 0.62214500 | 0.59324700 |
| O | 0.96315300 | 1.00519300 | -0.33796900 |
| N | -1.08328100 | 0.36483400 | 0.43777200 |
| C | -1.98209600 | 0.02897600 | 1.43141500 |
| O | -3.14411700 | -0.27573600 | 1.10482500 |
| C | -1.45974300 | 0.07626900 | 2.78250200 |
| C | -0.10718400 | 0.21168100 | 2.91971800 |
| H | 1.00383700 | 2.79764700 | -0.69496000 |
| H | -1.52332200 | 0.53692100 | -0.49460500 |
| H | 0.36875900 | 0.16421200 | 3.90024200 |
| H | -1.78170700 | 0.23269700 | 4.88755000 |
| H | -3.84921300 | 2.88130900 | 4.58656900 |
| H | -5.55450400 | 0.49076100 | 4.19011700 |
| H | -3.80676000 | -1.08286000 | 6.01755200 |
| H | -5.53358300 | -1.85561700 | 3.59845300 |
| H | -3.86883500 | 1.14773600 | 2.90229100 |
| H | -4.83238400 | -3.00795900 | 4.76127400 |
| P | -5.95551200 | 2.41949400 | 6.03428300 |
| O | -5.59773500 | 3.17399200 | 4.73955600 |
| O | -6.01594300 | 3.02107500 | 7.38676400 |
| O | -7.35279200 | 1.58644800 | 5.68408000 |
| C | 6.18485800 | -1.39565900 | 4.93225300 |
| C | 5.56072800 | -2.35412200 | 3.92123400 |
| C | 6.17625500 | -2.30892800 | 2.52471800 |
| C | 2.20445900 | 0.49757000 | 2.07335100 |
| S | 5.77101500 | -0.73904200 | 1.63164600 |
| C | 5.72240300 | -1.36661000 | -0.08921800 |
| C | 7.04606000 | -1.50812500 | -0.84322000 |
| O | 7.36136100 | -0.28182000 | -1.52721400 |
| C | 8.31230100 | -1.81591100 | -0.02711400 |
| O | 8.46172000 | -3.14588400 | 0.36524400 |
| C | 9.41081000 | -1.32550300 | -0.97727300 |
| O | 9.60339100 | -2.36515700 | -1.92070700 |
| C | 8.74735700 | -0.08999800 | -1.61994700 |
| C | 10.30523000 | 1.81914400 | -1.05834200 |
| C | 10.18799900 | 2.93496400 | -0.26243500 |
| N | 8.94641400 | 2.95744300 | 0.32332000 |
| C | 8.30970000 | 1.87807900 | -0.09620200 |
| N | 9.09829600 | 1.15329000 | -0.94185100 |
| H | 6.07101100 | -0.34685200 | 4.61153000 |
| H | 4.47952400 | -2.16110300 | 3.85131200 |
| H | 5.67032500 | -3.39101900 | 4.28677600 |
| H | 7.26854800 | -2.43138400 | 2.57734000 |
| H | 5.78154200 | -3.12977400 | 1.91512000 |
| H | 2.45059500 | -0.03585900 | 2.99720400 |
| H | 2.54582400 | 1.54221500 | 2.11246400 |

| | | | |
|---|---|---|---|
| H | 2.71934300 | 0.00700300 | 1.24072400 |
| H | 7.28835200 | 1.59712900 | 0.15901200 |
| H | 9.05379300 | 0.00435600 | -2.67700300 |
| H | 10.43673100 | -2.23178700 | -2.38789300 |
| H | 8.29825200 | -1.18566400 | 0.87483600 |
| H | 6.92200200 | -2.30363500 | -1.60191000 |
| H | 5.21350300 | -2.34084100 | -0.07166500 |
| H | 8.77201400 | -3.62520300 | -0.41932200 |
| H | 10.34638000 | -1.06595700 | -0.45566400 |
| H | 5.09864000 | -0.65086400 | -0.64608000 |
| C | -7.53080900 | 0.75583500 | -1.47636900 |
| C | -6.71547300 | -0.19640000 | -0.60078200 |
| C | -6.84796400 | 0.02034700 | 0.91573600 |
| C | -6.47729500 | 1.41352300 | 1.43015300 |
| N | -5.14568300 | 1.76981900 | 0.95360200 |
| C | -4.50873600 | 2.92203100 | 1.20172000 |
| N | -4.96753700 | 3.77050300 | 2.12429400 |
| N | -3.42590900 | 3.18867400 | 0.45170700 |
| H | -7.20075400 | 1.79997700 | -1.36528400 |
| H | -5.65754200 | -0.13665900 | -0.90220900 |
| H | -7.00364200 | -1.23777200 | -0.82010600 |
| H | -6.21878400 | -0.72371400 | 1.43437000 |
| H | -7.88411800 | -0.18130700 | 1.23306500 |
| H | -6.52576300 | 1.43366800 | 2.53188600 |
| H | -7.19075400 | 2.17277000 | 1.06980500 |
| H | -4.55336900 | 1.01596900 | 0.60620800 |
| H | -5.41920900 | 3.44120500 | 3.01445500 |
| H | -4.47885500 | 4.65176800 | 2.23304900 |
| H | -3.20895400 | 2.60975900 | -0.36938100 |
| H | -2.81607300 | 3.96824500 | 0.65944800 |
| H | 0.65289500 | 8.92784000 | -1.81387100 |
| H | -0.60429800 | 8.52069000 | -3.01677000 |
| H | -4.56015900 | 7.30311000 | -4.73253400 |
| H | -5.42121300 | 7.00006800 | -6.26448500 |
| H | 3.10570900 | -7.42173400 | 0.73585800 |
| H | 4.06071500 | -4.87132300 | 5.46666800 |
| H | 10.92318300 | 3.71864300 | -0.08766900 |
| H | 7.78258300 | 4.94058400 | 4.13895700 |
| H | 7.11065900 | 3.30736100 | 4.29314700 |
| H | 11.11048300 | 1.45365500 | -1.69054400 |
| H | 6.20569900 | 5.17785100 | -0.57258800 |
| H | 7.26399900 | -1.58842000 | 5.05290500 |
| H | 4.61557900 | -6.90130900 | -0.09106800 |
| H | 2.39392900 | -5.44982200 | 5.78535200 |
| H | -6.60815900 | -0.95629200 | 5.58340700 |
| H | -7.87671400 | 1.55460100 | 6.49765200 |
| H | -8.60408700 | 0.71401400 | -1.22797200 |
| H | -4.09787900 | -6.55987300 | -5.25761800 |
| H | 5.71080500 | -1.49232400 | 5.92170800 |
| H | 4.77292900 | 0.97829100 | -5.14950200 |
| H | -0.89152000 | -3.57592600 | -6.59123500 |
| H | -7.43183200 | 0.49214800 | -2.54173900 |
| H | 3.71401900 | -0.44414100 | -4.87722200 |
| H | -4.26764000 | -8.14612200 | -0.91043100 |

#### Model5 – Initial state

| | | | |
|---|---|---|---|
| O | -4.61133300 | -1.69397900 | -3.90540100 |
| H | -4.43645200 | -1.90398600 | -4.84287600 |
| H | -4.93317800 | -2.55633300 | -3.55219800 |
| O | -1.21163700 | 4.18676900 | 2.08463300 |
| H | -1.75413000 | 4.58439200 | 1.34927900 |
| H | -1.81403800 | 4.14685800 | 2.84018000 |
| O | -3.29772200 | 0.07874200 | -2.38283400 |
| H | -3.76731400 | -0.55437300 | -2.99063200 |
| H | -3.22811400 | 0.92847400 | -2.86277000 |
| O | -1.39465000 | -7.00325500 | 2.65218700 |
| H | -1.69475300 | -7.91990700 | 2.69972000 |
| H | -2.09919300 | -6.46449400 | 3.09867900 |
| O | -5.86429400 | 1.43468400 | 4.11328000 |
| H | -6.49202700 | 2.08949200 | 4.44925000 |
| H | -5.21835700 | 1.28030600 | 4.89762500 |
| O | -2.58653500 | 2.77342200 | -2.95792200 |
| H | -1.75887400 | 2.53593600 | -2.49622400 |
| H | -2.64799700 | 3.73819000 | -2.76355900 |
| C | -7.43231300 | 5.75236500 | -2.31158100 |
| C | -6.06089600 | 6.46257400 | -2.37063300 |
| O | -5.39638200 | 6.65967800 | -1.35208800 |
| C | -7.42335100 | 4.39432100 | -3.03444400 |
| C | -6.53288400 | 3.31791300 | -2.39058300 |
| C | -7.06396800 | 2.90926600 | -1.01134100 |
| C | -6.40336900 | 2.09927200 | -3.31007500 |
| H | -7.68647000 | 5.64317000 | -1.24958000 |
| H | -7.10454900 | 4.54523000 | -4.08057800 |
| H | -8.45833400 | 4.01242300 | -3.08918800 |
| H | -5.52084400 | 3.73897300 | -2.25565900 |
| H | -6.56645100 | 1.99517300 | -0.66299200 |
| H | -8.14533000 | 2.69342900 | -1.05557600 |
| H | -6.89335600 | 3.68452600 | -0.25106400 |
| H | -7.37671700 | 1.59768500 | -3.44773200 |
| H | -6.02805500 | 2.38609300 | -4.30554600 |
| H | -5.70305200 | 1.35673500 | -2.89886000 |
| N | -3.50577800 | 8.00441700 | 1.20136300 |
| C | -2.10993200 | 7.60266800 | 0.96318500 |
| C | -1.81707500 | 7.18933600 | -0.48612300 |
| C | -2.29178000 | 5.79892200 | -0.91122200 |
| O | -2.64735200 | 5.00237800 | 0.03980900 |
| O | -2.26215000 | 5.47873200 | -2.10830900 |
| H | -4.18013400 | 7.26887100 | 1.40997600 |
| H | -1.85545500 | 6.78841000 | 1.65437200 |
| H | -0.72356200 | 7.19471500 | -0.64903000 |
| H | -2.22951700 | 7.93584200 | -1.18036000 |
| C | -7.34126900 | -1.33316700 | -1.03197000 |
| C | -5.96771900 | -1.90157900 | -0.67309300 |
| C | -5.53058100 | -1.67086200 | 0.77896200 |
| C | -5.49391900 | -0.21241300 | 1.22081200 |
| N | -4.54959000 | 0.56353900 | 0.42181500 |
| C | -4.36448400 | 1.88800300 | 0.58421200 |
| N | -4.76666200 | 2.51488200 | 1.69585700 |
| N | -3.76374500 | 2.57874600 | -0.37846300 |
| H | -7.36912200 | -0.23455500 | -0.95868700 |
| H | -5.21705300 | -1.48360300 | -1.36292300 |
| H | -5.96124800 | -2.98296800 | -0.87243700 |
| H | -4.53623700 | -2.11714500 | 0.94807500 |
| H | -6.22373700 | -2.19387500 | 1.46031200 |
| H | -5.21224200 | -0.15659000 | 2.28086200 |
| H | -6.49903000 | 0.23988000 | 1.15173600 |
| H | -4.22579900 | 0.19175700 | -0.47574700 |
| H | -5.17044500 | 2.02364200 | 2.50741400 |
| H | -4.89933900 | 3.53294200 | 1.63641500 |
| H | -3.55781500 | 2.18126900 | -1.29156200 |
| H | -3.38851300 | 3.53615100 | -0.22979000 |
| C | -1.27181200 | -8.39248400 | -2.22513100 |
| C | -1.41759500 | -8.05173900 | -0.74198500 |
| C | -2.82633700 | -7.57542500 | -0.35012900 |
| N | -3.22345700 | -6.30770300 | -0.92502900 |
| C | -2.89348300 | -5.10699500 | -0.43122000 |
| N | -2.07190400 | -4.97535400 | 0.62862400 |
| N | -3.37794700 | -4.01453000 | -1.00876000 |
| H | -1.54470400 | -7.53144100 | -2.85432500 |
| H | -0.23856800 | -8.68408200 | -2.47200700 |
| H | -0.68785300 | -7.27544600 | -0.45633900 |
| H | -1.17598500 | -8.93837100 | -0.12840700 |
| H | -2.89832300 | -7.51756500 | 0.74659300 |
| H | -3.56765700 | -8.32270500 | -0.67568100 |
| H | -3.71571500 | -6.30847500 | -1.85696200 |
| H | -1.73801100 | -5.77966100 | 1.15172100 |
| H | -1.93022900 | -4.03369700 | 0.99957700 |
| H | -4.06881700 | -4.09071100 | -1.79267800 |
| H | -2.97683200 | -3.12006100 | -0.73155300 |
| C | -4.68972700 | -5.41504900 | -6.16998800 |

| | | | |
|---|---|---|---|
| C | -3.76418200 | -4.20054800 | -6.10630300 |
| O | -4.18183300 | -3.06390500 | -6.31793500 |
| C | -5.74025200 | -5.31623800 | -5.05513100 |
| C | -5.03921300 | -5.28068700 | -3.69728400 |
| O | -5.14282000 | -4.21825100 | -3.01712900 |
| O | -4.35419300 | -6.28396300 | -3.37700400 |
| H | -5.17947300 | -5.40844300 | -7.15609600 |
| H | -6.39904900 | -6.19762100 | -5.09807500 |
| H | -6.34398200 | -4.40853700 | -5.18643400 |
| N | -2.47559400 | -4.45480800 | -5.75358500 |
| C | -1.51229600 | -3.41058200 | -5.42116300 |
| C | -1.41633600 | -3.17435000 | -3.90401400 |
| C | -0.52204300 | -1.97661300 | -3.54163100 |
| C | -1.00179300 | -4.45107900 | -3.16897000 |
| C | 0.95880600 | -2.09674200 | -3.90950300 |
| H | -2.25897400 | -5.40476400 | -5.47742700 |
| H | -1.84727100 | -2.49595500 | -5.92974100 |
| H | -2.43254100 | -2.90545200 | -3.57675600 |
| H | -0.60515300 | -1.83551200 | -2.45236700 |
| H | -0.94534900 | -1.06659300 | -3.99892600 |
| H | -0.06036400 | -4.86705600 | -3.56881500 |
| H | -1.77894200 | -5.22482300 | -3.24280300 |
| H | -0.85979000 | -4.24135800 | -2.10078500 |
| H | 1.51360200 | -1.19998400 | -3.58748600 |
| H | 1.11412400 | -2.20245900 | -4.99521700 |
| H | 1.43408200 | -2.96422800 | -3.42219200 |
| C | 4.98012700 | -6.07784100 | -2.71974600 |
| C | 4.83188100 | -4.94168900 | -1.71931900 |
| O | 5.76295400 | -4.20107800 | -1.41920000 |
| H | 4.25223000 | -5.98673100 | -3.54118900 |
| N | 3.60453600 | -4.79220500 | -1.13017500 |
| C | 3.41848000 | -3.71998500 | -0.15259200 |
| C | 4.11670200 | -4.01279800 | 1.18162100 |
| O | 4.58098100 | -3.09942400 | 1.86125100 |
| C | 1.88352800 | -3.62957000 | 0.00569800 |
| C | 1.39820100 | -5.04141300 | -0.34452700 |
| C | 2.34737400 | -5.47814000 | -1.46701200 |
| H | 3.85612200 | -2.78198300 | -0.51956800 |
| H | 1.48767700 | -2.90660800 | -0.72429700 |
| H | 1.57544900 | -3.29391100 | 1.00493900 |
| H | 0.34416600 | -5.06082700 | -0.64345200 |
| H | 1.49472500 | -5.70360100 | 0.52951400 |
| H | 1.97237600 | -5.14639600 | -2.45086100 |
| H | 2.48391400 | -6.56945700 | -1.51557900 |
| N | 4.11970600 | -5.31533400 | 1.55607800 |
| C | 4.38637200 | -5.79061500 | 2.90117100 |
| C | 3.10545000 | -6.20999100 | 3.63866600 |
| C | 2.08429900 | -5.11756700 | 3.63426900 |
| C | 2.19261400 | -3.75075700 | 3.82447600 |
| N | 0.76425100 | -5.32052400 | 3.28778200 |
| C | 0.14784700 | -4.10547600 | 3.28210200 |
| N | 0.98635900 | -3.13616200 | 3.59523000 |
| H | 3.73077900 | -5.97273200 | 0.89153400 |
| H | 4.87882200 | -4.96471600 | 3.43264000 |
| H | 2.68117800 | -7.10894900 | 3.15846000 |
| H | 0.26997900 | -6.19681700 | 3.10143700 |
| H | 3.08642900 | -3.18180400 | 4.06994200 |
| H | -0.90053700 | -3.97293500 | 3.03281800 |
| H | 3.38325100 | -6.51540400 | 4.66323800 |
| C | 7.78269100 | 4.03017800 | 4.00376400 |
| C | 6.49924800 | 4.55100100 | 3.36851200 |
| O | 5.41316700 | 4.44286500 | 3.91426300 |
| H | 8.55131900 | 3.79341700 | 3.25340400 |
| N | 6.66900900 | 5.19688000 | 2.17282900 |
| C | 5.50832400 | 5.57224000 | 1.41276800 |
| C | 4.89785400 | 4.37994400 | 0.66263300 |
| O | 5.31714200 | 3.22740300 | 0.78764700 |
| H | 7.53935100 | 5.02688300 | 1.65518500 |
| H | 4.73790000 | 5.96190600 | 2.09562600 |
| N | 3.83922200 | 4.68086000 | -0.11434100 |

| | | | |
|---|---|---|---|
| C | 3.21991100 | 3.68330700 | -0.94551900 |
| C | 4.14056200 | 3.36409100 | -2.14451100 |
| O | 5.18675000 | 3.95313200 | -2.33722700 |
| C | 1.80160700 | 4.11580100 | -1.33478200 |
| C | 0.96343100 | 4.46718200 | -0.11485900 |
| O | 1.93561600 | 5.21916500 | -2.21782300 |
| H | 3.64710900 | 5.64841900 | -0.35236500 |
| H | 3.13089800 | 2.76025800 | -0.35243600 |
| H | 1.31408400 | 3.26271000 | -1.84142400 |
| H | 1.04566500 | 5.52720700 | -2.44244700 |
| H | 1.31142700 | 5.40014500 | 0.35046100 |
| H | 1.01122500 | 3.67731800 | 0.64484500 |
| H | -0.08700300 | 4.57684700 | -0.40559800 |
| N | 3.72943300 | 2.31055200 | -2.91642400 |
| C | 4.42709600 | 1.99677300 | -4.15126700 |
| H | 2.75944400 | 2.02410400 | -2.83383300 |
| H | 5.50897000 | 2.03266000 | -3.96249300 |
| C | -3.88967700 | -4.23592600 | 3.82007300 |
| O | -3.09911600 | -5.42824100 | 3.96911600 |
| C | -3.06694800 | -2.98102300 | 4.00150400 |
| O | -2.45035200 | -2.59125600 | 2.77893000 |
| C | -3.80792900 | -1.73247000 | 4.47222700 |
| O | -4.00025900 | -1.79707600 | 5.85205700 |
| C | -2.87004300 | -0.58558700 | 3.97017200 |
| O | -2.17965000 | 0.11037300 | 4.95584100 |
| C | -1.84651700 | -1.32883300 | 3.08162700 |
| N | -0.86456500 | 1.66846600 | 1.07940800 |
| C | -0.88908000 | 1.30065300 | -0.24222300 |
| O | -0.62591900 | 2.06424100 | -1.16680000 |
| N | -1.22126600 | -0.01859700 | -0.45241400 |
| C | -1.45596600 | -1.00052800 | 0.50903100 |
| O | -1.64692000 | -2.16026000 | 0.12736700 |
| C | -1.46349800 | -0.52041900 | 1.87234200 |
| C | -1.18400300 | 0.79948500 | 2.07692000 |
| H | -0.83981500 | 2.68072600 | 1.33877800 |
| H | -1.53694600 | -0.24020000 | -1.40519700 |
| H | -1.21871500 | 1.22172500 | 3.08537200 |
| H | -0.95229400 | -1.51120800 | 3.70370700 |
| H | -2.88681400 | 0.49768700 | 5.54179800 |
| H | -4.77905800 | -1.66243700 | 3.94842200 |
| H | -2.28974000 | -3.17239700 | 4.77095800 |
| H | -4.70405700 | -4.24021700 | 4.56281800 |
| H | -3.46563700 | 0.09930600 | 3.33963800 |
| H | -4.32323400 | -4.26597200 | 2.81107900 |
| P | -4.98909200 | -0.65275200 | 6.57932200 |
| O | -4.44012300 | 0.71736600 | 6.09145100 |
| O | -5.12059100 | -1.00820800 | 8.00756500 |
| O | -6.37951800 | -0.87395200 | 5.72851500 |
| C | 7.63036600 | -1.37099100 | 3.65344200 |
| C | 8.21704500 | -1.75701300 | 2.29654800 |
| C | 7.24687000 | -1.66120300 | 1.12553000 |
| C | 5.11196600 | 0.25086600 | 1.51763200 |
| S | 6.74097500 | 0.07835500 | 0.74630200 |
| C | 6.32542700 | -0.12122800 | -1.03230500 |
| C | 7.51275100 | 0.14559200 | -1.95819500 |
| O | 7.59648800 | 1.53058900 | -2.25790300 |
| C | 8.92002900 | -0.23061400 | -1.46981300 |
| O | 9.15611200 | -1.60287500 | -1.68135000 |
| C | 9.79184200 | 0.68723600 | -2.34329300 |
| O | 9.87526800 | 0.17638900 | -3.64435200 |
| C | 8.93803100 | 1.96205300 | -2.37522000 |
| C | 10.44401900 | 3.47689500 | -1.00855100 |
| C | 10.22214500 | 4.29477300 | 0.07524400 |
| N | 8.91038400 | 4.20002000 | 0.47328500 |
| C | 8.33014500 | 3.34485800 | -0.34664300 |
| N | 9.22744300 | 2.86434900 | -1.26423600 |
| H | 7.39448200 | -0.29474900 | 3.70461200 |
| H | 8.53898100 | -2.81254400 | 2.32240700 |
| H | 9.12651100 | -1.16635700 | 2.08654100 |
| H | 7.69886300 | -2.01999700 | 0.19066900 |

| | | | |
|---|---|---|---|
| H | 6.31350700 | -2.21940000 | 1.30185800 |
| H | 5.29141800 | 0.28249200 | 2.59962100 |
| H | 4.74319100 | 1.23276500 | 1.18338700 |
| H | 4.48609600 | -0.61413400 | 1.26273100 |
| H | 7.27786900 | 3.06845200 | -0.33462800 |
| H | 9.06840000 | 2.49251500 | -3.33076400 |
| H | 9.89601800 | -0.79133800 | -3.57198300 |
| H | 9.05172700 | 0.05599600 | -0.41135800 |
| H | 7.32058600 | -0.42201200 | -2.88616300 |
| H | 5.94921100 | -1.14800800 | -1.15403300 |
| H | 9.92856500 | -1.88060200 | -1.17023900 |
| H | 10.78745500 | 0.86976600 | -1.89829600 |
| H | 5.52404700 | 0.60094800 | -1.24977200 |
| C | -5.46101100 | 9.55663300 | 1.14522700 |
| C | -3.95829900 | 9.26610200 | 0.90879700 |
| O | -3.21172800 | 10.13754700 | 0.48423600 |
| H | -5.61251100 | 10.63749100 | 1.04626400 |
| N | -5.62954200 | 6.83471100 | -3.60699200 |
| C | -4.34276300 | 7.47940300 | -3.92855700 |
| H | -6.26663800 | 6.67059800 | -4.37653600 |
| H | -4.33883000 | 8.53916500 | -3.62463400 |
| O | -5.05275600 | 5.39683900 | 1.12069000 |
| H | -4.13379300 | 5.30771100 | 0.77235800 |
| H | -5.48774100 | 5.77827900 | 0.32851300 |
| H | -6.05531800 | 9.02331900 | 0.38546200 |
| H | -5.78934200 | 9.21082900 | 2.13764900 |
| H | -1.48526500 | 8.47330000 | 1.20798500 |
| H | -3.52474100 | 6.95546100 | -3.41062000 |
| H | -4.18856500 | 7.41365900 | -5.01344700 |
| H | -8.12860100 | -1.73281500 | -0.37063400 |
| H | -7.60081400 | -1.59515900 | -2.06888600 |
| H | -4.12681400 | -6.35378300 | -6.06217900 |
| H | -1.93557200 | -9.22545600 | -2.50791200 |
| H | -0.53174200 | -3.69432700 | -5.83959000 |
| H | 4.80525300 | -7.05033200 | -2.23238900 |
| H | 5.99810100 | -6.05212100 | -3.12516400 |
| H | 4.14636100 | 0.98743300 | -4.48592000 |
| H | 4.19779200 | 2.71514500 | -4.95758700 |
| H | 11.32771700 | 3.29254000 | -1.61377200 |
| H | 10.93397500 | 4.94284100 | 0.58373000 |
| H | 8.18502400 | 4.79654300 | 4.68588500 |
| H | 7.54077300 | 3.14207200 | 4.60315500 |
| H | 5.76460900 | 6.35957300 | 0.68584600 |
| H | -6.45493600 | -0.18629500 | 5.03919600 |
| H | 5.09927300 | -6.63098300 | 2.86648500 |
| H | 8.34297400 | -1.58146500 | 4.46412200 |
| H | 6.70392300 | -1.93292600 | 3.85071300 |
| H | -2.75849000 | -5.43174000 | 4.87550300 |
| H | -8.19165200 | 6.41046200 | -2.76764700 |

#### Model5 − TS1

| | | | |
|---|---|---|---|
| O | -4.64084500 | -1.58579400 | -3.90446700 |
| H | -4.44725900 | -1.83714100 | -4.82827400 |
| H | -5.01492500 | -2.41915100 | -3.53776300 |
| O | -1.23243900 | 3.11116900 | 2.17353800 |
| H | -1.76200900 | 3.88481700 | 1.70875000 |
| H | -1.83151300 | 2.58340900 | 2.73832100 |
| O | -3.14788700 | 0.20396400 | -2.50970800 |
| H | -3.70234900 | -0.40426400 | -3.06313600 |
| H | -2.90445900 | 1.01159200 | -3.01958900 |
| O | -1.65640800 | -6.81453400 | 2.74879800 |
| H | -1.99327700 | -7.71866900 | 2.79085700 |
| H | -2.35857800 | -6.24880100 | 3.15880300 |
| O | -6.26590100 | 1.78827400 | 3.94342300 |
| H | -6.86551300 | 2.46884600 | 4.27820600 |
| H | -5.56970400 | 1.67817900 | 4.68553900 |
| O | -1.99206400 | 2.58196000 | -3.09972800 |
| H | -1.23781300 | 2.24134000 | -2.57353800 |
| H | -2.23493400 | 3.38453300 | -2.60111100 |
| C | -7.14935000 | 6.00394000 | -2.70560200 |
| C | -5.74383300 | 6.65077400 | -2.74491100 |
| O | -5.06370600 | 6.79046100 | -1.72498900 |
| C | -7.11564900 | 4.56259100 | -3.24726400 |
| C | -6.41452300 | 3.53807600 | -2.33637100 |
| C | -7.26862800 | 3.23550300 | -1.10084800 |
| C | -6.09587300 | 2.26056900 | -3.11873200 |
| H | -7.48713600 | 6.03698800 | -1.66184800 |
| H | -6.62355300 | 4.56768100 | -4.23555600 |
| H | -8.15110100 | 4.22529100 | -3.42859200 |
| H | -5.45764500 | 3.96934500 | -1.99082800 |
| H | -6.80148000 | 2.46504000 | -0.47483300 |
| H | -8.26406200 | 2.86408000 | -1.39894100 |
| H | -7.40421700 | 4.12036300 | -0.46097200 |
| H | -7.01667800 | 1.78489900 | -3.49695800 |
| H | -5.44608800 | 2.47307300 | -3.98305300 |
| H | -5.57330600 | 1.52112800 | -2.49657900 |
| N | -3.19962300 | 8.27318100 | 0.41098600 |
| C | -1.82453300 | 7.77539900 | 0.60868000 |
| C | -1.51401100 | 6.70318800 | -0.42670000 |
| C | -2.12559800 | 5.33169300 | -0.12984800 |
| O | -2.34708700 | 5.06458000 | 1.09638800 |
| O | -2.33105400 | 4.53983100 | -1.07244600 |
| H | -3.79810300 | 7.78504600 | -0.25106500 |
| H | -1.70870400 | 7.35525100 | 1.61900000 |
| H | -0.42410700 | 6.53768400 | -0.45138900 |
| H | -1.80831600 | 7.01504100 | -1.44086700 |
| C | -7.32162200 | -1.03758100 | -1.20988300 |
| C | -6.21420100 | -1.83532300 | -0.53281200 |
| C | -6.08990900 | -1.58444800 | 0.97552200 |
| C | -5.95220900 | -0.11114900 | 1.35475500 |
| N | -4.82118100 | 0.46474300 | 0.64071900 |
| C | -4.49870100 | 1.76893300 | 0.58729200 |
| N | -4.99976800 | 2.63072900 | 1.46736200 |
| N | -3.66702100 | 2.15934000 | -0.38298100 |
| H | -7.14364600 | 0.04654400 | -1.14587100 |
| H | -5.26101100 | -1.60093100 | -1.03247700 |
| H | -6.36753500 | -2.91101300 | -0.70783200 |
| H | -5.21355000 | -2.13124500 | 1.36082400 |
| H | -6.97525600 | -1.98030700 | 1.50069600 |
| H | -5.83236700 | -0.00269900 | 2.44179100 |
| H | -6.86696200 | 0.44844600 | 1.09735000 |
| H | -4.25929800 | -0.17258900 | 0.08144100 |
| H | -5.48992700 | 2.30815800 | 2.31493700 |
| H | -5.01264600 | 3.63690500 | 1.22844700 |
| H | -3.50996700 | 1.51554000 | -1.16225200 |
| H | -3.25657800 | 3.10403100 | -0.45115400 |
| C | -1.47609100 | -8.32972800 | -2.07622600 |
| C | -1.63900100 | -7.95244300 | -0.60409800 |
| C | -3.03844200 | -7.41851300 | -0.26261900 |
| N | -3.37823500 | -6.16510600 | -0.89891100 |
| C | -3.14677600 | -4.94542100 | -0.39131700 |
| N | -2.38486500 | -4.76530400 | 0.71243700 |
| N | -3.65217200 | -3.89096400 | -1.00584700 |
| H | -1.70250700 | -7.47131000 | -2.72766400 |
| H | -0.45057700 | -8.66671100 | -2.29644400 |
| H | -0.89057100 | -7.19404000 | -0.31808300 |
| H | -1.44279100 | -8.83322700 | 0.03377200 |
| H | -3.13252600 | -7.31207200 | 0.82826700 |
| H | -3.79465700 | -8.15777800 | -0.57416100 |
| H | -3.84430200 | -6.18164800 | -1.84329300 |
| H | -2.03164000 | -5.55943100 | 1.23844500 |
| H | -2.42190100 | -3.85949800 | 1.18507500 |
| H | -4.33857900 | -4.00382600 | -1.79057200 |
| H | -3.38337800 | -2.96041500 | -0.67622200 |
| C | -4.72160300 | -5.36297300 | -6.17165400 |
| C | -3.76476600 | -4.17486300 | -6.08655300 |
| O | -4.14487500 | -3.02548700 | -6.30201200 |
| C | -5.81850500 | -5.21056700 | -5.11058100 |
| C | -5.18375800 | -5.15481600 | -3.71997900 |
| O | -5.37473700 | -4.10873200 | -3.03578700 |

| | | | |
|---|---|---|---|
| O | -4.46759100 | -6.13068400 | -3.37788600 |
| H | -5.16271500 | -5.36241700 | -7.18083100 |
| H | -6.49815600 | -6.07616500 | -5.16100300 |
| H | -6.39467600 | -4.29337300 | -5.28988400 |
| N | -2.48840200 | -4.46608900 | -5.72019100 |
| C | -1.49143600 | -3.44401900 | -5.42604900 |
| C | -1.44398000 | -3.06470400 | -3.93485700 |
| C | -0.49625500 | -1.88294000 | -3.66480900 |
| C | -1.12915100 | -4.28016700 | -3.06048600 |
| C | 0.98983500 | -2.12426400 | -3.94192800 |
| H | -2.30433900 | -5.41638000 | -5.42268800 |
| H | -1.75169500 | -2.56220100 | -6.02826300 |
| H | -2.45524900 | -2.71309900 | -3.67577700 |
| H | -0.61332200 | -1.59558200 | -2.60870000 |
| H | -0.84245300 | -1.01011400 | -4.24318700 |
| H | -0.17920100 | -4.76074600 | -3.35352500 |
| H | -1.92791600 | -5.03267100 | -3.12059900 |
| H | -1.05212400 | -3.98103700 | -2.00561800 |
| H | 1.57859300 | -1.22521200 | -3.69631700 |
| H | 1.18782900 | -2.36521100 | -4.99889400 |
| H | 1.39259000 | -2.95201000 | -3.33439600 |
| C | 4.85870000 | -6.24231900 | -2.52598700 |
| C | 4.71060600 | -5.07787600 | -1.55570100 |
| O | 5.65288200 | -4.35449600 | -1.24928800 |
| H | 4.16012500 | -6.14995600 | -3.37244400 |
| N | 3.47262700 | -4.87956100 | -1.00129600 |
| C | 3.29134200 | -3.77073700 | -0.06259600 |
| C | 3.95258100 | -4.03241500 | 1.29657300 |
| O | 4.42542300 | -3.10772900 | 1.95388300 |
| C | 1.75613200 | -3.63843000 | 0.06346200 |
| C | 1.23825800 | -5.04377600 | -0.26992500 |
| C | 2.20413900 | -5.53427600 | -1.35648800 |
| H | 3.75792100 | -2.85550800 | -0.45073700 |
| H | 1.39455100 | -2.91561800 | -0.68475800 |
| H | 1.44197300 | -3.27712500 | 1.05286200 |
| H | 0.19310300 | -5.04073500 | -0.60137000 |
| H | 1.29159200 | -5.68821400 | 0.62123700 |
| H | 1.86556400 | -5.21374000 | -2.35747000 |
| H | 2.30794500 | -6.63003800 | -1.37702200 |
| N | 3.91031200 | -5.31964500 | 1.71601700 |
| C | 4.17742500 | -5.76109600 | 3.07151000 |
| C | 2.90471700 | -6.17459700 | 3.81938800 |
| C | 1.90289700 | -5.06749900 | 3.82328900 |
| C | 2.04032700 | -3.70361100 | 4.01575000 |
| N | 0.57990600 | -5.24282400 | 3.47778800 |
| C | -0.01106700 | -4.01550700 | 3.47695200 |
| N | 0.84663600 | -3.06360500 | 3.79222600 |
| H | 3.52686200 | -5.99280800 | 1.06421200 |
| H | 4.66351100 | -4.91967500 | 3.58383800 |
| H | 2.46705500 | -7.06861400 | 3.34184800 |
| H | 0.06872400 | -6.10332800 | 3.27093700 |
| H | 2.94683300 | -3.15427500 | 4.26092600 |
| H | -1.05440000 | -3.86037100 | 3.22249500 |
| H | 3.19024400 | -6.48381200 | 4.84073200 |
| C | 7.88634800 | 3.96986300 | 3.93789400 |
| C | 6.63547300 | 4.51954600 | 3.26201000 |
| O | 5.53241400 | 4.44534200 | 3.77821700 |
| H | 8.65813300 | 3.68228700 | 3.20866500 |
| N | 6.85113100 | 5.14823000 | 2.06277600 |
| C | 5.71144700 | 5.50678700 | 1.25984300 |
| C | 5.09800100 | 4.27856400 | 0.57036900 |
| O | 5.53125400 | 3.13573700 | 0.73430700 |
| H | 7.72331400 | 4.93410100 | 1.56544900 |
| H | 4.93682300 | 5.95494200 | 1.90008300 |
| N | 4.02919100 | 4.53554000 | -0.20478400 |
| C | 3.42709200 | 3.50752200 | -1.01055600 |
| C | 4.34403200 | 3.17764500 | -2.20859200 |
| O | 5.37777200 | 3.77979300 | -2.42681900 |
| C | 2.01299900 | 3.93243100 | -1.41023300 |
| C | 1.13644900 | 4.19796000 | -0.19587000 |

| | | | |
|---|---|---|---|
| O | 2.16673400 | 5.08963400 | -2.22584900 |
| H | 3.81475000 | 5.49069000 | -0.47333100 |
| H | 3.33638200 | 2.59604300 | -0.40104700 |
| H | 1.55086300 | 3.10706300 | -1.97689700 |
| H | 1.28469600 | 5.40400400 | -2.46374900 |
| H | 1.44466400 | 5.11922500 | 0.31968400 |
| H | 1.18985200 | 3.36237900 | 0.51149900 |
| H | 0.09221000 | 4.27734900 | -0.51946000 |
| N | 3.93506300 | 2.10780800 | -2.96011900 |
| C | 4.60555800 | 1.79834100 | -4.21183700 |
| H | 2.97530100 | 1.79726200 | -2.84477400 |
| H | 5.69160600 | 1.83203700 | -4.04806300 |
| C | -4.12127900 | -3.94482800 | 3.87521600 |
| O | -3.30874800 | -5.11567700 | 3.99063200 |
| C | -3.26183400 | -2.72340700 | 4.03455600 |
| O | -2.48385600 | -2.52666300 | 2.86876800 |
| C | -3.97325500 | -1.40133000 | 4.32735700 |
| O | -4.01306200 | -1.27792500 | 5.72115600 |
| C | -3.10817500 | -0.36944100 | 3.52503400 |
| O | -2.64996000 | 0.77621100 | 4.17803200 |
| C | -1.91788500 | -1.22909500 | 3.04017200 |
| N | -0.10210500 | 1.26369600 | 0.96906800 |
| C | -0.46909200 | 0.97953200 | -0.32288000 |
| O | -0.12201100 | 1.66037400 | -1.29328300 |
| N | -1.28200100 | -0.12875100 | -0.51194000 |
| C | -1.84499900 | -0.88729400 | 0.49523500 |
| O | -2.79166300 | -1.64531000 | 0.22701600 |
| C | -1.28916400 | -0.68064700 | 1.80629100 |
| C | -0.44975000 | 0.38753900 | 1.95113700 |
| H | -0.57779600 | 2.37741000 | 1.49156300 |
| H | -1.79073900 | -0.17407000 | -1.41843800 |
| H | -0.12810100 | 0.67399000 | 2.95749400 |
| H | -1.16757700 | -1.26160400 | 3.85006700 |
| H | -3.32478500 | 1.03013500 | 4.87716000 |
| H | -4.99859300 | -1.41162600 | 3.92351000 |
| H | -2.58970500 | -2.87632600 | 4.90504800 |
| H | -4.91433200 | -3.94652900 | 4.64134800 |
| H | -3.68867000 | -0.10149100 | 2.62942300 |
| H | -4.58468200 | -3.97139700 | 2.87931700 |
| P | -5.08001400 | -0.20081800 | 6.41895400 |
| O | -4.65930400 | 1.16946200 | 5.81488200 |
| O | -5.13391100 | -0.47301900 | 7.87033600 |
| O | -6.47836600 | -0.58590700 | 5.63456700 |
| C | 7.55642900 | -1.43115500 | 3.74949800 |
| C | 8.12671800 | -1.88011500 | 2.40481700 |
| C | 7.16118200 | -1.78239700 | 1.22958600 |
| C | 5.08772600 | 0.20853400 | 1.51811600 |
| S | 6.72925600 | -0.03557200 | 0.79404300 |
| C | 6.35357100 | -0.26044700 | -0.99074300 |
| C | 7.57423800 | -0.07263400 | -1.89351600 |
| O | 7.72036500 | 1.29564900 | -2.24311500 |
| C | 8.95354200 | -0.48321200 | -1.35566900 |
| O | 9.14024900 | -1.87139500 | -1.50697700 |
| C | 9.88164700 | 0.36239000 | -2.24433000 |
| O | 9.97656600 | -0.20838400 | -3.51970600 |
| C | 9.08073100 | 1.66715900 | -2.35279000 |
| C | 10.61331900 | 3.20879500 | -1.04786300 |
| C | 10.40852500 | 4.06829700 | 0.00691700 |
| N | 9.09643500 | 4.01074500 | 0.41159900 |
| C | 8.49973900 | 3.13668300 | -0.37615900 |
| N | 9.38553600 | 2.60913900 | -1.27920100 |
| H | 7.35414300 | -0.34691300 | 3.76477700 |
| H | 8.40883600 | -2.94574300 | 2.46288400 |
| H | 9.05793900 | -1.33081400 | 2.17804500 |
| H | 7.59900200 | -2.19129000 | 0.30880300 |
| H | 6.20564600 | -2.29593700 | 1.42223400 |
| H | 5.23958300 | 0.24468000 | 2.60423900 |
| H | 4.76844900 | 1.19972700 | 1.16282800 |
| H | 4.43377700 | -0.63268100 | 1.25500700 |
| H | 7.44312000 | 2.87938800 | -0.34935100 |

| | | | |
|---|---|---|---|
| H | 9.25230800 | 2.14666300 | -3.32856400 |
| H | 9.95473700 | -1.17197000 | -3.40441000 |
| H | 9.07130600 | -0.15967600 | -0.30633700 |
| H | 7.38348800 | -0.66745900 | -2.80461600 |
| H | 5.93454100 | -1.27215100 | -1.09654400 |
| H | 9.88517500 | -2.15837100 | -0.96133300 |
| H | 10.87264900 | 0.52684400 | -1.78216300 |
| H | 5.59027800 | 0.48911300 | -1.24960300 |
| C | -5.11012600 | 9.84544700 | 0.66969200 |
| C | -3.66097700 | 9.42197100 | 0.99507600 |
| O | -2.96459000 | 10.09467500 | 1.73949800 |
| H | -5.66203700 | 9.91947100 | 1.61791700 |
| N | -5.29582200 | 7.02426800 | -3.97983500 |
| C | -3.97511500 | 7.57569200 | -4.31726200 |
| H | -5.92454400 | 6.84936200 | -4.75527900 |
| H | -3.45580900 | 7.80839500 | -3.37842100 |
| O | -5.13572800 | 5.37827100 | 0.82472400 |
| H | -4.22439500 | 5.54803600 | 1.12853500 |
| H | -5.17723800 | 5.78426900 | -0.06240800 |
| H | -5.08290600 | 10.84899600 | 0.22011000 |
| H | -5.62821600 | 9.14642300 | -0.00318900 |
| H | -1.13388200 | 8.62898300 | 0.52223300 |
| H | -3.37509000 | 6.84626700 | -4.88458200 |
| H | -4.08559000 | 8.49786100 | -4.90735100 |
| H | -8.30718500 | -1.24340600 | -0.75849100 |
| H | -7.37454000 | -1.29118900 | -2.27959000 |
| H | -4.19115100 | -6.31388000 | -6.01892200 |
| H | -2.16652400 | -9.14182000 | -2.35604700 |
| H | -0.51021700 | -3.81344400 | -5.76871400 |
| H | 4.64125800 | -7.19825300 | -2.02325400 |
| H | 5.88954800 | -6.25229200 | -2.89799900 |
| H | 4.31620300 | 0.79057600 | -4.54330700 |
| H | 4.35784800 | 2.52013100 | -5.00954000 |
| H | 11.49201600 | 2.98652900 | -1.64776000 |
| H | 11.13311000 | 4.72175200 | 0.48995300 |
| H | 8.30678300 | 4.73982700 | 4.60489500 |
| H | 7.59826900 | 3.10932800 | 4.55646200 |
| H | 6.00237000 | 6.23935100 | 0.49033700 |
| H | -6.68282200 | 0.10871700 | 4.97957200 |
| H | 4.89916300 | -6.59533800 | 3.05584200 |
| H | 8.26422200 | -1.63718400 | 4.56554000 |
| H | 6.61311800 | -1.95635400 | 3.96718400 |
| H | -2.89215400 | -5.08649800 | 4.86443100 |
| H | -7.84183200 | 6.61704700 | -3.30596100 |

#### Model5 − Intermediate

| | | | |
|---|---|---|---|
| O | -3.95224700 | -1.71854100 | -3.89467900 |
| H | -4.06272600 | -1.85494300 | -4.85355800 |
| H | -4.32810000 | -2.55940500 | -3.54819900 |
| O | -2.07137300 | 2.56492700 | 2.21799800 |
| H | -2.23182800 | 4.07953600 | 1.67684400 |
| H | -2.43862200 | 2.09800400 | 3.01866600 |
| O | -2.29155900 | -0.01011000 | -2.41933200 |
| H | -2.83727400 | -0.60353000 | -2.99132000 |
| H | -1.57539500 | 0.37004700 | -2.97198200 |
| O | -1.04491200 | -6.11654500 | 3.03437700 |
| H | -1.14801900 | -7.07341000 | 3.12381900 |
| H | -1.90803800 | -5.73585300 | 3.35182700 |
| O | -6.32700000 | 1.58366300 | 3.63610700 |
| H | -7.03491000 | 2.18777100 | 3.89863800 |
| H | -5.72290000 | 1.54565100 | 4.45867200 |
| O | 0.20490100 | 0.77758700 | -3.33070300 |
| H | 0.71903400 | 0.50468900 | -2.52383000 |
| H | 0.57726700 | 1.62972900 | -3.59146900 |
| C | -7.60077800 | 5.70153800 | -2.91400100 |
| C | -6.23400000 | 6.41946500 | -2.99108800 |
| O | -5.55975100 | 6.64706100 | -1.97852200 |
| C | -7.48357500 | 4.22022700 | -3.32312000 |
| C | -6.76395900 | 3.31456800 | -2.30796200 |
| C | -7.60834700 | 3.13298300 | -1.04197000 |
| C | -6.42070700 | 1.96414700 | -2.94382600 |
| H | -7.96095900 | 5.80919500 | -1.88308400 |
| H | -6.96786700 | 4.16191500 | -4.29762900 |
| H | -8.49848900 | 3.82165400 | -3.49463200 |
| H | -5.81171700 | 3.79432600 | -2.01762900 |
| H | -7.11423800 | 2.45343100 | -0.33604600 |
| H | -8.59686900 | 2.71098200 | -1.29169200 |
| H | -7.76227600 | 4.08234900 | -0.50625200 |
| H | -7.33299300 | 1.42449300 | -3.24869500 |
| H | -5.78912300 | 2.08770800 | -3.83815900 |
| H | -5.87045900 | 1.32097500 | -2.24256500 |
| N | -3.77190400 | 8.08042100 | 0.16571300 |
| C | -2.34372300 | 7.73895800 | 0.35555500 |
| C | -1.89625000 | 6.55403900 | -0.48593500 |
| C | -2.02829300 | 5.14852600 | 0.06590700 |
| O | -2.26959200 | 5.04292900 | 1.34738300 |
| O | -1.82288500 | 4.19286600 | -0.67852800 |
| H | -4.24795500 | 7.71531800 | -0.65863800 |
| H | -2.16379900 | 7.54569200 | 1.42261800 |
| H | -0.81437300 | 6.64703600 | -0.68725500 |
| H | -2.37547200 | 6.54593100 | -1.47782500 |
| C | -7.50107000 | -1.31144600 | -1.28372200 |
| C | -6.22411600 | -1.99632800 | -0.80438600 |
| C | -5.97244200 | -1.84431300 | 0.70003300 |
| C | -5.81207500 | -0.39741400 | 1.16704700 |
| N | -4.70382000 | 0.20895800 | 0.45046000 |
| C | -4.43918700 | 1.51857500 | 0.34515700 |
| N | -5.07239200 | 2.40770800 | 1.11109900 |
| N | -3.55972300 | 1.89610200 | -0.59054200 |
| H | -7.47109100 | -0.22330200 | -1.12092700 |
| H | -5.36964400 | -1.58995600 | -1.36942300 |
| H | -6.25509600 | -3.06639000 | -1.06206700 |
| H | -5.05805000 | -2.39875400 | 0.97207000 |
| H | -6.79883300 | -2.29840500 | 1.27317700 |
| H | -5.66254800 | -0.35932600 | 2.25686800 |
| H | -6.72412800 | 0.18791000 | 0.97356200 |
| H | -3.91941800 | -0.40994900 | 0.21613400 |
| H | -5.53049800 | 2.11337400 | 1.98554700 |
| H | -5.02045900 | 3.40643000 | 0.87980200 |
| H | -3.17928200 | 1.19630200 | -1.24204100 |
| H | -3.04362200 | 2.77260900 | -0.50563200 |
| C | -1.38130000 | -8.38953100 | -2.02199500 |
| C | -1.13371100 | -7.74926600 | -0.65470900 |
| C | -2.37803100 | -7.05921300 | -0.07963900 |
| N | -2.73447700 | -5.84094100 | -0.77926500 |
| C | -2.59025100 | -4.59690400 | -0.30077400 |
| N | -1.92515000 | -4.34385000 | 0.84473300 |
| N | -3.10580000 | -3.58395600 | -0.98930700 |
| H | -1.85910600 | -7.68432300 | -2.71694800 |
| H | -0.44398300 | -8.74749400 | -2.47634700 |
| H | -0.32347600 | -7.00259000 | -0.71557900 |
| H | -0.79531900 | -8.51825900 | 0.06301700 |
| H | -2.22899300 | -6.83670200 | 0.98503400 |
| H | -3.23633200 | -7.75147400 | -0.13762500 |
| H | -3.17641700 | -5.94629400 | -1.72695000 |
| H | -1.48586700 | -5.07743800 | 1.39386000 |
| H | -2.02314300 | -3.43172900 | 1.29373600 |
| H | -3.74713900 | -3.79590500 | -1.78661700 |
| H | -2.91872000 | -2.63267600 | -0.65962800 |
| C | -4.74320300 | -5.62845400 | -6.16708300 |
| C | -3.85529100 | -4.37182800 | -6.17484800 |
| O | -4.32402900 | -3.24940300 | -6.34139400 |
| C | -5.55533200 | -5.59864500 | -4.84993400 |
| C | -4.66239200 | -5.35390000 | -3.61948000 |
| O | -4.75545700 | -4.21952800 | -3.06159900 |
| O | -3.88203100 | -6.26916100 | -3.25981700 |
| H | -5.41970400 | -5.59251900 | -7.03268900 |
| H | -6.07796500 | -6.55980500 | -4.72368000 |
| H | -6.29603200 | -4.78994300 | -4.90418200 |
| N | -2.54578500 | -4.59707300 | -5.87658700 |

| | | | |
|---|---|---|---|
| C | -1.58787500 | -3.57308200 | -5.46431000 |
| C | -1.19014500 | -3.73461900 | -3.98891200 |
| C | -0.28157200 | -2.59435000 | -3.50296400 |
| C | -0.60870500 | -5.12232700 | -3.70669200 |
| C | 1.10271700 | -2.48486800 | -4.14260300 |
| H | -2.29925500 | -5.55531200 | -5.66294100 |
| H | -2.06582700 | -2.59901400 | -5.62273600 |
| H | -2.12453800 | -3.64485800 | -3.41945300 |
| H | -0.16828000 | -2.69839000 | -2.40990900 |
| H | -0.81042300 | -1.64335200 | -3.64473700 |
| H | 0.30205800 | -5.32167800 | -4.29697400 |
| H | -1.34317300 | -5.91201700 | -3.91520600 |
| H | -0.35620700 | -5.22469100 | -2.64279700 |
| H | 1.64881800 | -1.63421700 | -3.70836800 |
| H | 1.04092300 | -2.30983800 | -5.22842600 |
| H | 1.71111100 | -3.39163800 | -3.98969300 |
| C | 4.86822900 | -6.06968700 | -2.52383300 |
| C | 4.82486800 | -4.93297300 | -1.51261700 |
| O | 5.82806600 | -4.32695300 | -1.15839100 |
| H | 4.22093300 | -5.85641400 | -3.38912700 |
| N | 3.59451100 | -4.62168200 | -0.99437500 |
| C | 3.46522500 | -3.47646900 | -0.09058400 |
| C | 3.97494900 | -3.76277400 | 1.32409700 |
| O | 4.38191500 | -2.85940400 | 2.04752900 |
| C | 1.95128500 | -3.18366700 | -0.09566400 |
| C | 1.31865300 | -4.54273800 | -0.40374100 |
| C | 2.29294300 | -5.16465700 | -1.40955200 |
| H | 4.05406600 | -2.62509000 | -0.45756300 |
| H | 1.73746200 | -2.46523900 | -0.89921500 |
| H | 1.61318900 | -2.73904700 | 0.84749500 |
| H | 0.30035200 | -4.45440200 | -0.79857000 |
| H | 1.25874600 | -5.15476000 | 0.51002600 |
| H | 2.04751700 | -4.85405900 | -2.43812000 |
| H | 2.28826800 | -6.26563000 | -1.38816400 |
| N | 3.87841000 | -5.04991900 | 1.72435000 |
| C | 4.17522100 | -5.50756700 | 3.06500000 |
| C | 2.92100600 | -5.74379000 | 3.91081600 |
| C | 2.20738600 | -4.48193000 | 4.28253800 |
| C | 2.65207400 | -3.22119200 | 4.63728100 |
| N | 0.83695700 | -4.43062200 | 4.43926700 |
| C | 0.52297300 | -3.17579600 | 4.86726100 |
| N | 1.59406100 | -2.42064300 | 4.99619400 |
| H | 3.57514800 | -5.72428500 | 1.03337700 |
| H | 4.81538600 | -4.74815800 | 3.53437700 |
| H | 2.23447700 | -6.41033100 | 3.35919200 |
| H | 0.17526500 | -5.14062300 | 4.11358900 |
| H | 3.67318400 | -2.84810200 | 4.62565600 |
| H | -0.50083100 | -2.87368200 | 5.05796100 |
| H | 3.22044500 | -6.30169600 | 4.81820600 |
| C | 7.50992100 | 4.37286700 | 3.73788000 |
| C | 6.24551900 | 4.69993200 | 2.96081500 |
| O | 5.13468600 | 4.34501900 | 3.32711300 |
| H | 8.36476500 | 4.20605800 | 3.06470300 |
| N | 6.42625400 | 5.48206200 | 1.84774700 |
| C | 5.27405900 | 5.77307500 | 1.03517900 |
| C | 4.75987000 | 4.50068500 | 0.30727300 |
| O | 5.43172300 | 3.46523800 | 0.29262300 |
| H | 7.34065500 | 5.44401500 | 1.38877200 |
| H | 4.45963300 | 6.16943000 | 1.65908200 |
| N | 3.55408700 | 4.55092700 | -0.29278500 |
| C | 3.04459600 | 3.41316800 | -1.03716000 |
| C | 3.68366600 | 3.37312000 | -2.45350100 |
| O | 4.27490300 | 4.32866100 | -2.91954300 |
| C | 1.50986400 | 3.36382300 | -0.97983700 |
| C | 1.01954300 | 3.18263600 | 0.45581300 |
| O | 0.98635800 | 4.54381600 | -1.57124100 |
| H | 3.05699500 | 5.42851600 | -0.40347700 |
| H | 3.38009500 | 2.50731700 | -0.51624100 |
| H | 1.19894300 | 2.48049700 | -1.55874100 |
| H | 0.01623500 | 4.46666700 | -1.52524800 |
| H | 1.20370800 | 4.09375900 | 1.04506900 |
| H | 1.52037900 | 2.33545700 | 0.94310600 |
| H | -0.06087400 | 2.98628900 | 0.44362400 |
| N | 3.60050600 | 2.16653400 | -3.09804700 |
| C | 4.30491000 | 1.92138800 | -4.36249800 |
| H | 3.12122300 | 1.39884200 | -2.63368600 |
| H | 5.38866500 | 1.79738300 | -4.19631800 |
| C | -3.94019000 | -3.85810500 | 3.75118100 |
| O | -3.32846800 | -5.13181500 | 3.98604100 |
| C | -3.06922900 | -2.69552200 | 4.17102500 |
| O | -2.02979100 | -2.47217200 | 3.23473500 |
| C | -3.85115000 | -1.38121700 | 4.33948500 |
| O | -4.03948400 | -1.20206500 | 5.72036600 |
| C | -2.96691100 | -0.33873400 | 3.57565600 |
| O | -2.77549500 | 0.89488000 | 4.22084800 |
| C | -1.62813000 | -1.08935400 | 3.34207400 |
| N | 1.35971500 | -0.08691500 | 1.19240900 |
| C | 0.84613600 | -0.12651000 | -0.05651800 |
| O | 1.52239800 | 0.14051600 | -1.08468000 |
| N | -0.48469400 | -0.50480200 | -0.20419700 |
| C | -1.32732500 | -0.87714400 | 0.82579800 |
| O | -2.45960400 | -1.34161100 | 0.54611600 |
| C | -0.79396000 | -0.73165800 | 2.14501100 |
| C | 0.55157800 | -0.37718800 | 2.22746100 |
| H | -1.18314500 | 2.19048000 | 2.13180600 |
| H | -0.93957600 | -0.49875700 | -1.13055300 |
| H | 1.02141800 | -0.35970300 | 3.21909000 |
| H | -1.02253300 | -0.94157300 | 4.25460800 |
| H | -3.51864900 | 1.05307200 | 4.88331700 |
| H | -4.83222600 | -1.44895300 | 3.84185300 |
| H | -2.64226700 | -2.90470300 | 5.17283500 |
| H | -4.89548200 | -3.80367900 | 4.29865100 |
| H | -3.43387600 | -0.18642100 | 2.59156900 |
| H | -4.15143600 | -3.80123700 | 2.67432000 |
| P | -5.28444600 | -0.24304400 | 6.32348100 |
| O | -4.93215300 | 1.15368200 | 5.73422200 |
| O | -5.44240100 | -0.52351100 | 7.76434000 |
| O | -6.55825200 | -0.76795200 | 5.41775700 |
| C | 7.38730300 | -1.03968400 | 3.65297000 |
| C | 7.74165300 | -1.41917800 | 2.21367900 |
| C | 6.64233500 | -1.16875200 | 1.18543800 |
| C | 4.69418800 | 0.85328300 | 1.69452300 |
| S | 6.31403800 | 0.62449500 | 0.89423000 |
| C | 5.89818000 | 0.59041500 | -0.89331800 |
| C | 7.09682600 | 0.49329200 | -1.85555800 |
| O | 7.39464500 | 1.77476500 | -2.40542400 |
| C | 8.44412900 | -0.03815100 | -1.33217600 |
| O | 8.47266000 | -1.44462600 | -1.43100800 |
| C | 9.44333500 | 0.63334200 | -2.29351000 |
| O | 9.46451000 | -0.05168800 | -3.51366300 |
| C | 8.78457000 | 1.99793200 | -2.51018200 |
| C | 10.42344400 | 3.52752300 | -1.29404000 |
| C | 10.27951200 | 4.41648300 | -0.25140700 |
| N | 8.97645400 | 4.42330700 | 0.18952300 |
| C | 8.33062300 | 3.55594500 | -0.56732800 |
| N | 9.16731400 | 2.97601600 | -1.48535100 |
| H | 7.22192700 | 0.04599700 | 3.75858200 |
| H | 7.94378600 | -2.50306900 | 2.16322500 |
| H | 8.67770500 | -0.91782200 | 1.90531000 |
| H | 6.90955300 | -1.58616600 | 0.20678900 |
| H | 5.68483500 | -1.61883200 | 1.49244300 |
| H | 4.80397500 | 0.42079100 | 2.69699800 |
| H | 4.53682400 | 1.93580000 | 1.75537100 |
| H | 3.88586900 | 0.33562000 | 1.15331500 |
| H | 7.26757300 | 3.33904000 | -0.50056300 |
| H | 9.01889500 | 2.39628500 | -3.50863000 |
| H | 9.30575900 | -0.98867200 | -3.31159100 |
| H | 8.63260700 | 0.31090400 | -0.30296100 |
| H | 6.78562400 | -0.17503600 | -2.67643700 |
| H | 5.20271100 | -0.24943800 | -1.02492400 |

| | | | |
|---|---|---|---|
| H | 9.13317400 | -1.80528100 | -0.82436000 |
| H | 10.45135600 | 0.73218300 | -1.84873400 |
| H | 5.37059300 | 1.53589100 | -1.07017700 |
| C | -5.70630000 | 9.68235500 | 0.38333600 |
| C | -4.28460600 | 9.21438400 | 0.77163400 |
| O | -3.63254200 | 9.83417400 | 1.59298900 |
| H | -6.28373100 | 9.80975100 | 1.30948800 |
| N | -5.79045400 | 6.75707900 | -4.23290300 |
| C | -4.49112000 | 7.36264000 | -4.56158100 |
| H | -6.40015700 | 6.53093900 | -5.01067800 |
| H | -3.97428300 | 7.58972300 | -3.62100100 |
| O | -5.06052400 | 5.16396100 | 0.32143500 |
| H | -4.67936100 | 5.87378900 | 0.85879100 |
| H | -5.38883200 | 5.61980900 | -0.47954700 |
| H | -5.62034300 | 10.66833100 | -0.09707300 |
| H | -6.22534600 | 8.98471100 | -0.29041900 |
| H | -1.73477200 | 8.62258000 | 0.10005900 |
| H | -3.87350000 | 6.66767600 | -5.15192600 |
| H | -4.63300900 | 8.29501200 | -5.12878500 |
| H | -8.38940800 | -1.70234600 | -0.75856600 |
| H | -7.64854200 | -1.47365600 | -2.36271600 |
| H | -4.13668900 | -6.54507700 | -6.22223200 |
| H | -2.05999300 | -9.25289800 | -1.92745500 |
| H | -0.69984800 | -3.62439600 | -6.11879000 |
| H | 4.51381100 | -7.01127800 | -2.07507100 |
| H | 5.90401800 | -6.19374400 | -2.86025400 |
| H | 3.89305800 | 1.01852300 | -4.83329600 |
| H | 4.16553900 | 2.78036800 | -5.03342800 |
| H | 11.27871100 | 3.25426400 | -1.90690100 |
| H | 11.04450100 | 5.04851900 | 0.19717000 |
| H | 7.75518100 | 5.21656900 | 4.40284100 |
| H | 7.32649200 | 3.48537900 | 4.35698200 |
| H | 5.53412200 | 6.52636300 | 0.27525400 |
| H | -6.72889500 | -0.12457500 | 4.70408100 |
| H | 4.76119300 | -6.43901200 | 2.99995500 |
| H | 8.19698900 | -1.31492300 | 4.34467000 |
| H | 6.46815600 | -1.55599900 | 3.97126900 |
| H | -3.30619300 | -5.27008400 | 4.94379800 |
| H | -8.31248900 | 6.21973600 | -3.57766000 |

#### Model5 – TS2

| | | | |
|---|---|---|---|
| O | -3.92114700 | -1.65525300 | -3.87407300 |
| H | -4.02242000 | -1.77372700 | -4.83645900 |
| H | -4.30932900 | -2.49938700 | -3.54847700 |
| O | -1.98516400 | 2.61331700 | 2.27909300 |
| H | -2.18915300 | 4.13390100 | 1.72720400 |
| H | -2.37177500 | 2.12902300 | 3.05979300 |
| O | -2.24054900 | 0.00073700 | -2.38670000 |
| H | -2.79986200 | -0.58253100 | -2.95877500 |
| H | -1.54295800 | 0.39808200 | -2.95212400 |
| O | -1.11805200 | -6.18679700 | 2.98027600 |
| H | -1.27310200 | -7.13938100 | 3.03492000 |
| H | -1.96146100 | -5.77089500 | 3.30425200 |
| O | -6.27590100 | 1.66864500 | 3.62704100 |
| H | -6.97329100 | 2.28362100 | 3.89278600 |
| H | -5.67506100 | 1.61018900 | 4.45226500 |
| O | 0.21279200 | 0.83090500 | -3.34898900 |
| H | 0.74037900 | 0.50259600 | -2.58008300 |
| H | 0.56487400 | 1.71049400 | -3.53967400 |
| C | -7.52399400 | 5.79255900 | -2.93166400 |
| C | -6.15062100 | 6.49841200 | -2.99949800 |
| O | -5.47456100 | 6.70522200 | -1.98358300 |
| C | -7.41231700 | 4.30847200 | -3.33244500 |
| C | -6.69432400 | 3.40502300 | -2.31415100 |
| C | -7.53293900 | 3.23721500 | -1.04244100 |
| C | -6.36309500 | 2.04822500 | -2.94286300 |
| H | -7.89273500 | 5.90749300 | -1.90450900 |
| H | -6.89789800 | 4.24369900 | -4.30719300 |
| H | -8.42892100 | 3.91274000 | -3.50060600 |
| H | -5.73753200 | 3.88029600 | -2.03186600 |
| H | -7.04071900 | 2.55681100 | -0.33583000 |
| H | -8.52645700 | 2.82227300 | -1.28394600 |
| H | -7.67538900 | 4.19040100 | -0.51062500 |
| H | -7.28019100 | 1.51339400 | -3.24150800 |
| H | -5.73397400 | 2.16163500 | -3.84026700 |
| H | -5.81561200 | 1.40509600 | -2.23916500 |
| N | -3.69699600 | 8.12737700 | 0.18258500 |
| C | -2.27016800 | 7.77948700 | 0.37396500 |
| C | -1.83147600 | 6.58251300 | -0.45708200 |
| C | -1.96223500 | 5.18428100 | 0.11159500 |
| O | -2.22336700 | 5.09249700 | 1.39061900 |
| O | -1.74461300 | 4.21799400 | -0.61621600 |
| H | -4.16593100 | 7.77971600 | -0.65331800 |
| H | -2.09125600 | 7.59717500 | 1.44289600 |
| H | -0.75183500 | 6.66985400 | -0.67213900 |
| H | -2.32072800 | 6.56264000 | -1.44404300 |
| C | -7.47969600 | -1.22835500 | -1.33337400 |
| C | -6.20739700 | -1.90904800 | -0.83537300 |
| C | -5.97638700 | -1.75947200 | 0.67300000 |
| C | -5.80278700 | -0.31594700 | 1.14696000 |
| N | -4.67470400 | 0.27687700 | 0.44948700 |
| C | -4.38108000 | 1.58186400 | 0.35434300 |
| N | -4.99601100 | 2.47951200 | 1.12419100 |
| N | -3.48760900 | 1.94535900 | -0.57311200 |
| H | -7.45953000 | -0.14125200 | -1.16199000 |
| H | -5.34627200 | -1.50034700 | -1.38887600 |
| H | -6.23256800 | -2.97902700 | -1.09437100 |
| H | -5.07416400 | -2.32718500 | 0.95888900 |
| H | -6.81669900 | -2.20339400 | 1.23345800 |
| H | -5.66874600 | -0.28340800 | 2.23904600 |
| H | -6.70328700 | 0.28298900 | 0.94068500 |
| H | -3.90885900 | -0.35513300 | 0.20196600 |
| H | -5.47233400 | 2.18954500 | 1.99098100 |
| H | -4.93904800 | 3.47758200 | 0.88796000 |
| H | -3.12156800 | 1.24139100 | -1.22722400 |
| H | -2.95564700 | 2.81273700 | -0.48171500 |
| C | -1.40387700 | -8.34403300 | -2.07329500 |
| C | -1.25891500 | -7.82064000 | -0.64328900 |
| C | -2.49143600 | -7.05610400 | -0.14402300 |
| N | -2.73029600 | -5.80994500 | -0.84851900 |
| C | -2.60211600 | -4.57994000 | -0.33103400 |
| N | -1.96266200 | -4.35937100 | 0.83549400 |
| N | -3.10251500 | -3.54980100 | -1.00635000 |
| H | -1.70388600 | -7.54651100 | -2.76506600 |
| H | -0.46255600 | -8.78614100 | -2.43596300 |
| H | -0.38125600 | -7.15868000 | -0.56265900 |
| H | -1.07449900 | -8.66379300 | 0.04665300 |
| H | -2.39741300 | -6.85355900 | 0.93139900 |
| H | -3.38596700 | -7.69121700 | -0.26876500 |
| H | -3.16709700 | -5.89069400 | -1.80293700 |
| H | -1.53033000 | -5.11033400 | 1.36776600 |
| H | -2.05622100 | -3.45880400 | 1.30770100 |
| H | -3.73947600 | -3.74622400 | -1.81259900 |
| H | -2.91674000 | -2.60570100 | -0.66086800 |
| C | -4.72592100 | -5.54135600 | -6.22258500 |
| C | -3.83343800 | -4.28612300 | -6.21330600 |
| O | -4.30563800 | -3.15901500 | -6.32885900 |
| C | -5.54216600 | -5.51092900 | -4.90721200 |
| C | -4.65350600 | -5.28526400 | -3.66976300 |
| O | -4.74394700 | -4.15908000 | -3.09307500 |
| O | -3.87829900 | -6.20827300 | -3.31993400 |
| H | -5.40016300 | -5.49692300 | -7.08958400 |
| H | -6.07582900 | -6.46698800 | -4.78868000 |
| H | -6.27356500 | -4.69386800 | -4.95999300 |
| N | -2.51606900 | -4.51833400 | -5.95371800 |
| C | -1.56052700 | -3.51054100 | -5.49428300 |
| C | -1.20088600 | -3.71627900 | -4.01556300 |
| C | -0.29673700 | -2.60214400 | -3.46617700 |
| C | -0.63971400 | -5.11707500 | -3.76744700 |
| C | 1.13966500 | -2.54672000 | -3.98644900 |

| | | | |
|---|---|---|---|
| H | -2.26114700 | -5.48447900 | -5.79283400 |
| H | -2.02996300 | -2.53006200 | -5.63820600 |
| H | -2.14997500 | -3.63615300 | -3.46977300 |
| H | -0.27936800 | -2.70494700 | -2.36732400 |
| H | -0.77965400 | -1.63454800 | -3.65489000 |
| H | 0.27703700 | -5.30822100 | -4.35095800 |
| H | -1.38191200 | -5.88793300 | -4.01334000 |
| H | -0.40530400 | -5.25621500 | -2.70438000 |
| H | 1.67506300 | -1.70508100 | -3.52205500 |
| H | 1.17771700 | -2.39779800 | -5.07730600 |
| H | 1.70428600 | -3.46560000 | -3.75874400 |
| C | 4.86357100 | -6.06404200 | -2.53260500 |
| C | 4.70201600 | -4.90561000 | -1.55757300 |
| O | 5.62764000 | -4.16542200 | -1.25217500 |
| H | 4.18077600 | -5.95829000 | -3.39045000 |
| N | 3.45540500 | -4.73558300 | -1.00875100 |
| C | 3.21348900 | -3.60010900 | -0.11677700 |
| C | 3.74135400 | -3.82679600 | 1.30187500 |
| O | 4.04272100 | -2.88037900 | 2.02281000 |
| C | 1.67924800 | -3.45385200 | -0.12583500 |
| C | 1.18449300 | -4.87585900 | -0.40072500 |
| C | 2.21853600 | -5.43404800 | -1.38538700 |
| H | 3.71854800 | -2.70039000 | -0.49012500 |
| H | 1.39603300 | -2.78087200 | -0.94658400 |
| H | 1.29181100 | -3.03117100 | 0.80937600 |
| H | 0.16739800 | -4.89797300 | -0.80539000 |
| H | 1.17661100 | -5.46234900 | 0.53117100 |
| H | 1.93704900 | -5.20179200 | -2.42524500 |
| H | 2.33099100 | -6.52736000 | -1.31206000 |
| N | 3.78665200 | -5.11288700 | 1.71179100 |
| C | 4.14578700 | -5.52295000 | 3.05516200 |
| C | 2.93479000 | -5.81147500 | 3.94714200 |
| C | 2.16618200 | -4.58277200 | 4.32075200 |
| C | 2.55908300 | -3.30985700 | 4.69161200 |
| N | 0.79078300 | -4.57704100 | 4.43998200 |
| C | 0.42412400 | -3.33479100 | 4.86168900 |
| N | 1.46582000 | -2.54446900 | 5.01998500 |
| H | 3.58094100 | -5.82299400 | 1.02064500 |
| H | 4.75591700 | -4.72052000 | 3.49222400 |
| H | 2.26861900 | -6.52502900 | 3.43136700 |
| H | 0.15539000 | -5.30194900 | 4.09613400 |
| H | 3.56717400 | -2.90279500 | 4.71247500 |
| H | -0.61338400 | -3.06798600 | 5.02538600 |
| H | 3.29617700 | -6.33374700 | 4.85277500 |
| C | 7.54323000 | 4.33159100 | 3.79071300 |
| C | 6.29655400 | 4.90321700 | 3.12201500 |
| O | 5.19799900 | 4.83640600 | 3.65099500 |
| H | 8.47903900 | 4.69400100 | 3.34180700 |
| N | 6.48670800 | 5.48757000 | 1.89650300 |
| C | 5.33068000 | 5.75924500 | 1.08317400 |
| C | 4.77442300 | 4.44818900 | 0.46324300 |
| O | 5.34347600 | 3.37204800 | 0.62486100 |
| H | 7.36117200 | 5.30174100 | 1.38903500 |
| H | 4.53999900 | 6.21716800 | 1.69549100 |
| N | 3.62286400 | 4.53983200 | -0.25366600 |
| C | 3.12751600 | 3.39758100 | -0.99200000 |
| C | 3.75768600 | 3.36910900 | -2.40994600 |
| O | 4.36642000 | 4.32093600 | -2.86050700 |
| C | 1.59448500 | 3.30281800 | -0.94009300 |
| C | 1.09558400 | 3.14490200 | 0.49548000 |
| O | 1.03853600 | 4.44727900 | -1.56725400 |
| H | 3.25850300 | 5.44721900 | -0.52221300 |
| H | 3.51044000 | 2.51235600 | -0.46907300 |
| H | 1.31068900 | 2.39510700 | -1.49730900 |
| H | 0.07067000 | 4.37791800 | -1.47765400 |
| H | 1.27754500 | 4.06755300 | 1.06630700 |
| H | 1.60435100 | 2.31401300 | 1.00142000 |
| H | 0.01655300 | 2.93690000 | 0.48687200 |
| N | 3.64659600 | 2.17890700 | -3.08171700 |
| C | 4.36331000 | 1.93900000 | -4.33712400 |
| H | 3.15447900 | 1.41002300 | -2.63659000 |
| H | 5.42969800 | 1.72383200 | -4.15206400 |
| C | -3.95860000 | -3.81953500 | 3.71681800 |
| O | -3.35982500 | -5.10089100 | 3.93976000 |
| C | -3.07413400 | -2.67239500 | 4.14742500 |
| O | -2.02305900 | -2.46561700 | 3.21895800 |
| C | -3.83075600 | -1.34492400 | 4.31509700 |
| O | -4.05019300 | -1.18162400 | 5.69260800 |
| C | -2.90378100 | -0.31247900 | 3.58207500 |
| O | -2.69942400 | 0.90480500 | 4.25230300 |
| C | -1.57892400 | -1.09945000 | 3.35152500 |
| N | 1.42064900 | -0.22065700 | 1.14835200 |
| C | 0.88326300 | -0.22052200 | -0.09826400 |
| O | 1.55399700 | 0.02114500 | -1.12995600 |
| N | -0.46100200 | -0.52980200 | -0.21027100 |
| C | -1.29356500 | -0.88166600 | 0.83538700 |
| O | -2.43937600 | -1.30639300 | 0.57691900 |
| C | -0.72855800 | -0.76487000 | 2.15226400 |
| C | 0.62978400 | -0.48269700 | 2.21562800 |
| H | -1.04849100 | 2.37330700 | 2.31303100 |
| H | -0.93709200 | -0.49728400 | -1.13000900 |
| H | 1.12952500 | -0.50096100 | 3.19158600 |
| H | -0.97087500 | -0.98439900 | 4.26693400 |
| H | -3.45287300 | 1.06489500 | 4.90245700 |
| H | -4.79938700 | -1.38328900 | 3.79051200 |
| H | -2.65810200 | -2.89060400 | 5.15163100 |
| H | -4.91300000 | -3.75752900 | 4.26496700 |
| H | -3.35287800 | -0.13342600 | 2.59387200 |
| H | -4.17019100 | -3.75030900 | 2.64073600 |
| P | -5.29481500 | -0.20988700 | 6.28982700 |
| O | -4.90115600 | 1.18523100 | 5.72285200 |
| O | -5.48191800 | -0.50942100 | 7.72277900 |
| O | -6.55714800 | -0.70220100 | 5.35164500 |
| C | 7.38471300 | -1.07956300 | 3.68016800 |
| C | 7.79224600 | -1.36418200 | 2.23142900 |
| C | 6.63779000 | -1.40010400 | 1.22653400 |
| C | 3.66302300 | 0.06158100 | 1.05225100 |
| S | 5.99864600 | 0.27455400 | 0.74533600 |
| C | 6.01620200 | 0.10226400 | -1.08770800 |
| C | 7.35535400 | 0.25362900 | -1.83224000 |
| O | 7.47910700 | 1.56808800 | -2.38191600 |
| C | 8.66766300 | 0.04248400 | -1.07097200 |
| O | 8.96536500 | -1.33412400 | -0.96461100 |
| C | 9.66805800 | 0.77937100 | -1.98215700 |
| O | 9.98342400 | -0.02589600 | -3.08588000 |
| C | 8.83376800 | 1.97685400 | -2.46166300 |
| C | 10.06619000 | 4.03523900 | -1.65325500 |
| C | 9.84301700 | 4.92932300 | -0.63046500 |
| N | 8.67819500 | 4.61060000 | 0.02604800 |
| C | 8.19371200 | 3.54027400 | -0.57623700 |
| N | 9.00305500 | 3.15110800 | -1.60988300 |
| H | 6.92700200 | -0.08107500 | 3.77775500 |
| H | 8.27058700 | -2.35925700 | 2.18335500 |
| H | 8.55989900 | -0.63726500 | 1.90968700 |
| H | 6.94364000 | -1.88938600 | 0.29403800 |
| H | 5.78727700 | -1.96087600 | 1.63241300 |
| H | 3.74071700 | -0.90705800 | 1.54678900 |
| H | 3.59545600 | 0.98143600 | 1.62932000 |
| H | 3.45355100 | 0.10372700 | -0.01369000 |
| H | 7.25264600 | 3.05490000 | -0.32608600 |
| H | 9.09836700 | 2.24335200 | -3.49626200 |
| H | 9.98070500 | -0.94282700 | -2.76383600 |
| H | 8.61011700 | 0.53132700 | -0.08498300 |
| H | 7.34116500 | -0.47616300 | -2.66115000 |
| H | 5.58515200 | -0.88473600 | -1.30639000 |
| H | 9.49665000 | -1.48598200 | -0.17240900 |
| H | 10.57124000 | 1.11672100 | -1.44024100 |
| H | 5.32947300 | 0.87290600 | -1.46535700 |
| C | -5.61973100 | 9.74533200 | 0.39365400 |
| C | -4.20659500 | 9.26121600 | 0.79378500 |

| | | | |
|---|---|---|---|
| O | -3.55822000 | 9.86755800 | 1.62729100 |
| H | -6.20191400 | 9.88375800 | 1.31512900 |
| N | -5.70165000 | 6.84540200 | -4.23648500 |
| C | -4.39487500 | 7.44023100 | -4.55571100 |
| H | -6.31465100 | 6.64016900 | -5.01747200 |
| H | -3.86990200 | 7.63403800 | -3.61226800 |
| O | -4.98395700 | 5.22741400 | 0.31870900 |
| H | -4.61406600 | 5.94572000 | 0.85327000 |
| H | -5.31075500 | 5.67569800 | -0.48740000 |
| H | -5.51801600 | 10.72816700 | -0.09025900 |
| H | -6.14253700 | 9.05188600 | -0.28158100 |
| H | -1.65470400 | 8.65551600 | 0.10808800 |
| H | -3.79155100 | 6.75181200 | -5.16792200 |
| H | -4.52473800 | 8.38964700 | -5.09700700 |
| H | -8.37468200 | -1.62880800 | -0.82735900 |
| H | -7.60577800 | -1.38394000 | -2.41594300 |
| H | -4.12367600 | -6.46048300 | -6.28014700 |
| H | -2.18407700 | -9.12044800 | -2.12681400 |
| H | -0.65875400 | -3.55003000 | -6.13037200 |
| H | 4.63179500 | -7.02461300 | -2.04575100 |
| H | 5.90056500 | -6.07449500 | -2.88727500 |
| H | 3.89845200 | 1.09449700 | -4.86419700 |
| H | 4.30472600 | 2.83912800 | -4.96397800 |
| H | 10.85738700 | 3.95189800 | -2.39410000 |
| H | 10.45247900 | 5.78419900 | -0.34195800 |
| H | 7.51279700 | 4.57113700 | 4.86223000 |
| H | 7.50708300 | 3.23498400 | 3.68775900 |
| H | 5.59406700 | 6.44801300 | 0.26486200 |
| H | -6.71256400 | -0.03970200 | 4.65166800 |
| H | 4.78142400 | -6.42041900 | 2.98723700 |
| H | 8.25471900 | -1.11469400 | 4.35299800 |
| H | 6.64648600 | -1.81877500 | 4.02926600 |
| H | -3.35139600 | -5.25498300 | 4.89531900 |
| H | -8.22659100 | 6.31134600 | -3.60434500 |

#### Model5 – Final state

| | | | |
|---|---|---|---|
| O | -4.07972000 | -1.47869100 | -3.96151300 |
| H | -4.02596200 | -1.70859400 | -4.90910100 |
| H | -4.50784600 | -2.28832700 | -3.60319300 |
| O | -1.80607000 | 2.75719400 | 2.42374700 |
| H | -2.05759300 | 4.25433800 | 1.84598000 |
| H | -2.29307000 | 2.19198400 | 3.08645700 |
| O | -2.38269700 | 0.16080200 | -2.44986500 |
| H | -2.93133700 | -0.39153200 | -3.06189000 |
| H | -1.74848900 | 0.70996600 | -2.96226800 |
| O | -1.55594000 | -6.60424200 | 2.80513300 |
| H | -1.85716200 | -7.52206500 | 2.79851300 |
| H | -2.29625300 | -6.08106400 | 3.19986800 |
| O | -6.23219600 | 1.96913700 | 3.72048500 |
| H | -6.86782500 | 2.63246300 | 4.02200800 |
| H | -5.60359300 | 1.84730100 | 4.51498200 |
| O | -0.08086700 | 1.50818000 | -3.10140300 |
| H | 0.54853300 | 0.89308500 | -2.68127100 |
| H | 0.26955500 | 2.39355700 | -2.88827200 |
| C | -7.29980900 | 5.95021800 | -2.97068500 |
| C | -5.91278900 | 6.62987900 | -3.03148000 |
| O | -5.23733700 | 6.82090900 | -2.01162100 |
| C | -7.21245000 | 4.46755400 | -3.38212100 |
| C | -6.49773500 | 3.54707700 | -2.37676500 |
| C | -7.32872300 | 3.38201800 | -1.09972300 |
| C | -6.18954400 | 2.19135400 | -3.01946700 |
| H | -7.66802900 | 6.06491300 | -1.94330700 |
| H | -6.70725600 | 4.40214400 | -4.36150600 |
| H | -8.23567000 | 4.08670900 | -3.54424700 |
| H | -5.53275500 | 4.00777700 | -2.09762900 |
| H | -6.84595200 | 2.68706700 | -0.40155300 |
| H | -8.33081700 | 2.98537600 | -1.33613300 |
| H | -7.45126500 | 4.33243600 | -0.55820900 |
| H | -7.11562800 | 1.67386600 | -3.32082700 |
| H | -5.56186300 | 2.30283800 | -3.91795400 |
| H | -5.65101400 | 1.53152600 | -2.32462300 |
| N | -3.45674500 | 8.22805900 | 0.17335300 |
| C | -2.03638900 | 7.85461300 | 0.36806900 |
| C | -1.64275600 | 6.61129400 | -0.42302000 |
| C | -1.77090500 | 5.24120100 | 0.21268700 |
| O | -2.07491000 | 5.20442800 | 1.48418300 |
| O | -1.53115500 | 4.23777600 | -0.45740700 |
| H | -3.91603700 | 7.90664300 | -0.67850900 |
| H | -1.85643400 | 7.71119300 | 1.44232600 |
| H | -0.57590800 | 6.67629900 | -0.69784500 |
| H | -2.17883200 | 6.55073400 | -1.38407700 |
| C | -7.39370000 | -1.06450300 | -1.34751800 |
| C | -6.23470700 | -1.77798800 | -0.65923200 |
| C | -6.11677400 | -1.47030500 | 0.83909900 |
| C | -5.95980800 | 0.01512200 | 1.16745500 |
| N | -4.80935400 | 0.53317400 | 0.44375900 |
| C | -4.37274600 | 1.79974400 | 0.40817000 |
| N | -4.86681200 | 2.72230900 | 1.23323100 |
| N | -3.44960200 | 2.10675900 | -0.51218300 |
| H | -7.28506900 | 0.02920700 | -1.30353800 |
| H | -5.29910200 | -1.50614400 | -1.17406600 |
| H | -6.32922200 | -2.86618600 | -0.79872800 |
| H | -5.24794000 | -2.01088800 | 1.25184700 |
| H | -7.00604500 | -1.84179800 | 1.37519900 |
| H | -5.85446900 | 0.15966400 | 2.25283900 |
| H | -6.85870900 | 0.58174900 | 0.87410100 |
| H | -4.14299200 | -0.15752600 | 0.09713500 |
| H | -5.39739400 | 2.45333000 | 2.07400000 |
| H | -4.76281400 | 3.71753700 | 0.99526000 |
| H | -3.17123000 | 1.40188700 | -1.20669600 |
| H | -2.82937900 | 2.91016000 | -0.38851400 |
| C | -1.44117600 | -8.29056400 | -2.01597800 |
| C | -1.63633800 | -7.94047000 | -0.53833100 |
| C | -2.94154400 | -7.18970100 | -0.23485100 |
| N | -3.08464300 | -5.90911600 | -0.89926300 |
| C | -2.88422000 | -4.70429000 | -0.35399600 |
| N | -2.22713500 | -4.55405400 | 0.81667100 |
| N | -3.31965700 | -3.62931200 | -1.00273800 |
| H | -1.40299900 | -7.38241100 | -2.63474400 |
| H | -0.50612800 | -8.85109100 | -2.17160700 |
| H | -0.78503900 | -7.34304700 | -0.17206000 |
| H | -1.64067500 | -8.86661200 | 0.06465900 |
| H | -3.04024900 | -7.05251000 | 0.85188600 |
| H | -3.79417300 | -7.81162800 | -0.55549500 |
| H | -3.52993200 | -5.93073800 | -1.85923700 |
| H | -1.87674500 | -5.36215600 | 1.32609400 |
| H | -2.29988600 | -3.67569200 | 1.33252000 |
| H | -3.95750400 | -3.75334400 | -1.82661700 |
| H | -3.11193300 | -2.70006600 | -0.63370100 |
| C | -4.68078600 | -5.44391400 | -6.20039600 |
| C | -3.74363200 | -4.23029400 | -6.16627200 |
| O | -4.15055200 | -3.09324800 | -6.39426300 |
| C | -5.66871600 | -5.27974900 | -5.02621700 |
| C | -4.90053200 | -5.11125300 | -3.71000000 |
| O | -4.96247800 | -3.99006300 | -3.12567400 |
| O | -4.21135400 | -6.09125900 | -3.32501400 |
| H | -5.22009600 | -5.44517400 | -7.15903500 |
| H | -6.30821800 | -6.17350600 | -4.95525600 |
| H | -6.29718200 | -4.39580700 | -5.19636200 |
| N | -2.46872700 | -4.49285100 | -5.77508100 |
| C | -1.48507300 | -3.46756800 | -5.45498000 |
| C | -1.26542800 | -3.28500000 | -3.94769700 |
| C | -0.21986400 | -2.19944300 | -3.64186100 |
| C | -0.94268000 | -4.61120700 | -3.26327800 |
| C | 1.22989800 | -2.50625100 | -4.02048800 |
| H | -2.26366600 | -5.44523900 | -5.49972700 |
| H | -1.84264700 | -2.53149600 | -5.90450000 |
| H | -2.22040600 | -2.92010800 | -3.54552900 |
| H | -0.25871100 | -2.00566000 | -2.55943200 |
| H | -0.52849100 | -1.25476900 | -4.11871100 |

| | | | |
|---|---|---|---|
| H | -0.05856600 | -5.10235200 | -3.70528800 |
| H | -1.79199900 | -5.30456600 | -3.32230800 |
| H | -0.73981100 | -4.44506000 | -2.19742500 |
| H | 1.88176600 | -1.66963100 | -3.72665900 |
| H | 1.35743700 | -2.66692600 | -5.10286100 |
| H | 1.60828700 | -3.40443100 | -3.50569800 |
| C | 4.86952500 | -6.12502500 | -2.43535600 |
| C | 4.61653400 | -4.97265100 | -1.47465800 |
| O | 5.48713300 | -4.17181100 | -1.16604500 |
| H | 4.17927400 | -6.08849900 | -3.29304900 |
| N | 3.35457800 | -4.87835000 | -0.93928600 |
| C | 3.03993700 | -3.75605800 | -0.05186600 |
| C | 3.61910300 | -3.92586500 | 1.35730000 |
| O | 3.91070600 | -2.95524900 | 2.04220600 |
| C | 1.49645200 | -3.74423400 | -0.02509000 |
| C | 1.12442500 | -5.21095600 | -0.26704300 |
| C | 2.17248600 | -5.67791800 | -1.28483600 |
| H | 3.46248800 | -2.82183900 | -0.44293200 |
| H | 1.13042000 | -3.11713200 | -0.85229800 |
| H | 1.09238600 | -3.34460000 | 0.91524200 |
| H | 0.10012300 | -5.33878800 | -0.63206500 |
| H | 1.21170400 | -5.77822400 | 0.67233700 |
| H | 1.83775300 | -5.46607700 | -2.31515800 |
| H | 2.37923700 | -6.75781000 | -1.22076500 |
| N | 3.70415000 | -5.20653600 | 1.80229200 |
| C | 4.11918600 | -5.55080500 | 3.14478200 |
| C | 2.96580600 | -5.96168700 | 4.06822200 |
| C | 1.94266900 | -4.87907100 | 4.15936500 |
| C | 2.04891300 | -3.53342100 | 4.46079300 |
| N | 0.63561000 | -5.04811500 | 3.75727800 |
| C | 0.02093900 | -3.83790300 | 3.82954300 |
| N | 0.85061100 | -2.89751900 | 4.24529400 |
| H | 3.54571000 | -5.94358100 | 1.12773800 |
| H | 4.61955300 | -4.66310900 | 3.55599800 |
| H | 2.49503700 | -6.88701900 | 3.69471100 |
| H | 0.15390200 | -5.89976800 | 3.46631600 |
| H | 2.93754400 | -2.99077700 | 4.77602600 |
| H | -1.01325100 | -3.69132000 | 3.54253100 |
| H | 3.38891700 | -6.20682800 | 5.05776700 |
| C | 7.68735000 | 4.24386900 | 3.87178500 |
| C | 6.47066500 | 4.91508200 | 3.23077800 |
| O | 5.39440300 | 4.97875500 | 3.80437200 |
| H | 8.64052900 | 4.62190500 | 3.47458700 |
| N | 6.66734000 | 5.42112400 | 1.97044200 |
| C | 5.52136900 | 5.70107800 | 1.14239700 |
| C | 4.91694000 | 4.38807300 | 0.59120000 |
| O | 5.37124800 | 3.30003100 | 0.91099500 |
| H | 7.51999200 | 5.14420500 | 1.46360800 |
| H | 4.74785800 | 6.20890600 | 1.73814300 |
| N | 3.82929700 | 4.50387400 | -0.23829300 |
| C | 3.33832200 | 3.32165800 | -0.91842400 |
| C | 3.85949100 | 3.31417800 | -2.37368700 |
| O | 4.29297400 | 4.32765600 | -2.89377300 |
| C | 1.82296100 | 3.07998900 | -0.77343400 |
| C | 1.36344800 | 3.25464700 | 0.67021400 |
| O | 1.07693700 | 3.90095400 | -1.67151000 |
| H | 3.67848000 | 5.38663100 | -0.71640200 |
| H | 3.82427400 | 2.48326200 | -0.39738700 |
| H | 1.66758300 | 2.02823700 | -1.05592600 |
| H | 0.17327100 | 4.00720300 | -1.31507300 |
| H | 1.41678000 | 4.31370100 | 0.96328300 |
| H | 2.01540000 | 2.69319300 | 1.35441500 |
| H | 0.32717700 | 2.90329800 | 0.78577600 |
| N | 3.85472000 | 2.09817500 | -2.99490800 |
| C | 4.52668200 | 1.87911500 | -4.27167500 |
| H | 3.50859000 | 1.29433900 | -2.48213700 |
| H | 5.57819900 | 1.57899500 | -4.12112000 |
| C | -4.15536700 | -3.80307000 | 3.77592100 |
| O | -3.38409000 | -4.99102900 | 3.97051800 |
| C | -3.29974000 | -2.58891100 | 4.01724100 |
| O | -2.40239000 | -2.40335000 | 2.93951200 |
| C | -4.01116700 | -1.25193000 | 4.23232200 |
| O | -4.13323200 | -1.11103000 | 5.62051200 |
| C | -3.07335500 | -0.23557400 | 3.47593800 |
| O | -2.73451900 | 0.93103500 | 4.17261100 |
| C | -1.83053600 | -1.10261500 | 3.14756300 |
| N | 1.17791600 | -0.32805500 | 0.98359300 |
| C | 0.67342100 | -0.32104600 | -0.31163900 |
| O | 1.38213000 | -0.15282300 | -1.30327700 |
| N | -0.67986100 | -0.53584800 | -0.40482900 |
| C | -1.54227700 | -0.84927600 | 0.62811600 |
| O | -2.69763000 | -1.20648300 | 0.36331200 |
| C | -0.98318800 | -0.77202100 | 1.95840200 |
| C | 0.36131600 | -0.55880200 | 2.06084600 |
| H | -0.88877200 | 2.70784000 | 2.72725500 |
| H | -1.15956400 | -0.43040300 | -1.32772700 |
| H | 0.86278700 | -0.62417100 | 3.02875800 |
| H | -1.18990000 | -1.12259000 | 4.04670000 |
| H | -3.45211900 | 1.13744900 | 4.86116100 |
| H | -5.00792300 | -1.24633700 | 3.76273700 |
| H | -2.72099800 | -2.74767100 | 4.94983700 |
| H | -5.02061900 | -3.78668600 | 4.45945400 |
| H | -3.56708400 | 0.00825500 | 2.52337600 |
| H | -4.52248800 | -3.82503200 | 2.74036300 |
| P | -5.24878300 | -0.05760600 | 6.28854000 |
| O | -4.78389000 | 1.31810400 | 5.72205800 |
| O | -5.38094800 | -0.35656600 | 7.72739400 |
| O | -6.59705000 | -0.42646700 | 5.41624800 |
| C | 7.43255300 | -1.16395700 | 3.77900100 |
| C | 7.96985900 | -1.64106200 | 2.42687400 |
| C | 6.94371600 | -1.67638600 | 1.29306800 |
| C | 2.62357700 | -0.13857500 | 1.19676800 |
| S | 6.36519100 | 0.00050500 | 0.75115600 |
| C | 6.33147600 | -0.27879400 | -1.06237000 |
| C | 7.61876800 | -0.00295700 | -1.84927600 |
| O | 7.65776000 | 1.36734500 | -2.27108000 |
| C | 8.95219800 | -0.20423800 | -1.13022600 |
| O | 9.33224200 | -1.56786300 | -1.14943900 |
| C | 9.89907200 | 0.67116900 | -1.96901400 |
| O | 10.28092600 | -0.02072100 | -3.12978400 |
| C | 8.98273200 | 1.84363500 | -2.36215000 |
| C | 10.12063900 | 3.94725800 | -1.55620500 |
| C | 9.88636200 | 4.81567600 | -0.51485400 |
| N | 8.77390900 | 4.41566200 | 0.18501000 |
| C | 8.33183400 | 3.32114400 | -0.40805600 |
| N | 9.11983300 | 2.99662600 | -1.47764400 |
| H | 7.07548100 | -0.12266000 | 3.71975000 |
| H | 8.36152100 | -2.66909900 | 2.53955200 |
| H | 8.82989700 | -1.01414700 | 2.12708000 |
| H | 7.37562500 | -2.16452900 | 0.40956500 |
| H | 6.05390800 | -2.24804100 | 1.59214400 |
| H | 3.08911900 | -1.07194100 | 1.54243100 |
| H | 2.78783700 | 0.65466500 | 1.93710000 |
| H | 3.07868700 | 0.15938500 | 0.25005700 |
| H | 7.44642100 | 2.75820800 | -0.11906100 |
| H | 9.21419000 | 2.16744800 | -3.38952400 |
| H | 10.30890300 | -0.95983700 | -2.87968300 |
| H | 8.86210800 | 0.18363200 | -0.10337200 |
| H | 7.61625100 | -0.64983400 | -2.74569900 |
| H | 6.01068000 | -1.31698100 | -1.23266300 |
| H | 9.83079200 | -1.76547300 | -0.34679100 |
| H | 10.77492800 | 1.03045300 | -1.39685200 |
| H | 5.55202300 | 0.38323300 | -1.46692200 |
| C | -5.35014200 | 9.88028700 | 0.35521200 |
| C | -3.95395000 | 9.37084800 | 0.78170700 |
| O | -3.30954000 | 9.96009800 | 1.62927400 |
| H | -5.93982100 | 10.05050500 | 1.26638800 |
| N | -5.45071800 | 6.96903000 | -4.26502400 |
| C | -4.12939600 | 7.53560200 | -4.57650800 |
| H | -6.06328000 | 6.77735500 | -5.04985800 |

| | | | |
|---|---|---|---|
| H | -3.60753600 | 7.72316000 | -3.63021300 |
| O | -4.78186400 | 5.41648700 | 0.33561600 |
| H | -4.41654200 | 6.16886200 | 0.82617400 |
| H | -5.10424700 | 5.81759800 | -0.49693100 |
| H | -5.21984100 | 10.85070300 | -0.14691700 |
| H | -5.88026400 | 9.18637000 | -0.31373900 |
| H | -1.39888400 | 8.70086300 | 0.06063300 |
| H | -3.53563400 | 6.83224200 | -5.18096500 |
| H | -4.23629600 | 8.48472100 | -5.12314300 |
| H | -8.36049700 | -1.32486200 | -0.88409300 |
| H | -7.44036200 | -1.34431600 | -2.41107100 |
| H | -4.12438800 | -6.38675800 | -6.09741800 |
| H | -2.27505800 | -8.90466500 | -2.39195000 |
| H | -0.53517800 | -3.73077700 | -5.95240300 |
| H | 4.72074600 | -7.09527900 | -1.93478800 |
| H | 5.90346800 | -6.05396300 | -2.79177800 |
| H | 3.99916100 | 1.10482500 | -4.84752300 |
| H | 4.51463500 | 2.82233500 | -4.83267900 |
| H | 10.88130800 | 3.92026700 | -2.33227200 |
| H | 10.45094600 | 5.70534100 | -0.24059800 |
| H | 7.63785000 | 4.38083700 | 4.95972100 |
| H | 7.62780900 | 3.16554000 | 3.65079600 |
| H | 5.80525700 | 6.35350200 | 0.30090100 |
| H | -6.75046300 | 0.26880900 | 4.74924100 |
| H | 4.87049600 | -6.35751400 | 3.09761100 |
| H | 8.20968500 | -1.20868600 | 4.55754400 |
| H | 6.58273500 | -1.78700400 | 4.10249100 |
| H | -3.09275000 | -4.99667200 | 4.89419800 |
| H | -7.99185100 | 6.48573500 | -3.64108600 |

#### Model5' – Initial state

| | | | |
|---|---|---|---|
| O | -3.78842200 | -1.59971500 | -3.81989600 |
| H | -3.87303800 | -1.71674500 | -4.78384400 |
| H | -4.16111500 | -2.45232800 | -3.50577300 |
| O | -1.87302800 | 1.75372500 | 2.31937200 |
| H | -2.57824900 | 2.39305900 | 2.14925700 |
| H | -2.10945200 | 1.39724500 | 3.20641000 |
| O | -1.93709700 | 0.16004800 | -2.46786600 |
| H | -2.57415300 | -0.43179800 | -2.93216300 |
| H | -1.96623600 | 1.03667400 | -2.91890500 |
| O | -0.96897500 | -6.87890300 | 2.52946000 |
| H | -1.20330700 | -7.81170500 | 2.43918800 |
| H | -1.74137500 | -6.45050800 | 2.98157900 |
| O | -7.32336500 | 3.36405800 | 1.72468400 |
| H | -7.07354500 | 4.28389000 | 1.44498100 |
| H | -7.51516500 | 3.43798900 | 2.66912800 |
| O | -2.28205800 | 2.82156100 | -3.24869600 |
| H | -2.06977300 | 3.47879800 | -2.54588100 |
| H | -3.24698700 | 2.86953600 | -3.30218000 |
| C | -7.87817400 | 5.39251600 | -2.36823700 |
| C | -6.69001000 | 6.38034900 | -2.45431200 |
| O | -6.38861600 | 7.08465100 | -1.48901200 |
| C | -7.69180000 | 4.10255100 | -3.17236000 |
| C | -6.59407300 | 3.17328100 | -2.60583900 |
| C | -7.17413200 | 2.25433100 | -1.52784100 |
| C | -5.91789500 | 2.35428800 | -3.71054100 |
| H | -8.00513700 | 5.16937400 | -1.30003800 |
| H | -7.48403500 | 4.34311800 | -4.23101800 |
| H | -8.64815600 | 3.55432800 | -3.20147900 |
| H | -5.81704900 | 3.80127600 | -2.13090600 |
| H | -6.41289900 | 1.54504400 | -1.17862100 |
| H | -8.00909100 | 1.66270100 | -1.93960800 |
| H | -7.54181500 | 2.80487300 | -0.65016300 |
| H | -6.65106000 | 1.73874100 | -4.25877900 |
| H | -5.41702000 | 3.00515700 | -4.44821700 |
| H | -5.16975800 | 1.65903400 | -3.29455500 |
| N | -4.05585700 | 8.04777800 | 0.64899700 |
| C | -2.71484200 | 7.53361000 | 0.98379200 |
| C | -2.06073900 | 6.77829400 | -0.17374300 |
| C | -2.64514200 | 5.39943700 | -0.49989500 |
| O | -3.84533000 | 5.16574300 | -0.21004700 |
| O | -1.87503800 | 4.55806400 | -1.06661300 |
| H | -4.60135200 | 7.53573600 | -0.03767600 |
| H | -2.76955900 | 6.87887700 | 1.87287100 |
| H | -0.98805800 | 6.64007300 | 0.03139200 |
| H | -2.11665800 | 7.38281900 | -1.09758700 |
| C | -7.35894300 | -1.68757200 | -1.16390200 |
| C | -6.04850400 | -2.01869500 | -0.44728900 |
| C | -5.93373700 | -1.50529900 | 0.99844300 |
| C | -5.78829300 | 0.01316100 | 1.16899900 |
| N | -4.63326200 | 0.45617200 | 0.40332100 |
| C | -4.19234700 | 1.71485400 | 0.25506900 |
| N | -4.76696600 | 2.75421600 | 0.85454700 |
| N | -3.13181600 | 1.91568800 | -0.53021800 |
| H | -7.50965400 | -0.60167500 | -1.25787800 |
| H | -5.20838700 | -1.63171600 | -1.04777900 |
| H | -5.91339200 | -3.11291300 | -0.43525000 |
| H | -5.05232400 | -1.97555900 | 1.46669400 |
| H | -6.80738600 | -1.83237800 | 1.58686300 |
| H | -5.66905400 | 0.25217000 | 2.24282200 |
| H | -6.68816000 | 0.54578600 | 0.82285300 |
| H | -3.92485200 | -0.25373500 | 0.20897500 |
| H | -5.68554400 | 2.70724300 | 1.32123500 |
| H | -4.48578600 | 3.67704100 | 0.48836600 |
| H | -2.74679300 | 1.14614300 | -1.08224800 |
| H | -2.73080300 | 2.85133600 | -0.64846300 |
| C | -0.84963800 | -8.34192300 | -2.36626300 |
| C | -1.12640000 | -8.15828200 | -0.86969000 |
| C | -2.36917400 | -7.31689900 | -0.54535400 |
| N | -2.39222600 | -5.98779600 | -1.13551000 |
| C | -2.35804900 | -4.82464500 | -0.46614900 |
| N | -1.76383200 | -4.72455800 | 0.73922700 |
| N | -2.88794500 | -3.74698300 | -1.03650500 |
| H | -0.58195200 | -7.38841500 | -2.83899700 |
| H | -0.02311700 | -9.05099600 | -2.53052500 |
| H | -0.24740600 | -7.71521500 | -0.37190900 |
| H | -1.27599600 | -9.14658900 | -0.39856800 |
| H | -2.48353300 | -7.23923000 | 0.54536600 |
| H | -3.26037000 | -7.84565700 | -0.92485000 |
| H | -2.83091300 | -5.97087400 | -2.09856200 |
| H | -1.35478000 | -5.54828700 | 1.17553500 |
| H | -1.96259200 | -3.92878900 | 1.35392800 |
| H | -3.52438200 | -3.88919000 | -1.85704000 |
| H | -2.77215000 | -2.82632000 | -0.60759600 |
| C | -4.40623800 | -5.53838700 | -6.31607900 |
| C | -3.57585200 | -4.24066300 | -6.26818500 |
| O | -4.10144700 | -3.13645300 | -6.36743200 |
| C | -5.28032800 | -5.52458600 | -5.04110200 |
| C | -4.41110200 | -5.31939200 | -3.79222600 |
| O | -4.52924900 | -4.23176100 | -3.15621800 |
| O | -3.59589000 | -6.23671800 | -3.51003400 |
| H | -5.03770900 | -5.52873700 | -7.21611500 |
| H | -5.81527800 | -6.48288700 | -4.94929900 |
| H | -6.00889100 | -4.70642000 | -5.10969000 |
| N | -2.25230800 | -4.40832200 | -5.98900100 |
| C | -1.36689200 | -3.34851000 | -5.51118300 |
| C | -1.02495900 | -3.51841800 | -4.02601500 |
| C | -0.14847600 | -2.38111100 | -3.47866700 |
| C | -0.43541200 | -4.89945200 | -3.74588800 |
| C | 1.32509300 | -2.37101000 | -3.88755300 |
| H | -1.94938200 | -5.36063700 | -5.82908300 |
| H | -1.88577600 | -2.39670900 | -5.67667600 |
| H | -1.98307400 | -3.44853000 | -3.49591700 |
| H | -0.20959700 | -2.42853700 | -2.37773900 |
| H | -0.60793000 | -1.41720900 | -3.73827000 |
| H | 0.46941300 | -5.09471800 | -4.34744100 |
| H | -1.17145300 | -5.68933900 | -3.94535400 |
| H | -0.16822000 | -4.99115100 | -2.68584600 |
| H | 1.83764900 | -1.52008800 | -3.41221000 |
| H | 1.45532200 | -2.28091900 | -4.97834900 |

| | | | |
|---|---|---|---|
| H | 1.85302000 | -3.28564900 | -3.56915900 |
| C | 5.25071000 | -5.63756000 | -2.76792600 |
| C | 4.85846400 | -4.59815000 | -1.71832900 |
| O | 5.61980000 | -3.70256600 | -1.35906100 |
| H | 4.54821100 | -5.62517100 | -3.61576500 |
| N | 3.60938400 | -4.71791900 | -1.16930400 |
| C | 3.11900300 | -3.69511700 | -0.23774600 |
| C | 3.71934200 | -3.80599700 | 1.16950700 |
| O | 3.79827000 | -2.81305800 | 1.88887300 |
| C | 1.59422200 | -3.93205900 | -0.20635800 |
| C | 1.45334700 | -5.42152400 | -0.53001000 |
| C | 2.55603800 | -5.66803700 | -1.56530700 |
| H | 3.36594900 | -2.68898500 | -0.60129300 |
| H | 1.12063300 | -3.32367200 | -0.98957300 |
| H | 1.14123700 | -3.65577600 | 0.75444400 |
| H | 0.45873000 | -5.68055000 | -0.90868800 |
| H | 1.62762600 | -6.02357600 | 0.37567300 |
| H | 2.19336800 | -5.44823000 | -2.58296800 |
| H | 2.92662400 | -6.70444200 | -1.55939500 |
| N | 4.08120900 | -5.05811200 | 1.54297600 |
| C | 4.58233600 | -5.45154300 | 2.84896600 |
| C | 3.51556900 | -6.08185000 | 3.75842800 |
| C | 2.41176700 | -5.11838500 | 4.04787300 |
| C | 2.43223900 | -3.86328400 | 4.62715700 |
| N | 1.11958100 | -5.27309100 | 3.58968700 |
| C | 0.43058400 | -4.14214400 | 3.90937200 |
| N | 1.20141300 | -3.26725100 | 4.52852500 |
| H | 4.02259200 | -5.76711300 | 0.82240100 |
| H | 4.97745900 | -4.54520100 | 3.33010100 |
| H | 4.01266500 | -6.40974100 | 4.68739500 |
| H | 3.11207100 | -6.99233900 | 3.28441000 |
| H | 0.67554300 | -6.09237700 | 3.16848500 |
| H | 3.28074000 | -3.35312700 | 5.08083300 |
| H | -0.60973000 | -3.99667800 | 3.63105200 |
| C | 7.34930900 | 4.54815800 | 4.09360100 |
| C | 6.05106600 | 5.08611100 | 3.49164000 |
| O | 4.97403500 | 4.90708500 | 4.03435600 |
| H | 8.24565000 | 5.06612800 | 3.72137500 |
| N | 6.17955800 | 5.76250700 | 2.30685400 |
| C | 5.01031800 | 5.97529700 | 1.49482600 |
| C | 4.52323600 | 4.64236200 | 0.86739700 |
| O | 5.17643400 | 3.60589800 | 1.00549700 |
| H | 7.04277600 | 5.66451800 | 1.76388700 |
| H | 4.19413300 | 6.39786100 | 2.09962000 |
| N | 3.36256300 | 4.67264600 | 0.17469600 |
| C | 2.93568400 | 3.56740800 | -0.65781700 |
| C | 3.70446900 | 3.61612800 | -2.01089400 |
| O | 4.44869700 | 4.54031500 | -2.28697600 |
| C | 1.39434600 | 3.50661800 | -0.74685100 |
| C | 0.78092700 | 3.12002200 | 0.59889000 |
| O | 0.93527500 | 4.75716000 | -1.20217300 |
| H | 2.89418100 | 5.55488700 | -0.00916800 |
| H | 3.23920500 | 2.63802000 | -0.16163900 |
| H | 1.15050200 | 2.70822400 | -1.47936000 |
| H | -0.05052300 | 4.75084700 | -1.18430500 |
| H | 1.00358200 | 3.88715500 | 1.35533400 |
| H | 1.18022000 | 2.16151700 | 0.96361300 |
| H | -0.30931200 | 3.01343100 | 0.52824300 |
| N | 3.55791100 | 2.51573600 | -2.81605500 |
| C | 4.21062700 | 2.40259900 | -4.11906600 |
| H | 2.90242000 | 1.79318200 | -2.53209000 |
| H | 4.95900500 | 1.59093300 | -4.12700400 |
| C | -3.81661700 | -4.45891400 | 3.70537200 |
| O | -2.92121200 | -5.56744500 | 3.83937300 |
| C | -3.07482600 | -3.19304800 | 4.01301300 |
| O | -2.19722800 | -2.87062100 | 2.94383500 |
| C | -3.88166000 | -1.93723800 | 4.33402200 |
| O | -3.95630400 | -1.90067300 | 5.72941200 |
| C | -3.04119100 | -0.79901000 | 3.63682200 |
| O | -2.78613700 | 0.30212500 | 4.44425200 |
| C | -1.73607800 | -1.53917800 | 3.24748800 |
| N | 1.39930600 | -0.77704700 | 1.33726900 |
| C | 1.02848600 | -0.48192200 | 0.03665500 |
| O | 1.85473900 | -0.22298100 | -0.83348600 |
| N | -0.33223300 | -0.55230600 | -0.17225700 |
| C | -1.29043400 | -0.96166300 | 0.75856900 |
| O | -2.42590400 | -1.24438600 | 0.36118800 |
| C | -0.83388700 | -1.10237000 | 2.12976100 |
| C | 0.50913500 | -1.05589800 | 2.34038900 |
| H | 2.38853000 | -0.95794900 | 1.49823900 |
| H | -0.70457100 | -0.34259300 | -1.11844000 |
| H | 0.94329500 | -1.31036600 | 3.31084600 |
| H | -1.12745100 | -1.57672200 | 4.16912900 |
| H | -3.63153100 | 0.47957200 | 4.99182500 |
| H | -4.89264100 | -1.97269300 | 3.89223500 |
| H | -2.47234200 | -3.36165500 | 4.92817700 |
| H | -4.67199500 | -4.56588500 | 4.39244000 |
| H | -3.55815100 | -0.51368100 | 2.70514800 |
| H | -4.18602000 | -4.46636000 | 2.67004200 |
| P | -5.13227600 | -0.94824500 | 6.47962500 |
| O | -5.03597400 | 0.38590200 | 5.72800600 |
| O | -4.99997800 | -1.15210500 | 7.94023600 |
| O | -6.49678100 | -1.68198100 | 5.86884000 |
| C | 7.53754600 | -0.84762000 | 3.68405900 |
| C | 7.22632100 | -1.71006900 | 2.45674400 |
| C | 6.17377300 | -1.15268100 | 1.50265400 |
| C | 5.20734000 | 0.88542000 | -0.22503500 |
| S | 6.74700800 | 0.29801600 | 0.52639500 |
| C | 7.56269900 | -0.55278800 | -0.87255400 |
| C | 8.42650700 | 0.40195100 | -1.67457600 |
| O | 7.74370400 | 1.61334500 | -1.96599100 |
| C | 9.71726900 | 0.87452400 | -1.00381900 |
| O | 10.66841200 | -0.16365800 | -1.00041700 |
| C | 10.08452500 | 2.04658000 | -1.93288100 |
| O | 10.67338000 | 1.54676400 | -3.10423100 |
| C | 8.69639000 | 2.62840100 | -2.29413100 |
| C | 8.73930400 | 5.10640500 | -2.04769700 |
| C | 8.49730800 | 5.95931600 | -0.99839600 |
| N | 7.99991700 | 5.25481500 | 0.07184600 |
| C | 7.94019300 | 3.99576700 | -0.30434100 |
| N | 8.37815500 | 3.84778600 | -1.59441500 |
| H | 7.93722700 | 0.13996400 | 3.40017900 |
| H | 6.82206600 | -2.68319800 | 2.78141400 |
| H | 8.15059200 | -1.94809600 | 1.90062900 |
| H | 5.84219300 | -1.89323700 | 0.76232800 |
| H | 5.28662500 | -0.79078800 | 2.03860500 |
| H | 4.47620500 | 0.06935600 | -0.30672200 |
| H | 4.88214200 | 1.72758000 | 0.40742400 |
| H | 5.47861600 | 1.28915600 | -1.20736700 |
| H | 7.51827700 | 3.18572300 | 0.28348000 |
| H | 8.67430000 | 2.83644000 | -3.37474900 |
| H | 11.22285200 | 0.78948300 | -2.84377000 |
| H | 9.49772100 | 1.24175200 | 0.01435800 |
| H | 8.71429800 | -0.12389800 | -2.60399100 |
| H | 8.20684100 | -1.33419500 | -0.44340400 |
| H | 11.33511100 | 0.01237800 | -0.32345800 |
| H | 10.71183000 | 2.81322300 | -1.44352700 |
| H | 6.77559800 | -1.04471400 | -1.46237700 |
| C | -6.18292600 | 9.27255300 | 1.15017600 |
| C | -4.69383600 | 8.97224300 | 1.44437500 |
| O | -4.11494300 | 9.52213900 | 2.36888900 |
| H | -6.40397000 | 10.28015100 | 1.52204600 |
| N | -6.02812300 | 6.45824100 | -3.64077000 |
| C | -4.88538400 | 7.32692700 | -3.93333200 |
| H | -6.33441900 | 5.83549200 | -4.37752200 |
| H | -4.67914100 | 7.92790300 | -3.03820300 |
| O | -6.34399600 | 5.74271400 | 0.91845600 |
| H | -5.39032800 | 5.64245900 | 0.72101800 |
| H | -6.63625800 | 6.29494500 | 0.16543600 |
| H | -6.43084000 | 9.17604000 | 0.08334200 |

| | | | |
|---|---|---|---|
| H | -6.78964100 | 8.53776600 | 1.70300400 |
| H | -2.09623700 | 8.39538700 | 1.27253100 |
| H | -3.99240500 | 6.72908400 | -4.17598200 |
| H | -5.11100500 | 8.00012100 | -4.77524100 |
| H | -8.22718500 | -2.10008600 | -0.62310000 |
| H | -7.35657300 | -2.11248300 | -2.17943600 |
| H | -3.76349600 | -6.43064100 | -6.33026900 |
| H | -1.74089400 | -8.72032900 | -2.89126800 |
| H | -0.44914700 | -3.34006500 | -6.12665500 |
| H | 5.23479600 | -6.65254500 | -2.34078300 |
| H | 6.26159800 | -5.40605500 | -3.12245600 |
| H | 3.47435900 | 2.21483100 | -4.91617300 |
| H | 4.72153400 | 3.35442200 | -4.31267000 |
| H | 9.11804100 | 5.26756700 | -3.05348000 |
| H | 8.63710500 | 7.03808000 | -0.96220200 |
| H | 7.29224000 | 4.62689400 | 5.18727300 |
| H | 7.42657100 | 3.47940200 | 3.83352600 |
| H | 5.24534500 | 6.67258700 | 0.67644500 |
| H | -6.94819700 | -1.00737200 | 5.34366700 |
| H | 5.42458900 | -6.14889700 | 2.70601900 |
| H | 8.28373200 | -1.33932700 | 4.32511000 |
| H | 6.63126800 | -0.67764800 | 4.28594700 |
| H | -2.63390800 | -5.59052900 | 4.76388300 |
| H | -8.77763700 | 5.94104600 | -2.69614600 |

#### Model5' – TS1

| | | | |
|---|---|---|---|
| O | -5.21005200 | -1.75823800 | -3.58953500 |
| H | -5.01624900 | -1.94426300 | -4.53327300 |
| H | -5.48198600 | -2.65292500 | -3.25934800 |
| O | -1.09434100 | 2.63718400 | 0.13146000 |
| H | -1.01945300 | 3.48623200 | -0.44049400 |
| H | -1.89112000 | 2.14293400 | -0.19789700 |
| O | -3.03225600 | -0.70580000 | -2.66541100 |
| H | -3.89340600 | -1.15823100 | -2.96858800 |
| H | -2.66794100 | -0.32578600 | -3.47709500 |
| O | -1.09351200 | -6.31950400 | 2.60173200 |
| H | -1.19232200 | -7.27020200 | 2.74556000 |
| H | -1.83908100 | -5.90578800 | 3.11864600 |
| O | -6.43614500 | 3.67175000 | 2.33872600 |
| H | -6.10109100 | 4.56445000 | 2.09190500 |
| H | -6.19841000 | 3.54076700 | 3.26824200 |
| O | -3.29718600 | 3.93563100 | -2.54095700 |
| H | -2.40355500 | 4.23123400 | -2.26539800 |
| H | -3.87067800 | 4.63604900 | -2.17139000 |
| C | -7.29072300 | 6.01656400 | -2.17203000 |
| C | -5.83276700 | 6.45828100 | -2.36750300 |
| O | -4.94364900 | 6.05461500 | -1.60279800 |
| C | -7.57688300 | 4.72030400 | -2.96023300 |
| C | -6.94946300 | 3.43998200 | -2.38072700 |
| C | -7.63218200 | 3.02616100 | -1.07406300 |
| C | -6.99989900 | 2.30792500 | -3.41232000 |
| H | -7.44877900 | 5.86893000 | -1.09515100 |
| H | -7.22937500 | 4.86288000 | -3.99828100 |
| H | -8.67044300 | 4.58300500 | -3.02082600 |
| H | -5.88802300 | 3.64132000 | -2.16559200 |
| H | -7.23941100 | 2.06341000 | -0.72162100 |
| H | -8.71812500 | 2.90074100 | -1.22463500 |
| H | -7.47792600 | 3.74765400 | -0.25839100 |
| H | -8.04183100 | 2.04768800 | -3.66667500 |
| H | -6.48720100 | 2.59487100 | -4.34457900 |
| H | -6.51531300 | 1.39306800 | -3.03678100 |
| N | -3.30705300 | 8.14588600 | 1.12237400 |
| C | -1.90253800 | 7.71314800 | 1.07852000 |
| C | -1.46495400 | 7.01772300 | -0.20980300 |
| C | -1.81740800 | 5.54553100 | -0.30894100 |
| O | -2.74658100 | 5.06155000 | 0.36124800 |
| O | -1.09545000 | 4.84262500 | -1.11752400 |
| H | -4.03000400 | 7.44291800 | 1.26556500 |
| H | -1.70230000 | 7.05821800 | 1.94181200 |
| H | -0.37276700 | 7.09686300 | -0.33045500 |
| H | -1.91214700 | 7.52365700 | -1.08242200 |
| C | -7.35610900 | -1.07981900 | -0.95414300 |
| C | -6.61599000 | -1.79259800 | 0.17515000 |
| C | -6.51394900 | -1.01527200 | 1.49624300 |
| C | -5.93573100 | 0.40593400 | 1.41048400 |
| N | -4.85590700 | 0.48067400 | 0.42579000 |
| C | -4.30981400 | 1.61142900 | -0.00873800 |
| N | -4.64975800 | 2.79730200 | 0.46310200 |
| N | -3.34487200 | 1.56413700 | -0.98256400 |
| H | -6.84426600 | -0.15276400 | -1.25042600 |
| H | -5.60607200 | -2.07153400 | -0.16748300 |
| H | -7.10975900 | -2.75563800 | 0.38821800 |
| H | -5.91258600 | -1.60770600 | 2.20277600 |
| H | -7.51155100 | -0.92372200 | 1.95646500 |
| H | -5.56582400 | 0.72495500 | 2.39909100 |
| H | -6.71491000 | 1.12149500 | 1.10964500 |
| H | -4.38770000 | -0.38938600 | 0.16439100 |
| H | -5.31032600 | 2.93550800 | 1.25182300 |
| H | -4.06780900 | 3.59479100 | 0.19470000 |
| H | -3.30169600 | 0.69257500 | -1.53863700 |
| H | -3.35964300 | 2.40921100 | -1.59481600 |
| C | -1.46236000 | -8.26839000 | -2.25341700 |
| C | -1.59703800 | -8.05408500 | -0.74342400 |
| C | -2.91406400 | -7.38467400 | -0.31965500 |
| N | -3.20631200 | -6.13584800 | -0.99315200 |
| C | -3.01491800 | -4.90158200 | -0.50564300 |
| N | -2.26653500 | -4.67398200 | 0.59272700 |
| N | -3.54788300 | -3.87352000 | -1.15720100 |
| H | -1.48771400 | -7.31108500 | -2.79154200 |
| H | -0.51669800 | -8.77673900 | -2.49991900 |
| H | -0.75312800 | -7.45104900 | -0.37237600 |
| H | -1.53273600 | -9.02356500 | -0.21790200 |
| H | -2.91308700 | -7.22411000 | 0.77032200 |
| H | -3.75010000 | -8.07064100 | -0.53615200 |
| H | -3.70798400 | -6.17806200 | -1.91428800 |
| H | -1.81620400 | -5.42565700 | 1.11340800 |
| H | -2.35429900 | -3.77236000 | 1.06292100 |
| H | -4.35341600 | -4.05281200 | -1.80185500 |
| H | -3.40012200 | -2.94325700 | -0.75134000 |
| C | -4.83868000 | -5.17774300 | -6.14680300 |
| C | -3.92496500 | -3.96563300 | -5.97534700 |
| O | -4.36420500 | -2.81504900 | -5.99044400 |
| C | -5.90991100 | -5.18483700 | -5.05109000 |
| C | -5.26180600 | -5.23052600 | -3.66465000 |
| O | -5.58161500 | -4.31799900 | -2.84672200 |
| O | -4.42229000 | -6.13798700 | -3.44647800 |
| H | -5.31265800 | -5.10000300 | -7.13841600 |
| H | -6.54784000 | -6.07486900 | -5.17448700 |
| H | -6.54312500 | -4.29121000 | -5.12968500 |
| N | -2.61267700 | -4.24261400 | -5.77878700 |
| C | -1.60914600 | -3.25437300 | -5.40455200 |
| C | -1.29703400 | -3.25962400 | -3.90220100 |
| C | -0.15860000 | -2.28686300 | -3.55587000 |
| C | -1.04426800 | -4.67085600 | -3.37579200 |
| C | 1.24002000 | -2.64638500 | -4.05972900 |
| H | -2.36569300 | -5.21979900 | -5.68580900 |
| H | -1.99644300 | -2.27345100 | -5.71276000 |
| H | -2.19854200 | -2.88538600 | -3.39509600 |
| H | -0.11263300 | -2.19366900 | -2.46181800 |
| H | -0.41544500 | -1.27603200 | -3.91188400 |
| H | -0.21781400 | -5.17540700 | -3.90646800 |
| H | -1.94661900 | -5.29081100 | -3.46159900 |
| H | -0.78868500 | -4.63033500 | -2.30910000 |
| H | 1.95855600 | -1.87243500 | -3.74740900 |
| H | 1.29077900 | -2.72168400 | -5.15785900 |
| H | 1.59392100 | -3.60508500 | -3.64698400 |
| C | 4.83836400 | -6.09592400 | -2.77522700 |
| C | 4.57142700 | -5.00745000 | -1.73971200 |
| O | 5.42950800 | -4.19302400 | -1.40592700 |
| H | 4.11851700 | -6.03949800 | -3.60671800 |

| | | | |
|---|---|---|---|
| N | 3.32641900 | -4.98652600 | -1.17393500 |
| C | 2.98821400 | -3.93846300 | -0.20552300 |
| C | 3.63008900 | -4.15701300 | 1.17030300 |
| O | 3.93587500 | -3.20498900 | 1.87129400 |
| C | 1.44973000 | -3.99233800 | -0.12920200 |
| C | 1.12957200 | -5.45233800 | -0.46153200 |
| C | 2.16575500 | -5.81567600 | -1.53250700 |
| H | 3.35160000 | -2.96226900 | -0.55125000 |
| H | 1.02249600 | -3.32253500 | -0.89039300 |
| H | 1.06878700 | -3.67264200 | 0.84981700 |
| H | 0.10192800 | -5.59057000 | -0.81130900 |
| H | 1.25695700 | -6.08058100 | 0.43354600 |
| H | 1.79493600 | -5.55699100 | -2.53905500 |
| H | 2.41984900 | -6.88696900 | -1.53811900 |
| N | 3.78411800 | -5.45155300 | 1.55183400 |
| C | 4.29130000 | -5.84342700 | 2.85252500 |
| C | 3.19097700 | -6.13801000 | 3.89015500 |
| C | 2.35839300 | -4.93691000 | 4.19766200 |
| C | 2.64905800 | -3.78467700 | 4.90375900 |
| N | 1.11899900 | -4.69959400 | 3.63517900 |
| C | 0.72031100 | -3.45218300 | 4.03561500 |
| N | 1.62966500 | -2.87517200 | 4.79445200 |
| H | 3.56085900 | -6.17104000 | 0.87738300 |
| H | 4.91470300 | -5.01700900 | 3.22158300 |
| H | 3.68603000 | -6.50198100 | 4.80607500 |
| H | 2.55632100 | -6.96595200 | 3.52893500 |
| H | 0.51471600 | -5.37023400 | 3.15425500 |
| H | 3.55457300 | -3.56853900 | 5.47146900 |
| H | -0.24953900 | -3.04926600 | 3.74289700 |
| C | 7.92551900 | 3.88505500 | 4.01354200 |
| C | 6.67803100 | 4.58025100 | 3.46265800 |
| O | 5.61651800 | 4.55692600 | 4.06096900 |
| H | 8.86005600 | 4.37312700 | 3.69780300 |
| N | 6.83458600 | 5.20780700 | 2.25237900 |
| C | 5.66906200 | 5.50214400 | 1.45319200 |
| C | 5.11807100 | 4.24548900 | 0.79625900 |
| O | 5.54717200 | 3.11880800 | 1.05034600 |
| H | 7.66552600 | 5.01334100 | 1.68758600 |
| H | 4.88273900 | 5.91841000 | 2.10134600 |
| N | 4.09576600 | 4.48243600 | -0.03472000 |
| C | 3.40979500 | 3.44670600 | -0.75017800 |
| C | 4.12854100 | 3.26541900 | -2.10470800 |
| O | 5.10680700 | 3.93853300 | -2.39141500 |
| C | 1.90298500 | 3.74695000 | -0.71395400 |
| C | 1.48782700 | 3.91718000 | 0.75480000 |
| O | 1.67427400 | 4.91947000 | -1.46578700 |
| H | 3.85810900 | 5.43295900 | -0.29684300 |
| H | 3.54995000 | 2.51150300 | -0.19130000 |
| H | 1.36963000 | 2.88027300 | -1.15108400 |
| H | 0.70467200 | 5.06772800 | -1.47018800 |
| H | 2.06514600 | 4.72490600 | 1.22138900 |
| H | 1.66558600 | 2.99255200 | 1.32323700 |
| H | 0.42939700 | 4.17200100 | 0.84613900 |
| N | 3.72478100 | 2.24097600 | -2.90198200 |
| C | 4.46383600 | 2.00172100 | -4.13305000 |
| H | 2.85637600 | 1.74887400 | -2.71022000 |
| H | 5.52013700 | 1.78068900 | -3.90910300 |
| C | -3.90199600 | -4.09820900 | 3.82350100 |
| O | -3.17399200 | -5.31525500 | 3.98015700 |
| C | -3.07480400 | -2.89389700 | 4.19382800 |
| O | -2.15372200 | -2.56154500 | 3.15301300 |
| C | -3.92040000 | -1.65256800 | 4.49626900 |
| O | -3.68533900 | -1.30553300 | 5.83338300 |
| C | -3.43239400 | -0.63682100 | 3.43811600 |
| O | -3.54004500 | 0.70364800 | 3.76444600 |
| C | -1.99758300 | -1.12674500 | 3.15067700 |
| N | 0.22371500 | 0.65488300 | 0.62420600 |
| C | -0.08986900 | 0.03642700 | -0.56731000 |
| O | 0.45365200 | 0.31285300 | -1.63231400 |
| N | -1.15971500 | -0.86299300 | -0.52370600 |
| C | -1.90446300 | -1.16318300 | 0.59808100 |
| O | -2.99416300 | -1.76301300 | 0.45947300 |
| C | -1.36405300 | -0.71738500 | 1.85624900 |
| C | -0.35992200 | 0.21496900 | 1.77640700 |
| H | -0.24111000 | 1.85862200 | 0.37506900 |
| H | -1.62673700 | -1.06388400 | -1.41796900 |
| H | -0.02467700 | 0.71900100 | 2.68918500 |
| H | -1.35236200 | -0.81312900 | 3.99242500 |
| H | -4.19545800 | 0.78644000 | 4.53005400 |
| H | -4.99411000 | -1.85479900 | 4.34714100 |
| H | -2.51467100 | -3.10838300 | 5.12367300 |
| H | -4.80077900 | -4.12669200 | 4.46155800 |
| H | -3.99449000 | -0.88056200 | 2.52124000 |
| H | -4.21752200 | -4.04314800 | 2.77159900 |
| P | -4.93346500 | -0.52048400 | 6.64417300 |
| O | -5.34322000 | 0.60281400 | 5.68330600 |
| O | -4.52492400 | -0.35624700 | 8.05904200 |
| O | -6.12749200 | -1.67847600 | 6.47500300 |
| C | 7.64456100 | -1.50778600 | 3.61838200 |
| C | 7.17252400 | -2.30975900 | 2.40502900 |
| C | 6.18110500 | -1.60781300 | 1.48609900 |
| C | 5.40881700 | 0.45122100 | -0.28074200 |
| S | 6.89712900 | -0.23448900 | 0.49335500 |
| C | 7.63527000 | -1.19542600 | -0.87736400 |
| C | 8.56772000 | -0.34210600 | -1.71360500 |
| O | 7.96936300 | 0.90074100 | -2.05409800 |
| C | 9.88578800 | 0.06388300 | -1.05112400 |
| O | 10.76872700 | -1.03268500 | -1.02236700 |
| C | 10.33130800 | 1.18828500 | -2.00425100 |
| O | 10.89617100 | 0.62394600 | -3.15798600 |
| C | 8.98555100 | 1.84618200 | -2.39140500 |
| C | 9.11545200 | 4.33013900 | -2.22422800 |
| C | 8.92919900 | 5.22008300 | -1.19456800 |
| N | 8.44357900 | 4.56154800 | -0.09032400 |
| C | 8.33482700 | 3.29543500 | -0.42792100 |
| N | 8.73230400 | 3.09709500 | -1.72359700 |
| H | 8.18318000 | -0.59183600 | 3.32247300 |
| H | 6.63063100 | -3.20799400 | 2.74229800 |
| H | 8.03273800 | -2.68338300 | 1.82126300 |
| H | 5.74615700 | -2.29531000 | 0.75102100 |
| H | 5.35227100 | -1.14732200 | 2.04043800 |
| H | 4.65328200 | -0.33992600 | -0.37096000 |
| H | 5.11068100 | 1.29655200 | 0.36048200 |
| H | 5.71856700 | 0.84525000 | -1.25508900 |
| H | 7.91222600 | 2.51366900 | 0.19736000 |
| H | 8.98780600 | 2.03293800 | -3.47598400 |
| H | 11.38717200 | -0.16460200 | -2.87398600 |
| H | 9.69004100 | 0.46651300 | -0.04178000 |
| H | 8.82064400 | -0.92199400 | -2.62061600 |
| H | 8.21579300 | -2.00951100 | -0.41945200 |
| H | 11.43380400 | -0.89344100 | -0.33542200 |
| H | 11.00214200 | 1.92583500 | -1.52795300 |
| H | 6.81035600 | -1.64618900 | -1.44840600 |
| C | -5.20581800 | 9.74384400 | 1.30186500 |
| C | -3.69612200 | 9.46091700 | 1.09834100 |
| O | -2.91261200 | 10.38490200 | 0.93930200 |
| H | -5.31307100 | 10.36324200 | 2.20452200 |
| N | -5.53289400 | 7.23950600 | -3.43355000 |
| C | -4.17332000 | 7.68536600 | -3.79776000 |
| H | -6.30305700 | 7.55872100 | -4.00897100 |
| H | -3.80972100 | 8.45938100 | -3.10342800 |
| O | -5.16549200 | 5.82237900 | 1.17214100 |
| H | -4.22305000 | 5.53635300 | 1.14788900 |
| H | -5.32253100 | 5.89197500 | 0.20680000 |
| H | -5.56205000 | 10.33935300 | 0.44877300 |
| H | -5.81250100 | 8.83248600 | 1.40720300 |
| H | -1.31196300 | 8.63152200 | 1.20161100 |
| H | -3.48492500 | 6.82854500 | -3.76022700 |
| H | -4.20048200 | 8.08920200 | -4.81715700 |
| H | -8.38424900 | -0.81370500 | -0.65631300 |

| | | | |
|---|---|---|---|
| H | -7.40816900 | -1.71458600 | -1.85045000 |
| H | -4.26351200 | -6.11370200 | -6.10586000 |
| H | -2.28987600 | -8.88216300 | -2.64457600 |
| H | -0.69668800 | -3.45440100 | -5.99166400 |
| H | 4.74281300 | -7.09613500 | -2.32358100 |
| H | 5.85670700 | -5.96586100 | -3.15908700 |
| H | 4.01882100 | 1.14675200 | -4.66069200 |
| H | 4.44480900 | 2.88433600 | -4.79199500 |
| H | 9.46745000 | 4.45048300 | -3.24522000 |
| H | 9.10151900 | 6.29470200 | -1.19399500 |
| H | 7.86107800 | 3.85622800 | 5.10851000 |
| H | 7.93048200 | 2.84820800 | 3.63767700 |
| H | 5.91930200 | 6.24273500 | 0.67847900 |
| H | -6.84527200 | -1.24257600 | 5.99594500 |
| H | 4.93852600 | -6.72708100 | 2.72436400 |
| H | 8.32391000 | -2.10804500 | 4.24135000 |
| H | 6.79095000 | -1.20282300 | 4.24344600 |
| H | -2.95399800 | -5.39500600 | 4.91975500 |
| H | -7.97307100 | 6.81491000 | -2.50630100 |

#### Model5' – Intermediate

| | | | |
|---|---|---|---|
| O | -4.17980800 | -1.93179800 | -3.80172100 |
| H | -4.16739500 | -2.12659800 | -4.75993500 |
| H | -4.48104000 | -2.79352000 | -3.42446200 |
| O | -2.30352500 | 1.85046300 | 2.07695800 |
| H | -2.24048700 | 3.17182000 | 1.33673100 |
| H | -1.59445500 | 1.21212200 | 1.89449100 |
| O | -2.24332600 | -0.45057800 | -2.71131400 |
| H | -2.98312500 | -1.03105400 | -3.06098400 |
| H | -1.77913000 | -0.17154400 | -3.51215800 |
| O | -0.61640200 | -6.69106000 | 2.69828900 |
| H | -0.76925900 | -7.64471600 | 2.71285100 |
| H | -1.39503300 | -6.28897200 | 3.16752200 |
| O | -6.96609400 | 3.17655000 | 1.82336800 |
| H | -6.86741800 | 4.14065200 | 1.63246900 |
| H | -6.63663600 | 3.06979900 | 2.72699500 |
| O | -3.14822100 | 4.45537400 | -2.64953500 |
| H | -2.39916900 | 4.98515500 | -2.34656400 |
| H | -3.94634300 | 4.92766000 | -2.32825200 |
| C | -7.92009700 | 5.25664300 | -2.18480800 |
| C | -6.49455000 | 5.77223400 | -2.43557200 |
| O | -5.54706400 | 5.40017700 | -1.71050400 |
| C | -8.24527700 | 4.04055600 | -3.07589700 |
| C | -7.54432100 | 2.73580900 | -2.66437900 |
| C | -8.12374100 | 2.18704000 | -1.35854300 |
| C | -7.63011500 | 1.69106800 | -3.78107300 |
| H | -7.99129400 | 5.00187500 | -1.11949800 |
| H | -7.98732000 | 4.28942200 | -4.12013700 |
| H | -9.33709400 | 3.88027400 | -3.06060800 |
| H | -6.47587100 | 2.95740200 | -2.50084200 |
| H | -7.60412800 | 1.26621500 | -1.06328600 |
| H | -9.19182600 | 1.94061900 | -1.48491600 |
| H | -8.03060100 | 2.88174100 | -0.51242100 |
| H | -8.67911700 | 1.41788500 | -3.98785200 |
| H | -7.18926400 | 2.06534900 | -4.71938900 |
| H | -7.09309600 | 0.77035800 | -3.50375500 |
| N | -3.98893500 | 8.01472500 | 1.41287500 |
| C | -2.68174600 | 7.51625600 | 0.96642600 |
| C | -2.69527800 | 6.21169400 | 0.16145500 |
| C | -2.91024900 | 4.98125100 | 1.01359800 |
| O | -3.78114600 | 4.91851500 | 1.86976000 |
| O | -2.07334200 | 4.00306800 | 0.74580300 |
| H | -4.25376100 | 7.85003600 | 2.37603000 |
| H | -2.04518400 | 7.41515600 | 1.85689200 |
| H | -1.73876900 | 6.11907500 | -0.37444900 |
| H | -3.50183700 | 6.27773000 | -0.58396700 |
| C | -7.27015700 | -1.79553600 | -0.88438500 |
| C | -5.91756000 | -2.10702600 | -0.25262900 |
| C | -5.65594700 | -1.40707500 | 1.08554300 |
| C | -5.51456900 | 0.11856300 | 1.01467700 |
| N | -4.47522500 | 0.47538500 | 0.06094800 |
| C | -4.30863700 | 1.67528300 | -0.50112400 |
| N | -5.09271900 | 2.70481000 | -0.16828800 |
| N | -3.35289300 | 1.83374800 | -1.42373200 |
| H | -7.36670500 | -0.72937900 | -1.13890500 |
| H | -5.12124900 | -1.84426100 | -0.96690300 |
| H | -5.82715200 | -3.19519700 | -0.10189000 |
| H | -4.72953600 | -1.81666800 | 1.51471900 |
| H | -6.46502800 | -1.63569300 | 1.80031400 |
| H | -5.27182500 | 0.51168400 | 2.01941000 |
| H | -6.45977900 | 0.59325500 | 0.71242600 |
| H | -3.68562700 | -0.18304500 | 0.00187000 |
| H | -5.72401700 | 2.70632000 | 0.64886900 |
| H | -5.07263400 | 3.55389000 | -0.72279600 |
| H | -2.94068400 | 1.00500300 | -1.87111400 |
| H | -3.20801200 | 2.75499000 | -1.84783400 |
| C | -0.71537800 | -8.38630000 | -2.18713000 |
| C | -0.79374700 | -8.02082700 | -0.70131300 |
| C | -2.11911200 | -7.37317200 | -0.26954700 |
| N | -2.41277000 | -6.09581100 | -0.89058900 |
| C | -2.31552700 | -4.88722200 | -0.31437600 |
| N | -1.61425600 | -4.69972000 | 0.82219300 |
| N | -2.89582500 | -3.84823600 | -0.90401000 |
| H | -0.84207200 | -7.49596400 | -2.82069100 |
| H | 0.25354100 | -8.84887200 | -2.43253900 |
| H | 0.03595700 | -7.34743100 | -0.43020400 |
| H | -0.65835800 | -8.93240900 | -0.09108100 |
| H | -2.12728300 | -7.25889100 | 0.82359700 |
| H | -2.94856400 | -8.05501200 | -0.52397100 |
| H | -2.90862800 | -6.14453300 | -1.81796100 |
| H | -1.15689000 | -5.48180300 | 1.28502000 |
| H | -1.76625800 | -3.85779600 | 1.38577500 |
| H | -3.58568100 | -4.03279000 | -1.66404200 |
| H | -2.76754400 | -2.90583700 | -0.51213400 |
| C | -4.43175500 | -5.69296300 | -6.06588100 |
| C | -3.60595700 | -4.40506300 | -6.03553900 |
| O | -4.09950700 | -3.31137000 | -6.30907300 |
| C | -5.34433500 | -5.69314400 | -4.82108700 |
| C | -4.50842000 | -5.49646800 | -3.54966500 |
| O | -4.68939500 | -4.43368500 | -2.88802800 |
| O | -3.65548100 | -6.38112600 | -3.28128700 |
| H | -5.03398700 | -5.69492000 | -6.98561600 |
| H | -5.87506200 | -6.65617100 | -4.75675100 |
| H | -6.08155500 | -4.88295900 | -4.89947700 |
| N | -2.32772900 | -4.55038600 | -5.60031900 |
| C | -1.39752900 | -3.45211500 | -5.39337900 |
| C | -0.93485800 | -3.26906700 | -3.94236400 |
| C | 0.15915200 | -2.18575200 | -3.85376700 |
| C | -0.50308800 | -4.58573000 | -3.30299900 |
| C | 1.55762000 | -2.58312300 | -4.33152000 |
| H | -2.05800800 | -5.47272300 | -5.27950800 |
| H | -1.90176000 | -2.54421100 | -5.75041500 |
| H | -1.80523900 | -2.90320300 | -3.38153500 |
| H | 0.23986800 | -1.84024500 | -2.81241000 |
| H | -0.17024300 | -1.30460000 | -4.43328300 |
| H | 0.31130000 | -5.07542500 | -3.86418100 |
| H | -1.34433400 | -5.28663700 | -3.23188700 |
| H | -0.15207800 | -4.40485200 | -2.27910200 |
| H | 2.23659300 | -1.71781500 | -4.27843600 |
| H | 1.56389200 | -2.95068800 | -5.37041700 |
| H | 1.99583400 | -3.36971500 | -3.69747900 |
| C | 5.33325800 | -5.61208800 | -2.82648000 |
| C | 5.05455100 | -4.49294000 | -1.82412100 |
| O | 5.88394800 | -3.62769800 | -1.56254700 |
| H | 4.57573300 | -5.63012400 | -3.62564800 |
| N | 3.83419300 | -4.50580700 | -1.20199900 |
| C | 3.49863000 | -3.45314600 | -0.23420300 |
| C | 4.20944400 | -3.65029100 | 1.11065900 |
| O | 4.58755300 | -2.69941800 | 1.78751800 |
| C | 1.96794000 | -3.57273600 | -0.09233000 |

| | | | |
|---|---|---|---|
| C | 1.69193900 | -5.05294700 | -0.37322900 |
| C | 2.70468300 | -5.40841200 | -1.47095900 |
| H | 3.79607700 | -2.46510000 | -0.60495300 |
| H | 1.49832100 | -2.93887500 | -0.85841300 |
| H | 1.60915800 | -3.23128900 | 0.88698800 |
| H | 0.65884000 | -5.23994400 | -0.68324300 |
| H | 1.86570400 | -5.65119600 | 0.53435200 |
| H | 2.28000700 | -5.21766600 | -2.47067700 |
| H | 3.01522400 | -6.46507400 | -1.44225200 |
| N | 4.33509200 | -4.94020100 | 1.51081500 |
| C | 4.83995700 | -5.34190700 | 2.80540400 |
| C | 3.75546100 | -5.84023900 | 3.76892500 |
| C | 2.72703200 | -4.78851800 | 4.01984600 |
| C | 2.82825600 | -3.49414800 | 4.49714300 |
| N | 1.41026200 | -4.92027600 | 3.63513000 |
| C | 0.78468500 | -3.73811400 | 3.89792900 |
| N | 1.61770600 | -2.85481100 | 4.41402800 |
| H | 4.05845200 | -5.65960000 | 0.85649400 |
| H | 5.33480600 | -4.46037800 | 3.23493500 |
| H | 4.25133100 | -6.15351500 | 4.70404800 |
| H | 3.27491300 | -6.74455000 | 3.35759300 |
| H | 0.92995500 | -5.75030300 | 3.27937500 |
| H | 3.71883900 | -2.98911400 | 4.86833300 |
| H | -0.26080600 | -3.57450300 | 3.65137300 |
| C | 7.51411600 | 4.71082400 | 3.80064300 |
| C | 6.19820800 | 5.22375500 | 3.21905200 |
| O | 5.13354400 | 5.04165700 | 3.78708000 |
| H | 8.39555600 | 5.22301100 | 3.38893600 |
| N | 6.28574300 | 5.88547100 | 2.01902900 |
| C | 5.07513400 | 6.06509100 | 1.25523600 |
| C | 4.62981000 | 4.71299700 | 0.63136200 |
| O | 5.32154900 | 3.69888900 | 0.73943200 |
| H | 7.14759400 | 5.77579300 | 1.47085800 |
| H | 4.26990700 | 6.43392100 | 1.90686000 |
| N | 3.45054300 | 4.69731900 | -0.02889300 |
| C | 3.02320400 | 3.52755100 | -0.76616400 |
| C | 3.56013300 | 3.61005900 | -2.21929500 |
| O | 4.06305000 | 4.62807100 | -2.65893900 |
| C | 1.52067900 | 3.26488300 | -0.56289400 |
| C | 1.23395800 | 2.95791600 | 0.90354800 |
| O | 0.78630100 | 4.40629100 | -0.99699100 |
| H | 2.94182000 | 5.55608500 | -0.20723700 |
| H | 3.53775800 | 2.67427600 | -0.31095800 |
| H | 1.25567700 | 2.38116400 | -1.17134200 |
| H | -0.09973100 | 4.31462400 | -0.61267300 |
| H | 1.52357800 | 3.80964400 | 1.53671000 |
| H | 1.77261600 | 2.06060300 | 1.23404300 |
| H | 0.16201400 | 2.76922600 | 1.05559700 |
| N | 3.55385800 | 2.43848300 | -2.93058100 |
| C | 4.14507200 | 2.39136100 | -4.27187900 |
| H | 3.00136700 | 1.65372100 | -2.58958100 |
| H | 5.21639400 | 2.63827900 | -4.21540200 |
| C | -3.50178200 | -4.41138300 | 3.88394200 |
| O | -2.62514000 | -5.53534000 | 4.03645600 |
| C | -2.77491700 | -3.11965600 | 4.14824900 |
| O | -1.95278200 | -2.77046700 | 3.04497200 |
| C | -3.64322200 | -1.89813000 | 4.47536700 |
| O | -3.59357600 | -1.75675400 | 5.86871700 |
| C | -2.97563000 | -0.76689600 | 3.61755000 |
| O | -2.89788200 | 0.47929900 | 4.24621400 |
| C | -1.59772700 | -1.37678800 | 3.24180400 |
| N | 1.41121900 | -0.33237600 | 1.18320100 |
| C | 0.94217100 | -0.32021800 | -0.09442300 |
| O | 1.64772800 | -0.05290300 | -1.08447200 |
| N | -0.40906000 | -0.64813100 | -0.27798200 |
| C | -1.30192800 | -1.00866900 | 0.71931000 |
| O | -2.45514200 | -1.38608900 | 0.39015000 |
| C | -0.79767100 | -0.91947000 | 2.05358400 |
| C | 0.56793800 | -0.62089200 | 2.17536300 |
| H | -2.61429900 | 1.56710900 | 2.98693500 |

| | | | |
|---|---|---|---|
| H | -0.80539500 | -0.64415800 | -1.22442000 |
| H | 1.00865100 | -0.65865500 | 3.17882300 |
| H | -0.95451900 | -1.29677700 | 4.13768200 |
| H | -3.67980700 | 0.55463000 | 4.91402100 |
| H | -4.68645600 | -2.03854800 | 4.14278300 |
| H | -2.13864900 | -3.25090100 | 5.04650000 |
| H | -4.35049400 | -4.50029000 | 4.58232700 |
| H | -3.54588700 | -0.69950500 | 2.67867800 |
| H | -3.88467300 | -4.44291300 | 2.85401400 |
| P | -4.81202300 | -0.90274200 | 6.67053900 |
| O | -4.91887300 | 0.39118200 | 5.84907700 |
| O | -4.55139600 | -1.00379000 | 8.12441900 |
| O | -6.13221500 | -1.80667800 | 6.21286600 |
| C | 7.76012400 | -0.68817900 | 3.47188200 |
| C | 8.16579100 | -0.88797800 | 2.00727300 |
| C | 7.02362800 | -1.00422700 | 0.99508600 |
| C | 4.47620700 | 0.44457000 | 1.03457400 |
| S | 6.24197400 | 0.62540300 | 0.55494100 |
| C | 6.13684500 | 0.45145500 | -1.27691500 |
| C | 7.40057300 | 0.81035300 | -2.05958400 |
| O | 7.37320500 | 2.18052700 | -2.44956600 |
| C | 8.77168700 | 0.64495900 | -1.39247000 |
| O | 9.17325700 | -0.70665500 | -1.40999500 |
| C | 9.64210300 | 1.53940600 | -2.29389600 |
| O | 9.92725900 | 0.87033000 | -3.49224000 |
| C | 8.68321900 | 2.70344000 | -2.59520300 |
| C | 9.85049100 | 4.75625200 | -1.69426400 |
| C | 9.61694100 | 5.58150100 | -0.61762500 |
| N | 8.49132900 | 5.16999400 | 0.05522900 |
| C | 8.03993600 | 4.11029100 | -0.58888200 |
| N | 8.83445200 | 3.81656200 | -1.66592400 |
| H | 7.23291300 | 0.26953500 | 3.61770300 |
| H | 8.72612500 | -1.83562600 | 1.91750400 |
| H | 8.86940100 | -0.09388500 | 1.69741300 |
| H | 7.38546200 | -1.40516200 | 0.04018200 |
| H | 6.20637500 | -1.65022100 | 1.34791000 |
| H | 4.36339500 | -0.46577300 | 1.63582400 |
| H | 4.22945100 | 1.35998100 | 1.58286400 |
| H | 3.84876200 | 0.34615200 | 0.13701800 |
| H | 7.12937800 | 3.56966600 | -0.33941400 |
| H | 8.84069000 | 3.06741600 | -3.62217400 |
| H | 10.03433600 | -0.06933400 | -3.27126500 |
| H | 8.74617500 | 1.05878400 | -0.36954300 |
| H | 7.39677800 | 0.17256300 | -2.96176000 |
| H | 5.84279000 | -0.59218900 | -1.46131900 |
| H | 9.85426000 | -0.84771000 | -0.73868400 |
| H | 10.55652400 | 1.89783200 | -1.78576800 |
| H | 5.32210800 | 1.12016100 | -1.59385500 |
| C | -6.16557300 | 9.21531200 | 1.21484500 |
| C | -4.88600900 | 8.63157500 | 0.57638800 |
| O | -4.67204000 | 8.71255700 | -0.62761100 |
| H | -5.96173200 | 9.70159100 | 2.18041000 |
| N | -6.29404800 | 6.59231100 | -3.48489800 |
| C | -5.00429600 | 7.20246000 | -3.87540000 |
| H | -7.11671000 | 6.87139300 | -4.00639200 |
| H | -4.66602400 | 7.89103800 | -3.08428000 |
| O | -6.26909000 | 5.72363500 | 1.07250500 |
| H | -5.38949700 | 5.60620700 | 1.49530800 |
| H | -6.04862200 | 5.66891200 | 0.12325100 |
| H | -6.59887000 | 9.93252100 | 0.50816100 |
| H | -6.86849800 | 8.38254500 | 1.37920100 |
| H | -2.24088700 | 8.29396500 | 0.32447800 |
| H | -4.24642900 | 6.41956000 | -4.02333500 |
| H | -5.15426500 | 7.74931800 | -4.81377200 |
| H | -8.10266200 | -2.05097800 | -0.20721400 |
| H | -7.40127800 | -2.36584000 | -1.81704300 |
| H | -3.78470600 | -6.58201400 | -6.05442500 |
| H | -1.51269700 | -9.09470000 | -2.46334200 |
| H | -0.51935100 | -3.61206900 | -6.04632500 |
| H | 5.31212700 | -6.59518600 | -2.32966600 |

| | | | |
|---|---|---|---|
| H | 6.32534800 | -5.44390700 | -3.26077800 |
| H | 4.02404300 | 1.37687000 | -4.67569500 |
| H | 3.66255700 | 3.11284200 | -4.95006700 |
| H | 10.61884300 | 4.75478700 | -2.46336000 |
| H | 10.19343400 | 6.44922400 | -0.30107900 |
| H | 7.48280100 | 4.81333900 | 4.89331400 |
| H | 7.59239900 | 3.63827700 | 3.55890400 |
| H | 5.24098500 | 6.78874200 | 0.44203300 |
| H | -6.67461200 | -1.22512800 | 5.66294200 |
| H | 5.61132300 | -6.12004500 | 2.66847700 |
| H | 8.64483600 | -0.68357300 | 4.12528500 |
| H | 7.08513400 | -1.49290900 | 3.80235100 |
| H | -2.32110800 | -5.53398400 | 4.95603000 |
| H | -8.63751300 | 6.06931800 | -2.38309700 |

#### Model5' − TS2

| | | | |
|---|---|---|---|
| O | -4.14943800 | -1.92861300 | -3.76056800 |
| H | -4.14855900 | -2.10734400 | -4.72243500 |
| H | -4.44186100 | -2.80028700 | -3.39788700 |
| O | -2.26168200 | 1.83615600 | 2.13224100 |
| H | -2.27067700 | 3.17123100 | 1.38275200 |
| H | -1.47750400 | 1.28665000 | 1.97933600 |
| O | -2.22014600 | -0.46504100 | -2.66221800 |
| H | -2.95824300 | -1.04528400 | -3.02286000 |
| H | -1.72740500 | -0.20729600 | -3.45330000 |
| O | -0.53853100 | -6.66809200 | 2.67789300 |
| H | -0.68938200 | -7.62232800 | 2.68198800 |
| H | -1.31353400 | -6.27459600 | 3.15934900 |
| O | -6.92647700 | 3.17074100 | 1.85117300 |
| H | -6.84861800 | 4.13572600 | 1.65432500 |
| H | -6.58785400 | 3.07693700 | 2.75287900 |
| O | -3.14219200 | 4.47203500 | -2.63505600 |
| H | -2.40269100 | 5.04128300 | -2.38539600 |
| H | -3.94583300 | 4.93762300 | -2.31649200 |
| C | -7.91870400 | 5.25289300 | -2.14759800 |
| C | -6.49473300 | 5.76889800 | -2.40474600 |
| O | -5.54253900 | 5.38917600 | -1.68903400 |
| C | -8.24874600 | 4.03808200 | -3.03842400 |
| C | -7.54530000 | 2.73255300 | -2.63409300 |
| C | -8.11226200 | 2.18407900 | -1.32265100 |
| C | -7.64450900 | 1.68839500 | -3.75022400 |
| H | -7.98490700 | 4.99757400 | -1.08210200 |
| H | -7.99762200 | 4.28863800 | -4.08388600 |
| H | -9.34038800 | 3.87757200 | -3.01637600 |
| H | -6.47486400 | 2.95284500 | -2.48169600 |
| H | -7.59213400 | 1.26141100 | -1.03380400 |
| H | -9.18214900 | 1.94026300 | -1.43786300 |
| H | -8.00907700 | 2.87800200 | -0.47712300 |
| H | -8.69614500 | 1.41800700 | -3.94680800 |
| H | -7.21194600 | 2.06177800 | -4.69268800 |
| H | -7.10740600 | 0.76613200 | -3.47827800 |
| N | -3.97493200 | 8.01642700 | 1.44924300 |
| C | -2.67351700 | 7.51031700 | 0.99372700 |
| C | -2.70403200 | 6.20320700 | 0.19486500 |
| C | -2.93462500 | 4.98041900 | 1.05275800 |
| O | -3.80052200 | 4.93090300 | 1.91474700 |
| O | -2.11351900 | 3.99045300 | 0.78098500 |
| H | -4.22106100 | 7.87888900 | 2.42151600 |
| H | -2.03069900 | 7.40867100 | 1.87949500 |
| H | -1.74765100 | 6.09384800 | -0.33773200 |
| H | -3.50829500 | 6.27569500 | -0.55229300 |
| C | -7.25293500 | -1.80261700 | -0.87353600 |
| C | -5.88832200 | -2.09675400 | -0.25501500 |
| C | -5.62132800 | -1.40348900 | 1.08712600 |
| C | -5.47939500 | 0.12247500 | 1.02651600 |
| N | -4.44095400 | 0.48790300 | 0.07419100 |
| C | -4.28314200 | 1.68913600 | -0.49036800 |
| N | -5.07001800 | 2.71367900 | -0.15285200 |
| N | -3.33290100 | 1.85092400 | -1.41815400 |
| H | -7.37187100 | -0.73518900 | -1.11282000 |
| H | -5.10341300 | -1.81761400 | -0.97618600 |
| H | -5.78092000 | -3.18464800 | -0.11328900 |
| H | -4.69385400 | -1.81403600 | 1.51449100 |
| H | -6.42833800 | -1.63604500 | 1.80285600 |
| H | -5.23556400 | 0.50750000 | 2.03407900 |
| H | -6.42390600 | 0.60051800 | 0.72792300 |
| H | -3.65119100 | -0.16499400 | 0.01232500 |
| H | -5.69657700 | 2.71295200 | 0.66896400 |
| H | -5.06305900 | 3.56084200 | -0.71149000 |
| H | -2.92463200 | 1.02304300 | -1.86905700 |
| H | -3.18561200 | 2.77503200 | -1.83628100 |
| C | -0.69176000 | -8.37807200 | -2.22078000 |
| C | -0.81258300 | -8.10053300 | -0.71792400 |
| C | -2.10309300 | -7.38279700 | -0.29121500 |
| N | -2.33461900 | -6.09176000 | -0.91502000 |
| C | -2.25560100 | -4.88944000 | -0.32345300 |
| N | -1.57189100 | -4.70768900 | 0.82317100 |
| N | -2.83713000 | -3.84871900 | -0.91225100 |
| H | -0.66367700 | -7.44075300 | -2.79520000 |
| H | 0.22465500 | -8.94600900 | -2.44493800 |
| H | 0.05505300 | -7.51679900 | -0.36974500 |
| H | -0.77749400 | -9.05619300 | -0.16408800 |
| H | -2.10958100 | -7.26778200 | 0.80226800 |
| H | -2.96518300 | -8.02157500 | -0.54908900 |
| H | -2.83157000 | -6.13345000 | -1.84236400 |
| H | -1.10097300 | -5.48663700 | 1.27953400 |
| H | -1.72123500 | -3.86553200 | 1.38585200 |
| H | -3.52818600 | -4.03510500 | -1.67251400 |
| H | -2.72805100 | -2.91299000 | -0.50650400 |
| C | -4.42550000 | -5.67752500 | -6.07776900 |
| C | -3.60731800 | -4.38443500 | -6.03933800 |
| O | -4.11654700 | -3.28733400 | -6.26568000 |
| C | -5.31272500 | -5.70159700 | -4.81359400 |
| C | -4.45847100 | -5.50670400 | -3.55360500 |
| O | -4.62824700 | -4.44247400 | -2.88933100 |
| O | -3.60500900 | -6.39228800 | -3.29281100 |
| H | -5.04752700 | -5.66875500 | -6.98411200 |
| H | -5.83054500 | -6.67148900 | -4.74916900 |
| H | -6.06047600 | -4.89941100 | -4.87071700 |
| N | -2.31585600 | -4.53240500 | -5.64590800 |
| C | -1.39274400 | -3.43414300 | -5.40783400 |
| C | -0.92590200 | -3.30044500 | -3.95349400 |
| C | 0.16289200 | -2.21637700 | -3.82473300 |
| C | -0.48528800 | -4.63926000 | -3.37000000 |
| C | 1.56743600 | -2.59190900 | -4.30135300 |
| H | -2.02737400 | -5.46389500 | -5.37209200 |
| H | -1.90538500 | -2.51838600 | -5.73123700 |
| H | -1.79703500 | -2.96142600 | -3.37688500 |
| H | 0.23070500 | -1.90311900 | -2.77248800 |
| H | -0.16405200 | -1.31859500 | -4.37969100 |
| H | 0.33056000 | -5.09895700 | -3.95374000 |
| H | -1.32276500 | -5.34688100 | -3.32748400 |
| H | -0.13216600 | -4.50198600 | -2.34083600 |
| H | 2.24074100 | -1.72518100 | -4.21416700 |
| H | 1.58436900 | -2.92593000 | -5.35133000 |
| H | 2.00375000 | -3.39667400 | -3.68915400 |
| C | 5.35015900 | -5.59175400 | -2.87092400 |
| C | 5.01340500 | -4.51756800 | -1.84153000 |
| O | 5.79842400 | -3.62598600 | -1.54446300 |
| H | 4.61031800 | -5.60667700 | -3.68677800 |
| N | 3.78438900 | -4.59953800 | -1.23546400 |
| C | 3.38377700 | -3.56379600 | -0.27745300 |
| C | 4.05705500 | -3.72589200 | 1.08924900 |
| O | 4.29941900 | -2.75553100 | 1.79736600 |
| C | 1.85558200 | -3.73900500 | -0.17312900 |
| C | 1.64928600 | -5.22939900 | -0.46118800 |
| C | 2.69554200 | -5.53771900 | -1.54106400 |
| H | 3.66265800 | -2.56811000 | -0.64315900 |
| H | 1.37257800 | -3.12425600 | -0.94730200 |
| H | 1.46052800 | -3.42204100 | 0.80098500 |

| | | | |
|---|---|---|---|
| H | 0.63200300 | -5.46432700 | -0.78946700 |
| H | 1.84162200 | -5.81747400 | 0.44927900 |
| H | 2.28523800 | -5.35155000 | -2.54794700 |
| H | 3.04151100 | -6.58325000 | -1.51504900 |
| N | 4.29930700 | -5.00149000 | 1.47922200 |
| C | 4.87545400 | -5.34159000 | 2.76361000 |
| C | 3.86192100 | -5.82495400 | 3.81111500 |
| C | 2.81715300 | -4.79313800 | 4.07325300 |
| C | 2.89487300 | -3.50689700 | 4.57524300 |
| N | 1.51990800 | -4.91807300 | 3.62429800 |
| C | 0.88176800 | -3.74168800 | 3.87715700 |
| N | 1.68962400 | -2.86467400 | 4.44338900 |
| H | 4.15876600 | -5.73203500 | 0.79408400 |
| H | 5.37221400 | -4.43570600 | 3.13769600 |
| H | 4.42027000 | -6.07030400 | 4.73034200 |
| H | 3.38973900 | -6.76354400 | 3.47403300 |
| H | 1.05046800 | -5.74636200 | 3.25242900 |
| H | 3.76866800 | -3.00790900 | 4.99183900 |
| H | -0.15627600 | -3.58113200 | 3.60068600 |
| C | 7.53639300 | 4.71205400 | 3.78409000 |
| C | 6.21998900 | 5.22847200 | 3.20872000 |
| O | 5.15871500 | 5.06057900 | 3.78879200 |
| H | 8.41785600 | 5.21499900 | 3.36074000 |
| N | 6.30131000 | 5.87754800 | 2.00281600 |
| C | 5.08676300 | 6.07093300 | 1.25154100 |
| C | 4.62755700 | 4.73567900 | 0.61099400 |
| O | 5.30498400 | 3.71611100 | 0.70378300 |
| H | 7.15815500 | 5.77897600 | 1.44377200 |
| H | 4.28897700 | 6.43506800 | 1.91514200 |
| N | 3.43824300 | 4.73976000 | -0.04400300 |
| C | 3.01175500 | 3.57906800 | -0.79376100 |
| C | 3.55693100 | 3.66019300 | -2.24484100 |
| O | 4.08077600 | 4.67158900 | -2.67477000 |
| C | 1.50556800 | 3.32477300 | -0.62405700 |
| C | 1.16445600 | 3.04784300 | 0.83665800 |
| O | 0.78555000 | 4.45826200 | -1.10064500 |
| H | 2.96410000 | 5.61259700 | -0.24655000 |
| H | 3.52127700 | 2.72292400 | -0.33373000 |
| H | 1.25363400 | 2.43246400 | -1.22584700 |
| H | -0.11904700 | 4.35728700 | -0.76833900 |
| H | 1.42497600 | 3.91468500 | 1.46101100 |
| H | 1.70082000 | 2.16836800 | 1.21138500 |
| H | 0.08886300 | 2.85563900 | 0.94836700 |
| N | 3.51653400 | 2.49687100 | -2.96385900 |
| C | 4.14405100 | 2.41463800 | -4.28498600 |
| H | 2.98514700 | 1.70642700 | -2.60722100 |
| H | 5.23117500 | 2.56961900 | -4.19718900 |
| C | -3.42243900 | -4.39719300 | 3.87644600 |
| O | -2.58525800 | -5.55148000 | 4.01883000 |
| C | -2.67039600 | -3.12317500 | 4.16225400 |
| O | -1.83906300 | -2.76877800 | 3.06370600 |
| C | -3.53454600 | -1.89980900 | 4.49157800 |
| O | -3.52939400 | -1.78149700 | 5.88723700 |
| C | -2.84192200 | -0.76118100 | 3.65927600 |
| O | -2.75989400 | 0.47049500 | 4.31526100 |
| C | -1.46421000 | -1.38307700 | 3.27159200 |
| N | 1.47288800 | -0.32554100 | 1.07002200 |
| C | 0.93643000 | -0.29626400 | -0.18898000 |
| O | 1.58104800 | -0.00786700 | -1.20700900 |
| N | -0.41325900 | -0.63290700 | -0.29181600 |
| C | -1.24953000 | -1.00619700 | 0.74699800 |
| O | -2.40883200 | -1.38895000 | 0.47578000 |
| C | -0.67661700 | -0.92876600 | 2.06295200 |
| C | 0.68972200 | -0.63541000 | 2.12087100 |
| H | -2.54742200 | 1.54652300 | 3.05012900 |
| H | -0.85758400 | -0.62827900 | -1.22013800 |
| H | 1.19350500 | -0.69180300 | 3.09287500 |
| H | -0.80836900 | -1.31587800 | 4.15897400 |
| H | -3.55142400 | 0.53876700 | 4.97342400 |
| H | -4.56821600 | -2.02683700 | 4.12464500 |
| H | -2.03937200 | -3.27170900 | 5.06078600 |
| H | -4.27889500 | -4.46934100 | 4.56714100 |
| H | -3.40689300 | -0.66887200 | 2.71827800 |
| H | -3.80061600 | -4.40272400 | 2.84421800 |
| P | -4.77079300 | -0.93158500 | 6.67610500 |
| O | -4.82585200 | 0.38014700 | 5.87804400 |
| O | -4.55777100 | -1.07382400 | 8.13360600 |
| O | -6.08346900 | -1.80814600 | 6.15192200 |
| C | 7.79003600 | -0.68543300 | 3.43582800 |
| C | 8.05719800 | -0.89768100 | 1.94027400 |
| C | 6.81569900 | -1.00543900 | 1.04398300 |
| C | 3.71926200 | 0.16034800 | 0.93371500 |
| S | 6.03402600 | 0.62267800 | 0.60931700 |
| C | 6.06820900 | 0.54885800 | -1.23605600 |
| C | 7.37328200 | 0.87153700 | -1.99159100 |
| O | 7.36872900 | 2.21120600 | -2.49717600 |
| C | 8.71329500 | 0.75888100 | -1.26283800 |
| O | 9.12354100 | -0.59068800 | -1.18820000 |
| C | 9.62797000 | 1.59364900 | -2.18235800 |
| O | 9.98467700 | 0.83521700 | -3.30698000 |
| C | 8.68721000 | 2.72849500 | -2.62235300 |
| C | 9.75804100 | 4.88633600 | -1.84029900 |
| C | 9.51817000 | 5.72477100 | -0.77483500 |
| N | 8.43613600 | 5.27432900 | -0.05625000 |
| C | 8.01870900 | 4.17888300 | -0.66334700 |
| N | 8.79018200 | 3.90128800 | -1.76016700 |
| H | 7.26413200 | 0.26648900 | 3.61706300 |
| H | 8.60939400 | -1.84604600 | 1.81007700 |
| H | 8.72507600 | -0.09964300 | 1.57038300 |
| H | 7.06154600 | -1.48847000 | 0.08992800 |
| H | 6.03662300 | -1.61104400 | 1.52504300 |
| H | 3.88975400 | -0.83785400 | 1.33998300 |
| H | 3.58856000 | 1.01030700 | 1.59993000 |
| H | 3.46032000 | 0.27528500 | -0.11726800 |
| H | 7.14657100 | 3.59695200 | -0.37234000 |
| H | 8.89434500 | 3.02453600 | -3.66186200 |
| H | 10.06787800 | -0.08320600 | -3.00035300 |
| H | 8.63595400 | 1.22361500 | -0.26638100 |
| H | 7.41353500 | 0.17388400 | -2.84707600 |
| H | 5.73832700 | -0.46547300 | -1.50094000 |
| H | 9.67280700 | -0.71462800 | -0.40365700 |
| H | 10.51242100 | 1.99627000 | -1.65431800 |
| H | 5.29865900 | 1.25925900 | -1.57186600 |
| C | -6.15923100 | 9.20281300 | 1.25963800 |
| C | -4.88615900 | 8.61383300 | 0.61452800 |
| O | -4.69106300 | 8.67034300 | -0.59423400 |
| H | -5.95378100 | 9.67425600 | 2.23209000 |
| N | -6.29993000 | 6.59720800 | -3.44800200 |
| C | -5.01179900 | 7.20921400 | -3.84135100 |
| H | -7.12538300 | 6.88113400 | -3.96248600 |
| H | -4.66122700 | 7.87824300 | -3.03904900 |
| O | -6.28076900 | 5.72819900 | 1.09061000 |
| H | -5.40532800 | 5.62834800 | 1.52571800 |
| H | -6.04735300 | 5.67119500 | 0.14466300 |
| H | -6.58488100 | 9.93344700 | 0.56193100 |
| H | -6.87063400 | 8.37466600 | 1.40966800 |
| H | -2.23200000 | 8.28237700 | 0.34520100 |
| H | -4.26010800 | 6.42631400 | -4.01799900 |
| H | -5.17116700 | 7.77859500 | -4.76459900 |
| H | -8.07466500 | -2.08320300 | -0.19341300 |
| H | -7.37954400 | -2.36355700 | -1.81232000 |
| H | -3.77301000 | -6.56259200 | -6.09483600 |
| H | -1.55436800 | -8.95547000 | -2.58962700 |
| H | -0.51673200 | -3.56516900 | -6.06977100 |
| H | 5.35128800 | -6.59212800 | -2.40933200 |
| H | 6.34457700 | -5.37632800 | -3.27807100 |
| H | 3.94981600 | 1.41938800 | -4.70736600 |
| H | 3.74129700 | 3.18279500 | -4.96321600 |
| H | 10.50290100 | 4.90180000 | -2.63194800 |
| H | 10.05983300 | 6.62753700 | -0.49702200 |

| | | | |
|---|---|---|---|
| H | 7.51525900 | 4.82491000 | 4.87602700 |
| H | 7.60490200 | 3.63686500 | 3.55131500 |
| H | 5.24984800 | 6.80672100 | 0.44826200 |
| H | -6.59756800 | -1.21040700 | 5.59218600 |
| H | 5.65440600 | -6.10809300 | 2.61032000 |
| H | 8.73144600 | -0.66204200 | 4.00509300 |
| H | 7.16185400 | -1.49511900 | 3.83987200 |
| H | -2.32105600 | -5.59690700 | 4.94943900 |
| H | -8.63720100 | 6.06562100 | -2.34140200 |

#### Model5' – Final state

| | | | |
|---|---|---|---|
| O | -4.14143600 | -1.93394200 | -3.74136400 |
| H | -4.09596200 | -2.11666500 | -4.70223000 |
| H | -4.43997600 | -2.80982400 | -3.39280900 |
| O | -2.17719000 | 1.83577700 | 2.22725700 |
| H | -2.39651600 | 3.16446300 | 1.45468700 |
| H | -1.22965100 | 1.76972200 | 2.41082700 |
| O | -2.27626900 | -0.41435000 | -2.63323800 |
| H | -2.99285000 | -1.02738600 | -2.99089600 |
| H | -1.79686900 | -0.13936400 | -3.42722400 |
| O | -0.63541600 | -6.86002600 | 2.66412900 |
| H | -0.80582500 | -7.81084200 | 2.65611300 |
| H | -1.40955100 | -6.45084100 | 3.12680100 |
| O | -7.08008500 | 3.25208700 | 1.80286800 |
| H | -6.96861700 | 4.21286700 | 1.59906000 |
| H | -6.80452100 | 3.16495200 | 2.72632400 |
| O | -3.11254600 | 4.49453900 | -2.64366000 |
| H | -2.39767700 | 5.11826500 | -2.46276100 |
| H | -3.92783000 | 4.94995300 | -2.33819900 |
| C | -7.88222100 | 5.23427400 | -2.21777500 |
| C | -6.45996200 | 5.76063900 | -2.45548100 |
| O | -5.51716100 | 5.39042800 | -1.72155200 |
| C | -8.19555500 | 4.02734700 | -3.12600600 |
| C | -7.49826100 | 2.71890400 | -2.71981100 |
| C | -8.10486400 | 2.15031800 | -1.43467700 |
| C | -7.55711200 | 1.68957200 | -3.85249800 |
| H | -7.95840700 | 4.96711800 | -1.15596400 |
| H | -7.92530600 | 4.28745900 | -4.16422700 |
| H | -9.28710300 | 3.86562200 | -3.12599700 |
| H | -6.43376500 | 2.94170600 | -2.53215100 |
| H | -7.58815600 | 1.22760000 | -1.13924700 |
| H | -9.16778100 | 1.89904800 | -1.59032600 |
| H | -8.03819600 | 2.83623200 | -0.57875900 |
| H | -8.60055100 | 1.41731600 | -4.08623200 |
| H | -7.09660300 | 2.07788200 | -4.77541700 |
| H | -7.02419600 | 0.76597200 | -3.57641300 |
| N | -3.97335200 | 8.03562200 | 1.42289100 |
| C | -2.67660300 | 7.50785100 | 0.97726200 |
| C | -2.72623500 | 6.18216100 | 0.21283600 |
| C | -3.01425900 | 4.98815600 | 1.09302100 |
| O | -3.88098600 | 4.98865400 | 1.95630200 |
| O | -2.24036200 | 3.96000900 | 0.83193600 |
| H | -4.21179300 | 7.93289900 | 2.40130500 |
| H | -2.03417600 | 7.42127400 | 1.86479400 |
| H | -1.76127900 | 6.02307600 | -0.29083600 |
| H | -3.50950400 | 6.25751100 | -0.55585100 |
| C | -7.20482000 | -1.81973000 | -0.94128000 |
| C | -5.89155600 | -2.08138300 | -0.21358900 |
| C | -5.73326500 | -1.34490600 | 1.12097400 |
| C | -5.61024900 | 0.18270200 | 1.03592300 |
| N | -4.55334700 | 0.55410900 | 0.10445100 |
| C | -4.38906900 | 1.75203000 | -0.46946100 |
| N | -5.15489300 | 2.78699600 | -0.12174700 |
| N | -3.45181900 | 1.89629600 | -1.41232700 |
| H | -7.30915700 | -0.76221500 | -1.22726300 |
| H | -5.05587000 | -1.82071700 | -0.88217900 |
| H | -5.78832100 | -3.16280800 | -0.02580400 |
| H | -4.83750100 | -1.73512200 | 1.62499400 |
| H | -6.58684000 | -1.57072700 | 1.78256200 |
| H | -5.40165500 | 0.58339300 | 2.04547100 |
| H | -6.55387400 | 0.64139700 | 0.70582000 |
| H | -3.78439100 | -0.11504900 | 0.01319800 |
| H | -5.80147600 | 2.78126100 | 0.68464400 |
| H | -5.15638300 | 3.62795200 | -0.69229300 |
| H | -3.03978500 | 1.06554500 | -1.85213200 |
| H | -3.24966100 | 2.82222100 | -1.80281700 |
| C | -0.60776100 | -8.37161100 | -2.22666600 |
| C | -0.74235600 | -8.06279700 | -0.73291800 |
| C | -2.09512100 | -7.45954300 | -0.32612900 |
| N | -2.40396300 | -6.17155500 | -0.92000100 |
| C | -2.32402400 | -4.97984100 | -0.31515800 |
| N | -1.62456700 | -4.80984000 | 0.82578400 |
| N | -2.92475300 | -3.93493600 | -0.87928400 |
| H | -0.72365000 | -7.45989300 | -2.83103700 |
| H | 0.37553700 | -8.81193100 | -2.45442700 |
| H | 0.06367900 | -7.38434500 | -0.40872700 |
| H | -0.60944500 | -8.99279600 | -0.15096800 |
| H | -2.14106400 | -7.37668600 | 0.76903600 |
| H | -2.90028000 | -8.15063500 | -0.62752200 |
| H | -2.89102200 | -6.20463600 | -1.85559800 |
| H | -1.15688300 | -5.59789300 | 1.26901600 |
| H | -1.78572500 | -3.98363900 | 1.40715000 |
| H | -3.60065500 | -4.10866700 | -1.65884400 |
| H | -2.83148800 | -3.00531800 | -0.46536100 |
| C | -4.31201400 | -5.68153800 | -6.11921400 |
| C | -3.50136300 | -4.38512100 | -6.05283200 |
| O | -4.02302200 | -3.28553500 | -6.23606800 |
| C | -5.25374000 | -5.69385400 | -4.89587300 |
| C | -4.45180600 | -5.51516800 | -3.60046000 |
| O | -4.64662400 | -4.46125100 | -2.92497000 |
| O | -3.61180300 | -6.40676700 | -3.31902100 |
| H | -4.89459400 | -5.68173800 | -7.05184500 |
| H | -5.78751400 | -6.65632500 | -4.85432500 |
| H | -5.98649200 | -4.88105400 | -4.98512000 |
| N | -2.20236700 | -4.53161900 | -5.68682400 |
| C | -1.29392600 | -3.42813200 | -5.41765300 |
| C | -0.92238700 | -3.26359900 | -3.93790400 |
| C | 0.14098800 | -2.16356700 | -3.75598200 |
| C | -0.50670600 | -4.58591000 | -3.29891800 |
| C | 1.56968100 | -2.50657800 | -4.18167200 |
| H | -1.89580600 | -5.46815200 | -5.45382300 |
| H | -1.78454000 | -2.51927000 | -5.79160700 |
| H | -1.83526800 | -2.92592300 | -3.42771500 |
| H | 0.15905600 | -1.86834100 | -2.69671100 |
| H | -0.18074600 | -1.26296200 | -4.30934100 |
| H | 0.34779700 | -5.04985900 | -3.82048100 |
| H | -1.33914800 | -5.30106500 | -3.29309400 |
| H | -0.21666500 | -4.42223600 | -2.25294000 |
| H | 2.22637200 | -1.63458500 | -4.03989900 |
| H | 1.63617100 | -2.80711800 | -5.23954200 |
| H | 1.99088600 | -3.32448000 | -3.57695400 |
| C | 5.43066100 | -5.56402800 | -2.81460100 |
| C | 5.12271100 | -4.47697700 | -1.79672400 |
| O | 5.92890900 | -3.60740000 | -1.50122300 |
| H | 4.68772300 | -5.57306600 | -3.62787500 |
| N | 3.88835700 | -4.52396600 | -1.19437700 |
| C | 3.52496500 | -3.47743800 | -0.23669100 |
| C | 4.19327500 | -3.66063900 | 1.13042200 |
| O | 4.43273700 | -2.70239400 | 1.85045400 |
| C | 1.99143700 | -3.61070200 | -0.12942500 |
| C | 1.74229600 | -5.09663100 | -0.41017900 |
| C | 2.77670500 | -5.43514100 | -1.49269100 |
| H | 3.83907600 | -2.49091000 | -0.59992900 |
| H | 1.52482700 | -2.98913800 | -0.90911800 |
| H | 1.60534900 | -3.28435400 | 0.84589000 |
| H | 0.71660500 | -5.30616900 | -0.73152000 |
| H | 1.92485400 | -5.68358500 | 0.50259600 |
| H | 2.36660500 | -5.24485100 | -2.49950600 |
| H | 3.09500700 | -6.48946100 | -1.46313400 |
| N | 4.41476400 | -4.94879200 | 1.50308000 |

| | | | |
|---|---|---|---|
| C | 4.89880300 | -5.31903300 | 2.81498700 |
| C | 3.80311000 | -5.83890700 | 3.75618600 |
| C | 2.71021700 | -4.83433800 | 3.92360600 |
| C | 2.73286400 | -3.49810400 | 4.27939500 |
| N | 1.40292600 | -5.08053200 | 3.56118300 |
| C | 0.70667200 | -3.92110100 | 3.70904900 |
| N | 1.48530800 | -2.94253200 | 4.13535300 |
| H | 4.28238500 | -5.66638800 | 0.80292900 |
| H | 5.35718400 | -4.41878600 | 3.24748500 |
| H | 4.27258800 | -6.09271100 | 4.72241800 |
| H | 3.38341200 | -6.77929300 | 3.35986300 |
| H | 0.97351300 | -5.95756800 | 3.26113300 |
| H | 3.59204600 | -2.90511500 | 4.58492200 |
| H | -0.34856900 | -3.83741300 | 3.47073100 |
| C | 7.51447900 | 4.74309600 | 3.86808100 |
| C | 6.20618600 | 5.30137600 | 3.30646700 |
| O | 5.15179900 | 5.20462800 | 3.91601800 |
| H | 8.40169800 | 5.25533900 | 3.46859100 |
| N | 6.29356100 | 5.89932300 | 2.07488100 |
| C | 5.08566800 | 6.09512200 | 1.31190600 |
| C | 4.61578900 | 4.75898400 | 0.69074800 |
| O | 5.25219300 | 3.72671800 | 0.83973500 |
| H | 7.14623500 | 5.74104500 | 1.51865500 |
| H | 4.28861700 | 6.47546600 | 1.96789000 |
| N | 3.44134900 | 4.78357400 | -0.00893400 |
| C | 3.04526600 | 3.62446700 | -0.77610900 |
| C | 3.61651000 | 3.71231000 | -2.21280400 |
| O | 4.09946100 | 4.74152400 | -2.64905100 |
| C | 1.53170700 | 3.37938900 | -0.71520000 |
| C | 1.04743400 | 3.14446000 | 0.71144900 |
| O | 0.86654800 | 4.49989400 | -1.29302500 |
| H | 3.06440100 | 5.67345800 | -0.31633900 |
| H | 3.53448500 | 2.76612500 | -0.29285000 |
| H | 1.32897700 | 2.47288200 | -1.31368600 |
| H | -0.07982300 | 4.35432000 | -1.15886900 |
| H | 1.18629800 | 4.05371500 | 1.31348400 |
| H | 1.60843700 | 2.33227700 | 1.18889200 |
| H | -0.02081000 | 2.88384800 | 0.70710100 |
| N | 3.57744100 | 2.54914800 | -2.92706200 |
| C | 4.21109700 | 2.43904700 | -4.23597000 |
| H | 3.20146800 | 1.71471100 | -2.48848000 |
| H | 5.30532700 | 2.32952600 | -4.14040300 |
| C | -3.50467400 | -4.47899700 | 3.86954300 |
| O | -2.57342200 | -5.55554700 | 3.99540800 |
| C | -2.80372000 | -3.17455100 | 4.12005000 |
| O | -1.99523800 | -2.82266500 | 3.00616100 |
| C | -3.66431500 | -1.95559700 | 4.45381800 |
| O | -3.60505100 | -1.82692200 | 5.84594000 |
| C | -2.98304800 | -0.81752300 | 3.61765600 |
| O | -2.88371200 | 0.41262000 | 4.26495400 |
| C | -1.61839100 | -1.45203900 | 3.24270400 |
| N | 1.28446800 | -0.31869900 | 1.09921800 |
| C | 0.77311900 | -0.25679400 | -0.19729500 |
| O | 1.44274100 | 0.03855100 | -1.17497000 |
| N | -0.56873200 | -0.58266400 | -0.29026500 |
| C | -1.40670700 | -0.99538200 | 0.72968700 |
| O | -2.55314300 | -1.37652400 | 0.44933300 |
| C | -0.83272300 | -0.97871100 | 2.05607100 |
| C | 0.49790300 | -0.68209500 | 2.15966500 |
| H | -2.58573700 | 1.44404800 | 3.06483800 |
| H | -1.01102100 | -0.55785700 | -1.22480100 |
| H | 1.01577000 | -0.78804400 | 3.11657700 |
| H | -0.96827000 | -1.40258400 | 4.13474400 |
| H | -3.69054900 | 0.49834200 | 4.92956800 |
| H | -4.70909600 | -2.08260900 | 4.12233200 |
| H | -2.15189300 | -3.29629500 | 5.00791300 |
| H | -4.33145000 | -4.59700600 | 4.58975100 |
| H | -3.54723700 | -0.72658600 | 2.67717000 |
| H | -3.91270900 | -4.52242000 | 2.84964900 |
| P | -4.80147200 | -0.94484900 | 6.63337300 |

| | | | |
|---|---|---|---|
| O | -4.88985200 | 0.34446100 | 5.79584400 |
| O | -4.54135100 | -1.02102300 | 8.08748500 |
| O | -6.13470200 | -1.82800000 | 6.18200500 |
| C | 7.79015600 | -0.65336400 | 3.51921900 |
| C | 8.31461900 | -0.99338800 | 2.12138500 |
| C | 7.24715800 | -1.08053400 | 1.02920200 |
| C | 2.71372200 | -0.03522100 | 1.31668600 |
| S | 6.46835400 | 0.54904500 | 0.61005100 |
| C | 6.35859100 | 0.35222200 | -1.21133700 |
| C | 7.57012700 | 0.78272500 | -2.04744500 |
| O | 7.46583700 | 2.16935900 | -2.39628900 |
| C | 8.95506800 | 0.66712400 | -1.41092900 |
| O | 9.45377100 | -0.65329500 | -1.52524400 |
| C | 9.77255000 | 1.66520100 | -2.24829200 |
| O | 10.14796600 | 1.06880100 | -3.46334300 |
| C | 8.73635000 | 2.76765800 | -2.52774100 |
| C | 9.74722800 | 4.90892100 | -1.64918300 |
| C | 9.48379000 | 5.70574600 | -0.55854200 |
| N | 8.42805800 | 5.18949400 | 0.15369200 |
| C | 8.04978600 | 4.09383600 | -0.48073700 |
| N | 8.82314100 | 3.88184100 | -1.58817700 |
| H | 7.32011100 | 0.34364700 | 3.53562400 |
| H | 8.82025000 | -1.97632600 | 2.15831700 |
| H | 9.08703300 | -0.25941100 | 1.82603500 |
| H | 7.68844100 | -1.47111800 | 0.10290100 |
| H | 6.44059500 | -1.76308400 | 1.33075600 |
| H | 3.26313300 | -0.95358900 | 1.56611800 |
| H | 2.82634500 | 0.69259300 | 2.13082200 |
| H | 3.12854100 | 0.38873200 | 0.39912600 |
| H | 7.21490200 | 3.45672100 | -0.19702200 |
| H | 8.87999200 | 3.16237700 | -3.54646100 |
| H | 10.27120000 | 0.12494900 | -3.26533700 |
| H | 8.88910900 | 0.99385900 | -0.36122600 |
| H | 7.57383800 | 0.18298100 | -2.97630300 |
| H | 6.12742800 | -0.70325400 | -1.41721400 |
| H | 10.01228300 | -0.84371600 | -0.76132600 |
| H | 10.64456000 | 2.07264400 | -1.70318400 |
| H | 5.49930300 | 0.95442600 | -1.54110100 |
| C | -6.17063500 | 9.18808600 | 1.20926800 |
| C | -4.89587800 | 8.59356800 | 0.57512700 |
| O | -4.71295600 | 8.60736600 | -0.63728600 |
| H | -5.97628000 | 9.65068200 | 2.18811200 |
| N | -6.25482700 | 6.58441000 | -3.49935900 |
| C | -4.96530000 | 7.20167600 | -3.88115500 |
| H | -7.07402800 | 6.85468400 | -4.03105200 |
| H | -4.60878900 | 7.84242300 | -3.05896400 |
| O | -6.33098500 | 5.77550400 | 1.04946500 |
| H | -5.46979100 | 5.67689700 | 1.51338600 |
| H | -6.06745200 | 5.73263900 | 0.11130100 |
| H | -6.58257900 | 9.92745800 | 0.51221600 |
| H | -6.88964100 | 8.36371600 | 1.34087700 |
| H | -2.22650100 | 8.25868800 | 0.30988500 |
| H | -4.21885300 | 6.42157300 | -4.09004200 |
| H | -5.12834000 | 7.80331900 | -4.78292400 |
| H | -8.07436300 | -2.08189400 | -0.31556200 |
| H | -7.25708000 | -2.41392300 | -1.86651600 |
| H | -3.65868700 | -6.56566600 | -6.10049200 |
| H | -1.38448900 | -9.07977000 | -2.55679600 |
| H | -0.38031200 | -3.57732700 | -6.02059200 |
| H | 5.41415500 | -6.56009100 | -2.34352500 |
| H | 6.42764200 | -5.37429000 | -3.22832600 |
| H | 3.80075100 | 1.56988000 | -4.76919000 |
| H | 4.01109300 | 3.35102900 | -4.81538200 |
| H | 10.48119200 | 4.97769000 | -2.44807800 |
| H | 9.99006400 | 6.62151400 | -0.25731400 |
| H | 7.48799400 | 4.80745700 | 4.96349900 |
| H | 7.57508500 | 3.68070100 | 3.58018000 |
| H | 5.25837900 | 6.82179400 | 0.50170800 |
| H | -6.69970600 | -1.22975800 | 5.67437700 |
| H | 5.69594900 | -6.07412400 | 2.70969900 |

| | | | | | | | |
|---|---|---|---|---|---|---|---|
| H | 8.60164500 | -0.65457300 | 4.26332600 | H | -2.23146800 | -5.53389700 | 4.90129400 |
| H | 7.02778300 | -1.38320400 | 3.83726400 | H | -8.60354700 | 6.04473000 | -2.41031000 |

### 4 Experimental procedures

#### 4.1 Purification and expression of Nep1

The EMG1 gene from *S. cerevisiae* and Nep1 gene from *A. flugidus*, which code for the Nep1 protein was cloned into a kanamycin resistance–plasmid containing an N–terminal 6×histidine tag followed by a Tobacco Etch Virus (TEV) protease cleavage site. The recombinant proteins were overexpressed in competent cells *Escherichia coli* BL21 (DE3) RIL. Expression cultures were grown in 37°C in LB–medium kanamycin–containing (100 mg/L). Protein expression was induced by 1 mM isopropyl 1–thio–*beta*–D–galactopyranoside (IPTG) at an OD600 = 0.8–1.0 and then cooled to 24°C for *Sc*Nep1 and 20°C for *Af*Nep1[1]. After overnight bacterial cultivations, cells with induced proteins were harvested by centrifugation (30 min, 4000 rpm, 4°C for each culture) and stored in –80°C. The cells were resuspended in lysis buffer (25 mM Tris/HCl, pH 8.0 200 mM NaCl, 20 mM imidazole, 1 mM DTT for *Sc*Nep1, and 25 mM Tris/HCl, pH 8.0 300 mM NaCl, 20 mM imidazole, 1 mM DTT for *Af*Nep1) and were lysed using a sonicator in an ice bath. The cell lysates were cleared by centrifugation (30 min, 20 000 rpm, 4°C for each culture).

After centrifugation, the supernatant of each protein was filtered through a 0,45 $\mu$M filter and loaded onto 5 ml HisTrap$^{TM}$ HP column (GE Healthcare). The resin was then washed twice, first with a lysis then with a buffer containing 25 mM Tris/HCl, pH 8.0 500 mM NaCl, 40 mM imidazole and eluted with lysis buffer containing 300 mM imidazole. The fractions containing Nep1 were dialyzed overnight with TEV proteases into 25 mM Tris pH 8.0, 300 mM NaCl, 1 mM EDTA and 1 mM DTT for *Sc*Nep1 and 25 mM Tris pH 8.0, 200 mM NaCl, 1 mM EDTA and 1 mM DTT for *Af*Nep1. After dialysis, it was checked that the tag was cleaved. In the case of *Sc*Nep1 tag was successfully cleaved. The *Sc*Nep1 protein was reapplied to the HisTrapTM HP column for the removal of TEV proteases and uncleaved protein. For *Af*Nep1, protease attempts to cleave the tag by TEV proteases have been unsuccessful. The proteins were concentrated to 5 ml volume and further purified by size exclusion chromatography using a Superdex 75$^{TM}$ column (GE Healthcare) equilibrated in 20 mM Tris pH 8.0, 300 mM NaCl, 1 mM EDTA and 1 mM TCEP for *Sc*Nep1 and 20 mM Tris pH 8.0, 200 mM NaCl, 1 mM EDTA and 1 mM TCEP for *Af*Nep1[1]. The proteins were concentrated again and stored at –20°C.

Mutagenesis was used to prepare the single mutants *Sc*Nep1–S233M, *Sc*Nep1_D101A, *Sc*Nep1_S233A. The template for the PCR reaction was used EMG1 gene encoding full–length *Sc*Nep1 protein. *Sc*Nep1_S233M, *Sc*Nep1_S233A and *Sc*Nep1_D101A were overexpressed and purified as previously described for *Sc*Nep1 [1]. To determine the biological activity rRNA with the following sequences was used: AAUUUGACΨCAACACGGGGA (Horizon Discovery).

#### 4.2 Sequences of expressed proteins

***Sc*Nep1**:
MGSSHHHHHHSSGLEVLFQGPHMENLYFQGMVEDSRVRDALKGGDQKALPASLVPQAPPVLTSK
DKITKRMIVVLAMASLETHKISSNGPGGDKYVLLNCDDHQGLLKKMGRDISEARPDITHQCLLTL
LASPINKAGKLQVYIQTSRGILIEVNPTVRIPRTFKRFSGLMVQLLHKLSIRSVNSEEKLLKVIKNPI
TDHLPTKCRKVTLSFDAPVIRVQDYIEKLDDDESICVFVGAMARGKDNFADEYVDEKVGLSNYPL
SASVACSKFCHGAEDAWNIL–

***Af*Nep1**:
MGSSHHYHHHSSGLEVLFQGPHMENLYFQGMGSAFVFLEASLELIPQKIRGHPAVRADAIRRGKR
PEKILLDDSKHHTAMKSLEFREKRGRPDIVHQCLLLLLDSPLRDFEVYVHTLNGEIIWVNRETRLP
RNYNRFVGLMEKLFEERRITAGDTTLIEFKDVGLRDIVRGRDVLLFREKGGRFEFSELLDGDVAV
CIGAFPHGDFFEETLRELGEFKEVSLGTESYTSLYVTSRVLCEYERVRAHKVG–

For *Af*Nep1 plasmid was made by Eurofins based on a commercial vector. The sequence cut from that plasmid was cloned with the Gibson method to mentioned pET28b vector giving the sequence above. To the *Sc*Nep1 wild type, point mutations were introduced using KLD (Kinase, Ligase and DpnI enzymes) reaction with the following starters:

**S233A**
Forward: GCG GCC AGC GTT GCG TGT TCG AAG
Reverse: CAG CGG ATA ATT GGA CAG GCC

**S233M**
Forward: ATG GCC AGC GTT GCG TGT TCG AAG
Reverse: CAG CGG ATA ATT GGA CAG GCC

**D101A**
Forward: GCC AGT CCG ATT AAC AAA GCC GGG
Reverse: AAG CAG GGT TAA CAG ACA TTG G
